## supplementary methods and figures for "MGPfact^XMBD^: A Model-Based Factorization Method for scRNA Data Unveils Bifurcating Transcriptional Modules Underlying Cell Fate Determination"

### 1 Methods

#### 1.1 Overview of MGPfact

The analytical pipeline of MGPfact consists two major steps (Fig. 1): first, we perform downsampling with preprocessed data based on the "minimum unbiased representative points" (MURPs) as described previously(Ren et al., 2022). The resulted MURPs serve as landmarks of the trajectory and help to mitigate the impact of random noises. Then, MGPfact infers the trajectory of cell fate as a mixture Gaussian process. To address the branching events in cell fate, each Gaussian process is defined with a bifurcation point. We then performed factorization of the scRNA expression profile to generate multiple independent differentiation trajectories, along with corresponding gene modules. Finally, multiple factorized trajectories are combined to form a coherent diffusion tree to represent the trajectory of cell fate.

#### 1.2 MURP Downsampling

Single-cell transcriptome (scRNA-seq) data often suffer from imbalance due to technical factors. In MGPfact, we use a downsampling technique called "minimum unbiased representative points (MURPs)(Ren et al., 2022) " to address this issue. This method aims to reduce noise while retaining valuable information by selecting a subset of points that can unbiasedly represent the data.

We performed downsampling of the preprocessed scRNA-seq data $\boldsymbol{Y}$ to yield a $M$-by-$N$ expression matrix $\boldsymbol{Y}'$ based on the MURP(Ren et al., 2022), where $M$ representative points were considered as landmarks of the cellular trajectory and $N$ is the number of genes. Then, we computed $L$ Principal Components (PC) of the downsampled expression matrix to obtain the matrix $\boldsymbol{Y}^* = \{\boldsymbol{y}_1^*, \boldsymbol{y}_2^*, \boldsymbol{y}_3^*, \dots \boldsymbol{y}_L^*\}$ ($M$-by-$L$),

$$\boldsymbol{y}_l^* = \boldsymbol{Y}' \cdot \mathbf{v}_l \qquad (1)$$

where $\mathbf{v}_l$ is projection vector, $\boldsymbol{y}_l^*$ serve as the $l$-th initial state of embedding.

#### 1.3 Trajectory inference

We used typical Gaussian Process Regression of $\boldsymbol{y}_l^*$ on pseudotime $\boldsymbol{T}$:

$$\boldsymbol{y}_l^* = f(\boldsymbol{T}) + \varepsilon \qquad (2)$$

Here, $f(\boldsymbol{T})$ is modeled as a Gaussian Process (GP) with covariance matrix $\boldsymbol{S}$, where ε denotes the error term.

$$f(\boldsymbol{T}) = \mathcal{GP}(0, \boldsymbol{S} + \sigma_S^2 \cdot \boldsymbol{I}) \tag{3}$$

The term $\sigma_S^2$ represents the variance, and $\boldsymbol{I}$ is the identity matrix. We specified $\sigma_S^2$ to 1e-6.

$$p(\varepsilon) = \mathcal{N}(0, \sigma_S^2) \tag{4}$$

For all features:

$$p\left(\boldsymbol{Y}^*, f(\boldsymbol{T})\right) = p\left(\boldsymbol{Y}^* \middle| f(\boldsymbol{T})\right) \cdot p(\boldsymbol{f}(\boldsymbol{T})) \tag{5}$$

where $p\left(\mathbf{Y}^* \middle| f(\mathbf{T})\right)$ is defined as follow:

$$p\left(\boldsymbol{Y}^* \middle| f(\boldsymbol{T})\right) = \prod_{l=1}^{L} p\left(\boldsymbol{y}_l^* \middle| f(\boldsymbol{T})\right) = \prod_{l=1}^{L} \mathcal{N}(\boldsymbol{y}_l^* | 0, \boldsymbol{S} + \sigma_S^2 \cdot \boldsymbol{I}) \tag{6}$$

For any prior of $t_x$, it satisfies:

$$p(t_x) = \mathcal{N}(\mu_T, \sigma_T) \tag{7}$$

$$p(\mu_T) = \mathcal{N}(0.5,\ 99) \tag{8}$$

$$p(\sigma_T) = \mathcal{IG}(0.01, 0.01) \tag{9}$$

We consider $\boldsymbol{S}$ as a mixture of $L$ independent bifurcating Gaussian processes(Schulz et al., 2018),

$$\boldsymbol{S} = \sum_{l=1}^{L} \boldsymbol{s}_l \tag{10}$$

To account for the bifurcation events in various differentiation trajectories, we introduce bifurcation points $\mathbf{B} = \{b_1, b_2, \cdots, b_L\}$. The prior distribution of $b_l$ satisfies:

$$p(b_l) = \Gamma\left(l, \frac{10}{L}\right) \tag{11}$$

Additionally, we introduce branching labels $\mathbf{C} = \{\boldsymbol{c_1}, \boldsymbol{c_2}, \ldots \boldsymbol{c_L}\}$ ($M$-by-$L$). The branching labels $\boldsymbol{c_l} \in \{0,1,2\}$, corresponds to different phase and states of cell fate, where $\boldsymbol{c_l} = \mathbf{0}$ corresponds the phase before branching, and $\boldsymbol{c_l} \in \{1,2\}$, corresponds to the two cellular states of the bifurcating process, respectively. For any one landmark (MURP) $x$,

$$\begin{cases} c_{l,x} = 0, & if\ t_x < b_l \\ c_{l,x} \in \{1,2\}, & if\ t_x \ge b_l \end{cases} \quad (12)$$

For $t_x \ge b_l$, we introduce the continuous variable $\mathbf{\Pi} = \{\boldsymbol{\pi_1}, \boldsymbol{\pi_2}, \dots, \boldsymbol{\pi}_L\}$ ($M$-by-$L$) to map $\boldsymbol{c_l}$ into a continuous space.

$$c_{l,x} = \begin{cases} 1, & \pi_{l,x} \ge 1 - \pi_{l,x} \\ 2, & \pi_{l,x} < 1 - \pi_{l,x} \end{cases} \quad (13)$$

The prior distribution of the $\pi_{l,x}$ satisfies:

$$p(\pi_{l,x}) = \mathcal{B}(v_{l,x} \cdot (1-\rho), v_{l,x} \cdot \rho) \quad (14)$$

Where $\rho$ satisfies:

$$p(\rho) = \mathcal{B}(1,1) \quad (15)$$

The activation parameter $v_{l,x}$ links $t_x$ and $b_l$, and is derived from $\mathbf{\Upsilon} = \{\boldsymbol{v_1}, \boldsymbol{v_2}, \dots, \boldsymbol{v_L}\}$ ($M$-by-$L$). We use the sigmoid function map $v_{l,x}$ to the range $[0,1]$:

$$v_{l,x} = \frac{1}{1 + e^{-(b_l - t_x)}} \quad (16)$$

Then, the covariance matrix $\boldsymbol{s_l}$ for any trajectories can be expressed as follows:

$$[s_l]_{x,y} = \mathcal{K}(t_x, t_y) \quad (17)$$

We create the $T$ component of the covariance matrix as follows:

$$\mathcal{K}(t_x, t_y) = \begin{cases} k_{rbf}(t_x, t_y) + k_{pl}(t_x, t_y) & t_x, t_y < b_l \\ k_{rbf}(t_x, t_y) + k_{pl}(t_x - b_l, t_y - b_l) & t_x, t_y > b_l, c_{l,x} = c_{l,y} \\ \dfrac{k_{rbf}(t_x, b_l) \cdot k_{rbf}(b_l, t_y)}{k_{rbf}(b_l, b_l)} & c_{l,x} \ne c_{l,y} \end{cases} \quad (18)$$

where $k_{rbf}$ is the Radial Basis Function (RBF) kernel, $k_{pl}$ is the Polynomial kernel function (PL) kernel. The RBF kernel is chosen for its ability to effectively model smooth functions and capture local variations in the data, making it well-suited to the continuous and smooth characteristics of biological processes; its hyperparameters offer modeling flexibility. The polynomial kernel is used to capture more complex nonlinear relationships between input features, with its hyperparameters also allowing further customization of the model.

$$k_{rbf}(t_x, t_y) = \lambda_{rbf} \cdot e^{\left(-\alpha_{rbf} \left\| t_x - t_y \right\|^2\right)} \quad (19)$$

$$k_{pl}(t_x, t_y) = \left(\lambda_{pl} {t_x}^T t_y + c_{pl}\right)^{d_{pl}} \quad (20)$$

The hyperparameters for the RBF kernel are: $\alpha_{\mathrm{rbf}}$ - determines input variable similarity and smoothness of changes, default value is $10^5$. $\lambda_{rbf}$ - length scale parameter influencing the variance of the GP, with a prior distribution as follows:

$$p(\lambda_{rbf}) = \mathcal{N}(0,99) \tag{21}$$

The hyperparameters for the PL kernel are: $c_{pl}$ - a constant term, influencing the 0-th order feature, is set to 0 in the MGPfact; $d_{pl}$ - controls the order of the PL, default value is 2; $\lambda_{pl}$ - scales the distance in the feature space and controls feature correlation, with a prior distribution as follows:

$$p(\lambda_{pl}) = \mathcal{N}(0,99) \tag{22}$$

Additionally, we insert a variable $\boldsymbol{Z} = \{z_1, z_2, \dots, z_M\}$ on the covariance matrix that represents the unknown components.

$$p(z_x) = \mathcal{N}(\mu_Z, \sigma_Z) \tag{23}$$

$$p(\mu_Z) = \mathcal{N}(0.5, 99) \tag{24}$$

$$p(\sigma_Z) = \mathcal{IG}(0.01, 0.01) \tag{25}$$

The final composition of the covariance matrix is as follows after applying the RBF to quantify $\mathbf{Z}$:

$$\mathcal{K}^*(t_x, t_y) = \mathcal{K}(t_x, t_y) + \lambda_{rbf} \cdot k_{rbf}(z_x, z_y) \tag{26}$$

$$[\mathbf{s}_l^*]_{x,y} = \mathcal{K}^*(t_x, t_y) \tag{27}$$

Therefore, $p(\mathbf{Y}^* | f(\mathbf{T}))$ is updated as follows:

$$p(\boldsymbol{Y}^* | f(\boldsymbol{T})) = \prod_{l=1}^{L} \mathcal{N}(\boldsymbol{y}_l^* | 0, \textstyle\sum_{l=1}^{L} \boldsymbol{s}_l^* + \sigma_S^2 \cdot \boldsymbol{I}) \tag{28}$$

We infer parameters by maximizing the posterior likelihood using Markov Chain Monte Carlo (MCMC) method available in Mamba (B J, 2014). The posterior distribution of pseudotime $\boldsymbol{T}$ can be represented as:

$$p(\boldsymbol{T} \mid \boldsymbol{Y}^*) \propto p(\boldsymbol{Y}^* \mid f(\boldsymbol{T})) \cdot p(f(\boldsymbol{T})) \tag{29}$$

where $p(\boldsymbol{Y}^* \mid f(\boldsymbol{T}))$ is the likelihood function of the observed data $\boldsymbol{Y}^*$, and $p(f(\boldsymbol{T}))$ is the prior distribution of the Gaussian process. This posterior distribution seamlessly integrates the observed data with model priors, enabling robust inference of pseudotime. Due to the high autocorrelation of $T$ in the posterior distribution, we use Adaptive Metropolis within Gibbs (AMWG) sampling (Roberts and Rosenthal, 2009; Tierney, 1994). Other parameters are estimated using the more efficient SLICE sampling technique(Neal, 2003).

##### 1.4 Extract gene modules

Through a factorization process, MGPfact can identify genes that have significant impacts during cell differentiation. We introduce a rotation matrix $\boldsymbol{R} = \{r_1, r_2, \ldots, r_L\}$ to obtain factor score $\boldsymbol{w_l}$ for each trajectory $l$ by rotating $\mathbf{Y}^*$.

$$\mathbf{w}_l = \mathbf{Y}^* \cdot \mathbf{r}_l + e_l^2 \tag{30}$$

$e_l$ represents the error term for the $l$-th trajectory, which satisfies:

$$p(e_l) = \mathcal{N}(0, \sigma_{error}^2) \tag{31}$$

$\sigma_{error}^2$ follows:

$$p(\sigma_{error}^2) = \mathcal{IG}(0.01,0.01) \tag{32}$$

The prior of any $\mathbf{w}_l$ follows an independent GP, where the covariance matrix $\mathbf{s}_l$ is obtained through previous computations. We have removed the unknown variables $\mathbf{Z}$ and retained only the part related to $\mathbf{T}$.

$$p(\mathbf{w_l}) = \mathcal{N}(0, \mathbf{s}_l) \tag{33}$$

Therefore, for all trajectories,

$$p(\boldsymbol{W}|\boldsymbol{Y}^*) = \prod_{l=1}^{L}[\mathcal{N}(\boldsymbol{Y}^* \cdot \boldsymbol{r}_l + e_l|0, \boldsymbol{s}_l) \cdot \mathcal{N}(e_l|0, \sigma_{error}^2)] \tag{34}$$

Specifically, the loading matrix for each gene onto the $l$-th trajectory can be expressed using equations (1) and (30) as follows, which is used to represent the contribution of each gene to a specific trajectory.

$$\mathbf{w}_l = [\boldsymbol{Y}' \cdot \mathbf{v}_l] \cdot \mathbf{r}_l + e_l^2 = \mathbf{Y}' \cdot \mathbf{u}_l + e_l^2 \tag{35}$$

which enables factorization and gene-selection based on the inferred trajectories.

##### 1.5 Generating the consensus trajectory

After conducting MGPfact factorization, we obtained $L$ independent differentiation trajectories, with each trajectory representing a binary tree of branching events in the time domain. We merge multiple factorized trajectories to form a coherent diffusion tree, representing the consensus trajectory of cell fate. The specific merging process is as follows:

1) First, we sort all trajectories based on bifurcation point $B$ to ensure that each trajectory's subtree is processed before its parent node.

2) Then, we select the first trajectory as the initial trajectory.

3) For the subsequent trajectories, we connect them to the previous trajectory based on their branching time points, following the inheritance relationships. The specific procedure is as follows:

a. For trajectory $l$, based on its bifurcation point $b_l$ and the bifurcation point of the previous trajectory $b_{l-1}$, all points can be divided into three parts:

- The first part consists of points before $b_{l-1}$, denoted as $\mathcal{G}_{l,1} = \{\mathbf{T} \mid \mathbf{T} < b_{l-1}\}$
- The second part consists of points between $b_{l-1}$ and $b_l$, denoted as $\mathcal{G}_{l,2} = \{\mathbf{T} \mid b_{l-1} \leq \mathbf{T} < b_l\}$
- The third part consists of points after $b_l$, denoted as $\mathcal{G}_{l,3} = \{\mathbf{T} \mid \mathbf{T} \geq b_l\}$.

b. These three parts inherit the branching labels $[c_{l-1}]$ from the previous $(l-1)$-th trajectory. For the set $\mathcal{G}_{i,3}$, these points need to inherit the branching relationships from the previous trajectory and are located after the current trajectory's branching time point. Therefore, based on the current trajectory's branching relationships $[c_l]_{\mathcal{G}_{l,3}}$, we update the branching labels to form a new subtree.

c. Through iterative execution of step b, we effectively consolidate multiple trajectories into a comprehensive binary tree known as the consensus trajectory.

### 2 Supplementary Figures

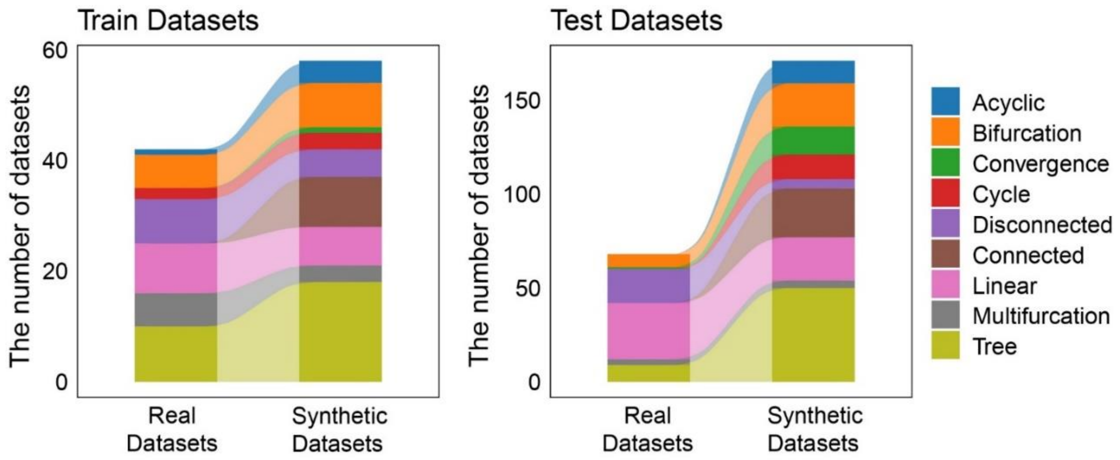


**Supplementary Fig. 1. The distribution of trajectory types among training set and test set**. The 339 datasets were randomly split into two groups, serving as the training set and test set, respectively. The training set consists of 100 datasets, while the testing set includes 239 datasets.

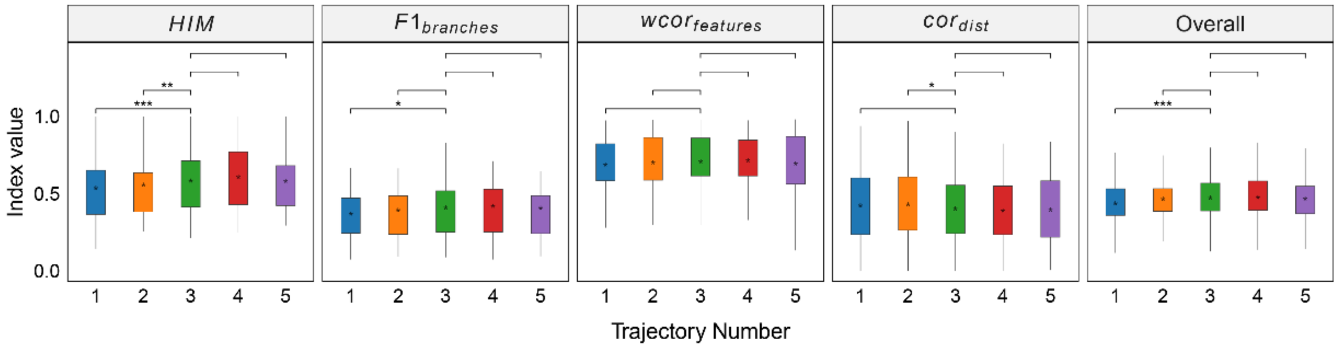


**Supplementary Fig. 2. Robustness testing of the number of independent trajectories on 100 training datasets.** With L=3 set as the default, we tested the impact of different L values (1, 2, 4, and 5) on the prediction results. In all box plots, the asterisk represents the mean value, while the whiskers extend to the farthest data points within 1.5 times the interquartile range. Significance is denoted as follows: not annotated indicates non-significant; * P < 0.05; ** P < 0.01; *** P < 0.001; two-sided paired Student's T-tests.

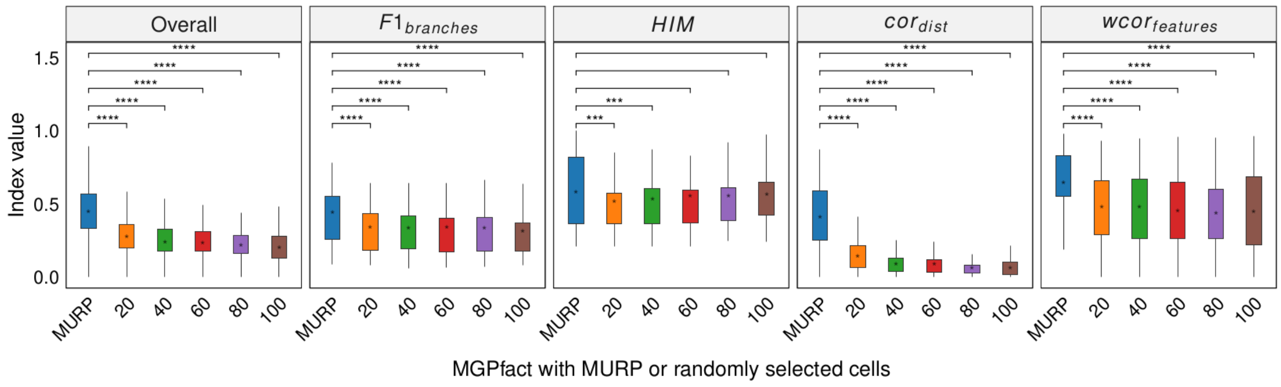


**Supplementary Fig. 3. MURP downsampling enhances the performance of MGPfact.** Trajectory inference was conducted by randomly selecting 20, 40, 60, 80, and 100 cells from the original dataset. The results were mapped back to the original data using a KNN graph structure to obtain the final predictions, which were then compared with those obtained through MURP downsampling. In all box plots, the asterisk represents the mean value, while the whiskers extend to the farthest data points within 1.5 times the interquartile range. Significance is denoted as follows: not annotated indicates non-significant; * P < 0.05; ** P < 0.01; *** P < 0.001; two-sided paired Student's T-tests.

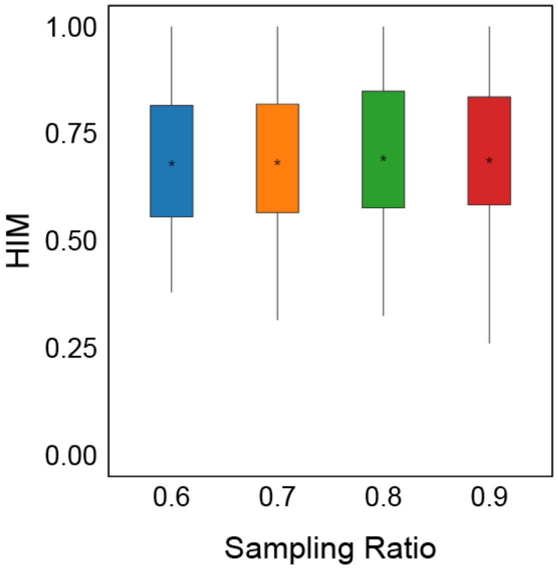


**Supplementary Fig. 4. Robustness analysis of consensus trees.** Random sampling is performed on the original data at different proportions of 60%, 70%, 80%, and 90%. The MGPfact consensus trajectory predictions are compared with that based on the original, unsampled data using $HIM$. In all box plots, the asterisk represents the mean value, while the whiskers extend to the farthest data points within 1.5 times the interquartile range.

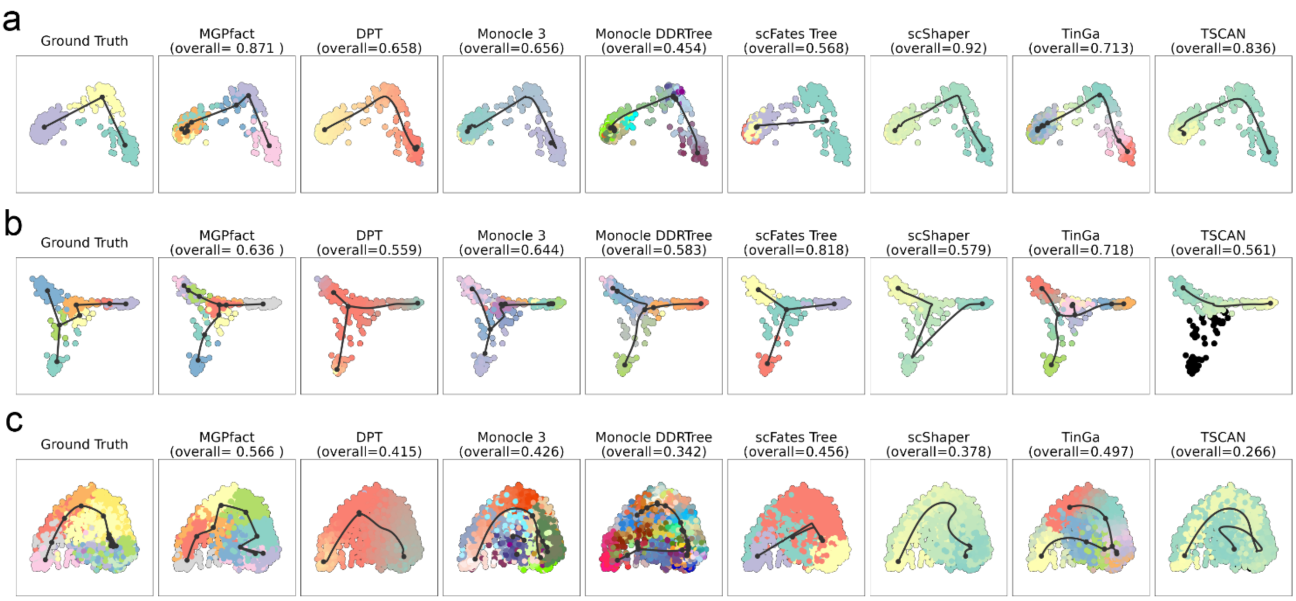


**Supplementary Fig. 5. Trajectories identified by different methods on 3 real-world datasets, with reference structures being linear (a, dataset-id=real/silver/germline-human-female_li), bifurcation (b, dataset-id=real/silver/fibroblast-reprogramming_treutlein), and multifurcation (c, dataset-id= real/silver/oligodendrocyte-differentiation-subclusters_marques).** The first column of each row presents the actual cell development trajectory ("ground truth"), with black lines representing the trajectory structure and the colored dots indicating cells at different stages. These elements highlight the main paths and key bifurcation points of cell differentiation as a reference for evaluating prediction results. Subsequent columns show the trajectory reconstruction results of MGPfact and other methods. The ovreall score for each method reflects its accuracy in trajectory inference relative to the gold standard, which is represented in the top-left figure.

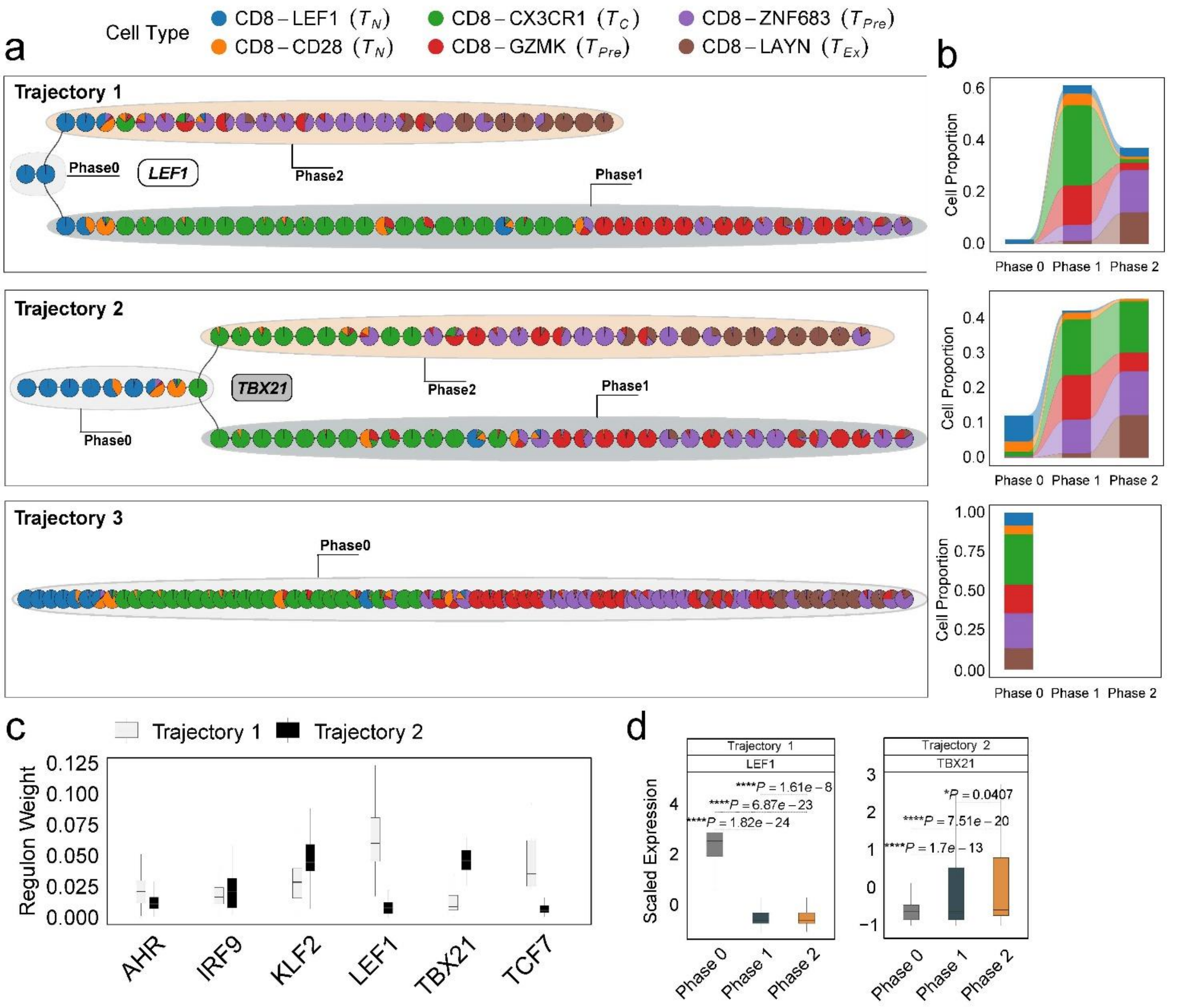


**Supplementary Fig. 6. Differentiation process and determinants of CD8$^+$ T cells from NSCLC environment. a.** The independent trajectories, where phase 0 represents the juncture before bifurcation, while phases 1 and 2 represent the diverging branches post-bifurcation. **b.** Proportions of cell types in different phases of independent trajectories; **c.** The regulon weight of top regulons of two trajectories. **d.** The differential expression of top transcription factors among different phases for each trajectory. In all box plots, the horizontal line represents the median value, and the whisker extends to the furthest data point within 1.5 times the interquartile range. Significance denoted as: * $P < 0.05$, **** $P < 0.0001$, two-sided unpaired Student’s t test.

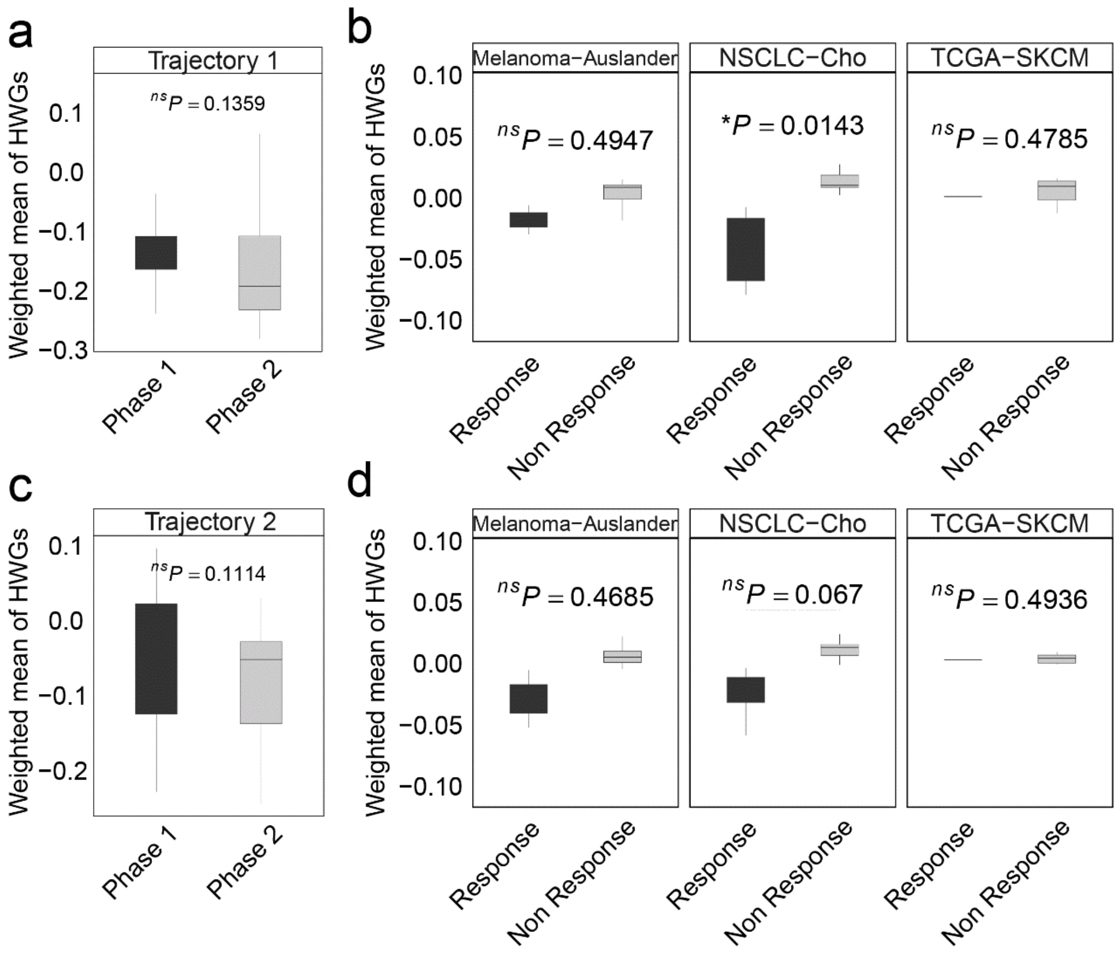


**Supplementary Fig. 7. In NSCLC, the weighted mean of highly weighted genes (HWG) from independent differentiation trajectories differs between phases (a, c) and ICI treatment groups.** a-b. The HWG obtained from the first branching event. c-d. The HWG obtained from the first branching event. In all box plots, the horizontal line represents the median value, and the whisker extends to the furthest data point within 1.5 times the interquartile range. Significance denoted as: ns, not significant, * P < 0.05, two-sided unpaired Student's t test.

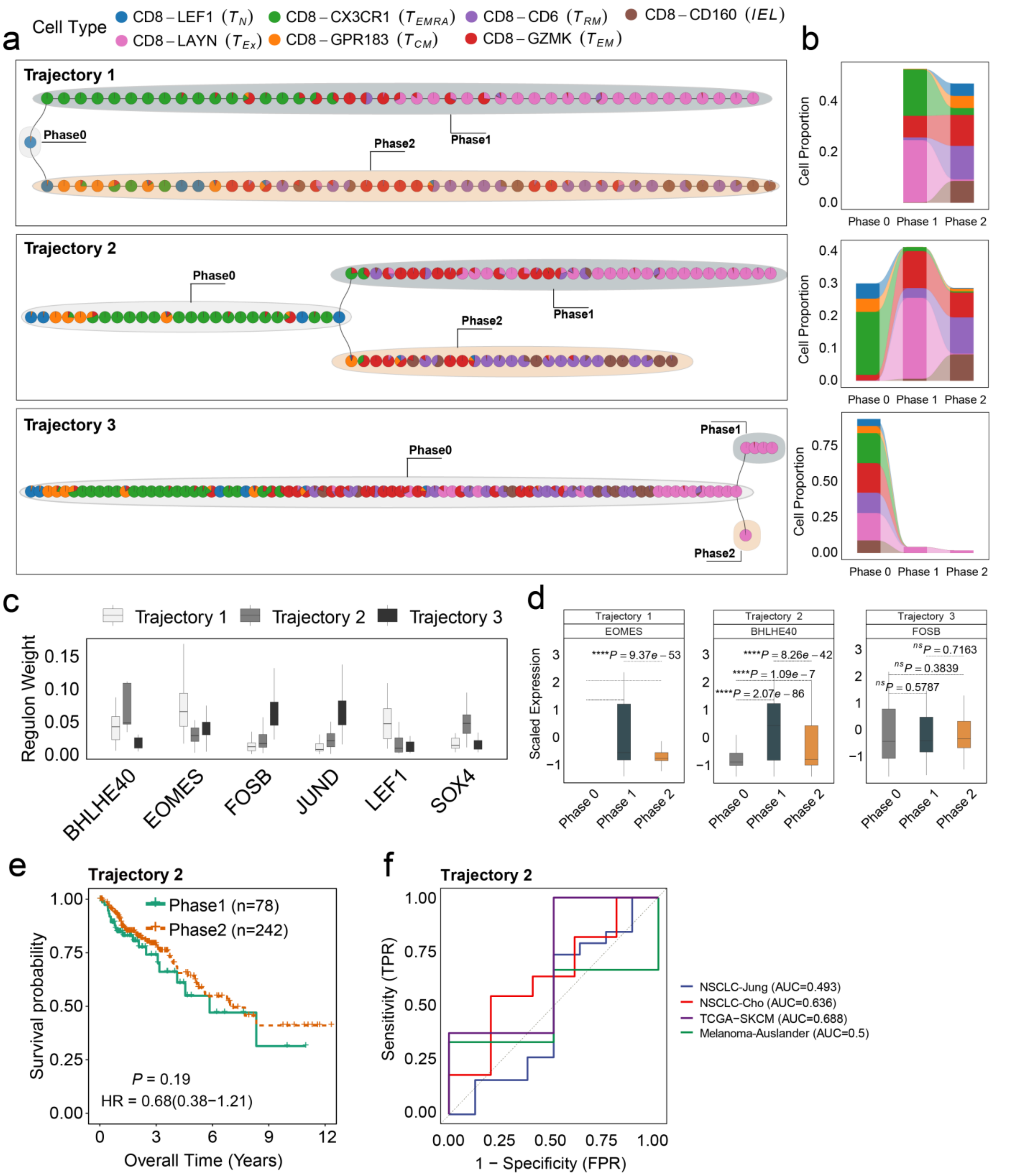


**Supplementary Fig. 8. Differentiation process and determinants of CD8$^{+}$ T cells from CRC environment. a.** The independent trajectories, where phase 0 represents the juncture before bifurcation, while phases 1 and 2 represent the diverging branches post-bifurcation. **b.** Proportions of cell types in different phases of independent trajectories; **c**. The regulon weight of top regulons of two trajectories. **d.** The differential expression of top transcription factors among different phases for each trajectory. **e.** Survival analysis of the patients from TCGA-COAD cohort. The patients are stratified by utilizing the branch event within trajectory 2 from MGPfact analysis. P-values are calculated using multivariate Cox regression, with HR representing the hazard ratio. **f.** ROC curve for ICI treatment response associated with highly weighted genes related to Trajectories 2 in CRC across 4 independent studies. In all box plots, the horizontal line represents the median value, and the whisker extends to the furthest data point within 1.5 times the interquartile range. Significance denoted as: ns, not significant, * P < 0.05, ** P < 0.01, *** P < 0.001, **** P < 0.0001, two-sided unpaired Student's t test.

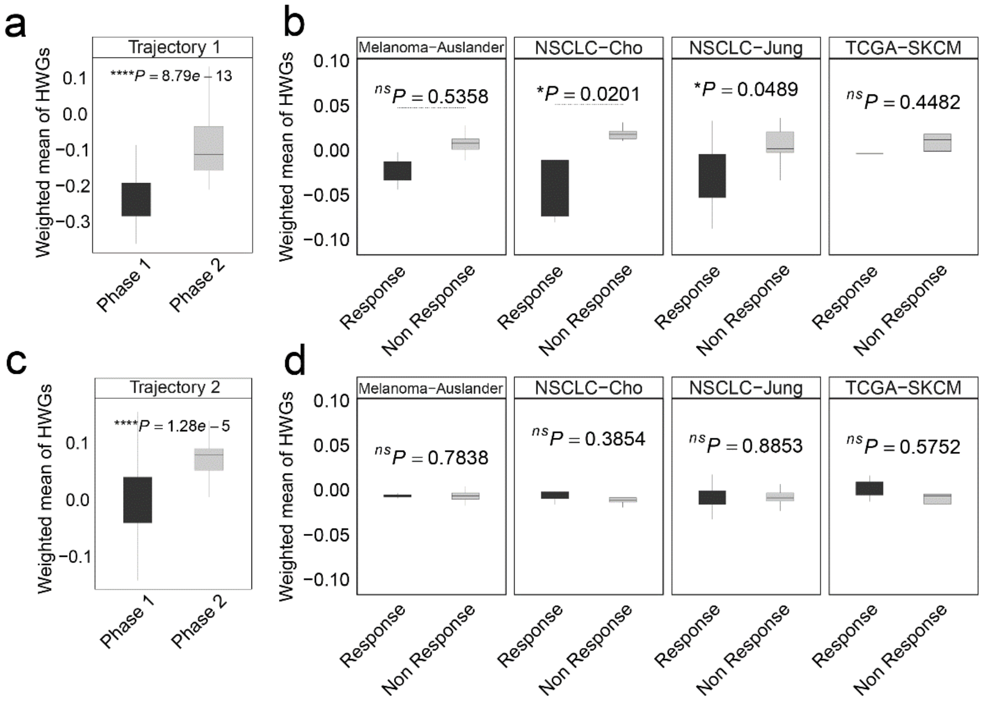


**Supplementary Fig. 9. In CRC, the weighted mean of highly weighted genes (HWG) from independent differentiation trajectories differs between phases (a, c) and ICI treatment groups.** a-b. The HWG obtained from the first branching event. c-d. The HWG obtained from the first branching event. In all box plots, the horizontal line represents the median value, and the whisker extends to the furthest data point within 1.5 times the interquartile range. Significance denoted as: ns, not significant, * P < 0.05, **** P < 0.0001, two-sided unpaired Student's t test.

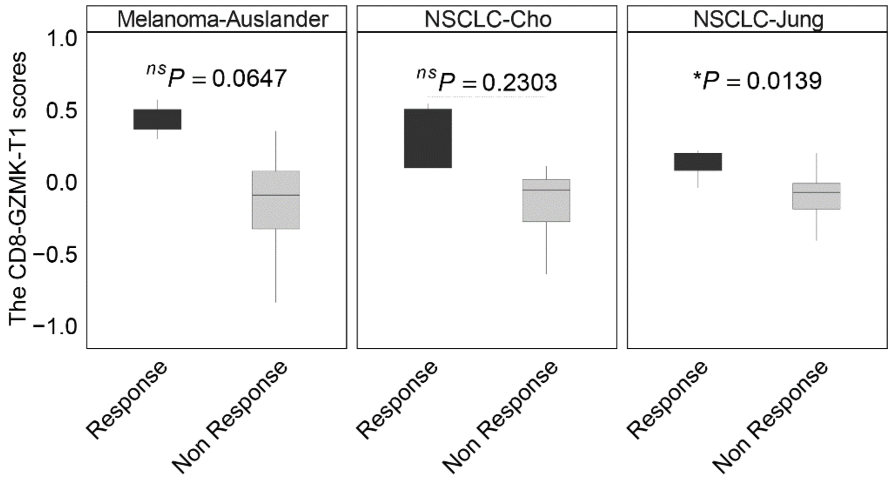


**Supplementary Fig. 10. The CD8-GZMK-T1 scores between ICI treatment response groups (Methods).** In all box plots, the horizontal line represents the median value, and the whisker extends to the furthest data point within 1.5 times the interquartile range. Significance denoted as: ns, not significant, * P < 0.05, two-sided unpaired Student's t test.
