## supplementary tables for "MGPfact^XMBD^: A Model-Based Factorization Method for scRNA Data Unveils Bifurcating Transcriptional Modules Underlying Cell Fate Determination": supplementary_table_1_model.pdf

**Supplementary Table 1: The specific configuration of parameters contained in the model.**

| | | |
|---|---|---|
| $p(\boldsymbol{Y}^{*}\|f(\boldsymbol{T})) = \prod_{l=1}^{L} \mathcal{N}\left(\boldsymbol{y}_l^{*}\middle\|0, \sum_{l=1}^{L} \boldsymbol{s}_l^{*} + \sigma_S^2 \cdot \boldsymbol{I}\right)$ | $p(\varepsilon) = \mathcal{N}\left(0, \sigma_S^2\right)$ | |
| | $\sigma_S^2$=1e-6 | |
| $[\boldsymbol{s}_l^{*}]_{x,y} = \mathcal{K}\left(t_x, t_y\right) + \lambda_{rbf} \cdot k_{rbf}\left(z_x, z_y\right.$ | | |
| $\mathcal{K}\left(t_x, t_y\right) = \begin{cases} k_{rbf}\left(t_x, t_y\right) + k_{pl}\left(t_x, t_y\right) & t_x, t_y < b_l \\ k_{rbf}\left(t_x, t_y\right) + k_{pl}\left(t_x - b_l, t_y - b_l\right) & t_x, t_y > b_l, c_{l,x} = c_{l,y} \\ \dfrac{k_{rbf}(t_x, b_l) \cdot k_{rbf}\left(b_l, t_y\right)}{k_{rbf}(b_l, b_l)} & c_{l,x} \neq c_{l,y} \end{cases}$ | $k_{rbf}\left(t_x, t_y\right) = \lambda_{rbf} \cdot e^{\left(-\alpha_{rbf}\left\|\left\|t_x - t_y\right\|\right\|^2\right)}$ | $\alpha_{rbf}$=1e+5 |
| | | $p\left(\lambda_{rbf}\right) = \mathcal{N}(0,99)$ |
| | $k_{pl}\left(t_x, t_y\right) = \left(\lambda_{pl} {t_x}^T t_y + c_{pl}\right)^{d_{pl}}$ | $c_{pl}$=0 |
| | | $d_{pl}$=2 |
| | | $p\left(\lambda_{pl}\right) = \mathcal{N}(0,99)$ |
| $p\left(t_x\right) = \mathcal{N}(\mu_T, \sigma_T)$ | $p(\mu_T) = \mathcal{N}(0.5,\ 99)$ | |
| | $p(\sigma_T) = \mathcal{IG}(0.01{,}0.01)$ | |
| $p(b_l) = \Gamma\left(l, \dfrac{10}{L}\right)$ | | |
| $p\left(\pi_{l,x}\right) = \mathcal{B}\left(v_{l,x} \cdot (1 - \rho), v_{l,x} \cdot \rho\right)$ | $p(\rho) = \mathcal{B}(1{,}1)$ | |
| | $v_{l,x} = \dfrac{1}{1 + e^{-(b_l - t_x)}}$ | |
| $p(z_x) = \mathcal{N}(\mu_Z, \sigma_Z)$ | $p(\mu_Z) = \mathcal{N}(0.5, 99)$ | |
| | $p(\sigma_Z) = \mathcal{IG}(0.01, 0.01)$ | |
