## supplementary tables for "MGPfact^XMBD^: A Model-Based Factorization Method for scRNA Data Unveils Bifurcating Transcriptional Modules Underlying Cell Fate Determination": supplementary_table_2_pvalue.pdf

**Supplementary Table 2: P-values associated with one-sided paired t-tests assessing whether the overall performance score of MGPfact significantly higher than the other methods on the different trajectory types in test set.**

| Dataset Trajectory Type | DPT | Monocle DDRTree | Monocle 3 | scFates Tree | scShaper | TInGa | TSCAN |
|---|---|---|---|---|---|---|---|
| Acyclic Graph | **0.003** | 0.974 | 0.652 | 0.168 | **0.005** | 0.861 | **0.002** |
| Bifurcation | **0.012** | **0.074** | 0.166 | 0.224 | **0.045** | 0.942 | **0.000** |
| Convergence | 0.276 | 0.235 | **0.001** | **0.071** | **0.049** | 0.940 | **0.002** |
| Cycle | **0.000** | 0.195 | 0.463 | **0.092** | 0.984 | 1.000 | 0.861 |
| Disconnected Graph | **0.000** | **0.049** | 0.915 | 0.126 | **0.001** | 0.939 | **0.000** |
| Connected Graph | **0.001** | 1.000 | 0.988 | 0.224 | **0.002** | 0.924 | **0.000** |
| Linear | **0.000** | **0.000** | **0.001** | **0.000** | 1.000 | **0.013** | **0.059** |
| Multifurcation | **0.003** | **0.012** | 0.120 | 0.559 | **0.040** | 0.246 | **0.001** |
| Tree | **0.000** | 1.000 | 1.000 | 0.069 | **0.000** | 1.000 | **0.000** |

**Supplementary Table 2: P-values associated with one-sided paired t-tests assessing whether the HIM performance score of MGPfact significantly higher than the other methods on the different trajectory types in test set.**

| Dataset Trajectory Type | DPT | Monocle DDRTree | Monocle 3 | scFates.Tree | scShaper | TInGa | TSCAN |
|---|---|---|---|---|---|---|---|
| Acyclic Graph | 0.786 | 1.000 | 0.857 | 0.326 | **0.001** | 0.796 | **0.001** |
| Bifurcation | 0.920 | 0.211 | **0.076** | 0.618 | **0.004** | 0.605 | **0.004** |
| Convergence | 0.674 | 0.150 | **0.000** | 0.592 | **0.007** | 0.711 | **0.007** |
| Cycle | 0.182 | 1.000 | 0.846 | 0.787 | **0.083** | 1.000 | **0.083** |
| Disconnected Graph | **0.013** | 0.858 | 0.919 | 0.991 | **0.009** | 0.938 | **0.009** |
| Connected Graph | 0.969 | 1.000 | 0.995 | 0.976 | **0.000** | 0.792 | **0.000** |
| Linear | **0.000** | **0.000** | **0.000** | **0.000** | 0.998 | **0.000** | 0.998 |
| Multifurcation | 0.733 | 0.895 | 0.957 | 0.728 | **0.019** | **0.011** | **0.019** |
| Tree | 1.000 | 1.000 | 1.000 | 1.000 | **0.000** | 1.000 | **0.000** |

**Supplementary Table 2: P-values associated with one-sided paired t-tests assessing whether the $wcor_{features}$ performance score of MGPfact significantly higher than the other methods on the different trajectory types in test set.**

| Dataset Trajectory Type | DPT | Monocle DDRTree | Monocle 3 | scFates.Tree | scShaper | TInGa | TSCAN |
|---|---|---|---|---|---|---|---|
| Acyclic Graph | **0.001** | 0.892 | 0.573 | 0.219 | **0.024** | 0.887 | **0.068** |
| Bifurcation | **0.000** | **0.073** | 0.350 | **0.077** | **0.002** | 0.773 | **0.001** |
| Convergence | **0.095** | 0.246 | 0.477 | 0.123 | **0.049** | 0.784 | 0.209 |
| Cycle | **0.001** | 0.952 | 0.782 | 0.732 | 0.767 | 0.706 | 0.322 |
| Disconnected Graph | **0.008** | 0.257 | 0.993 | **0.091** | **0.001** | 0.917 | **0.068** |
| Connected Graph | **0.000** | 0.965 | 0.893 | 0.001 | **0.010** | 0.785 | **0.010** |
| Linear | **0.000** | 0.352 | 0.964 | 0.144 | 0.998 | 0.958 | 0.170 |
| Multifurcation | **0.003** | **0.046** | 0.379 | **0.084** | **0.016** | **0.086** | **0.027** |
| Tree | **0.000** | 0.868 | 1.000 | 0.106 | **0.000** | 0.995 | **0.011** |

**Supplementary Table 2: P-values associated with one-sided paired t-tests assessing whether the $cor_{dist}$ performance score of MGPfact significantly higher than the other methods on the different trajectory types in test set.**

| Dataset Trajectory Type | DPT | Monocle DDRTree | Monocle 3 | scFates.Tree | scShaper | TInGa | TSCAN |
|---|---|---|---|---|---|---|---|
| Acyclic Graph | **0.021** | 0.772 | 0.678 | 0.276 | 0.505 | 0.996 | 0.204 |
| Bifurcation | 0.131 | 0.987 | 0.937 | 0.749 | 0.994 | 0.999 | **0.002** |
| Convergence | 0.595 | 0.999 | 0.800 | 0.445 | 0.987 | 0.991 | **0.017** |
| Cycle | **0.001** | 0.999 | 0.998 | 0.358 | 0.994 | 1.000 | 0.658 |
| Disconnected Graph | **0.011** | 1.000 | 1.000 | 0.935 | 0.527 | 0.992 | **0.065** |
| Connected Graph | **0.056** | 0.999 | 0.969 | 0.415 | 0.976 | 1.000 | **0.028** |
| Linear | **0.027** | 0.999 | 0.999 | 0.875 | 1.000 | 1.000 | **0.028** |
| Multifurcation | **0.064** | 0.514 | 0.120 | 0.758 | 0.569 | 0.731 | **0.015** |
| Tree | **0.003** | 1.000 | 0.993 | 0.223 | 0.699 | 1.000 | **0.000** |

**Supplementary Table 2: P-values associated with one-sided paired t-tests assessing whether the $F1_{branches}$ performance score of MGPfact significantly higher than the other methods on the different trajectory types in test set.**

| Dataset Trajectory Type | DPT | Monocle DDRTree | Monocle 3 | scFates.Tree | scShaper | TInGa | TSCAN |
|---|---|---|---|---|---|---|---|
| Acyclic Graph | 0.169 | 0.735 | 0.341 | 0.167 | **0.006** | 0.485 | **0.002** |
| Bifurcation | 0.559 | **0.001** | **0.015** | 0.298 | **0.010** | 0.772 | **0.002** |
| Convergence | 0.275 | **0.059** | **0.000** | **0.027** | 0.120 | 0.871 | **0.008** |
| Cycle | **0.000** | **0.000** | **0.002** | 0.114 | 0.991 | 0.659 | 0.982 |
| Disconnected Graph | **0.020** | **0.001** | 0.108 | **0.007** | **0.001** | 0.475 | **0.000** |
| Connected Graph | **0.053** | 0.214 | 0.114 | 0.285 | **0.005** | **0.057** | **0.001** |
| Linear | **0.000** | **0.000** | **0.000** | **0.000** | 1.000 | **0.000** | 0.980 |
| Multifurcation | **0.033** | **0.001** | **0.041** | 0.717 | 0.552 | 0.758 | **0.051** |
| Tree | **0.021** | 0.809 | 0.918 | **0.086** | **0.000** | 0.993 | **0.000** |
