## supplementary tables for "MGPfact^XMBD^: A Model-Based Factorization Method for scRNA Data Unveils Bifurcating Transcriptional Modules Underlying Cell Fate Determination": supplementary_table_3_efficiency.pdf

**Supplementary Table 3: Comparison of memory utilization (GB) and temporal efficiency (min)**

| | The average maximum memory usage (GB) | Elapsed time (min) |
|---|---|---|
| TinGa | 0.55 | 0.16 |
| Monocle 3 | 0.68 | 0.54 |
| TSCAN | 0.68 | 0.64 |
| ScShaper | 0.91 | 0.83 |
| scFates Tree | 0.68 | 0.93 |
| DPT | 0.68 | 2.42 |
| **MGPfact** | **0.75** | **3.42** |
| Monocle DDRTree | 0.67 | 7.50 |
