## supplementary tables for "MGPfact^XMBD^: A Model-Based Factorization Method for scRNA Data Unveils Bifurcating Transcriptional Modules Underlying Cell Fate Determination": supplementary_table_4_gse123025_gene_weight_0.05.pdf

**Supplementary Table 4: The highly weighted genes associated with independent bifurcating processes in microglia development (absolute gene weight > 0.05).**

| Trajectory 1 | | Trajectory 2 | | Trajectory 3 | |
|---|---|---|---|---|---|
| Gene | Weight | Gene | Weight | Gene | Weight |
| Apoe | 0.24360146 | Psen1 | -0.05010239 | ERCC-00002 | 0.24509556 |
| Lyz2 | 0.14801909 | Arhgap19 | -0.05044649 | Cst3 | 0.21810378 |
| Lpl | 0.13976103 | Ctsl | -0.05055875 | Tmem119 | 0.20383928 |
| Cd63 | 0.13227215 | Nusap1 | -0.05064861 | Selplg | 0.18556118 |
| Spp1 | 0.11858374 | Pcyox1 | -0.05071016 | P2ry12 | 0.18211465 |
| Pkm2 | 0.11161999 | ERCC-00108 | -0.0507653 | ERCC-00046 | 0.174178 |
| Ftl1 | 0.1102561 | Ncaph | -0.05105601 | ERCC-00108 | 0.1573142 |
| Ctsl | 0.11019648 | Kif2c | -0.05124333 | Siglech | 0.13047841 |
| Ldha | 0.10898228 | Incenp | -0.05130036 | 4632428N05Rik | 0.12299337 |
| Plek | 0.09838422 | Ivns1abp | -0.05165504 | P2ry13 | 0.11914988 |
| Hif1a | 0.09828405 | Kif15 | -0.05168464 | Sepp1 | 0.11636426 |
| Gatm | 0.09351161 | Iqgap3 | -0.05226651 | Gpr56 | 0.11464711 |
| Abca1 | 0.0904025 | Nuf2 | -0.05249763 | Slco2b1 | 0.10488559 |
| Pld3 | 0.09004353 | Lrrc33 | -0.05251642 | Itgb5 | 0.10203121 |
| Sepp1 | 0.08971533 | Fam64a | -0.05283197 | Gpr34 | 0.10071877 |
| Plin2 | 0.08870985 | Atf7ip | -0.05363316 | ERCC-00022 | 0.09404665 |
| Mcm6 | 0.08822107 | Asf1b | -0.05462439 | Ctsh | 0.09306572 |
| Atp6v1b2 | 0.0829249 | Spc25 | -0.05482592 | Fgd2 | 0.09274442 |
| Wdr1 | 0.0827963 | Ncapg | -0.05510579 | Cmtm6 | 0.08959441 |
| Mcm3 | 0.08126813 | Aurka | -0.05517708 | Slc2a5 | 0.08877948 |
| Cd83 | 0.07983332 | Ogdh | -0.05567751 | Ptgs1 | 0.08778003 |
| C3ar1 | 0.07206513 | Kif20b | -0.05625308 | Rps6ka1 | 0.08679537 |
| Igf1 | 0.07113068 | Hmmr | -0.0566873 | ERCC-00092 | 0.08675221 |
| Anxa5 | 0.06991745 | Anln | -0.05696095 | Itgam | 0.08494034 |
| Cd9 | 0.06762955 | Gatm | -0.05697461 | Entpd1 | 0.08492381 |
| Sgpl1 | 0.067552 | Mis18bp1 | -0.05713845 | Ivns1abp | 0.08446367 |
| Gas6 | 0.06481251 | P2ry13 | -0.05722955 | Ftl1 | 0.08437287 |
| Serpine2 | 0.06238303 | Bub1b | -0.05746302 | Jun | 0.08244943 |
| Abcg1 | 0.06197675 | Tln1 | -0.05780222 | Serpine2 | 0.07361138 |
| Stab1 | 0.06097427 | Pom121 | -0.05813355 | ERCC-00112 | 0.0732127 |
| Soat1 | 0.06049531 | C3ar1 | -0.05825621 | Gal3st4 | 0.07278804 |
| Tpi1 | 0.05888454 | Fen1 | -0.05839838 | Lair1 | 0.07264729 |
| Sec61a1 | 0.05828091 | Ldha | -0.05914716 | Cd164 | 0.07228344 |
| Vps35 | 0.05719019 | Pbk | -0.05957543 | Il6ra | 0.07189311 |
| Myo1f | 0.05536775 | P2ry12 | -0.06007048 | Rab3il1 | 0.07113571 |
| Slc23a2 | 0.05429312 | Casc5 | -0.06027612 | Junb | 0.07100601 |
| Glt25d1 | 0.05393831 | Cenpf | -0.06030782 | Tmem173 | 0.07081865 |
| Gpx3 | 0.05374018 | Nde1 | -0.0606653 | Ssh2 | 0.06785463 |
| Hspa5 | 0.05361964 | Lpl | -0.06067363 | Ctsl | 0.06712546 |
| Itgb1 | 0.05343024 | Sec61a1 | -0.06081441 | Csf3r | 0.06671725 |
| Dab2 | 0.05286636 | Lbr | -0.06086407 | ERCC-00116 | 0.06422983 |
| Txndc5 | 0.05235416 | Neil3 | -0.06271371 | Hspa5 | 0.06343891 |
| Lrp1 | 0.0518027 | Plk1 | -0.06281239 | Sgk1 | 0.0594574 |
| Gpnmb | 0.05162566 | E130306D19Rik | -0.06282177 | Dusp6 | 0.05928076 |
| Asph | 0.05014166 | Actn4 | -0.06313936 | Macf1 | 0.05924534 |
| Tgfbr1 | -0.05020589 | Kif20a | -0.0635821 | Tgfbr1 | 0.05846194 |
| H1f0 | -0.05060331 | Tpx2 | -0.06358541 | Tcn2 | 0.0582778 |
| Iqgap3 | -0.05100379 | Itgb1 | -0.06368898 | Gna15 | 0.05820413 |
| E130306D19Rik | -0.05113265 | Anapc5 | -0.06426519 | Mertk | 0.05815588 |
| 4632428N05Rik | -0.05189701 | Cdca2 | -0.06453008 | Pisd-ps3 | 0.05742441 |
| Kpna2 | -0.05199001 | Cdca8 | -0.06474161 | Pla2g15 | 0.05734156 |
| Ier5 | -0.05269489 | Wdr1 | -0.06534941 | Lrrc3 | 0.05666185 |
| Ivns1abp | -0.05284625 | Ckap2l | -0.06589881 | Btg2 | 0.05574766 |
| Casc5 | -0.05318194 | Sepp1 | -0.06673108 | Slc7a8 | 0.05523304 |
| Kif20a | -0.0534329 | Ccnb2 | -0.06690692 | 1500010J02Rik | 0.054214 |
| Lmf2 | -0.05487428 | Mrc1 | -0.06738835 | Hmha1 | 0.05394787 |

| | | | | | |
|---|---|---|---|---|---|
| Ccnb1 | -0.05686139 | Wsb1 | -0.06757753 | Slc40a1 | 0.05232134 |
| Aurkb | -0.05844697 | H1f0 | -0.06825331 | Il10ra | 0.05202447 |
| Cdc20 | -0.0618077 | Ftl1 | -0.0685254 | Cd9 | 0.050893 |
| Ccna2 | -0.06198533 | Lmf2 | -0.06911619 | Tpst2 | 0.0507708 |
| Jun | -0.06200613 | Kif22 | -0.06946336 | Txnip | 0.05038161 |
| St3gal6 | -0.06227149 | Cenpe | -0.0701196 | Top2a | -0.06003 |
| P2ry12 | -0.0628894 | Mgat4a | -0.07064616 | Mki67 | -0.06663234 |
| Top2a | -0.06315959 | Kif11 | -0.07218601 | | |
| Gpr165 | -0.06608178 | Aurkb | -0.07311392 | | |
| Prc1 | -0.06968387 | Smc4 | -0.07320556 | | |
| Arl6ip1 | -0.07348925 | Rrm1 | -0.07334864 | | |
| Ccnf | -0.07475049 | Racgap1 | -0.07474855 | | |
| Slco2b1 | -0.08024412 | Tubb4b | -0.07517668 | | |
| Mki67 | -0.10309226 | Rrm2 | -0.07702516 | | |
| Tmem119 | -0.1364109 | Slc29a1 | -0.07750179 | | |
| Selplg | -0.16095245 | Ccnb1 | -0.0780361 | | |
| | | Ncapd2 | -0.07936265 | | |
| | | Tmpo | -0.07965469 | | |
| | | Tacc3 | -0.08305091 | | |
| | | Prc1 | -0.08332551 | | |
| | | Ccnf | -0.08478689 | | |
| | | Pkm2 | -0.08685558 | | |
| | | Cdc20 | -0.09010535 | | |
| | | Kpna2 | -0.09401147 | | |
| | | Ccna2 | -0.09558965 | | |
| | | Arsb | -0.09652996 | | |
| | | ERCC-00002 | -0.09915862 | | |
| | | Hnrnpu | -0.10309481 | | |
| | | Cd164 | -0.10504114 | | |
| | | Top2a | -0.11609805 | | |
| | | Arl6ip1 | -0.11763113 | | |
| | | Tubb5 | -0.13170358 | | |
| | | Mki67 | -0.1381645 | | |
| | | Apoe | -0.15568154 | | |
