## supplementary tables for "MGPfact^XMBD^: A Model-Based Factorization Method for scRNA Data Unveils Bifurcating Transcriptional Modules Underlying Cell Fate Determination": supplementary_table_5_PAM_seg8_pamt1_seg9_pamt2.pdf

**Supplementary Table 5: List of genes specifically expressed in HM.**
**PAM-T1 cells (n = 113) vs. PAM-T2 cells (n = 298), two-sided moderated t-test with limma**

| Gene | logFC | AveExpr | t | P.Value | adj.P.Val | B |
|---|---|---|---|---|---|---|
| Spp1 | -2.516742756 | 3.541603023 | -9.961491038 | 4.32E-21 | 1.06E-17 | 36.84283356 |
| Gpnmb | -1.733073185 | 1.834825054 | -10.57800704 | 2.66E-23 | 1.30E-19 | 41.7295689 |
| Lgals3 | -0.96045566 | 1.004195617 | -7.605408899 | 1.94E-13 | 1.05E-10 | 19.95335274 |
| Igf1 | -0.863437052 | 1.550059413 | -8.466204246 | 4.48E-16 | 3.65E-13 | 25.76353781 |
| Lgals1 | -0.793147793 | 0.905351401 | -8.997094751 | 8.51E-18 | 1.39E-14 | 29.56230441 |
| Anxa5 | -0.753806811 | 1.743710594 | -7.018805683 | 9.24E-12 | 3.23E-09 | 16.26143185 |
| Soat1 | -0.751489757 | 1.465706896 | -7.048970214 | 7.62E-12 | 2.86E-09 | 16.445709 |
| Anxa2 | -0.730859647 | 1.041583684 | -7.455856548 | 5.30E-13 | 2.59E-10 | 18.99073268 |
| Cd9 | -0.730838389 | 3.049843197 | -7.179587131 | 3.28E-12 | 1.33E-09 | 17.25073082 |
| Capg | -0.696154641 | 0.743621287 | -7.706748042 | 9.71E-14 | 5.93E-11 | 20.61374921 |
| Pkm2 | -0.610674217 | 2.752027331 | -6.884273628 | 2.17E-11 | 6.63E-09 | 15.44710656 |
| Ldha | -0.571123826 | 2.42875813 | -5.264352375 | 2.27E-07 | 4.62E-05 | 6.666585517 |
| Aplp2 | -0.543965355 | 1.468166981 | -5.187776306 | 3.34E-07 | 5.84E-05 | 6.300991397 |
| Cd63 | -0.540293425 | 2.636781081 | -7.208430695 | 2.72E-12 | 1.21E-09 | 17.43003944 |
| Abcg1 | -0.52707727 | 1.453908634 | -5.24951501 | 2.45E-07 | 4.64E-05 | 6.595383258 |
| Lilrb4 | -0.515450695 | 0.778454653 | -5.138531224 | 4.28E-07 | 7.22E-05 | 6.068357134 |
| Plin2 | -0.513566407 | 2.091392357 | -4.519298679 | 8.11E-06 | 0.000991658 | 3.311499377 |
| Apoe | -0.512223673 | 5.305230968 | -6.520024692 | 2.06E-10 | 5.91E-08 | 13.30523952 |
| Tpi1 | -0.501702423 | 1.546788346 | -5.009528535 | 8.10E-07 | 0.000120062 | 5.468190115 |
| Hpse | -0.473511692 | 0.756792141 | -5.015013535 | 7.89E-07 | 0.000120062 | 5.493434804 |
| Atp6v0d2 | -0.471270302 | 0.400574779 | -5.280952005 | 2.08E-07 | 4.43E-05 | 6.746451944 |
| Ctsl | -0.470461096 | 3.470270544 | -6.087395308 | 2.62E-09 | 7.13E-07 | 10.88438654 |
| Colec12 | -0.445624699 | 0.52776175 | -5.247741285 | 2.47E-07 | 4.64E-05 | 6.586883136 |
| Gpx3 | -0.419800306 | 1.583128179 | -4.276920222 | 2.36E-05 | 0.002304364 | 2.319217908 |
| Dab2 | -0.416600398 | 1.157748398 | -4.313670522 | 2.01E-05 | 0.002137499 | 2.466471766 |
| Ftl1 | -0.411549557 | 4.08654463 | -8.278227438 | 1.75E-15 | 1.22E-12 | 24.45678016 |
| Pld3 | -0.410445848 | 2.494165972 | -3.994182042 | 7.69E-05 | 0.006368558 | 1.22499169 |
| Tubb5 | -0.396996987 | 2.140599995 | -4.212215917 | 3.11E-05 | 0.002761883 | 2.062753404 |
| Lpl | -0.369126777 | 2.320558657 | -3.078278716 | 0.002220885 | 0.073351717 | -1.83827859 |
| Ccl9 | -0.366352259 | 1.123546225 | -3.219724572 | 0.001384549 | 0.06098251 | -1.414159479 |
| Clec7a | -0.338807194 | 0.886647334 | -3.415684062 | 0.000699258 | 0.037159472 | -0.796748459 |
| BC021767 | -0.31531504 | 0.267103482 | -5.542127375 | 5.34E-08 | 1.30E-05 | 8.031657972 |
| Lair1 | -0.313982615 | 1.08646265 | -3.239606902 | 0.001293783 | 0.059205299 | -1.353091719 |
| Ms4a6c | -0.307070945 | 0.31888324 | -5.453679017 | 8.51E-08 | 1.98E-05 | 7.590423212 |
| Serpinb6a | -0.302668204 | 0.323038315 | -5.108010915 | 4.98E-07 | 8.12E-05 | 5.925155041 |
| Mcm6 | -0.302000912 | 0.598902412 | -3.002049568 | 0.002844554 | 0.084338861 | -2.059311924 |
| Itgax | -0.287291863 | 0.404562722 | -3.705116216 | 0.000240054 | 0.017009013 | 0.177906859 |
| Gla | -0.28589498 | 0.384666341 | -4.390992852 | 1.44E-05 | 0.001632037 | 2.780037385 |
| Rpa1 | -0.269165483 | 0.449746151 | -3.468969753 | 0.000577417 | 0.032448167 | -0.62290582 |
| Rasal3 | 0.257519776 | 0.545372657 | 3.224724562 | 0.001361184 | 0.060498452 | -1.39883581 |
| Cybasc3 | 0.270468747 | 0.603124284 | 2.934627562 | 0.003525874 | 0.091691474 | -2.250390239 |
| Gpr34 | 0.288810561 | 2.190802701 | 3.083787121 | 0.002181094 | 0.073036764 | -1.822101552 |
| Eif3b | 0.291562703 | 0.605902636 | 3.421288104 | 0.000685397 | 0.036823161 | -0.778584884 |
| Egr1 | 0.292412627 | 0.292488658 | 4.296337292 | 2.17E-05 | 0.0022078 | 2.396876506 |
| Gcn1l1 | 0.296656896 | 0.416978598 | 3.887635517 | 0.000117914 | 0.009150476 | 0.830555814 |
| Abcc3 | 0.297437374 | 0.732243285 | 3.185852542 | 0.001552866 | 0.061895171 | -1.517373501 |
| Asph | 0.300089656 | 1.333205868 | 3.185027736 | 0.001557191 | 0.061895171 | -1.519873889 |
| Il6ra | 0.304367192 | 1.149224739 | 3.147359739 | 0.001766995 | 0.063991408 | -1.633407627 |

| | | | | | | |
|---|---|---|---|---|---|---|
| Slc38a10 | 0.314257552 | 0.745389831 | 3.593443668 | 0.000365716 | 0.023947256 | -0.20695569 |
| Selplg | 0.315650415 | 0.80564386 | 3.450037633 | 0.000618233 | 0.033898145 | -0.684961674 |
| Dnm2 | 0.325132736 | 0.864250797 | 3.552981676 | 0.000424864 | 0.025978542 | -0.343677078 |
| Fscn1 | 0.330096877 | 0.462152084 | 3.933751086 | 9.81E-05 | 0.007861539 | 1.000064342 |
| Slc2a5 | 0.336235204 | 0.378027422 | 4.236419041 | 2.80E-05 | 0.002647285 | 2.158267423 |
| Ptgs1 | 0.340222961 | 0.979750491 | 3.152155418 | 0.001738906 | 0.063444125 | -1.619024544 |
| Itgam | 0.36286896 | 1.060263759 | 3.787126266 | 0.000175025 | 0.012917927 | 0.467543301 |
| Fgd2 | 0.368882751 | 1.061877337 | 3.510807075 | 0.000495981 | 0.029100472 | -0.484637379 |
| Slc7a8 | 0.373553333 | 1.17904674 | 4.028572151 | 6.68E-05 | 0.005729686 | 1.354407377 |
| Pisd-ps3 | 0.404645414 | 0.539552307 | 4.622200081 | 5.08E-06 | 0.000671819 | 3.747712718 |
| Cd164 | 0.421253674 | 2.045273129 | 4.536402265 | 7.51E-06 | 0.000941635 | 3.383390266 |
| Junb | 0.422622703 | 0.487646563 | 5.326677031 | 1.65E-07 | 3.66E-05 | 6.967580433 |
| Entpd1 | 0.444704814 | 1.394168716 | 4.555283557 | 6.90E-06 | 0.000887337 | 3.463037846 |
| Slco2b1 | 0.500454682 | 1.162032082 | 4.396103857 | 1.40E-05 | 0.001632037 | 2.800942486 |
| Tmem119 | 0.555800912 | 0.774176841 | 4.928505548 | 1.20E-06 | 0.000172832 | 5.09812649 |
| P2ry12 | 0.818497132 | 2.088348937 | 6.988105987 | 1.12E-11 | 3.66E-09 | 16.07451894 |
| Siglech | 0.881439375 | 1.4517903 | 8.691842295 | 8.47E-17 | 8.29E-14 | 27.35889957 |
