## supplementary tables for "MGPfact^XMBD^: A Model-Based Factorization Method for scRNA Data Unveils Bifurcating Transcriptional Modules Underlying Cell Fate Determination": supplementary_table_6_rsqured.pdf

**Supplementary Table 6: Comparison of the explanatory power for CD8+ T cell fate for MGPfact and three other different methods. Adjusted R-squared values and P-values based on F-tests demonstrate the relative performance of MGPfact, Monocle 2, Monocle 3, and scFates Tree in fitting the experimentally characterized and annotated CD8+ T cell subtypes.**

| | | **MGPfact** | | | **Monocle 2** | | | **Monocle 3** | | | **scFates Tree** | | |
|---|---|---|---|---|---|---|---|---|---|---|---|---|---|
| | | **Adjust R-squared** | **P-value** | **F** | **Adjust R-squared** | **P-value** | **F** | **Adjust R-squared** | **P-value** | **F** | **Adjust R-squared** | **P-value** | **F** |
| NSCLC (GSE99254) | CD8-LEF1 | **0.935** | 0.000 | 80.657 | 0.176 | 0.000 | 4.007 | 0.089 | 0.08 | 3.3466 | 0.902 | 0.000 | 227.898 |
| | CD8-CD28 | **0.195** | 0.002 | 3.906 | 0.170 | 0.000 | 3.887 | 0.108 | 0.06 | 3.9062 | 0.006 | 0.145 | 2.142 |
| | CD8-CX3CR1 | 0.634 | 0.000 | 10.581 | 0.259 | 0.000 | 5.924 | 0.629 | 0.000 | 41.636 | **0.882** | 0.000 | 246.703 |
| | CD8-GZMK | 0.259 | 0.000 | 4.597 | 0.189 | 0.000 | 8.386 | **0.855** | 0.000 | 15.202 | 0.547 | 0.000 | 40.657 |
| | CD8-ZNF683 | 0.232 | 0.001 | 4.105 | 0.051 | 0.043 | 2.375 | **0.625** | 0.003 | 5.001 | 0.039 | 0.003 | 8.89 |
| | CD8-LAYN | 0.435 | 0.000 | 5.262 | 0.031 | 0.027 | 5.008 | 0.503 | 0.018 | 3.432 | **0.523** | 0.000 | 37.032 |
| CRC (GSE108989) | CD8-LEF1 | 0.311 | 0.000 | 4.021 | 0.027 | 0.036 | 4.479 | 0.461 | 0.007 | 4.767 | 0.99 | 0.000 | 1591.340 |
| | CD8-GPR183 | 0.380 | 0.000 | 5.105 | 0.032 | 0.025 | 5.124 | **0.474** | 0.006 | 4.958 | 0.139 | 0.0001 | 3.677 |
| | CD8-CX3CR1 | 0.648 | 0.000 | 13.326 | 0.047 | 0.008 | 7.161 | 0.454 | 0.007 | 4.67 | **0.817** | 0.000 | 74.781 |
| | CD8-GZMK | 0.130 | 0.013 | 2.628 | 0.550 | 0.000 | 18.106 | **0.855** | 0.000 | 26.993 | 0.236 | 0.000 | 62.443 |
| | CD8-CD6 | 0.277 | 0.000 | 5.158 | 0.109 | 0.007 | 2.716 | **0.45** | 0.008 | 4.605 | 0.054 | 0.0006 | 12.300 |
| | CD8-CD160 | 0.124 | 0.016 | 2.544 | 0.080 | 0.025 | 2.224 | **0.856** | 0.000 | 27.249 | 0.707 | 0.000 | 35.246 |
| | CD8-LAYN | **0.741** | 0.000 | 20.148 | 0.172 | 0.000 | 3.916 | 0.373 | 0.021 | 3.621 | 0.505 | 0.000 | 19.489 |
