## supplementary tables for "MGPfact^XMBD^: A Model-Based Factorization Method for scRNA Data Unveils Bifurcating Transcriptional Modules Underlying Cell Fate Determination": supplementary_table_7_gse99254_gene_weight_0.05_nonabs.pdf

**Supplementary Table 7: The highly weighted genes associated with the CD8+ T cell independent bifurcating processes in NSCLC (absolute gene weight > 0.05).**

| Trajectory 1 | | Trajectory 2 | | Trajectory 3 | |
|---|---|---|---|---|---|
| Gene | Weight | Gene | Weight | Gene | Weight |
| SELL | 0.12277783 | RGS1 | 0.10252922 | FOS | 0.13679729 |
| IL7R | 0.11912339 | GPR183 | 0.05864842 | NR4A2 | 0.13132333 |
| CCR7 | 0.09793922 | GZMK | 0.05424001 | FOSB | 0.12545785 |
| LEF1 | 0.09211993 | S100A10 | -0.05023594 | ITGA1 | 0.12353989 |
| RIPOR2 | 0.07347148 | A2M | -0.05031572 | RGS1 | 0.10643883 |
| S1PR1 | 0.06906865 | RAD9A | -0.05042131 | PTGER4 | 0.10598461 |
| TCF7 | 0.06905916 | PSMA2 | -0.05055453 | CD69 | 0.10513546 |
| NOSIP | 0.06607784 | MTSS1 | -0.05069126 | TNFAIP3 | 0.1044454 |
| PIM2 | 0.0651974 | STK38 | -0.05084639 | BTG2 | 0.10351572 |
| LDHB | 0.06424098 | TBX21 | -0.0508471 | NR4A1 | 0.10237259 |
| PLAC8 | 0.06344933 | ITGB1 | -0.05097499 | CAPG | 0.10229029 |
| TXK | 0.06301315 | LIMD2 | -0.05102802 | ZNF331 | 0.09898785 |
| DGKA | 0.06100156 | RPL5 | -0.05114202 | GPR15 | 0.09848856 |
| SERINC5 | 0.06078206 | CD47 | -0.05134788 | ITGAE | 0.0975589 |
| MAL | 0.0598235 | CD52 | -0.05152893 | RGS2 | 0.09726967 |
| FCMR | 0.05973351 | PSMB8 | -0.05156328 | PPP1R15A | 0.09416101 |
| LRRC75A-AS1 | 0.0570353 | CYTH1 | -0.05159705 | NR4A3 | 0.0939202 |
| BIRC3 | 0.05598137 | GZMA | -0.05163451 | CREM | 0.09334473 |
| MYC | 0.05575529 | CDK2AP2 | -0.05165054 | JAML | 0.09246488 |
| LDLRAP1 | 0.05520815 | GTF3A | -0.05174336 | MYADM | 0.09104999 |
| THEM4 | 0.05449355 | ITGAL | -0.0524397 | ZFP36 | 0.08752144 |
| TMEM123 | 0.05372218 | ATP5G3 | -0.05275551 | CD160 | 0.08633946 |
| RPS5 | 0.05348627 | ARPC2 | -0.05275686 | PELO | 0.08470251 |
| ADD3 | 0.0534615 | SUB1 | -0.05308477 | CSRNP1 | 0.08295651 |
| EEF1G | 0.05267825 | EIF3L | -0.05310938 | PFKFB3 | 0.0826525 |
| ICAM2 | 0.05238508 | C1orf21 | -0.053331 | ANKRD28 | 0.08097853 |
| EEF2 | 0.05125448 | ITGB7 | -0.05344836 | KLRC2 | 0.08084336 |
| RACK1 | 0.0506414 | SYNE1 | -0.05381151 | YPEL5 | 0.08037651 |
| RPLP0 | 0.05007907 | RASSF1 | -0.05412827 | TMIGD2 | 0.07796671 |
| CAPG | -0.05090886 | LILRB1 | -0.05413067 | ITM2C | 0.0777218 |
| HLA-DQB1 | -0.05111884 | FTL | -0.05418794 | PTPN22 | 0.07751772 |
| OASL | -0.0511849 | TTC38 | -0.0543726 | LDLRAD4 | 0.07673066 |
| LYST | -0.05126527 | IDH2 | -0.05438132 | FAM46C | 0.07652411 |
| KLRC3 | -0.0513786 | NACA | -0.05480981 | SAMSN1 | 0.0762235 |
| TNIP3 | -0.05281256 | HLA-DPB1 | -0.05510259 | KLRC3 | 0.07531079 |
| HAVCR2 | -0.05284193 | ABI3 | -0.05598021 | MCL1 | 0.07379315 |
| CHST12 | -0.0531795 | ADRB2 | -0.05633323 | JUN | 0.07350829 |
| CCL3 | -0.0536317 | ATP5B | -0.05637771 | RANBP2 | 0.07240191 |
| CD74 | -0.05371019 | CD8B | -0.05670939 | PER1 | 0.07125798 |
| IL2RB | -0.05381337 | ID2 | -0.05706682 | FOSL2 | 0.07103898 |
| PELO | -0.05383944 | BTF3 | -0.05741629 | GPR65 | 0.07028736 |
| F2R | -0.05480732 | MYO1G | -0.05753213 | JUNB | 0.07025593 |
| KLRC2 | -0.05500017 | ARL6IP1 | -0.05764815 | DNAJA1 | 0.0697703 |
| HLA-DQA1 | -0.05572369 | CD8A | -0.05806677 | EGR1 | 0.06876391 |
| CD69 | -0.05597412 | ADD3 | -0.05813834 | NFKBIZ | 0.06809996 |
| ALOX5AP | -0.05638912 | RHOA | -0.0583966 | HOPX | 0.06807426 |
| KLRC1 | -0.05665359 | PATL2 | -0.05865958 | SYTL3 | 0.06804121 |
| CTSD | -0.05680028 | KLF2 | -0.05894567 | SDCBP | 0.06710357 |
| ADGRE5 | -0.058539 | TARP | -0.05912359 | ABCB1 | 0.06709273 |
| TBCD | -0.05880677 | SPON2 | -0.05960364 | CD55 | 0.06683529 |
| APOBEC3C | -0.05985941 | SLAMF7 | -0.05960885 | RASGEF1B | 0.06626586 |
| ID2 | -0.06250169 | SPN | -0.05962565 | CLDND1 | 0.06606434 |
| CRTAM | -0.06339606 | PTP4A2 | -0.0596508 | SERTAD1 | 0.06589464 |
| ITM2C | -0.06478816 | CCND3 | -0.06007424 | MAP3K8 | 0.06500334 |
| PTPN22 | -0.06575952 | SLC9A3R1 | -0.06051653 | CD96 | 0.06382304 |
| ANAPC1P1 | -0.06642552 | EMP3 | -0.0625229 | LOC100130476 | 0.06330417 |
| ABI3 | -0.0666702 | MYO1F | -0.06400365 | NFKBIA | 0.06296314 |
| LAG3 | -0.06888376 | FCGR3B | -0.06415346 | LINC-PINT | 0.06295145 |
| FASLG | -0.06962184 | SAMD3 | -0.06424079 | KLRC1 | 0.06248458 |
| HLA-DPB1 | -0.06969182 | ACTR3 | -0.06426521 | ID2 | 0.06212784 |
| HLA-DRA | -0.07062932 | ITGB2 | -0.06446334 | TIPARP | 0.06197539 |
| SLAMF7 | -0.07164892 | CTSC | -0.06473376 | GPR171 | 0.06197189 |
| CD63 | -0.07285543 | ARPC4 | -0.06531693 | REL | 0.06163944 |
| RGS1 | -0.07428344 | TGFBR3 | -0.06548807 | DNAJB1 | 0.06097907 |
| HLA-DPA1 | -0.07472285 | TPST2 | -0.06719822 | SIK1 | 0.0606986 |
| HLA-DRB6 | -0.08082488 | EFHD2 | -0.06747864 | DUSP2 | 0.06001204 |
| ITGAE | -0.08414212 | PPP1CA | -0.06761421 | CXCR4 | 0.05958199 |
| GZMK | -0.0856226 | APMAP | -0.06838524 | PDE4B | 0.05942223 |

| | | | | | |
|---|---|---|---|---|---|
| HLA-DRB5 | -0.08623641 | CST7 | -0.07016853 | LOC284454 | 0.05873695 |
| CXCR6 | -0.08667095 | AES | -0.07019411 | TSPYL2 | 0.05853272 |
| ZNF683 | -0.0867767 | ZEB2 | -0.07042714 | SKIL | 0.05826068 |
| ITGA1 | -0.08731439 | S1PR5 | -0.07093995 | SLA2 | 0.05773461 |
| TARP | -0.08937704 | RIPOR2 | -0.07116137 | CD8A | 0.05750839 |
| HLA-DRB1 | -0.09215427 | RAP1B | -0.07210229 | SYTL2 | 0.05696918 |
| CST7 | -0.09650377 | PRSS23 | -0.07247075 | SPRY1 | 0.05661457 |
| AOAH | -0.09767995 | BIN2 | -0.07701697 | IL18RAP | 0.05642521 |
| APOBEC3G | -0.10073416 | PXN | -0.07829418 | DDX3X | 0.05600051 |
| CTSW | -0.1009556 | FGR | -0.07919277 | CCL4 | 0.05574412 |
| PRF1 | -0.10131273 | C12orf75 | -0.08008906 | CLK1 | 0.05520014 |
| KLRD1 | -0.10222589 | FLNA | -0.08066581 | IVNS1ABP | 0.05506366 |
| CCL4L1 | -0.10545153 | KLRG1 | -0.08168838 | BRE-AS1 | 0.05495641 |
| CD8B | -0.12908598 | CTSW | -0.08392219 | PRMT9 | 0.05474413 |
| GZMH | -0.12958302 | PLAC8 | -0.08483126 | ZFAND5 | 0.05411584 |
| GZMB | -0.1402741 | S1PR1 | -0.08785517 | AOAH | 0.05383833 |
| GZMA | -0.14051908 | FCRL6 | -0.09099204 | PDE4D | 0.05358349 |
| CCL4 | -0.15068753 | UCP2 | -0.09314049 | CD7 | 0.05314107 |
| NKG7 | -0.15696927 | NKG7 | -0.10058139 | CCL5 | 0.05285404 |
| CCL5 | -0.16530581 | GZMB | -0.10386484 | PDCD4 | 0.05233882 |
| CD8A | -0.19519364 | PRF1 | -0.10731906 | ABLIM1 | 0.05192225 |
| | | ADGRG1 | -0.10772137 | FAM53C | 0.05103623 |
| | | PLEK | -0.10848652 | SLC2A3 | 0.05101168 |
| | | GZMH | -0.11502765 | PDE4A | 0.05076777 |
| | | FCGR3A | -0.11548208 | G3BP2 | 0.05030398 |
| | | LITAF | -0.11569373 | ARHGAP9 | 0.05022487 |
| | | CX3CR1 | -0.1166124 | PATL2 | -0.05008036 |
| | | GNLY | -0.11747741 | TMEM173 | -0.05056077 |
| | | FGFBP2 | -0.12352172 | CD27-AS1 | -0.050661 |
| | | KLRD1 | -0.12958919 | SH2D1A | -0.05181995 |
| | | | | GIMAP7 | -0.05270029 |
| | | | | SYNE1 | -0.05287288 |
| | | | | SELL | -0.05289495 |
| | | | | ADGRG1 | -0.05325115 |
| | | | | HLA-DPB1 | -0.05371068 |
| | | | | ITGB2-AS1 | -0.05410236 |
| | | | | FCGR3A | -0.05415227 |
| | | | | ADD3 | -0.05545101 |
| | | | | CD27 | -0.0554842 |
| | | | | UCP2 | -0.05890536 |
| | | | | PXN | -0.05933983 |
| | | | | A2M | -0.06128179 |
| | | | | FGFBP2 | -0.06251175 |
| | | | | GIMAP4 | -0.06313162 |
| | | | | S1PR1 | -0.06422009 |
| | | | | ITGB1 | -0.06667606 |
| | | | | LINC00861 | -0.06713255 |
| | | | | CX3CR1 | -0.06773757 |
| | | | | GZMH | -0.06837104 |
| | | | | SAMD3 | -0.06903684 |
| | | | | RIPOR2 | -0.07203213 |
| | | | | KLRG1 | -0.07412077 |
| | | | | PLEK | -0.0748601 |
| | | | | ITGB2 | -0.11604065 |
