## supplementary tables for "MGPfact^XMBD^: A Model-Based Factorization Method for scRNA Data Unveils Bifurcating Transcriptional Modules Underlying Cell Fate Determination": supplementary_table_8_gse108989_gene_weight_0.05_nonabs.pdf

**Supplementary Table 8: The highly weighted genes associated with the CD8+ T cell independent bifurcating processes in CRC (absolute gene weight > 0.05).**

| Trajectory 1 | | Trajectory 2 | | Trajectory 3 | |
|---|---|---|---|---|---|
| Gene | Weight | Gene | Weight | Gene | Weight |
| IL7R | 0.13298435 | CD8A | 0.150626 | FOS | 0.13679729 |
| LTB | 0.10841824 | TARP | 0.13528138 | NR4A2 | 0.13132333 |
| SELL | 0.09279064 | KLRD1 | 0.12232609 | FOSB | 0.12545785 |
| GPR183 | 0.0916535 | FCRL6 | 0.11964697 | ITGA1 | 0.12353989 |
| CCR7 | 0.08903049 | CCL5 | 0.11263488 | RGS1 | 0.10643883 |
| TMEM123 | 0.07793223 | NKG7 | 0.10874077 | PTGER4 | 0.10598461 |
| LEF1 | 0.07302568 | AOAH | 0.10866534 | CD69 | 0.10513546 |
| CD28 | 0.06836185 | SLAMF7 | 0.1082855 | TNFAIP3 | 0.1044454 |
| TCF7 | 0.06605279 | PLEK | 0.10488948 | BTG2 | 0.10351572 |
| FCMR | 0.0624517 | CD160 | 0.10063399 | NR4A1 | 0.10237259 |
| DGKA | 0.06209366 | FCGR3A | 0.09870179 | CAPG | 0.10229029 |
| SERINC5 | 0.05320843 | CTSW | 0.0899321 | ZNF331 | 0.09898785 |
| LDLRAP1 | 0.05281235 | HOPX | 0.08901752 | GPR15 | 0.09848856 |
| PIK3IP1 | 0.05023896 | CX3CR1 | 0.08767622 | ITGAE | 0.0975589 |
| PTPN22 | -0.05001684 | FGFBP2 | 0.08549309 | RGS2 | 0.09726967 |
| ARAP2 | -0.05023222 | STOM | 0.07969273 | PPP1R15A | 0.09416101 |
| APMAP | -0.05043737 | FGR | 0.0788306 | NR4A3 | 0.0939202 |
| THEMIS | -0.0504602 | KLRG1 | 0.07770192 | CREM | 0.09334473 |
| KLRG1 | -0.05079659 | PTGER2 | 0.07316796 | JAML | 0.09246488 |
| GABARAPL1 | -0.05092233 | S1PR5 | 0.06943705 | MYADM | 0.09104999 |
| TNFSF4 | -0.05113945 | MBP | 0.06903565 | ZFP36 | 0.08752144 |
| RAB27A | -0.05142474 | SCML4 | 0.06865519 | CD160 | 0.08633946 |
| CCL3L1 | -0.05149744 | A2M | 0.06505644 | PELO | 0.08470251 |
| KLRC4 | -0.0527624 | KLRC3 | 0.06500614 | CSRNP1 | 0.08295651 |
| MIR155HG | -0.05284935 | KLRC2 | 0.06272967 | PFKFB3 | 0.0826525 |
| CCL3L3 | -0.05428937 | ZEB2 | 0.06175142 | ANKRD28 | 0.08097853 |
| ADGRG5 | -0.05447833 | PLAC8 | 0.06159511 | KLRC2 | 0.08084336 |
| ITM2C | -0.05448603 | PRF1 | 0.06159372 | YPEL5 | 0.08037651 |
| TNIP3 | -0.05514648 | PATL2 | 0.06151737 | TMIGD2 | 0.07796671 |
| FCRL6 | -0.05568229 | TGFBR3 | 0.06106651 | ITM2C | 0.0777218 |
| CD244 | -0.05578783 | CD244 | 0.06088757 | PTPN22 | 0.07751772 |
| BHLHE40 | -0.05606846 | MYO1F | 0.06062513 | LDLRAD4 | 0.07673066 |
| PDCD1 | -0.05632214 | GPR15 | 0.06045013 | FAM46C | 0.07652411 |
| RBPJ | -0.0574984 | LYAR | 0.06042588 | SAMSN1 | 0.0762235 |
| CXCR6 | -0.05758775 | GZMA | 0.06027249 | KLRC3 | 0.07531079 |
| DUSP4 | -0.05771099 | CST7 | 0.05973188 | MCL1 | 0.07379315 |
| PLEK | -0.05773746 | ANXA1 | 0.05962489 | JUN | 0.07350829 |
| CTSC | -0.05821452 | ABI3 | 0.05903792 | RANBP2 | 0.07240191 |
| ID2 | -0.05825088 | XCL2 | 0.05620558 | PER1 | 0.07125798 |
| CD74 | -0.05893733 | BIN2 | 0.05586858 | FOSL2 | 0.07103898 |
| ITGA1 | -0.05943219 | S1PR1 | 0.0542717 | GPR65 | 0.07028736 |
| ANXA5 | -0.05972505 | ADGRG1 | 0.05425219 | JUNB | 0.07025593 |
| F2R | -0.06015661 | PTGDR | 0.05273247 | DNAJA1 | 0.0697703 |
| IDH2 | -0.0605085 | GPR65 | 0.0524179 | EGR1 | 0.06876391 |
| CLEC2B | -0.06083732 | ADRB2 | 0.05236872 | NFKBIZ | 0.06809996 |
| SH2D1A | -0.06119614 | CD300A | 0.05218803 | HOPX | 0.06807426 |
| DTHD1 | -0.06120573 | RASSF1 | 0.05123601 | SYTL3 | 0.06804121 |
| ABI3 | -0.06124065 | TTC38 | 0.05095901 | SDCBP | 0.06710357 |
| SAMD3 | -0.06485483 | FCGR3B | 0.05032246 | ABCB1 | 0.06709273 |
| CHST12 | -0.06529378 | INPP4B | -0.05006784 | CD55 | 0.06683529 |
| ZNF683 | -0.06621011 | SNX9 | -0.05061957 | RASGEF1B | 0.06626586 |
| HLA-DMA | -0.06739139 | PLP2 | -0.05117104 | CLDND1 | 0.06606434 |
| EOMES | -0.06740262 | TFRC | -0.05124327 | SERTAD1 | 0.06589464 |
| ADGRG1 | -0.06774398 | RNF19A | -0.05128149 | MAP3K8 | 0.06500334 |
| CD63 | -0.06860755 | HLA-DQA1 | -0.05190809 | CD96 | 0.06382304 |
| GNLY | -0.07107329 | SMC4 | -0.05208698 | LOC100130476 | 0.06330417 |
| CRTAM | -0.07285167 | TOX | -0.05275491 | NFKBIA | 0.06296314 |
| ZEB2 | -0.07297214 | HIF1A | -0.05282856 | LINC-PINT | 0.06295145 |
| LYST | -0.07620944 | PELI1 | -0.05305456 | KLRC1 | 0.06248458 |
| LAG3 | -0.07656486 | TRAT1 | -0.05323519 | ID2 | 0.06212784 |
| C12orf75 | -0.07662767 | PAG1 | -0.05475959 | TIPARP | 0.06197539 |
| VCAM1 | -0.07703371 | TNFSF10 | -0.05503417 | GPR171 | 0.06197189 |
| ITGAE | -0.07756817 | FKBP1A | -0.05539432 | REL | 0.06163944 |
| AOAH | -0.07898583 | GALM | -0.05589634 | DNAJB1 | 0.06097907 |
| FASLG | -0.08028904 | ENTPD1 | -0.05594273 | SIK1 | 0.0606986 |
| OASL | -0.08148413 | MYO7A | -0.05633295 | DUSP2 | 0.06001204 |
| APOBEC3C | -0.08225132 | NAMPT | -0.0565777 | CXCR4 | 0.05958199 |
| HLA-DQB1 | -0.08419791 | TANK | -0.05720541 | PDE4B | 0.05942223 |

| | | | | | |
|---|---|---|---|---|---|
| HLA-DQA1 | -0.08430235 | LGALS3 | -0.05755052 | LOC284454 | 0.05873695 |
| SLAMF7 | -0.08498413 | TNFSF4 | -0.05831843 | TSPYL2 | 0.05853272 |
| TARP | -0.08528143 | RGCC | -0.05951019 | SKIL | 0.05826068 |
| IFNG | -0.08556293 | TIAM1 | -0.05986704 | SLA2 | 0.05773461 |
| CXCL13 | -0.08656011 | MIR155HG | -0.06069517 | CD8A | 0.05750839 |
| CCL3 | -0.0887134 | PDE4DIP | -0.06160095 | SYTL2 | 0.05696918 |
| GZMK | -0.09127376 | BATF | -0.06270309 | SPRY1 | 0.05661457 |
| HLA-DPA1 | -0.09262842 | ITM2A | -0.06473874 | IL18RAP | 0.05642521 |
| HLA-DPB1 | -0.09348449 | IL6ST | -0.06556109 | DDX3X | 0.05600051 |
| HAVCR2 | -0.09510836 | ACP5 | -0.06571955 | CCL4 | 0.05574412 |
| CST7 | -0.10083295 | CD28 | -0.06624501 | CLK1 | 0.05520014 |
| CCL4L1 | -0.10744835 | BIRC3 | -0.06766238 | IVNS1ABP | 0.05506366 |
| KLRD1 | -0.11023813 | VCAM1 | -0.06916228 | BRE-AS1 | 0.05495641 |
| CTSW | -0.1164164 | NAB1 | -0.07021131 | PRMT9 | 0.05474413 |
| HLA-DRB6 | -0.11823782 | SIRPG | -0.0703857 | ZFAND5 | 0.05411584 |
| APOBEC3G | -0.11856668 | LYST | -0.07069774 | AOAH | 0.05383833 |
| HLA-DRB5 | -0.11924546 | TNFRSF18 | -0.07208177 | PDE4D | 0.05358349 |
| PRF1 | -0.12468491 | HLA-DRA | -0.07239457 | CD7 | 0.05314107 |
| HLA-DRA | -0.12548464 | ARID5B | -0.07292086 | CCL5 | 0.05285404 |
| HLA-DRB1 | -0.12878604 | SAMSN1 | -0.07413002 | PDCD4 | 0.05233882 |
| CCL4 | -0.14152013 | NDFIP2 | -0.07529167 | ABLIM1 | 0.05192225 |
| CCL5 | -0.14669454 | HNRNPLL | -0.07548146 | FAM53C | 0.05103623 |
| GZMA | -0.14835982 | CXCR6 | -0.07562927 | SLC2A3 | 0.05101168 |
| CD8A | -0.16080035 | PHLDA1 | -0.08056112 | PDE4A | 0.05076777 |
| NKG7 | -0.16329636 | ADAM19 | -0.08074802 | G3BP2 | 0.05030398 |
| GZMB | -0.16599309 | TNFRSF9 | -0.08337827 | ARHGAP9 | 0.05022487 |
| GZMH | -0.1684426 | RGS1 | -0.08418174 | PATL2 | -0.05008036 |
| | | TBC1D4 | -0.09389029 | TMEM173 | -0.05056077 |
| | | DUSP4 | -0.10723249 | CD27-AS1 | -0.050661 |
| | | ICOS | -0.10791782 | SH2D1A | -0.05181995 |
| | | RBPJ | -0.10849932 | GIMAP7 | -0.05270029 |
| | | CXCL13 | -0.10995167 | SYNE1 | -0.05287288 |
| | | PDCD1 | -0.11044705 | SELL | -0.05289495 |
| | | CTLA4 | -0.11091745 | ADGRG1 | -0.05325115 |
| | | HAVCR2 | -0.11456914 | HLA-DPB1 | -0.05371068 |
| | | CD27-AS1 | -0.11584246 | ITGB2-AS1 | -0.05410236 |
| | | CD27 | -0.12559839 | FCGR3A | -0.05415227 |
| | | CD82 | -0.14858959 | ADD3 | -0.05545101 |
| | | | | CD27 | -0.0554842 |
| | | | | UCP2 | -0.05890536 |
| | | | | PXN | -0.05933983 |
| | | | | A2M | -0.06128179 |
| | | | | FGFBP2 | -0.06251175 |
| | | | | GIMAP4 | -0.06313162 |
| | | | | S1PR1 | -0.06422009 |
| | | | | ITGB1 | -0.06667606 |
| | | | | LINC00861 | -0.06713255 |
| | | | | CX3CR1 | -0.06773757 |
| | | | | GZMH | -0.06837104 |
| | | | | SAMD3 | -0.06903684 |
| | | | | RIPOR2 | -0.07203213 |
| | | | | KLRG1 | -0.07412077 |
| | | | | PLEK | -0.0748601 |
| | | | | ITGB2 | -0.11604065 |
