## supplementary tables for "MGPfact^XMBD^: A Model-Based Factorization Method for scRNA Data Unveils Bifurcating Transcriptional Modules Underlying Cell Fate Determination": supplementary_table_9_gse99254_enrich_go_combine.pdf

**Supplementary Table 9: GO Enrichment Analysis Results for Highly Weighted Genes Associated with Trajectory 1 of NSCLC (Benjamini–Hochberg-adjusted P value < 0.05)**

| ID | Description | GeneRatio | BgRatio | pvalue | p.adjust | qvalue | geneID | Count | ONTOLOGY |
|---|---|---|---|---|---|---|---|---|---|
| GO:0019886 | antigen processing and presentation of exogenous peptide antigen via MHC class II | 9/83 | 30/18723 | 5.61105E-15 | 7.60587E-12 | 5.50863E-12 | HLA-DQB1/CD74/HLA-DQA1/CTSD/HLA-DPB1/HLA-DRA/HLA-DPA1/HLA-DRB5/HLA-DRB1 | 9 | BP |
| GO:0001909 | leukocyte mediated cytotoxicity | 13/83 | 124/18723 | 8.86982E-15 | 7.60587E-12 | 5.50863E-12 | IL7R/LYST/HAVCR2/KLRC2/KLRC1/CRTAM/LAG3/HLA-DRA/SLAMF7/HLA-DRB1/PRF1/KLRD1/GZMB | 13 | BP |
| GO:0002495 | antigen processing and presentation of peptide antigen via MHC class II | 9/83 | 34/18723 | 2.02793E-14 | 1.1593E-11 | 8.39633E-12 | HLA-DQB1/CD74/HLA-DQA1/CTSD/HLA-DPB1/HLA-DRA/HLA-DPA1/HLA-DRB5/HLA-DRB1 | 9 | BP |
| GO:0002504 | antigen processing and presentation of peptide or polysaccharide antigen via MHC class II | 9/83 | 36/18723 | 3.614E-14 | 1.5495E-11 | 1.12224E-11 | HLA-DQB1/CD74/HLA-DQA1/CTSD/HLA-DPB1/HLA-DRA/HLA-DPA1/HLA-DRB5/HLA-DRB1 | 9 | BP |
| GO:0002478 | antigen processing and presentation of exogenous peptide antigen | 9/83 | 38/18723 | 6.21326E-14 | 2.13115E-11 | 1.5435E-11 | HLA-DQB1/CD74/HLA-DQA1/CTSD/HLA-DPB1/HLA-DRA/HLA-DPA1/HLA-DRB5/HLA-DRB1 | 9 | BP |
| GO:0002399 | MHC class II protein complex assembly | 7/83 | 16/18723 | 2.87775E-13 | 7.05048E-11 | 5.10638E-11 | HLA-DQB1/HLA-DQA1/HLA-DPB1/HLA-DRA/HLA-DPA1/HLA-DRB5/HLA-DRB1 | 7 | BP |
| GO:0002503 | peptide antigen assembly with MHC class II protein complex | 7/83 | 16/18723 | 2.87775E-13 | 7.05048E-11 | 5.10638E-11 | HLA-DQB1/HLA-DQA1/HLA-DPB1/HLA-DRA/HLA-DPA1/HLA-DRB5/HLA-DRB1 | 7 | BP |
| GO:0019884 | antigen processing and presentation of exogenous antigen | 9/83 | 47/18723 | 5.02973E-13 | 1.07825E-10 | 7.80931E-11 | HLA-DQB1/CD74/HLA-DQA1/CTSD/HLA-DPB1/HLA-DRA/HLA-DPA1/HLA-DRB5/HLA-DRB1 | 9 | BP |
| GO:0050870 | positive regulation of T cell activation | 14/83 | 216/18723 | 6.35556E-13 | 1.21109E-10 | 8.77142E-11 | IL7R/CCR7/LEF1/HLA-DQB1/HAVCR2/CD74/HLA-DQA1/PTPN22/HLA-DPB1/HLA-DRA/HLA-DPA1/HLA-DRB5/HLA-DRB1/CCL5 | 14 | BP |
| GO:0002501 | peptide antigen assembly with MHC protein complex | 7/83 | 18/18723 | 7.94858E-13 | 1.36318E-10 | 9.87298E-11 | HLA-DQB1/HLA-DQA1/HLA-DPB1/HLA-DRA/HLA-DPA1/HLA-DRB5/HLA-DRB1 | 7 | BP |
| GO:0002396 | MHC protein complex assembly | 7/83 | 19/18723 | 1.25405E-12 | 1.85109E-10 | 1.34067E-10 | HLA-DQB1/HLA-DQA1/HLA-DPB1/HLA-DRA/HLA-DPA1/HLA-DRB5/HLA-DRB1 | 7 | BP |
| GO:0019882 | antigen processing and presentation | 11/83 | 106/18723 | 1.29522E-12 | 1.85109E-10 | 1.34067E-10 | CCR7/HLA-DQB1/CD74/HLA-DQA1/CTSD/HLA-DPB1/HLA-DRA/HLA-DPA1/HLA-DRB5/HLA-DRB1/CD8A | 11 | BP |
| GO:0001906 | cell killing | 13/83 | 188/18723 | 1.99003E-12 | 2.62531E-10 | 1.90141E-10 | IL7R/LYST/HAVCR2/KLRC2/KLRC1/CRTAM/LAG3/HLA-DRA/SLAMF7/HLA-DRB1/PRF1/KLRD1/GZMB | 13 | BP |
| GO:1903039 | positive regulation of leukocyte cell-cell adhesion | 14/83 | 239/18723 | 2.52358E-12 | 3.09139E-10 | 2.23897E-10 | IL7R/CCR7/LEF1/HLA-DQB1/HAVCR2/CD74/HLA-DQA1/PTPN22/HLA-DPB1/HLA-DRA/HLA-DPA1/HLA-DRB5/HLA-DRB1/CCL5 | 14 | BP |
| GO:0048002 | antigen processing and presentation of peptide antigen | 9/83 | 62/18723 | 7.09793E-12 | 8.1153E-10 | 5.87759E-10 | HLA-DQB1/CD74/HLA-DQA1/CTSD/HLA-DPB1/HLA-DRA/HLA-DPA1/HLA-DRB5/HLA-DRB1 | 9 | BP |
| GO:0042267 | natural killer cell mediated cytotoxicity | 9/83 | 68/18723 | 1.68771E-11 | 1.80901E-09 | 1.3102E-09 | LYST/HAVCR2/KLRC2/KLRC1/CRTAM/LAG3/SLAMF7/KLRD1/GZMB | 9 | BP |
| GO:0031341 | regulation of cell killing | 10/83 | 99/18723 | 1.89753E-11 | 1.91428E-09 | 1.38643E-09 | IL7R/HAVCR2/KLRC2/KLRC1/CRTAM/LAG3/HLA-DRA/HLA-DRB1/PRF1/KLRD1 | 10 | BP |
| GO:0002228 | natural killer cell mediated immunity | 9/83 | 71/18723 | 2.52338E-11 | 2.32064E-09 | 1.68074E-09 | LYST/HAVCR2/KLRC2/KLRC1/CRTAM/LAG3/SLAMF7/KLRD1/GZMB | 9 | BP |
| GO:0022409 | positive regulation of cell-cell adhesion | 14/83 | 284/18723 | 2.57097E-11 | 2.32064E-09 | 1.68074E-09 | IL7R/CCR7/LEF1/HLA-DQB1/HAVCR2/CD74/HLA-DQA1/PTPN22/HLA-DPB1/HLA-DRA/HLA-DPA1/HLA-DRB5/HLA-DRB1/CCL5 | 14 | BP |
| GO:0001910 | regulation of leukocyte mediated cytotoxicity | 9/83 | 82/18723 | 9.54724E-11 | 8.18676E-09 | 5.92934E-09 | IL7R/HAVCR2/KLRC2/KLRC1/CRTAM/LAG3/HLA-DRA/HLA-DRB1/KLRD1 | 9 | BP |
| GO:0030217 | T cell differentiation | 13/83 | 257/18723 | 1.02162E-10 | 8.34323E-09 | 6.04267E-09 | IL7R/CCR7/LEF1/TCF7/CD74/KLRC1/CRTAM/PTPN22/LAG3/HLA-DRA/ZNF683/HLA-DRB1/CD8A | 13 | BP |
| GO:1902105 | regulation of leukocyte differentiation | 12/83 | 279/18723 | 3.57954E-09 | 2.79042E-07 | 2.02099E-07 | IL7R/LEF1/TCF7/MYC/CCL3/CD74/ID2/CRTAM/LAG3/HLA-DRA/ZNF683/HLA-DRB1 | 12 | BP |
| GO:0045619 | regulation of lymphocyte differentiation | 10/83 | 174/18723 | 5.02307E-09 | 3.74546E-07 | 2.71269E-07 | IL7R/LEF1/TCF7/CD74/ID2/CRTAM/LAG3/HLA-DRA/ZNF683/HLA-DRB1 | 10 | BP |
| GO:0031343 | positive regulation of cell killing | 7/83 | 63/18723 | 1.17742E-08 | 8.41367E-07 | 6.09368E-07 | KLRC2/CRTAM/LAG3/HLA-DRA/HLA-DRB1/PRF1/KLRD1 | 7 | BP |
| GO:0045580 | regulation of T cell differentiation | 9/83 | 146/18723 | 1.67459E-08 | 1.14877E-06 | 8.32009E-07 | IL7R/LEF1/TCF7/CD74/CRTAM/LAG3/HLA-DRA/ZNF683/HLA-DRB1 | 9 | BP |
| GO:0030593 | neutrophil chemotaxis | 8/83 | 103/18723 | 1.78663E-08 | 1.17849E-06 | 8.53529E-07 | CCR7/RIPOR2/CCL3/CD74/ITGA1/CCL4L1/CCL4/CCL5 | 8 | BP |
| GO:0002381 | immunoglobulin production involved in immunoglobulin-mediated immune response | 7/83 | 70/18723 | 2.48839E-08 | 1.58059E-06 | 1.14476E-06 | HLA-DQB1/HLA-DQA1/HLA-DPB1/HLA-DRA/HLA-DPA1/HLA-DRB5/HLA-DRB1 | 7 | BP |
| GO:0032609 | interferon-gamma production | 8/83 | 112/18723 | 3.46005E-08 | 2.0462E-06 | 1.48198E-06 | CCR7/TXK/HAVCR2/CRTAM/PTPN22/HLA-DPB1/HLA-DPA1/HLA-DRB1 | 8 | BP |
| GO:0032649 | regulation of interferon-gamma production | 8/83 | 112/18723 | 3.46005E-08 | 2.0462E-06 | 1.48198E-06 | CCR7/TXK/HAVCR2/CRTAM/PTPN22/HLA-DPB1/HLA-DPA1/HLA-DRB1 | 8 | BP |
| GO:0042269 | regulation of natural killer cell mediated cytotoxicity | 6/83 | 44/18723 | 3.8973E-08 | 2.22796E-06 | 1.61362E-06 | HAVCR2/KLRC2/KLRC1/CRTAM/LAG3/KLRD1 | 6 | BP |
| GO:0070374 | positive regulation of ERK1 and ERK2 cascade | 10/83 | 217/18723 | 4.13549E-08 | 2.28786E-06 | 1.65701E-06 | CCR7/HAVCR2/CCL3/CD74/F2R/PTPN22/HLA-DRB1/CCL4L1/CCL4/CCL5 | 10 | BP |
| GO:0045088 | regulation of innate immune response | 10/83 | 218/18723 | 4.31889E-08 | 2.31465E-06 | 1.67641E-06 | TXK/BIRC3/HAVCR2/KLRC2/KLRC1/CRTAM/PTPN22/LAG3/KLRD1/CCL5 | 10 | BP |
| GO:0072676 | lymphocyte migration | 8/83 | 117/18723 | 4.87448E-08 | 2.53325E-06 | 1.83473E-06 | CCR7/RIPOR2/S1PR1/CCL3/CRTAM/CCL4L1/CCL4/CCL5 | 8 | BP |
| GO:0002706 | regulation of lymphocyte mediated immunity | 9/83 | 168/18723 | 5.66179E-08 | 2.77428E-06 | 2.0093E-06 | IL7R/HAVCR2/KLRC2/KLRC1/CRTAM/LAG3/HLA-DRA/HLA-DRB1/KLRD1 | 9 | BP |
| GO:0002833 | positive regulation of response to biotic stimulus | 9/83 | 168/18723 | 5.66179E-08 | 2.77428E-06 | 2.0093E-06 | TXK/OASL/HAVCR2/KLRC2/CRTAM/LAG3/HLA-DRB1/KLRD1/CCL5 | 9 | BP |
| GO:0002715 | regulation of natural killer cell mediated immunity | 6/83 | 48/18723 | 6.68015E-08 | 3.13385E-06 | 2.26972E-06 | HAVCR2/KLRC2/KLRC1/CRTAM/LAG3/KLRD1 | 6 | BP |
| GO:1990266 | neutrophil migration | 8/83 | 122/18723 | 6.76108E-08 | 3.13385E-06 | 2.26972E-06 | CCR7/RIPOR2/CCL3/CD74/ITGA1/CCL4L1/CCL4/CCL5 | 8 | BP |
| GO:0030595 | leukocyte chemotaxis | 10/83 | 230/18723 | 7.15016E-08 | 3.22698E-06 | 2.33717E-06 | CCR7/RIPOR2/S1PR1/LYST/CCL3/CD74/ITGA1/CCL4L1/CCL4/CCL5 | 10 | BP |
| GO:0001913 | T cell mediated cytotoxicity | 6/83 | 49/18723 | 7.58545E-08 | 3.33565E-06 | 2.41588E-06 | IL7R/KLRC1/HLA-DRA/HLA-DRB1/PRF1/KLRD1 | 6 | BP |
| GO:0071621 | granulocyte chemotaxis | 8/83 | 125/18723 | 8.17073E-08 | 3.5032E-06 | 2.53723E-06 | CCR7/RIPOR2/CCL3/CD74/ITGA1/CCL4L1/CCL4/CCL5 | 8 | BP |
| GO:0002695 | negative regulation of leukocyte activation | 9/83 | 187/18723 | 1.4192E-07 | 5.93642E-06 | 4.29951E-06 | RIPOR2/HAVCR2/CD74/ID2/CRTAM/PTPN22/LAG3/HLA-DRB1/CST7 | 9 | BP |
| GO:0001912 | positive regulation of leukocyte mediated cytotoxicity | 6/83 | 56/18723 | 1.71827E-07 | 7.01628E-06 | 5.08161E-06 | KLRC2/CRTAM/LAG3/HLA-DRA/HLA-DRB1/KLRD1 | 6 | BP |
| GO:0034341 | response to interferon-gamma | 8/83 | 141/18723 | 2.07686E-07 | 8.28329E-06 | 5.99925E-06 | TXK/CCL3/CD74/FASLG/HLA-DPA1/CCL4L1/CCL4/CCL5 | 8 | BP |
| GO:1990868 | response to chemokine | 7/83 | 97/18723 | 2.42244E-07 | 9.2322E-06 | 6.68651E-06 | CCR7/RIPOR2/CCL3/CXCR6/CCL4L1/CCL4/CCL5 | 7 | BP |
| GO:1990869 | cellular response to chemokine | 7/83 | 97/18723 | 2.42244E-07 | 9.2322E-06 | 6.68651E-06 | CCR7/RIPOR2/CCL3/CXCR6/CCL4L1/CCL4/CCL5 | 7 | BP |
| GO:0097530 | granulocyte migration | 8/83 | 148/18723 | 3.01386E-07 | 1.12365E-05 | 8.13812E-06 | CCR7/RIPOR2/CCL3/CD74/ITGA1/CCL4L1/CCL4/CCL5 | 8 | BP |

| | | | | | | | | | |
|---|---|---|---|---|---|---|---|---|---|
| GO:0050866 | negative regulation of cell activation | 9/83 | 210/18723 | 3.79483E-07 | 1.38471E-05 | 1.00289E-05 | RIPOR2/HAVCR2/CD74/ID2/CRTAM/PTPN22/LAG3/HLA-DRB1/CST7 | 9 | BP |
| GO:0031349 | positive regulation of defense response | 10/83 | 278/18723 | 4.15117E-07 | 1.48318E-05 | 1.07421E-05 | CCR7/TXK/HAVCR2/CCL3/KLRC2/ALOX5AP/CRTAM/LAG3/KLRD1/CCL5 | 10 | BP |
| GO:0002468 | dendritic cell antigen processing and presentation | 4/83 | 15/18723 | 4.72159E-07 | 1.62346E-05 | 1.17581E-05 | CCR7/CD74/HLA-DRA/HLA-DRB1 | 4 | BP |
| GO:0051250 | negative regulation of lymphocyte activation | 8/83 | 157/18723 | 4.73312E-07 | 1.62346E-05 | 1.17581E-05 | RIPOR2/HAVCR2/CD74/ID2/CRTAM/PTPN22/LAG3/HLA-DRB1 | 8 | BP |
| GO:0002456 | T cell mediated immunity | 7/83 | 109/18723 | 5.38414E-07 | 1.81055E-05 | 1.31131E-05 | IL7R/KLRC1/HLA-DRA/HLA-DRB1/PRF1/KLRD1/CD8A | 7 | BP |
| GO:0046651 | lymphocyte proliferation | 10/83 | 288/18723 | 5.73647E-07 | 1.89193E-05 | 1.37025E-05 | IL7R/LEF1/HAVCR2/CD74/CRTAM/PTPN22/HLA-DPB1/HLA-DPA1/HLA-DRB1/CCL5 | 10 | BP |
| GO:0032943 | mononuclear cell proliferation | 10/83 | 291/18723 | 6.30543E-07 | 2.04034E-05 | 1.47774E-05 | IL7R/LEF1/HAVCR2/CD74/CRTAM/PTPN22/HLA-DPB1/HLA-DPA1/HLA-DRB1/CCL5 | 10 | BP |
| GO:0002703 | regulation of leukocyte mediated immunity | 9/83 | 226/18723 | 7.0272E-07 | 2.23179E-05 | 1.61639E-05 | IL7R/HAVCR2/KLRC2/KLRC1/CRTAM/LAG3/HLA-DRA/HLA-DRB1/KLRD1 | 9 | BP |
| GO:0001914 | regulation of T cell mediated cytotoxicity | 5/83 | 39/18723 | 7.7512E-07 | 2.35128E-05 | 1.70294E-05 | IL7R/KLRC1/HLA-DRA/HLA-DRB1/KLRD1 | 5 | BP |
| GO:0002347 | response to tumor cell | 5/83 | 39/18723 | 7.7512E-07 | 2.35128E-05 | 1.70294E-05 | HAVCR2/CRTAM/ABI3/HLA-DRB1/PRF1 | 5 | BP |
| GO:0032729 | positive regulation of interferon-gamma production | 6/83 | 72/18723 | 7.81475E-07 | 2.35128E-05 | 1.70294E-05 | TXK/HAVCR2/CRTAM/PTPN22/HLA-DPB1/HLA-DPA1 | 6 | BP |
| GO:0071346 | cellular response to interferon-gamma | 7/83 | 118/18723 | 9.2267E-07 | 2.72824E-05 | 1.97595E-05 | TXK/CCL3/FASLG/HLA-DPA1/CCL4L1/CCL4/CCL5 | 7 | BP |
| GO:0050868 | negative regulation of T cell activation | 7/83 | 122/18723 | 1.1557E-06 | 3.35937E-05 | 2.43305E-05 | RIPOR2/HAVCR2/CD74/CRTAM/PTPN22/LAG3/HLA-DRB1 | 7 | BP |
| GO:0048245 | eosinophil chemotaxis | 4/83 | 19/18723 | 1.32273E-06 | 3.78081E-05 | 2.73829E-05 | CCL3/CCL4L1/CCL4/CCL5 | 4 | BP |
| GO:0045089 | positive regulation of innate immune response | 7/83 | 131/18723 | 1.86488E-06 | 5.24307E-05 | 3.79735E-05 | TXK/HAVCR2/KLRC2/CRTAM/LAG3/KLRD1/CCL5 | 7 | BP |
| GO:0043547 | positive regulation of GTPase activity | 9/83 | 255/18723 | 1.91305E-06 | 5.29174E-05 | 3.83259E-05 | CCR7/S1PR1/RACK1/CCL3/F2R/RGS1/CCL4L1/CCL4/CCL5 | 9 | BP |
| GO:0071674 | mononuclear cell migration | 8/83 | 196/18723 | 2.52011E-06 | 6.8603E-05 | 4.96864E-05 | CCR7/RIPOR2/S1PR1/CCL3/CRTAM/CCL4L1/CCL4/CCL5 | 8 | BP |
| GO:0070098 | chemokine-mediated signaling pathway | 6/83 | 88/18723 | 2.56199E-06 | 6.86532E-05 | 4.97228E-05 | CCR7/CCL3/CXCR6/CCL4L1/CCL4/CCL5 | 6 | BP |
| GO:0072677 | eosinophil migration | 4/83 | 23/18723 | 2.98134E-06 | 7.80086E-05 | 5.64985E-05 | CCL3/CCL4L1/CCL4/CCL5 | 4 | BP |
| GO:0050921 | positive regulation of chemotaxis | 7/83 | 141/18723 | 3.04757E-06 | 7.80086E-05 | 5.64985E-05 | CCR7/RIPOR2/S1PR1/CCL3/CD74/CCL4/CCL5 | 7 | BP |
| GO:1903038 | negative regulation of leukocyte cell-cell adhesion | 7/83 | 141/18723 | 3.04757E-06 | 7.80086E-05 | 5.64985E-05 | RIPOR2/HAVCR2/CD74/CRTAM/PTPN22/LAG3/HLA-DRB1 | 7 | BP |
| GO:0001911 | negative regulation of leukocyte mediated cytotoxicity | 4/83 | 24/18723 | 3.56555E-06 | 8.99252E-05 | 6.51292E-05 | IL7R/HAVCR2/KLRC1/KLRD1 | 4 | BP |
| GO:0002690 | positive regulation of leukocyte chemotaxis | 6/83 | 94/18723 | 3.7687E-06 | 9.26046E-05 | 6.70698E-05 | CCR7/RIPOR2/CCL3/CD74/CCL4/CCL5 | 6 | BP |
| GO:0016064 | immunoglobulin mediated immune response | 8/83 | 207/18723 | 3.77978E-06 | 9.26046E-05 | 6.70698E-05 | HLA-DQB1/CD74/HLA-DQA1/HLA-DPB1/HLA-DRA/HLA-DPA1/HLA-DRB5/HLA-DRB1 | 8 | BP |
| GO:0019724 | B cell mediated immunity | 8/83 | 210/18723 | 4.20406E-06 | 0.000101549 | 7.35477E-05 | HLA-DQB1/CD74/HLA-DQA1/HLA-DPB1/HLA-DRA/HLA-DPA1/HLA-DRB5/HLA-DRB1 | 8 | BP |
| GO:0002418 | immune response to tumor cell | 4/83 | 26/18723 | 4.98271E-06 | 0.00011706 | 8.47816E-05 | HAVCR2/CRTAM/HLA-DRB1/PRF1 | 4 | BP |
| GO:0045954 | positive regulation of natural killer cell mediated cytotoxicity | 4/83 | 26/18723 | 4.98271E-06 | 0.00011706 | 8.47816E-05 | KLRC2/CRTAM/LAG3/KLRD1 | 4 | BP |
| GO:0097529 | myeloid leukocyte migration | 8/83 | 220/18723 | 5.92234E-06 | 0.000137254 | 9.94077E-05 | CCR7/RIPOR2/CCL3/CD74/ITGA1/CCL4L1/CCL4/CCL5 | 8 | BP |
| GO:0031342 | negative regulation of cell killing | 4/83 | 28/18723 | 6.77825E-06 | 0.000154996 | 0.000112257 | IL7R/HAVCR2/KLRC1/KLRD1 | 4 | BP |
| GO:0050670 | regulation of lymphocyte proliferation | 8/83 | 225/18723 | 6.98361E-06 | 0.000157591 | 0.000114137 | HAVCR2/CD74/CRTAM/PTPN22/HLA-DPB1/HLA-DPA1/HLA-DRB1/CCL5 | 8 | BP |
| GO:0032944 | regulation of mononuclear cell proliferation | 8/83 | 227/18723 | 7.451E-06 | 0.000165954 | 0.000120194 | HAVCR2/CD74/CRTAM/PTPN22/HLA-DPB1/HLA-DPA1/HLA-DRB1/CCL5 | 8 | BP |
| GO:0002699 | positive regulation of immune effector process | 8/83 | 235/18723 | 9.59439E-06 | 0.000210954 | 0.000152785 | CD74/KLRC2/CRTAM/PTPN22/LAG3/HLA-DRA/HLA-DRB1/KLRD1 | 8 | BP |
| GO:0001915 | negative regulation of T cell mediated cytotoxicity | 3/83 | 10/18723 | 9.85705E-06 | 0.000211311 | 0.000153044 | IL7R/KLRC1/KLRD1 | 3 | BP |
| GO:0035747 | natural killer cell chemotaxis | 3/83 | 10/18723 | 9.85705E-06 | 0.000211311 | 0.000153044 | CCL3/CCL4/CCL5 | 3 | BP |
| GO:0002717 | positive regulation of natural killer cell mediated immunity | 4/83 | 31/18723 | 1.03116E-05 | 0.000218326 | 0.000158125 | KLRC2/CRTAM/LAG3/KLRD1 | 4 | BP |
| GO:0042129 | regulation of T cell proliferation | 7/83 | 171/18723 | 1.08587E-05 | 0.000225998 | 0.000163681 | HAVCR2/CRTAM/PTPN22/HLA-DPB1/HLA-DPA1/HLA-DRB1/CCL5 | 7 | BP |
| GO:0002708 | positive regulation of lymphocyte mediated immunity | 6/83 | 113/18723 | 1.09375E-05 | 0.000225998 | 0.000163681 | KLRC2/CRTAM/LAG3/HLA-DRA/HLA-DRB1/KLRD1 | 6 | BP |
| GO:0019835 | cytolysis | 4/83 | 32/18723 | 1.1745E-05 | 0.000239794 | 0.000173673 | PRF1/GZMH/GZMB/GZMA | 4 | BP |
| GO:0070663 | regulation of leukocyte proliferation | 8/83 | 245/18723 | 1.29857E-05 | 0.000262005 | 0.00018976 | HAVCR2/CD74/CRTAM/PTPN22/HLA-DPB1/HLA-DPA1/HLA-DRB1/CCL5 | 8 | BP |
| GO:0043122 | regulation of I-kappaB kinase/NF-kappaB signaling | 8/83 | 249/18723 | 1.45998E-05 | 0.000291147 | 0.000210866 | CCR7/PIM2/BIRC3/TNIP3/CD74/F2R/FASLG/HLA-DRB1 | 8 | BP |
| GO:0002761 | regulation of myeloid leukocyte differentiation | 6/83 | 120/18723 | 1.54278E-05 | 0.000304123 | 0.000220264 | LEF1/MYC/CCL3/CD74/ID2/HLA-DRB1 | 6 | BP |
| GO:0032814 | regulation of natural killer cell activation | 4/83 | 35/18723 | 1.69293E-05 | 0.000326671 | 0.000236595 | HAVCR2/KLRC2/PTPN22/ZNF683 | 4 | BP |
| GO:0002688 | regulation of leukocyte chemotaxis | 6/83 | 122/18723 | 1.69526E-05 | 0.000326671 | 0.000236595 | CCR7/RIPOR2/CCL3/CD74/CCL4/CCL5 | 6 | BP |
| GO:0050852 | T cell receptor signaling pathway | 6/83 | 123/18723 | 1.77594E-05 | 0.000338414 | 0.0002451 | CCR7/TXK/HLA-DQB1/PTPN22/HLA-DPB1/HLA-DRB1 | 6 | BP |
| GO:0043123 | positive regulation of I-kappaB kinase/NF-kappaB signaling | 7/83 | 186/18723 | 1.87357E-05 | 0.000353097 | 0.000255734 | CCR7/PIM2/BIRC3/CD74/F2R/FASLG/HLA-DRB1 | 7 | BP |
| GO:0051607 | defense response to virus | 8/83 | 265/18723 | 2.28526E-05 | 0.000420045 | 0.000304221 | SERINC5/BIRC3/OASL/LYST/APOBEC3C/PTPN22/APOBEC3G/PRF1 | 8 | BP |
| GO:0140546 | defense response to symbiont | 8/83 | 265/18723 | 2.28526E-05 | 0.000420045 | 0.000304221 | SERINC5/BIRC3/OASL/LYST/APOBEC3C/PTPN22/APOBEC3G/PRF1 | 8 | BP |
| GO:0001768 | establishment of T cell polarity | 3/83 | 13/18723 | 2.32678E-05 | 0.000420045 | 0.000304221 | CCR7/RIPOR2/CRTAM | 3 | BP |
| GO:0042492 | gamma-delta T cell differentiation | 3/83 | 13/18723 | 2.32678E-05 | 0.000420045 | 0.000304221 | LEF1/TCF7/KLRC1 | 3 | BP |
| GO:0002285 | lymphocyte activation involved in immune response | 7/83 | 194/18723 | 2.45711E-05 | 0.000438953 | 0.000317916 | LEF1/HAVCR2/CD74/KLRC2/HLA-DRA/ZNF683/HLA-DRB1 | 7 | BP |
| GO:0022408 | negative regulation of cell-cell adhesion | 7/83 | 196/18723 | 2.62429E-05 | 0.000463986 | 0.000336046 | RIPOR2/HAVCR2/CD74/CRTAM/PTPN22/LAG3/HLA-DRB1 | 7 | BP |
| GO:0002705 | positive regulation of leukocyte mediated immunity | 6/83 | 134/18723 | 2.88596E-05 | 0.000495751 | 0.000359052 | KLRC2/CRTAM/LAG3/HLA-DRA/HLA-DRB1/KLRD1 | 6 | BP |
| GO:0042098 | T cell proliferation | 7/83 | 199/18723 | 2.89256E-05 | 0.000495751 | 0.000359052 | HAVCR2/CRTAM/PTPN22/HLA-DPB1/HLA-DPA1/HLA-DRB1/CCL5 | 7 | BP |
| GO:0001767 | establishment of lymphocyte polarity | 3/83 | 14/18723 | 2.95188E-05 | 0.000495751 | 0.000359052 | CCR7/RIPOR2/CRTAM | 3 | BP |
| GO:0001771 | immunological synapse formation | 3/83 | 14/18723 | 2.95188E-05 | 0.000495751 | 0.000359052 | CCR7/HAVCR2/PRF1 | 3 | BP |
| GO:0043922 | negative regulation by host of viral transcription | 3/83 | 14/18723 | 2.95188E-05 | 0.000495751 | 0.000359052 | CCL3/CCL4/CCL5 | 3 | BP |
| GO:0002366 | leukocyte activation involved in immune response | 8/83 | 275/18723 | 2.9774E-05 | 0.000495751 | 0.000359052 | LEF1/HAVCR2/CCL3/CD74/KLRC2/HLA-DRA/ZNF683/HLA-DRB1 | 8 | BP |
| GO:0002687 | positive regulation of leukocyte migration | 6/83 | 135/18723 | 3.00958E-05 | 0.000496291 | 0.000359444 | CCR7/RIPOR2/CCL3/CD74/CCL4/CCL5 | 6 | BP |
| GO:0050671 | positive regulation of lymphocyte proliferation | 6/83 | 137/18723 | 3.2696E-05 | 0.000533846 | 0.000386643 | HAVCR2/CD74/PTPN22/HLA-DPB1/HLA-DPA1/CCL5 | 6 | BP |
| GO:0002263 | cell activation involved in immune response | 8/83 | 279/18723 | 3.29957E-05 | 0.000533846 | 0.000386643 | LEF1/HAVCR2/CCL3/CD74/KLRC2/HLA-DRA/ZNF683/HLA-DRB1 | 8 | BP |
| GO:0032946 | positive regulation of mononuclear cell proliferation | 6/83 | 138/18723 | 3.40622E-05 | 0.00054595 | 0.000395409 | HAVCR2/CD74/PTPN22/HLA-DPB1/HLA-DPA1/CCL5 | 6 | BP |
| GO:0007249 | I-kappaB kinase/NF-kappaB signaling | 8/83 | 281/18723 | 3.4713E-05 | 0.000551229 | 0.000399233 | CCR7/PIM2/BIRC3/TNIP3/CD74/F2R/FASLG/HLA-DRB1 | 8 | BP |
| GO:0002709 | regulation of T cell mediated immunity | 5/83 | 85/18723 | 3.76488E-05 | 0.000592364 | 0.000429025 | IL7R/KLRC1/HLA-DRA/HLA-DRB1/KLRD1 | 5 | BP |
| GO:0002573 | myeloid leukocyte differentiation | 7/83 | 208/18723 | 3.83624E-05 | 0.000598105 | 0.000433183 | CCR7/LEF1/MYC/CCL3/CD74/ID2/HLA-DRB1 | 7 | BP |
| GO:0002429 | immune response-activating cell surface receptor signaling pathway | 8/83 | 291/18723 | 4.44639E-05 | 0.000675381 | 0.000489151 | CCR7/TXK/HLA-DQB1/KLRC2/PTPN22/HLA-DPB1/HLA-DRB1/KLRD1 | 8 | BP |
| GO:0002757 | immune response-activating signal transduction | 8/83 | 291/18723 | 4.44639E-05 | 0.000675381 | 0.000489151 | CCR7/TXK/HLA-DQB1/KLRC2/PTPN22/HLA-DPB1/HLA-DRB1/KLRD1 | 8 | BP |

| | | | | | | | | | |
|---|---|---|---|---|---|---|---|---|---|
| GO:0030101 | natural killer cell activation | 5/83 | 88/18723 | 4.45003E-05 | 0.000675381 | 0.000489151 | HAVCR2/KLRC2/PTPN22/SLAMF7/ZNF683 | 5 | BP |
| GO:0002377 | immunoglobulin production | 7/83 | 216/18723 | 4.8749E-05 | 0.000733373 | 0.000531152 | HLA-DQB1/HLA-DQA1/HLA-DPB1/HLA-DRA/HLA-DPA1/HLA-DRB5/HLA-DRB1 | 7 | BP |
| GO:1903900 | regulation of viral life cycle | 6/83 | 148/18723 | 5.04158E-05 | 0.00075167 | 0.000544404 | OASL/CD74/APOBEC3C/HLA-DRB1/APOBEC3G/CCL5 | 6 | BP |
| GO:2000107 | negative regulation of leukocyte apoptotic process | 4/83 | 46/18723 | 5.08418E-05 | 0.00075167 | 0.000544404 | IL7R/CCR7/CD74/CCL5 | 4 | BP |
| GO:0045582 | positive regulation of T cell differentiation | 5/83 | 91/18723 | 5.22794E-05 | 0.000766317 | 0.000555013 | IL7R/LEF1/CD74/HLA-DRA/HLA-DRB1 | 5 | BP |
| GO:0070665 | positive regulation of leukocyte proliferation | 6/83 | 150/18723 | 5.43364E-05 | 0.00078972 | 0.000571963 | HAVCR2/CD74/PTPN22/HLA-DPB1/HLA-DPA1/CCL5 | 6 | BP |
| GO:0050920 | regulation of chemotaxis | 7/83 | 223/18723 | 5.96331E-05 | 0.000859418 | 0.000622442 | CCR7/RIPOR2/S1PR1/CCL3/CD74/CCL4/CCL5 | 7 | BP |
| GO:0045953 | negative regulation of natural killer cell mediated cytotoxicity | 3/83 | 18/18723 | 6.53315E-05 | 0.000933696 | 0.000676238 | HAVCR2/KLRC1/KLRD1 | 3 | BP |
| GO:0046631 | alpha-beta T cell activation | 6/83 | 156/18723 | 6.75835E-05 | 0.000957899 | 0.000693768 | LEF1/CRTAM/PTPN22/HLA-DRA/ZNF683/HLA-DRB1 | 6 | BP |
| GO:1902107 | positive regulation of leukocyte differentiation | 6/83 | 157/18723 | 7.00224E-05 | 0.000976328 | 0.000707115 | IL7R/LEF1/CD74/ID2/HLA-DRA/HLA-DRB1 | 6 | BP |
| GO:1903708 | positive regulation of hemopoiesis | 6/83 | 157/18723 | 7.00224E-05 | 0.000976328 | 0.000707115 | IL7R/LEF1/CD74/ID2/HLA-DRA/HLA-DRB1 | 6 | BP |
| GO:0019083 | viral transcription | 4/83 | 50/18723 | 7.07922E-05 | 0.000979101 | 0.000709124 | LEF1/CCL3/CCL4/CCL5 | 4 | BP |
| GO:0071622 | regulation of granulocyte chemotaxis | 4/83 | 51/18723 | 7.65589E-05 | 0.001044302 | 0.000756346 | CCR7/RIPOR2/CD74/CCL5 | 4 | BP |
| GO:0002716 | negative regulation of natural killer cell mediated immunity | 3/83 | 19/18723 | 7.73331E-05 | 0.001044302 | 0.000756346 | HAVCR2/KLRC1/KLRD1 | 3 | BP |
| GO:0140131 | positive regulation of lymphocyte chemotaxis | 3/83 | 19/18723 | 7.73331E-05 | 0.001044302 | 0.000756346 | CCL3/CCL4/CCL5 | 3 | BP |
| GO:2000116 | regulation of cysteine-type endopeptidase activity | 7/83 | 235/18723 | 8.28831E-05 | 0.001110504 | 0.000804293 | BIRC3/MYC/RACK1/F2R/CTSD/FASLG/CST7 | 7 | BP |
| GO:0042102 | positive regulation of T cell proliferation | 5/83 | 101/18723 | 8.60163E-05 | 0.00114355 | 0.000828227 | HAVCR2/PTPN22/HLA-DPB1/HLA-DPA1/CCL5 | 5 | BP |
| GO:0002707 | negative regulation of lymphocyte mediated immunity | 4/83 | 53/18723 | 8.91074E-05 | 0.001152091 | 0.000834413 | IL7R/HAVCR2/KLRC1/KLRD1 | 4 | BP |
| GO:0050792 | regulation of viral process | 6/83 | 164/18723 | 8.9131E-05 | 0.001152091 | 0.000834413 | OASL/CD74/APOBEC3C/HLA-DRB1/APOBEC3G/CCL5 | 6 | BP |
| GO:1902106 | negative regulation of leukocyte differentiation | 5/83 | 102/18723 | 9.01378E-05 | 0.001152091 | 0.000834413 | MYC/CCL3/CD74/ID2/LAG3 | 5 | BP |
| GO:0002834 | regulation of response to tumor cell | 3/83 | 20/18723 | 9.06894E-05 | 0.001152091 | 0.000834413 | HAVCR2/CRTAM/HLA-DRB1 | 3 | BP |
| GO:0002837 | regulation of immune response to tumor cell | 3/83 | 20/18723 | 9.06894E-05 | 0.001152091 | 0.000834413 | HAVCR2/CRTAM/HLA-DRB1 | 3 | BP |
| GO:0046629 | gamma-delta T cell activation | 3/83 | 20/18723 | 9.06894E-05 | 0.001152091 | 0.000834413 | LEF1/TCF7/KLRC1 | 3 | BP |
| GO:0002823 | negative regulation of adaptive immune response based on somatic recombination of immune receptors built from immunoglobulin superfamily domains | 4/83 | 54/18723 | 9.59127E-05 | 0.001200659 | 0.000869589 | IL7R/HAVCR2/KLRC1/KLRD1 | 4 | BP |
| GO:0006968 | cellular defense response | 4/83 | 54/18723 | 9.59127E-05 | 0.001200659 | 0.000869589 | FCMR/KLRC3/KLRC2/PRF1 | 4 | BP |
| GO:0045621 | positive regulation of lymphocyte differentiation | 5/83 | 104/18723 | 9.88352E-05 | 0.001219442 | 0.000883192 | IL7R/LEF1/CD74/HLA-DRA/HLA-DRB1 | 5 | BP |
| GO:0046634 | regulation of alpha-beta T cell activation | 5/83 | 104/18723 | 9.88352E-05 | 0.001219442 | 0.000883192 | CRTAM/PTPN22/HLA-DRA/ZNF683/HLA-DRB1 | 5 | BP |
| GO:0002822 | regulation of adaptive immune response based on somatic recombination of immune receptors built from immunoglobulin superfamily domains | 6/83 | 168/18723 | 0.000101785 | 0.001246866 | 0.000903055 | IL7R/HAVCR2/KLRC1/HLA-DRA/HLA-DRB1/KLRD1 | 6 | BP |
| GO:0045655 | regulation of monocyte differentiation | 3/83 | 21/18723 | 0.000105466 | 0.001282798 | 0.000929078 | MYC/CD74/HLA-DRB1 | 3 | BP |
| GO:0035821 | modulation of process of other organism | 5/83 | 106/18723 | 0.000108164 | 0.0012882 | 0.000932991 | LEF1/CCL3/PRF1/CCL4/CCL5 | 5 | BP |
| GO:0071887 | leukocyte apoptotic process | 5/83 | 106/18723 | 0.000108164 | 0.0012882 | 0.000932991 | IL7R/CCR7/CD74/FASLG/CCL5 | 5 | BP |
| GO:1903707 | negative regulation of hemopoiesis | 5/83 | 106/18723 | 0.000108164 | 0.0012882 | 0.000932991 | MYC/CCL3/CD74/ID2/LAG3 | 5 | BP |
| GO:0045071 | negative regulation of viral genome replication | 4/83 | 56/18723 | 0.000110645 | 0.001308659 | 0.000947809 | OASL/APOBEC3C/APOBEC3G/CCL5 | 4 | BP |
| GO:0002763 | positive regulation of myeloid leukocyte differentiation | 4/83 | 58/18723 | 0.000126953 | 0.001491258 | 0.001080058 | LEF1/CD74/ID2/HLA-DRB1 | 4 | BP |
| GO:0002820 | negative regulation of adaptive immune response | 4/83 | 59/18723 | 0.000135728 | 0.001583496 | 0.001146862 | IL7R/HAVCR2/KLRC1/KLRD1 | 4 | BP |
| GO:0051851 | modulation by host of symbiont process | 4/83 | 60/18723 | 0.000144935 | 0.001679481 | 0.00121638 | LEF1/CCL3/CCL4/CCL5 | 4 | BP |
| GO:0002286 | T cell activation involved in immune response | 5/83 | 114/18723 | 0.000152413 | 0.001754278 | 0.001270553 | LEF1/HAVCR2/CD74/HLA-DRA/HLA-DRB1 | 5 | BP |
| GO:0043030 | regulation of macrophage activation | 4/83 | 61/18723 | 0.000154585 | 0.001755716 | 0.001271594 | HAVCR2/CCL3/CD74/CST7 | 4 | BP |
| GO:2000401 | regulation of lymphocyte migration | 4/83 | 61/18723 | 0.000154585 | 0.001755716 | 0.001271594 | RIPOR2/CCL3/CCL4/CCL5 | 4 | BP |
| GO:0071675 | regulation of mononuclear cell migration | 5/83 | 115/18723 | 0.000158791 | 0.001781869 | 0.001290536 | CCR7/RIPOR2/CCL3/CCL4/CCL5 | 5 | BP |
| GO:0090023 | positive regulation of neutrophil chemotaxis | 3/83 | 24/18723 | 0.000158966 | 0.001781869 | 0.001290536 | CCR7/RIPOR2/CD74 | 3 | BP |
| GO:0002819 | regulation of adaptive immune response | 6/83 | 183/18723 | 0.000162516 | 0.001809836 | 0.001310791 | IL7R/HAVCR2/KLRC1/HLA-DRA/HLA-DRB1/KLRD1 | 6 | BP |
| GO:0002704 | negative regulation of leukocyte mediated immunity | 4/83 | 63/18723 | 0.000175266 | 0.001939239 | 0.001404512 | IL7R/HAVCR2/KLRC1/KLRD1 | 4 | BP |
| GO:1901623 | regulation of lymphocyte chemotaxis | 3/83 | 25/18723 | 0.000180066 | 0.00197957 | 0.001433723 | CCL3/CCL4/CCL5 | 3 | BP |
| GO:0048247 | lymphocyte chemotaxis | 4/83 | 64/18723 | 0.000186323 | 0.002035317 | 0.001474098 | CCL3/CCL4L1/CCL4/CCL5 | 4 | BP |
| GO:0071677 | positive regulation of mononuclear cell migration | 4/83 | 65/18723 | 0.000197875 | 0.002147822 | 0.001555581 | CCR7/CCL3/CCL4/CCL5 | 4 | BP |
| GO:0002710 | negative regulation of T cell mediated immunity | 3/83 | 26/18723 | 0.000202903 | 0.002174864 | 0.001575166 | IL7R/KLRC1/KLRD1 | 3 | BP |
| GO:0019883 | antigen processing and presentation of endogenous antigen | 3/83 | 26/18723 | 0.000202903 | 0.002174864 | 0.001575166 | CD74/HLA-DRA/HLA-DRB1 | 3 | BP |
| GO:0072678 | T cell migration | 4/83 | 66/18723 | 0.000209934 | 0.002236258 | 0.001619632 | RIPOR2/S1PR1/CCL3/CCL5 | 4 | BP |
| GO:0071624 | positive regulation of granulocyte chemotaxis | 3/83 | 27/18723 | 0.000227537 | 0.002408802 | 0.001744598 | CCR7/RIPOR2/CD74 | 3 | BP |
| GO:0045589 | regulation of regulatory T cell differentiation | 3/83 | 28/18723 | 0.000254028 | 0.002656451 | 0.00192396 | LAG3/HLA-DRA/HLA-DRB1 | 3 | BP |
| GO:1902624 | positive regulation of neutrophil migration | 3/83 | 28/18723 | 0.000254028 | 0.002656451 | 0.00192396 | CCR7/RIPOR2/CD74 | 3 | BP |
| GO:0002548 | monocyte chemotaxis | 4/83 | 70/18723 | 0.000263511 | 0.002738921 | 0.00198369 | CCL3/CCL4L1/CCL4/CCL5 | 4 | BP |
| GO:0043280 | positive regulation of cysteine-type endopeptidase activity involved in apoptotic process | 5/83 | 129/18723 | 0.00027132 | 0.002803096 | 0.002030169 | MYC/RACK1/F2R/CTSD/FASLG | 5 | BP |
| GO:0019079 | viral genome replication | 5/83 | 131/18723 | 0.000291351 | 0.002992021 | 0.002167 | OASL/APOBEC3C/CXCR6/APOBEC3G/CCL5 | 5 | BP |
| GO:0070227 | lymphocyte apoptotic process | 4/83 | 72/18723 | 0.00029369 | 0.002998082 | 0.00217139 | IL7R/CD74/FASLG/CCL5 | 4 | BP |
| GO:0070229 | negative regulation of lymphocyte apoptotic process | 3/83 | 30/18723 | 0.000312815 | 0.003174425 | 0.002299108 | IL7R/CD74/CCL5 | 3 | BP |
| GO:0043281 | regulation of cysteine-type endopeptidase activity involved in apoptotic process | 6/83 | 209/18723 | 0.000332617 | 0.003355522 | 0.002430269 | BIRC3/MYC/RACK1/F2R/CTSD/FASLG | 6 | BP |
| GO:0002685 | regulation of leukocyte migration | 6/83 | 210/18723 | 0.000341201 | 0.003402093 | 0.002463998 | CCR7/RIPOR2/CCL3/CD74/CCL4/CCL5 | 6 | BP |
| GO:0045637 | regulation of myeloid cell differentiation | 6/83 | 210/18723 | 0.000341201 | 0.003402093 | 0.002463998 | LEF1/MYC/CCL3/CD74/ID2/HLA-DRB1 | 6 | BP |
| GO:0045066 | regulatory T cell differentiation | 3/83 | 31/18723 | 0.000345226 | 0.003422329 | 0.002478655 | LAG3/HLA-DRA/HLA-DRB1 | 3 | BP |
| GO:0090022 | regulation of neutrophil chemotaxis | 3/83 | 32/18723 | 0.000379724 | 0.003742684 | 0.002710675 | CCR7/RIPOR2/CD74 | 3 | BP |
| GO:0032633 | interleukin-4 production | 3/83 | 33/18723 | 0.000416365 | 0.004034268 | 0.002921857 | LEF1/HAVCR2/HLA-DRB1 | 3 | BP |
| GO:0032673 | regulation of interleukin-4 production | 3/83 | 33/18723 | 0.000416365 | 0.004034268 | 0.002921857 | LEF1/HAVCR2/HLA-DRB1 | 3 | BP |
| GO:0071353 | cellular response to interleukin-4 | 3/83 | 33/18723 | 0.000416365 | 0.004034268 | 0.002921857 | LEF1/TCF7/RPLP0 | 3 | BP |
| GO:0051817 | modulation of process of other organism involved in symbiotic interaction | 4/83 | 81/18723 | 0.000460813 | 0.004415057 | 0.003197647 | LEF1/CCL3/CCL4/CCL5 | 4 | BP |
| GO:2000106 | regulation of leukocyte apoptotic process | 4/83 | 81/18723 | 0.000460813 | 0.004415057 | 0.003197647 | IL7R/CCR7/CD74/CCL5 | 4 | BP |
| GO:2000403 | positive regulation of lymphocyte migration | 3/83 | 35/18723 | 0.00049629 | 0.004728542 | 0.003424692 | CCL3/CCL4/CCL5 | 3 | BP |
| GO:2001056 | positive regulation of cysteine-type endopeptidase activity | 5/83 | 148/18723 | 0.000510253 | 0.004834714 | 0.003501588 | MYC/RACK1/F2R/CTSD/FASLG | 5 | BP |
| GO:0030224 | monocyte differentiation | 3/83 | 36/18723 | 0.000539682 | 0.005057673 | 0.003663069 | MYC/CD74/HLA-DRB1 | 3 | BP |
| GO:0070670 | response to interleukin-4 | 3/83 | 36/18723 | 0.000539682 | 0.005057673 | 0.003663069 | LEF1/TCF7/RPLP0 | 3 | BP |
| GO:0045069 | regulation of viral genome replication | 4/83 | 85/18723 | 0.000553338 | 0.005157472 | 0.003735349 | OASL/APOBEC3C/APOBEC3G/CCL5 | 4 | BP |
| GO:0050851 | antigen receptor-mediated signaling pathway | 6/83 | 240/18723 | 0.000690599 | 0.006402037 | 0.004636737 | CCR7/TXK/HLA-DQB1/PTPN22/HLA-DPB1/HLA-DRB1 | 6 | BP |
| GO:0048525 | negative regulation of viral process | 4/83 | 92/18723 | 0.00074588 | 0.006877339 | 0.004980979 | OASL/APOBEC3C/APOBEC3G/CCL5 | 4 | BP |
| GO:1902622 | regulation of neutrophil migration | 3/83 | 41/18723 | 0.000792991 | 0.007272622 | 0.005267266 | CCR7/RIPOR2/CD74 | 3 | BP |
| GO:0019080 | viral gene expression | 4/83 | 94/18723 | 0.000808605 | 0.007337346 | 0.005314143 | LEF1/CCL3/CCL4/CCL5 | 4 | BP |
| GO:0051702 | biological process involved in interaction with symbiont | 4/83 | 94/18723 | 0.000808605 | 0.007337346 | 0.005314143 | LEF1/CCL3/CCL4/CCL5 | 4 | BP |
| GO:0070383 | DNA cytosine deamination | 2/83 | 10/18723 | 0.000853792 | 0.007706595 | 0.005581575 | APOBEC3C/APOBEC3G | 2 | BP |
| GO:0042088 | T-helper 1 type immune response | 3/83 | 43/18723 | 0.000912204 | 0.008190734 | 0.005932218 | LEF1/HAVCR2/HLA-DRB1 | 3 | BP |
| GO:0031348 | negative regulation of defense response | 6/83 | 258/18723 | 0.001005125 | 0.008922468 | 0.006462184 | HAVCR2/KLRC1/HLA-DRB1/CST7/AOAH/KLRD1 | 6 | BP |
| GO:0002221 | pattern recognition receptor signaling pathway | 5/83 | 172/18723 | 0.001005486 | 0.008922468 | 0.006462184 | BIRC3/OASL/TNIP3/HAVCR2/PTPN22 | 5 | BP |
| GO:0002357 | defense response to tumor cell | 2/83 | 11/18723 | 0.001040521 | 0.008922468 | 0.006462184 | ABI3/PRF1 | 2 | BP |

| | | | | | | | | | |
|---|---|---|---|---|---|---|---|---|---|
| GO:0002604 | regulation of dendritic cell antigen processing and presentation | 2/83 | 11/18723 | 0.001040521 | 0.008922468 | 0.006462184 | CCR7/CD74 | 2 | BP |
| GO:0045060 | negative thymic T cell selection | 2/83 | 11/18723 | 0.001040521 | 0.008922468 | 0.006462184 | CCR7/CD74 | 2 | BP |
| GO:0045657 | positive regulation of monocyte differentiation | 2/83 | 11/18723 | 0.001040521 | 0.008922468 | 0.006462184 | CD74/HLA-DRB1 | 2 | BP |
| GO:0046598 | positive regulation of viral entry into host cell | 2/83 | 11/18723 | 0.001040521 | 0.008922468 | 0.006462184 | CD74/HLA-DRB1 | 2 | BP |
| GO:0046643 | regulation of gamma-delta T cell activation | 2/83 | 11/18723 | 0.001040521 | 0.008922468 | 0.006462184 | LEF1/TCF7 | 2 | BP |
| GO:0075294 | positive regulation by symbiont of entry into host | 2/83 | 11/18723 | 0.001040521 | 0.008922468 | 0.006462184 | CD74/HLA-DRB1 | 2 | BP |
| GO:0045639 | positive regulation of myeloid cell differentiation | 4/83 | 103/18723 | 0.001137534 | 0.009705829 | 0.00702954 | LEF1/CD74/ID2/HLA-DRB1 | 4 | BP |
| GO:0048661 | positive regulation of smooth muscle cell proliferation | 4/83 | 104/18723 | 0.001179082 | 0.010010527 | 0.00725022 | S1PR1/LDLRAP1/ID2/CCL5 | 4 | BP |
| GO:0010950 | positive regulation of endopeptidase activity | 5/83 | 179/18723 | 0.00120099 | 0.010146299 | 0.007348554 | MYC/RACK1/F2R/CTSD/FASLG | 5 | BP |
| GO:0006216 | cytidine catabolic process | 2/83 | 12/18723 | 0.001245034 | 0.010167779 | 0.007364112 | APOBEC3C/APOBEC3G | 2 | BP |
| GO:0009972 | cytidine deamination | 2/83 | 12/18723 | 0.001245034 | 0.010167779 | 0.007364112 | APOBEC3C/APOBEC3G | 2 | BP |
| GO:0016554 | cytidine to uridine editing | 2/83 | 12/18723 | 0.001245034 | 0.010167779 | 0.007364112 | APOBEC3C/APOBEC3G | 2 | BP |
| GO:0034154 | toll-like receptor 7 signaling pathway | 2/83 | 12/18723 | 0.001245034 | 0.010167779 | 0.007364112 | HAVCR2/PTPN22 | 2 | BP |
| GO:0043380 | regulation of memory T cell differentiation | 2/83 | 12/18723 | 0.001245034 | 0.010167779 | 0.007364112 | HLA-DRA/HLA-DRB1 | 2 | BP |
| GO:0043383 | negative T cell selection | 2/83 | 12/18723 | 0.001245034 | 0.010167779 | 0.007364112 | CCR7/CD74 | 2 | BP |
| GO:0046087 | cytidine metabolic process | 2/83 | 12/18723 | 0.001245034 | 0.010167779 | 0.007364112 | APOBEC3C/APOBEC3G | 2 | BP |
| GO:0042116 | macrophage activation | 4/83 | 106/18723 | 0.00126535 | 0.01028472 | 0.007448808 | HAVCR2/CCL3/CD74/CST7 | 4 | BP |
| GO:0007229 | integrin-mediated signaling pathway | 4/83 | 107/18723 | 0.001310099 | 0.010598207 | 0.007675853 | TXK/CD63/ITGAE/ITGA1 | 4 | BP |
| GO:1904894 | positive regulation of receptor signaling pathway via STAT | 3/83 | 49/18723 | 0.001336051 | 0.010757405 | 0.007791154 | IL7R/F2R/CCL5 | 3 | BP |
| GO:0070231 | T cell apoptotic process | 3/83 | 50/18723 | 0.001416812 | 0.011354359 | 0.008223504 | IL7R/FASLG/CCL5 | 3 | BP |
| GO:0002698 | negative regulation of immune effector process | 4/83 | 110/18723 | 0.001450975 | 0.011574053 | 0.008382619 | IL7R/HAVCR2/KLRC1/KLRD1 | 4 | BP |
| GO:0043379 | memory T cell differentiation | 2/83 | 13/18723 | 0.001467173 | 0.011595403 | 0.008398083 | HLA-DRA/HLA-DRB1 | 2 | BP |
| GO:0070424 | regulation of nucleotide-binding oligomerization domain containing signaling pathway | 2/83 | 13/18723 | 0.001467173 | 0.011595403 | 0.008398083 | BIRC3/PTPN22 | 2 | BP |
| GO:0046632 | alpha-beta T cell differentiation | 4/83 | 112/18723 | 0.001550559 | 0.012198203 | 0.008834666 | LEF1/HLA-DRA/ZNF683/HLA-DRB1 | 4 | BP |
| GO:0071347 | cellular response to interleukin-1 | 4/83 | 113/18723 | 0.001602094 | 0.012546076 | 0.009086616 | CCL3/CCL4L1/CCL4/CCL5 | 4 | BP |
| GO:0045006 | DNA deamination | 2/83 | 14/18723 | 0.001706783 | 0.013057672 | 0.009457145 | APOBEC3C/APOBEC3G | 2 | BP |
| GO:0046131 | pyrimidine ribonucleoside metabolic process | 2/83 | 14/18723 | 0.001706783 | 0.013057672 | 0.009457145 | APOBEC3C/APOBEC3G | 2 | BP |
| GO:0046133 | pyrimidine ribonucleoside catabolic process | 2/83 | 14/18723 | 0.001706783 | 0.013057672 | 0.009457145 | APOBEC3C/APOBEC3G | 2 | BP |
| GO:0090715 | immunological memory formation process | 2/83 | 14/18723 | 0.001706783 | 0.013057672 | 0.009457145 | HLA-DRA/HLA-DRB1 | 2 | BP |
| GO:0051209 | release of sequestered calcium ion into cytosol | 4/83 | 115/18723 | 0.00170872 | 0.013057672 | 0.009457145 | CCR7/CCL3/F2R/FASLG | 4 | BP |
| GO:0050777 | negative regulation of immune response | 5/83 | 194/18723 | 0.001713106 | 0.013057672 | 0.009457145 | IL7R/HAVCR2/KLRC1/HLA-DRB1/KLRD1 | 5 | BP |
| GO:0051283 | negative regulation of sequestering of calcium ion | 4/83 | 116/18723 | 0.00176384 | 0.013374795 | 0.009686824 | CCR7/CCL3/F2R/FASLG | 4 | BP |
| GO:0070228 | regulation of lymphocyte apoptotic process | 3/83 | 54/18723 | 0.001770308 | 0.013374795 | 0.009686824 | IL7R/CD74/CCL5 | 3 | BP |
| GO:0044403 | biological process involved in symbiotic interaction | 6/83 | 290/18723 | 0.001824297 | 0.013721554 | 0.009937968 | LEF1/CCL3/CD74/HLA-DRB1/CCL4/CCL5 | 6 | BP |
| GO:0010952 | positive regulation of peptidase activity | 5/83 | 197/18723 | 0.001832208 | 0.013721554 | 0.009937968 | MYC/RACK1/F2R/CTSD/FASLG | 5 | BP |
| GO:0006414 | translational elongation | 3/83 | 55/18723 | 0.001866494 | 0.013857303 | 0.010036285 | EEF1G/EEF2/RACK1 | 3 | BP |
| GO:0045620 | negative regulation of lymphocyte differentiation | 3/83 | 55/18723 | 0.001866494 | 0.013857303 | 0.010036285 | CD74/ID2/LAG3 | 3 | BP |
| GO:0051282 | regulation of sequestering of calcium ion | 4/83 | 118/18723 | 0.001877762 | 0.013880871 | 0.010053355 | CCR7/CCL3/F2R/FASLG | 4 | BP |
| GO:0002836 | positive regulation of response to tumor cell | 2/83 | 15/18723 | 0.001963706 | 0.014392119 | 0.010423631 | CRTAM/HLA-DRB1 | 2 | BP |
| GO:0002839 | positive regulation of immune response to tumor cell | 2/83 | 15/18723 | 0.001963706 | 0.014392119 | 0.010423631 | CRTAM/HLA-DRB1 | 2 | BP |
| GO:0002224 | toll-like receptor signaling pathway | 4/83 | 121/18723 | 0.002058065 | 0.015019497 | 0.010878015 | BIRC3/TNIP3/HAVCR2/PTPN22 | 4 | BP |
| GO:0051208 | sequestering of calcium ion | 4/83 | 122/18723 | 0.002120734 | 0.015346238 | 0.011114661 | CCR7/CCL3/F2R/FASLG | 4 | BP |
| GO:0051928 | positive regulation of calcium ion transport | 4/83 | 122/18723 | 0.002120734 | 0.015346238 | 0.011114661 | CCL3/F2R/CCL4/CCL5 | 4 | BP |
| GO:0043525 | positive regulation of neuron apoptotic process | 3/83 | 58/18723 | 0.002174412 | 0.015668554 | 0.011348101 | CCL3/FASLG/ITGA1 | 3 | BP |
| GO:0034138 | toll-like receptor 3 signaling pathway | 2/83 | 16/18723 | 0.002237789 | 0.015924517 | 0.011533485 | HAVCR2/PTPN22 | 2 | BP |
| GO:0045869 | negative regulation of single stranded viral RNA replication via double stranded DNA intermediate | 2/83 | 16/18723 | 0.002237789 | 0.015924517 | 0.011533485 | APOBEC3C/APOBEC3G | 2 | BP |
| GO:0046135 | pyrimidine nucleoside catabolic process | 2/83 | 16/18723 | 0.002237789 | 0.015924517 | 0.011533485 | APOBEC3C/APOBEC3G | 2 | BP |
| GO:0071222 | cellular response to lipopolysaccharide | 5/83 | 209/18723 | 0.002370046 | 0.016795989 | 0.012164656 | TNIP3/HAVCR2/CCL3/PTPN22/CCL5 | 5 | BP |
| GO:0002753 | cytoplasmic pattern recognition receptor signaling pathway | 3/83 | 60/18723 | 0.00239619 | 0.01684207 | 0.012198031 | BIRC3/OASL/PTPN22 | 3 | BP |
| GO:0031663 | lipopolysaccharide-mediated signaling pathway | 3/83 | 60/18723 | 0.00239619 | 0.01684207 | 0.012198031 | CCL3/PTPN22/CCL5 | 3 | BP |
| GO:0043491 | protein kinase B signaling | 5/83 | 211/18723 | 0.002469763 | 0.017278993 | 0.012514477 | CCR7/THEM4/RACK1/CCL3/CCL5 | 5 | BP |
| GO:0032612 | interleukin-1 production | 4/83 | 128/18723 | 0.002524677 | 0.017278993 | 0.012514477 | CCR7/HAVCR2/CCL3/F2R | 4 | BP |
| GO:0032652 | regulation of interleukin-1 production | 4/83 | 128/18723 | 0.002524677 | 0.017278993 | 0.012514477 | CCR7/HAVCR2/CCL3/F2R | 4 | BP |
| GO:0033151 | V(D)J recombination | 2/83 | 17/18723 | 0.002528879 | 0.017278993 | 0.012514477 | LEF1/TCF7 | 2 | BP |
| GO:0045591 | positive regulation of regulatory T cell differentiation | 2/83 | 17/18723 | 0.002528879 | 0.017278993 | 0.012514477 | HLA-DRA/HLA-DRB1 | 2 | BP |
| GO:0070206 | protein trimerization | 2/83 | 17/18723 | 0.002528879 | 0.017278993 | 0.012514477 | CD74/ALOX5AP | 2 | BP |
| GO:0090713 | immunological memory process | 2/83 | 17/18723 | 0.002528879 | 0.017278993 | 0.012514477 | HLA-DRA/HLA-DRB1 | 2 | BP |
| GO:0070265 | necrotic cell death | 3/83 | 62/18723 | 0.002631496 | 0.017908793 | 0.012970616 | BIRC3/TMEM123/FASLG | 3 | BP |
| GO:0002223 | stimulatory C-type lectin receptor signaling pathway | 2/83 | 18/18723 | 0.002836822 | 0.018971089 | 0.013739994 | KLRC2/KLRD1 | 2 | BP |
| GO:0034162 | toll-like receptor 9 signaling pathway | 2/83 | 18/18723 | 0.002836822 | 0.018971089 | 0.013739994 | HAVCR2/PTPN22 | 2 | BP |
| GO:1990840 | response to lectin | 2/83 | 18/18723 | 0.002836822 | 0.018971089 | 0.013739994 | KLRC2/KLRD1 | 2 | BP |
| GO:1990858 | cellular response to lectin | 2/83 | 18/18723 | 0.002836822 | 0.018971089 | 0.013739994 | KLRC2/KLRD1 | 2 | BP |
| GO:0007163 | establishment or maintenance of cell polarity | 5/83 | 218/18723 | 0.002842898 | 0.018971089 | 0.013739994 | CCR7/RIPOR2/RACK1/CRTAM/CCL4 | 5 | BP |
| GO:0048524 | positive regulation of viral process | 3/83 | 65/18723 | 0.003010411 | 0.019962203 | 0.014457818 | CD74/HLA-DRB1/CCL5 | 3 | BP |
| GO:0071219 | cellular response to molecule of bacterial origin | 5/83 | 221/18723 | 0.0030147 | 0.019962203 | 0.014457818 | TNIP3/HAVCR2/CCL3/PTPN22/CCL5 | 5 | BP |
| GO:0002274 | myeloid leukocyte activation | 5/83 | 223/18723 | 0.00313333 | 0.020306808 | 0.014707401 | HAVCR2/CCL3/CD74/CST7/CCL5 | 5 | BP |
| GO:0042093 | T-helper cell differentiation | 3/83 | 66/18723 | 0.003143767 | 0.020306808 | 0.014707401 | LEF1/HLA-DRA/HLA-DRB1 | 3 | BP |
| GO:0050918 | positive chemotaxis | 3/83 | 66/18723 | 0.003143767 | 0.020306808 | 0.014707401 | S1PR1/CCL3/CCL5 | 3 | BP |
| GO:0002483 | antigen processing and presentation of endogenous peptide antigen | 2/83 | 19/18723 | 0.003161468 | 0.020306808 | 0.014707401 | HLA-DRA/HLA-DRB1 | 2 | BP |
| GO:0045091 | regulation of single stranded viral RNA replication via double stranded DNA intermediate | 2/83 | 19/18723 | 0.003161468 | 0.020306808 | 0.014707401 | APOBEC3C/APOBEC3G | 2 | BP |
| GO:0070233 | negative regulation of T cell apoptotic process | 2/83 | 19/18723 | 0.003161468 | 0.020306808 | 0.014707401 | IL7R/CCL5 | 2 | BP |
| GO:1903978 | regulation of microglial cell activation | 2/83 | 19/18723 | 0.003161468 | 0.020306808 | 0.014707401 | CCL3/CST7 | 2 | BP |
| GO:2001185 | regulation of CD8-positive, alpha-beta T cell activation | 2/83 | 19/18723 | 0.003161468 | 0.020306808 | 0.014707401 | CRTAM/PTPN22 | 2 | BP |
| GO:0042130 | negative regulation of T cell proliferation | 3/83 | 67/18723 | 0.003280701 | 0.020915994 | 0.01514861 | HAVCR2/CRTAM/HLA-DRB1 | 3 | BP |
| GO:0046635 | positive regulation of alpha-beta T cell activation | 3/83 | 67/18723 | 0.003280701 | 0.020915994 | 0.01514861 | PTPN22/HLA-DRA/HLA-DRB1 | 3 | BP |
| GO:0002294 | CD4-positive, alpha-beta T cell differentiation involved in immune response | 3/83 | 68/18723 | 0.003421243 | 0.021571439 | 0.015623323 | LEF1/HLA-DRA/HLA-DRB1 | 3 | BP |
| GO:0009988 | cell-cell recognition | 3/83 | 68/18723 | 0.003421243 | 0.021571439 | 0.015623323 | CCR7/HAVCR2/PRF1 | 3 | BP |
| GO:0046637 | regulation of alpha-beta T cell differentiation | 3/83 | 68/18723 | 0.003421243 | 0.021571439 | 0.015623323 | HLA-DRA/ZNF683/HLA-DRB1 | 3 | BP |
| GO:0002577 | regulation of antigen processing and presentation | 2/83 | 20/18723 | 0.003502666 | 0.021728248 | 0.015736893 | CCR7/CD74 | 2 | BP |
| GO:0016553 | base conversion or substitution editing | 2/83 | 20/18723 | 0.003502666 | 0.021728248 | 0.015736893 | APOBEC3C/APOBEC3G | 2 | BP |
| GO:0039535 | regulation of RIG-I signaling pathway | 2/83 | 20/18723 | 0.003502666 | 0.021728248 | 0.015736893 | BIRC3/OASL | 2 | BP |
| GO:0039692 | single stranded viral RNA replication via double stranded DNA intermediate | 2/83 | 20/18723 | 0.003502666 | 0.021728248 | 0.015736893 | APOBEC3C/APOBEC3G | 2 | BP |
| GO:0071356 | cellular response to tumor necrosis factor | 5/83 | 229/18723 | 0.003509461 | 0.021728248 | 0.015736893 | BIRC3/CCL3/CCL4L1/CCL4/CCL5 | 5 | BP |
| GO:0002287 | alpha-beta T cell activation involved in immune response | 3/83 | 69/18723 | 0.003565423 | 0.021916486 | 0.015873226 | LEF1/HLA-DRA/HLA-DRB1 | 3 | BP |

| | | | | | | | | | |
|---|---|---|---|---|---|---|---|---|---|
| GO:0002293 | alpha-beta T cell differentiation involved in immune response | 3/83 | 69/18723 | 0.003565423 | 0.021916486 | 0.015873226 | LEF1/HLA-DRA/HLA-DRB1 | 3 | BP |
| GO:0097553 | calcium ion transmembrane import into cytosol | 4/83 | 142/18723 | 0.003666759 | 0.022458897 | 0.016266073 | CCR7/CCL3/F2R/FASLG | 4 | BP |
| GO:0045123 | cellular extravasation | 3/83 | 70/18723 | 0.003713269 | 0.022662836 | 0.016413777 | SELL/RIPOR2/ITGA1 | 3 | BP |
| GO:0030010 | establishment of cell polarity | 4/83 | 143/18723 | 0.003759727 | 0.022784211 | 0.016501684 | CCR7/RIPOR2/RACK1/CRTAM | 4 | BP |
| GO:0070555 | response to interleukin-1 | 4/83 | 143/18723 | 0.003759727 | 0.022784211 | 0.016501684 | CCL3/CCL4L1/CCL4/CCL5 | 4 | BP |
| GO:0042454 | ribonucleoside catabolic process | 2/83 | 21/18723 | 0.003860266 | 0.023175357 | 0.016784976 | APOBEC3C/APOBEC3G | 2 | BP |
| GO:0070269 | pyroptosis | 2/83 | 21/18723 | 0.003860266 | 0.023175357 | 0.016784976 | GZMB/GZMA | 2 | BP |
| GO:0045824 | negative regulation of innate immune response | 3/83 | 71/18723 | 0.003864812 | 0.023175357 | 0.016784976 | HAVCR2/KLRC1/KLRD1 | 3 | BP |
| GO:0002220 | innate immune response activating cell surface receptor signaling pathway | 2/83 | 22/18723 | 0.004234119 | 0.025039705 | 0.018135248 | KLRC2/KLRD1 | 2 | BP |
| GO:0032069 | regulation of nuclease activity | 2/83 | 22/18723 | 0.004234119 | 0.025039705 | 0.018135248 | OASL/GZMA | 2 | BP |
| GO:0045061 | thymic T cell selection | 2/83 | 22/18723 | 0.004234119 | 0.025039705 | 0.018135248 | CCR7/CD74 | 2 | BP |
| GO:2000114 | regulation of establishment of cell polarity | 2/83 | 22/18723 | 0.004234119 | 0.025039705 | 0.018135248 | RIPOR2/RACK1 | 2 | BP |
| GO:0002292 | T cell differentiation involved in immune response | 3/83 | 75/18723 | 0.00450849 | 0.026479661 | 0.01917815 | LEF1/HLA-DRA/HLA-DRB1 | 3 | BP |
| GO:0033077 | T cell differentiation in thymus | 3/83 | 75/18723 | 0.00450849 | 0.026479661 | 0.01917815 | IL7R/CCR7/CD74 | 3 | BP |
| GO:0002758 | innate immune response-activating signal transduction | 2/83 | 23/18723 | 0.004624079 | 0.027065856 | 0.019602707 | KLRC2/KLRD1 | 2 | BP |
| GO:0071216 | cellular response to biotic stimulus | 5/83 | 246/18723 | 0.004750831 | 0.027713182 | 0.020071539 | TNIP3/HAVCR2/CCL3/PTPN22/CCL5 | 5 | BP |
| GO:0006213 | pyrimidine nucleoside metabolic process | 2/83 | 24/18723 | 0.005029999 | 0.029053198 | 0.021042059 | APOBEC3C/APOBEC3G | 2 | BP |
| GO:0039531 | regulation of viral-induced cytoplasmic pattern recognition receptor signaling pathway | 2/83 | 24/18723 | 0.005029999 | 0.029053198 | 0.021042059 | BIRC3/OASL | 2 | BP |
| GO:0006919 | activation of cysteine-type endopeptidase activity involved in apoptotic process | 3/83 | 78/18723 | 0.00503137 | 0.029053198 | 0.021042059 | RACK1/F2R/FASLG | 3 | BP |
| GO:0034612 | response to tumor necrosis factor | 5/83 | 253/18723 | 0.005342995 | 0.030749115 | 0.022270343 | BIRC3/CCL3/CCL4L1/CCL4/CCL5 | 5 | BP |
| GO:0032272 | negative regulation of protein polymerization | 3/83 | 80/18723 | 0.005399436 | 0.030970007 | 0.022430326 | ADD3/CAPG/TBCD | 3 | BP |
| GO:0032878 | regulation of establishment or maintenance of cell polarity | 2/83 | 25/18723 | 0.005451732 | 0.031062193 | 0.022497092 | RIPOR2/RACK1 | 2 | BP |
| GO:0070423 | nucleotide-binding oligomerization domain containing signaling pathway | 2/83 | 25/18723 | 0.005451732 | 0.031062193 | 0.022497092 | BIRC3/PTPN22 | 2 | BP |
| GO:0060402 | calcium ion transport into cytosol | 4/83 | 160/18723 | 0.00559347 | 0.031764239 | 0.023005556 | CCR7/CCL3/F2R/FASLG | 4 | BP |
| GO:0002407 | dendritic cell chemotaxis | 2/83 | 26/18723 | 0.005889134 | 0.032898584 | 0.023827116 | CCR7/CCL5 | 2 | BP |
| GO:0009164 | nucleoside catabolic process | 2/83 | 26/18723 | 0.005889134 | 0.032898584 | 0.023827116 | APOBEC3C/APOBEC3G | 2 | BP |
| GO:0035872 | nucleotide-binding domain, leucine rich repeat containing receptor signaling pathway | 2/83 | 26/18723 | 0.005889134 | 0.032898584 | 0.023827116 | BIRC3/PTPN22 | 2 | BP |
| GO:0051016 | barbed-end actin filament capping | 2/83 | 26/18723 | 0.005889134 | 0.032898584 | 0.023827116 | ADD3/CAPG | 2 | BP |
| GO:1905523 | positive regulation of macrophage migration | 2/83 | 26/18723 | 0.005889134 | 0.032898584 | 0.023827116 | CCL3/CCL5 | 2 | BP |
| GO:0051494 | negative regulation of cytoskeleton organization | 4/83 | 163/18723 | 0.005969307 | 0.03308964 | 0.02396549 | S1PR1/ADD3/CAPG/TBCD | 4 | BP |
| GO:0043367 | CD4-positive, alpha-beta T cell differentiation | 3/83 | 83/18723 | 0.005981218 | 0.03308964 | 0.02396549 | LEF1/HLA-DRA/HLA-DRB1 | 3 | BP |
| GO:0050672 | negative regulation of lymphocyte proliferation | 3/83 | 83/18723 | 0.005981218 | 0.03308964 | 0.02396549 | HAVCR2/CRTAM/HLA-DRB1 | 3 | BP |
| GO:0032945 | negative regulation of mononuclear cell proliferation | 3/83 | 84/18723 | 0.006183149 | 0.033987502 | 0.024615775 | HAVCR2/CRTAM/HLA-DRB1 | 3 | BP |
| GO:0098586 | cellular response to virus | 3/83 | 84/18723 | 0.006183149 | 0.033987502 | 0.024615775 | BIRC3/OASL/CCL5 | 3 | BP |
| GO:0032635 | interleukin-6 production | 4/83 | 165/18723 | 0.006229005 | 0.034021479 | 0.024640384 | HAVCR2/CD74/F2R/PTPN22 | 4 | BP |
| GO:0032675 | regulation of interleukin-6 production | 4/83 | 165/18723 | 0.006229005 | 0.034021479 | 0.024640384 | HAVCR2/CD74/F2R/PTPN22 | 4 | BP |
| GO:0010528 | regulation of transposition | 2/83 | 27/18723 | 0.006342062 | 0.034096039 | 0.024694384 | APOBEC3C/APOBEC3G | 2 | BP |
| GO:0010529 | negative regulation of transposition | 2/83 | 27/18723 | 0.006342062 | 0.034096039 | 0.024694384 | APOBEC3C/APOBEC3G | 2 | BP |
| GO:0032703 | negative regulation of interleukin-2 production | 2/83 | 27/18723 | 0.006342062 | 0.034096039 | 0.024694384 | HAVCR2/LAG3 | 2 | BP |
| GO:0036037 | CD8-positive, alpha-beta T cell activation | 2/83 | 27/18723 | 0.006342062 | 0.034096039 | 0.024694384 | CRTAM/PTPN22 | 2 | BP |
| GO:0039529 | RIG-I signaling pathway | 2/83 | 27/18723 | 0.006342062 | 0.034096039 | 0.024694384 | BIRC3/OASL | 2 | BP |
| GO:1902904 | negative regulation of supramolecular fiber organization | 4/83 | 167/18723 | 0.006496123 | 0.034815157 | 0.025215213 | S1PR1/ADD3/CAPG/TBCD | 4 | BP |
| GO:0002675 | positive regulation of acute inflammatory response | 2/83 | 28/18723 | 0.006810372 | 0.03604873 | 0.02610864 | CCR7/ALOX5AP | 2 | BP |
| GO:0007205 | protein kinase C-activating G protein-coupled receptor signaling pathway | 2/83 | 28/18723 | 0.006810372 | 0.03604873 | 0.02610864 | DGKA/F2R | 2 | BP |
| GO:0010818 | T cell chemotaxis | 2/83 | 28/18723 | 0.006810372 | 0.03604873 | 0.02610864 | CCL3/CCL5 | 2 | BP |
| GO:0080111 | DNA demethylation | 2/83 | 28/18723 | 0.006810372 | 0.03604873 | 0.02610864 | APOBEC3C/APOBEC3G | 2 | BP |
| GO:0048872 | homeostasis of number of cells | 5/83 | 272/18723 | 0.007211461 | 0.038054327 | 0.027561213 | IL7R/CCR7/CD74/F2R/ID2 | 5 | BP |
| GO:0001916 | positive regulation of T cell mediated cytotoxicity | 2/83 | 29/18723 | 0.007293923 | 0.038254063 | 0.027705873 | HLA-DRA/HLA-DRB1 | 2 | BP |
| GO:1903902 | positive regulation of viral life cycle | 2/83 | 29/18723 | 0.007293923 | 0.038254063 | 0.027705873 | CD74/HLA-DRB1 | 2 | BP |
| GO:0070664 | negative regulation of leukocyte proliferation | 3/83 | 90/18723 | 0.007480217 | 0.039111503 | 0.028326883 | HAVCR2/CRTAM/HLA-DRB1 | 3 | BP |
| GO:1905475 | regulation of protein localization to membrane | 4/83 | 175/18723 | 0.007640557 | 0.039828434 | 0.028846127 | LDLRAP1/RACK1/ABI3/GZMB | 4 | BP |
| GO:0043032 | positive regulation of macrophage activation | 2/83 | 30/18723 | 0.007792574 | 0.040375421 | 0.029242287 | HAVCR2/CCL3 | 2 | BP |
| GO:0051491 | positive regulation of filopodium assembly | 2/83 | 30/18723 | 0.007792574 | 0.040375421 | 0.029242287 | CCR7/RIPOR2 | 2 | BP |
| GO:0002230 | positive regulation of defense response to virus by host | 2/83 | 31/18723 | 0.008306184 | 0.042522704 | 0.030797477 | PTPN22/APOBEC3G | 2 | BP |
| GO:0002323 | natural killer cell activation involved in immune response | 2/83 | 31/18723 | 0.008306184 | 0.042522704 | 0.030797477 | KLRC2/ZNF683 | 2 | BP |
| GO:0034656 | nucleobase-containing small molecule catabolic process | 2/83 | 31/18723 | 0.008306184 | 0.042522704 | 0.030797477 | APOBEC3C/APOBEC3G | 2 | BP |
| GO:0035510 | DNA dealkylation | 2/83 | 31/18723 | 0.008306184 | 0.042522704 | 0.030797477 | APOBEC3C/APOBEC3G | 2 | BP |
| GO:0048660 | regulation of smooth muscle cell proliferation | 4/83 | 180/18723 | 0.0084193 | 0.042973513 | 0.03112398 | S1PR1/LDLRAP1/ID2/CCL5 | 4 | BP |
| GO:0060401 | cytosolic calcium ion transport | 4/83 | 182/18723 | 0.008744834 | 0.044502641 | 0.032231467 | CCR7/CCL3/F2R/FASLG | 4 | BP |
| GO:0032196 | transposition | 2/83 | 32/18723 | 0.008834615 | 0.044562836 | 0.032275063 | APOBEC3C/APOBEC3G | 2 | BP |
| GO:0043372 | positive regulation of CD4-positive, alpha-beta T cell differentiation | 2/83 | 32/18723 | 0.008834615 | 0.044562836 | 0.032275063 | HLA-DRA/HLA-DRB1 | 2 | BP |
| GO:0050850 | positive regulation of calcium-mediated signaling | 2/83 | 32/18723 | 0.008834615 | 0.044562836 | 0.032275063 | CCL3/CCL4 | 2 | BP |
| GO:0048659 | smooth muscle cell proliferation | 4/83 | 184/18723 | 0.009078509 | 0.045658776 | 0.033068808 | S1PR1/LDLRAP1/ID2/CCL5 | 4 | BP |
| GO:1901216 | positive regulation of neuron death | 3/83 | 97/18723 | 0.009182584 | 0.046047168 | 0.033350105 | CCL3/FASLG/ITGA1 | 3 | BP |
| GO:0036336 | dendritic cell migration | 2/83 | 33/18723 | 0.009377728 | 0.046888638 | 0.033959548 | CCR7/CCL5 | 2 | BP |
| GO:0043473 | pigmentation | 3/83 | 98/18723 | 0.009442716 | 0.047076329 | 0.034095485 | RACK1/LYST/CD63 | 3 | BP |
| GO:0042100 | B cell proliferation | 3/83 | 99/18723 | 0.009707126 | 0.048254264 | 0.034948616 | IL7R/LEF1/CD74 | 3 | BP |
| GO:0039528 | cytoplasmic pattern recognition receptor signaling pathway in response to virus | 2/83 | 34/18723 | 0.009935385 | 0.048963178 | 0.035462053 | BIRC3/OASL | 2 | BP |
| GO:0051482 | positive regulation of cytosolic calcium ion concentration involved in phospholipase C-activating G protein-coupled signaling pathway | 2/83 | 34/18723 | 0.009935385 | 0.048963178 | 0.035462053 | S1PR1/F2R | 2 | BP |
| GO:0070232 | regulation of T cell apoptotic process | 2/83 | 34/18723 | 0.009935385 | 0.048963178 | 0.035462053 | IL7R/CCL5 | 2 | BP |
| GO:0044728 | DNA methylation or demethylation | 3/83 | 100/18723 | 0.009975829 | 0.049021624 | 0.035504383 | MYC/APOBEC3C/APOBEC3G | 3 | BP |
| GO:0042613 | MHC class II protein complex | 8/85 | 17/19550 | 2.14489E-15 | 2.53098E-13 | 1.80623E-13 | HLA-DQB1/CD74/HLA-DQA1/HLA-DPB1/HLA-DRA/HLA-DPA1/HLA-DRB5/HLA-DRB1 | 8 | CC |
| GO:0042611 | MHC protein complex | 8/85 | 25/19550 | 9.2787E-14 | 5.47443E-12 | 3.90682E-12 | HLA-DQB1/CD74/HLA-DQA1/HLA-DPB1/HLA-DRA/HLA-DPA1/HLA-DRB5/HLA-DRB1 | 8 | CC |
| GO:0071556 | integral component of lumenal side of endoplasmic reticulum membrane | 8/85 | 29/19550 | 3.63086E-13 | 1.0711E-11 | 7.64392E-12 | HLA-DQB1/CD74/HLA-DQA1/HLA-DPB1/HLA-DRA/HLA-DPA1/HLA-DRB5/HLA-DRB1 | 8 | CC |
| GO:0098553 | lumenal side of endoplasmic reticulum membrane | 8/85 | 29/19550 | 3.63086E-13 | 1.0711E-11 | 7.64392E-12 | HLA-DQB1/CD74/HLA-DQA1/HLA-DPB1/HLA-DRA/HLA-DPA1/HLA-DRB5/HLA-DRB1 | 8 | CC |
| GO:0030669 | clathrin-coated endocytic vesicle membrane | 10/85 | 72/19550 | 6.02408E-13 | 1.42168E-11 | 1.01458E-11 | IL7R/LDLRAP1/HLA-DQB1/CD74/HLA-DQA1/HLA-DPB1/HLA-DRA/HLA-DPA1/HLA-DRB5/HLA-DRB1 | 10 | CC |
| GO:0098576 | lumenal side of membrane | 8/85 | 36/19550 | 2.49771E-12 | 4.91217E-11 | 3.50556E-11 | HLA-DQB1/CD74/HLA-DQA1/HLA-DPB1/HLA-DRA/HLA-DPA1/HLA-DRB5/HLA-DRB1 | 8 | CC |
| GO:0045334 | clathrin-coated endocytic vesicle | 10/85 | 91/19550 | 6.75581E-12 | 1.13884E-10 | 8.12729E-11 | IL7R/LDLRAP1/HLA-DQB1/CD74/HLA-DQA1/HLA-DPB1/HLA-DRA/HLA-DPA1/HLA-DRB5/HLA-DRB1 | 10 | CC |

| | | | | | | | | | |
|---|---|---|---|---|---|---|---|---|---|
| GO:0030665 | clathrin-coated vesicle membrane | 10/85 | 116/19550 | 7.85613E-11 | 1.15878E-09 | 8.26961E-10 | IL7R/LDLRAP1/HLA-DQB1/CD74/HLA-DQA1/HLA-DPB1/HLA-DRA/HLA-DPA1/HLA-DRB5/HLA-DRB1 | 10 | CC |
| GO:0012507 | ER to Golgi transport vesicle membrane | 8/85 | 62/19550 | 2.54745E-10 | 3.34E-09 | 2.38358E-09 | HLA-DQB1/CD74/HLA-DQA1/HLA-DPB1/HLA-DRA/HLA-DPA1/HLA-DRB5/HLA-DRB1 | 8 | CC |
| GO:0030662 | coated vesicle membrane | 10/85 | 181/19550 | 6.19264E-09 | 7.30732E-08 | 5.21485E-08 | IL7R/LDLRAP1/HLA-DQB1/CD74/HLA-DQA1/HLA-DPB1/HLA-DRA/HLA-DPA1/HLA-DRB5/HLA-DRB1 | 10 | CC |
| GO:0030134 | COPII-coated ER to Golgi transport vesicle | 8/85 | 94/19550 | 7.49616E-09 | 8.04134E-08 | 5.73869E-08 | HLA-DQB1/CD74/HLA-DQA1/HLA-DPB1/HLA-DRA/HLA-DPA1/HLA-DRB5/HLA-DRB1 | 8 | CC |
| GO:0030666 | endocytic vesicle membrane | 10/85 | 193/19550 | 1.14642E-08 | 1.11378E-07 | 7.94846E-08 | IL7R/LDLRAP1/HLA-DQB1/CD74/HLA-DQA1/HLA-DPB1/HLA-DRA/HLA-DPA1/HLA-DRB5/HLA-DRB1 | 10 | CC |
| GO:0032588 | trans-Golgi network membrane | 8/85 | 100/19550 | 1.22704E-08 | 1.11378E-07 | 7.94846E-08 | HLA-DQB1/CD74/HLA-DQA1/HLA-DPB1/HLA-DRA/HLA-DPA1/HLA-DRB5/HLA-DRB1 | 8 | CC |
| GO:0030136 | clathrin-coated vesicle | 10/85 | 196/19550 | 1.32852E-08 | 1.11975E-07 | 7.99108E-08 | IL7R/LDLRAP1/HLA-DQB1/CD74/HLA-DQA1/HLA-DPB1/HLA-DRA/HLA-DPA1/HLA-DRB5/HLA-DRB1 | 10 | CC |
| GO:0001772 | immunological synapse | 6/85 | 44/19550 | 3.49234E-08 | 2.74731E-07 | 1.96061E-07 | HAVCR2/CRTAM/HLA-DRA/HLA-DRB1/GZMB/GZMA | 6 | CC |
| GO:0030176 | integral component of endoplasmic reticulum membrane | 8/85 | 162/19550 | 5.23623E-07 | 3.86172E-06 | 2.75591E-06 | HLA-DQB1/CD74/HLA-DQA1/HLA-DPB1/HLA-DRA/HLA-DPA1/HLA-DRB5/HLA-DRB1 | 8 | CC |
| GO:0030135 | coated vesicle | 10/85 | 299/19550 | 6.85299E-07 | 4.75678E-06 | 3.39467E-06 | IL7R/LDLRAP1/HLA-DQB1/CD74/HLA-DQA1/HLA-DPB1/HLA-DRA/HLA-DPA1/HLA-DRB5/HLA-DRB1 | 10 | CC |
| GO:0031227 | intrinsic component of endoplasmic reticulum membrane | 8/85 | 170/19550 | 7.54994E-07 | 4.94941E-06 | 3.53214E-06 | HLA-DQB1/CD74/HLA-DQA1/HLA-DPB1/HLA-DRA/HLA-DPA1/HLA-DRB5/HLA-DRB1 | 8 | CC |
| GO:0030658 | transport vesicle membrane | 8/85 | 212/19550 | 3.94104E-06 | 2.44759E-05 | 1.74672E-05 | HLA-DQB1/CD74/HLA-DQA1/HLA-DPB1/HLA-DRA/HLA-DPA1/HLA-DRB5/HLA-DRB1 | 8 | CC |
| GO:0005802 | trans-Golgi network | 8/85 | 259/19550 | 1.69969E-05 | 0.000100282 | 7.1566E-05 | HLA-DQB1/CD74/HLA-DQA1/HLA-DPB1/HLA-DRA/HLA-DPA1/HLA-DRB5/HLA-DRB1 | 8 | CC |
| GO:0005770 | late endosome | 7/85 | 282/19550 | 0.00022801 | 0.001281201 | 0.000914327 | CD74/F2R/HLA-DRA/CD63/HLA-DRB5/HLA-DRB1/CST7 | 7 | CC |
| GO:1990907 | beta-catenin-TCF complex | 2/85 | 13/19550 | 0.001412541 | 0.007576359 | 0.005406857 | LEF1/TCF7 | 2 | CC |
| GO:0005771 | multivesicular body | 3/85 | 64/19550 | 0.002729707 | 0.014004585 | 0.009994351 | CD74/CD63/CST7 | 3 | CC |
| GO:0042827 | platelet dense granule | 2/85 | 21/19550 | 0.003718115 | 0.017863661 | 0.012748375 | CD63/CTSW | 2 | CC |
| GO:0031902 | late endosome membrane | 4/85 | 146/19550 | 0.003784674 | 0.017863661 | 0.012748375 | HLA-DRA/CD63/HLA-DRB5/HLA-DRB1 | 4 | CC |
| GO:0005775 | vacuolar lumen | 4/85 | 174/19550 | 0.007014789 | 0.031836351 | 0.022719965 | PLAC8/CD74/CTSD/FASLG | 4 | CC |
| GO:0008305 | integrin complex | 2/85 | 31/19550 | 0.008004576 | 0.03498296 | 0.024965538 | ITGAE/ITGA1 | 2 | CC |
| GO:0043202 | lysosomal lumen | 3/85 | 97/19550 | 0.008718139 | 0.036740727 | 0.026219966 | CD74/CTSD/FASLG | 3 | CC |
| GO:0031904 | endosome lumen | 2/85 | 35/19550 | 0.010128059 | 0.041210722 | 0.029409971 | CD63/PRF1 | 2 | CC |
| GO:0098636 | protein complex involved in cell adhesion | 2/85 | 36/19550 | 0.010693793 | 0.042062252 | 0.030017664 | ITGAE/ITGA1 | 2 | CC |
| GO:0042470 | melanosome | 3/85 | 109/19550 | 0.011960827 | 0.04410555 | 0.031475861 | CAPG/CTSD/CD63 | 3 | CC |
| GO:0048770 | pigment granule | 3/85 | 109/19550 | 0.011960827 | 0.04410555 | 0.031475861 | CAPG/CTSD/CD63 | 3 | CC |
| GO:0023023 | MHC protein complex binding | 12/85 | 36/18368 | 4.91416E-20 | 1.02214E-17 | 7.55228E-18 | HLA-DQB1/CD74/KLRC2/HLA-DQA1/KLRC1/HLA-DPB1/HLA-DRA/HLA-DPA1/HLA-DRB5/HLA-DRB1/KLRD1/CD8A | 12 | MF |
| GO:0140375 | immune receptor activity | 13/85 | 144/18368 | 1.11143E-13 | 1.15589E-11 | 8.54048E-12 | IL7R/CCR7/HLA-DQB1/CD74/IL2RB/KLRC2/HLA-DQA1/KLRC1/HLA-DRA/HLA-DPA1/CXCR6/HLA-DRB1/KLRD1 | 13 | MF |
| GO:0023026 | MHC class II protein complex binding | 8/85 | 27/18368 | 3.10172E-13 | 2.15052E-11 | 1.58895E-11 | HLA-DQB1/CD74/HLA-DQA1/HLA-DPB1/HLA-DRA/HLA-DPA1/HLA-DRB5/HLA-DRB1 | 8 | MF |
| GO:0042605 | peptide antigen binding | 7/85 | 36/18368 | 2.64561E-10 | 1.37572E-08 | 1.01647E-08 | HLA-DQB1/HLA-DQA1/HLA-DPB1/HLA-DRA/HLA-DPA1/HLA-DRB5/HLA-DRB1 | 7 | MF |
| GO:0003823 | antigen binding | 11/85 | 171/18368 | 3.87534E-10 | 1.61214E-08 | 1.19116E-08 | IL7R/HLA-DQB1/KLRC2/HLA-DQA1/LAG3/HLA-DPB1/HLA-DRA/HLA-DPA1/HLA-DRB5/HLA-DRB1/KLRD1 | 11 | MF |
| GO:0032395 | MHC class II receptor activity | 5/85 | 10/18368 | 4.66129E-10 | 1.61591E-08 | 1.19394E-08 | HLA-DQB1/HLA-DQA1/HLA-DRA/HLA-DPA1/HLA-DRB1 | 5 | MF |
| GO:0042287 | MHC protein binding | 5/85 | 40/18368 | 1.09145E-06 | 3.24316E-05 | 2.39626E-05 | CD74/LAG3/KLRD1/CD8B/CD8A | 5 | MF |
| GO:0030246 | carbohydrate binding | 8/85 | 271/18368 | 3.65141E-05 | 0.000949367 | 0.000701455 | SELL/KLRC3/KLRC2/CD69/KLRC1/HLA-DRA/HLA-DRB1/KLRD1 | 8 | MF |
| GO:0048020 | CCR chemokine receptor binding | 4/85 | 48/18368 | 7.11519E-05 | 0.001605851 | 0.001186509 | CCL3/CCL4L1/CCL4/CCL5 | 4 | MF |
| GO:0008009 | chemokine activity | 4/85 | 49/18368 | 7.72044E-05 | 0.001605851 | 0.001186509 | CCL3/CCL4L1/CCL4/CCL5 | 4 | MF |
| GO:0004896 | cytokine receptor activity | 5/85 | 97/18368 | 8.69517E-05 | 0.001644178 | 0.001214828 | IL7R/CCR7/CD74/IL2RB/CXCR6 | 5 | MF |
| GO:0042379 | chemokine receptor binding | 4/85 | 72/18368 | 0.000345771 | 0.005993358 | 0.004428291 | CCL3/CCL4L1/CCL4/CCL5 | 4 | MF |
| GO:0047844 | deoxycytidine deaminase activity | 2/85 | 10/18368 | 0.000929693 | 0.014875091 | 0.010990705 | APOBEC3C/APOBEC3G | 2 | MF |
| GO:0042608 | T cell receptor binding | 2/85 | 11/18368 | 0.001132877 | 0.016831323 | 0.012436099 | HLA-DRA/HLA-DRB1 | 2 | MF |
| GO:0004126 | cytidine deaminase activity | 2/85 | 12/18368 | 0.00135537 | 0.018503773 | 0.013671816 | APOBEC3C/APOBEC3G | 2 | MF |
| GO:0015026 | coreceptor activity | 3/85 | 48/18368 | 0.001423367 | 0.018503773 | 0.013671816 | CXCR6/CD8B/CD8A | 3 | MF |
| GO:0016004 | phospholipase activator activity | 2/85 | 16/18368 | 0.002434859 | 0.029791216 | 0.022011729 | CCL3/CCL5 | 2 | MF |
| GO:0060229 | lipase activator activity | 2/85 | 18/18368 | 0.003085859 | 0.035658814 | 0.026347099 | CCL3/CCL5 | 2 | MF |
| GO:0003746 | translation elongation factor activity | 2/85 | 19/18368 | 0.003438567 | 0.03581164 | 0.026460018 | EEF1G/EEF2 | 2 | MF |
| GO:0016505 | peptidase activator activity involved in apoptotic process | 2/85 | 20/18368 | 0.003809185 | 0.03581164 | 0.026460018 | MAL/RACK1 | 2 | MF |
| GO:0042288 | MHC class I protein binding | 2/85 | 20/18368 | 0.003809185 | 0.03581164 | 0.026460018 | CD8B/CD8A | 2 | MF |
| GO:0019843 | rRNA binding | 3/85 | 68/18368 | 0.003859296 | 0.03581164 | 0.026460018 | RPS5/EEF2/RPLP0 | 3 | MF |
| GO:0019955 | cytokine binding | 4/85 | 139/18368 | 0.003959941 | 0.03581164 | 0.026460018 | CCR7/CD74/IL2RB/CXCR6 | 4 | MF |
| GO:0140416 | transcription regulator inhibitor activity | 2/85 | 21/18368 | 0.004197545 | 0.036378724 | 0.026879017 | LEF1/ID2 | 2 | MF |
| GO:0005125 | cytokine activity | 5/85 | 235/18368 | 0.004698685 | 0.039093057 | 0.028884546 | CCL3/FASLG/CCL4L1/CCL4/CCL5 | 5 | MF |
| GO:0016493 | C-C chemokine receptor activity | 2/85 | 23/18368 | 0.005026818 | 0.040214541 | 0.029713173 | CCR7/CXCR6 | 2 | MF |
| GO:0001965 | G-protein alpha-subunit binding | 2/85 | 24/18368 | 0.005467397 | 0.040614952 | 0.030009023 | F2R/RGS1 | 2 | MF |
| GO:0019957 | C-C chemokine binding | 2/85 | 24/18368 | 0.005467397 | 0.040614952 | 0.030009023 | CCR7/CXCR6 | 2 | MF |
| GO:0030247 | polysaccharide binding | 2/85 | 25/18368 | 0.005925053 | 0.042496929 | 0.031399553 | HLA-DRA/HLA-DRB1 | 2 | MF |
| GO:0001637 | G protein-coupled chemoattractant receptor activity | 2/85 | 26/18368 | 0.00639962 | 0.042939383 | 0.031726467 | CCR7/CXCR6 | 2 | MF |
| GO:0004950 | chemokine receptor activity | 2/85 | 26/18368 | 0.00639962 | 0.042939383 | 0.031726467 | CCR7/CXCR6 | 2 | MF |
| GO:0001540 | amyloid-beta binding | 3/85 | 84/18368 | 0.006959236 | 0.045235033 | 0.033422646 | LDLRAP1/CD74/ITM2C | 3 | MF |
| GO:0046625 | sphingolipid binding | 2/85 | 29/18368 | 0.007923172 | 0.049939993 | 0.036898983 | SELL/S1PR1 | 2 | MF |

**Supplementary Table 9: GO Enrichment Analysis Results for Highly Weighted Genes Associated with Trajectory 2 of NSCLC (Benjamini–Hochberg-adjusted P value < 0.05)**

| ID | Description | GeneRatio | BgRatio | pvalue | p.adjust | qvalue | geneID | Count | ONTOLOGY |
|---|---|---|---|---|---|---|---|---|---|
| GO:0072678 | T cell migration | 8/87 | 66/18723 | 7.21636E-10 | 1.34441E-06 | 1.08169E-06 | GPR183/ITGAL/ITGB7/MYO1G/RHOA/SPN/RIPOR2/S1PR1 | 8 | BP |
| GO:0071674 | mononuclear cell migration | 11/87 | 196/18723 | 1.74902E-09 | 1.62921E-06 | 1.31085E-06 | GPR183/TBX21/CD47/ITGAL/ITGB7/MYO1G/RHOA/SPN/RIPOR2/S1PR1/CX3CR1 | 11 | BP |
| GO:0072676 | lymphocyte migration | 9/87 | 117/18723 | 3.62929E-09 | 2.25379E-06 | 1.81337E-06 | GPR183/TBX21/ITGAL/ITGB7/MYO1G/RHOA/SPN/RIPOR2/S1PR1 | 9 | BP |
| GO:0110053 | regulation of actin filament organization | 12/87 | 278/18723 | 5.9724E-09 | 2.78164E-06 | 2.23808E-06 | S100A10/MTSS1/CD47/ARPC2/ADD3/RHOA/ACTR3/ARPC4/PXN/FLNA/S1PR1/PLEK | 12 | BP |
| GO:0032233 | positive regulation of actin filament bundle assembly | 7/87 | 63/18723 | 1.6398E-08 | 6.10989E-06 | 4.91594E-06 | S100A10/MTSS1/CD47/RHOA/PXN/FLNA/PLEK | 7 | BP |
| GO:0032231 | regulation of actin filament bundle assembly | 8/87 | 105/18723 | 3.02544E-08 | 8.17475E-06 | 6.57731E-06 | S100A10/MTSS1/CD47/RHOA/PXN/FLNA/S1PR1/PLEK | 8 | BP |
| GO:0045123 | cellular extravasation | 7/87 | 70/18723 | 3.46105E-08 | 8.17475E-06 | 6.57731E-06 | ITGB1/CD47/ITGAL/ITGB7/SPN/RIPOR2/CX3CR1 | 7 | BP |
| GO:0007229 | integrin-mediated signaling pathway | 8/87 | 107/18723 | 3.51036E-08 | 8.17475E-06 | 6.57731E-06 | ITGB1/CD47/ITGAL/ITGB7/ITGB2/FGR/FLNA/PLEK | 8 | BP |
| GO:1902905 | positive regulation of supramolecular fiber organization | 10/87 | 209/18723 | 4.58687E-08 | 9.49481E-06 | 7.63941E-06 | S100A10/MTSS1/CD47/ARPC2/RHOA/ACTR3/ARPC4/PXN/FLNA/PLEK | 10 | BP |
| GO:0051495 | positive regulation of cytoskeleton organization | 10/87 | 226/18723 | 9.56368E-08 | 1.78171E-05 | 1.43355E-05 | S100A10/MTSS1/CD47/ARPC2/RHOA/ACTR3/ARPC4/PXN/FLNA/PLEK | 10 | BP |
| GO:0006968 | cellular defense response | 6/87 | 54/18723 | 1.82544E-07 | 3.09163E-05 | 2.48749E-05 | ITGB1/SPN/KLRG1/PRF1/CX3CR1/GNLY | 6 | BP |
| GO:0061756 | leukocyte adhesion to vascular endothelial cell | 6/87 | 57/18723 | 2.53643E-07 | 3.93781E-05 | 3.16831E-05 | ITGB1/ITGB7/RHOA/SPN/ITGB2/CX3CR1 | 6 | BP |
| GO:0051017 | actin filament bundle assembly | 8/87 | 157/18723 | 6.81621E-07 | 9.76815E-05 | 7.85934E-05 | S100A10/MTSS1/CD47/RHOA/PXN/FLNA/S1PR1/PLEK | 8 | BP |
| GO:0002456 | T cell mediated immunity | 7/87 | 109/18723 | 7.43441E-07 | 9.89308E-05 | 7.95985E-05 | TBX21/LILRB1/MYO1G/CD8A/CTSC/PRF1/KLRD1 | 7 | BP |
| GO:0061572 | actin filament bundle organization | 8/87 | 161/18723 | 8.24919E-07 | 0.000102455 | 8.2434E-05 | S100A10/MTSS1/CD47/RHOA/PXN/FLNA/S1PR1/PLEK | 8 | BP |
| GO:0051015 | actin filament binding | 9/91 | 217/18368 | 1.2868E-06 | 0.000150973 | 0.000128264 | ARPC2/SYNE1/ABI3/MYO1G/ADD3/MYO1F/ACTR3/ARPC4/FLNA | 9 | MF |
| GO:0042287 | MHC protein binding | 5/91 | 40/18368 | 1.53272E-06 | 0.000150973 | 0.000128264 | LILRB1/CD8B/CD8A/FCRL6/KLRD1 | 5 | MF |
| GO:0001909 | leukocyte mediated cytotoxicity | 7/87 | 124/18723 | 1.77565E-06 | 0.000206753 | 0.000166351 | LILRB1/SLAMF7/CTSC/GZMB/PRF1/CX3CR1/KLRD1 | 7 | BP |
| GO:0001906 | cell killing | 8/87 | 188/18723 | 2.64346E-06 | 0.000289692 | 0.000233083 | LILRB1/SLAMF7/CTSC/GZMB/PRF1/CX3CR1/GNLY/KLRD1 | 8 | BP |
| GO:0007163 | establishment or maintenance of cell polarity | 8/87 | 218/18723 | 7.88382E-06 | 0.000815976 | 0.000656524 | ITGB1/CYTH1/RHOA/SPN/SLC9A3R1/ACTR3/RIPOR2/RAP1B | 8 | BP |
| GO:0034446 | substrate adhesion-dependent cell spreading | 6/87 | 108/18723 | 1.10712E-05 | 0.001085564 | 0.000873432 | S100A10/ARPC2/ITGB7/RHOA/PXN/FLNA | 6 | BP |
| GO:0002698 | negative regulation of immune effector process | 6/87 | 110/18723 | 1.23007E-05 | 0.001145806 | 0.000921902 | A2M/TBX21/CD47/LILRB1/CX3CR1/KLRD1 | 6 | BP |
| GO:0019835 | cytolysis | 4/87 | 32/18723 | 1.41587E-05 | 0.001256078 | 0.001010625 | GZMA/GZMB/PRF1/GZMH | 4 | BP |
| GO:0050901 | leukocyte tethering or rolling | 4/87 | 33/18723 | 1.60546E-05 | 0.001359535 | 0.001093865 | ITGB1/ITGB7/SPN/CX3CR1 | 4 | BP |
| GO:0030032 | lamellipodium assembly | 5/87 | 72/18723 | 2.11256E-05 | 0.001711172 | 0.001376788 | ITGB1/ARPC2/ABI3/ACTR3/S1PR1 | 5 | BP |
| GO:0023023 | MHC protein complex binding | 4/91 | 36/18368 | 2.94121E-05 | 0.001931393 | 0.001640884 | LILRB1/HLA-DPB1/CD8A/KLRD1 | 4 | MF |
| GO:0002695 | negative regulation of leukocyte activation | 7/87 | 187/18723 | 2.63961E-05 | 0.002049001 | 0.001648602 | TBX21/LILRB1/ID2/SPN/CST7/RIPOR2/FGR | 7 | BP |
| GO:0008305 | integrin complex | 4/91 | 31/19550 | 1.25534E-05 | 0.00211244 | 0.001664678 | ITGB1/ITGAL/ITGB7/ITGB2 | 4 | CC |
| GO:0098636 | protein complex involved in cell adhesion | 4/91 | 36/19550 | 2.30868E-05 | 0.00211244 | 0.001664678 | ITGB1/ITGAL/ITGB7/ITGB2 | 4 | CC |
| GO:0030027 | lamellipodium | 7/91 | 202/19550 | 4.4042E-05 | 0.002686561 | 0.002117106 | ITGB1/ARPC2/ABI3/MYO1G/RHOA/ACTR3/PXN | 7 | CC |
| GO:0002366 | leukocyte activation involved in immune response | 8/87 | 275/18723 | 4.19346E-05 | 0.003124969 | 0.002514313 | GPR183/TBX21/ITGAL/LILRB1/SPN/ITGB2/FGR/CX3CR1 | 8 | BP |
| GO:0002263 | cell activation involved in immune response | 8/87 | 279/18723 | 4.64374E-05 | 0.003327416 | 0.002677199 | GPR183/TBX21/ITGAL/LILRB1/SPN/ITGB2/FGR/CX3CR1 | 8 | BP |
| GO:2000177 | regulation of neural precursor cell proliferation | 5/87 | 87/18723 | 5.28251E-05 | 0.003549392 | 0.002855798 | ID2/RHOA/FLNA/ADGRG1/CX3CR1 | 5 | BP |
| GO:0030010 | establishment of cell polarity | 6/87 | 143/18723 | 5.42795E-05 | 0.003549392 | 0.002855798 | ITGB1/CYTH1/RHOA/SLC9A3R1/RIPOR2/RAP1B | 6 | BP |
| GO:0050866 | negative regulation of cell activation | 7/87 | 210/18723 | 5.52509E-05 | 0.003549392 | 0.002855798 | TBX21/LILRB1/ID2/SPN/CST7/RIPOR2/FGR | 7 | BP |
| GO:0097581 | lamellipodium organization | 5/87 | 90/18723 | 6.21498E-05 | 0.0038595 | 0.003105307 | ITGB1/ARPC2/ABI3/ACTR3/S1PR1 | 5 | BP |
| GO:0051492 | regulation of stress fiber assembly | 5/87 | 91/18723 | 6.55243E-05 | 0.003877154 | 0.003119512 | S100A10/CD47/RHOA/PXN/S1PR1 | 5 | BP |
| GO:0001774 | microglial cell activation | 4/87 | 47/18723 | 6.65963E-05 | 0.003877154 | 0.003119512 | ITGB2/CTSC/CST7/CX3CR1 | 4 | BP |
| GO:0001913 | T cell mediated cytotoxicity | 4/87 | 49/18723 | 7.85507E-05 | 0.004434545 | 0.003567982 | LILRB1/CTSC/PRF1/KLRD1 | 4 | BP |
| GO:0015026 | coreceptor activity | 4/91 | 48/18368 | 9.28472E-05 | 0.004572723 | 0.003884921 | ITGB1/CD8B/CD8A/TGFBR3 | 4 | MF |
| GO:1903975 | regulation of glial cell migration | 3/87 | 19/18723 | 8.89839E-05 | 0.00485119 | 0.003903209 | GPR183/IDH2/CX3CR1 | 3 | BP |
| GO:0051250 | negative regulation of lymphocyte activation | 6/87 | 157/18723 | 9.11388E-05 | 0.00485119 | 0.003903209 | TBX21/LILRB1/ID2/SPN/RIPOR2/FGR | 6 | BP |
| GO:0051496 | positive regulation of stress fiber assembly | 4/87 | 52/18723 | 9.93093E-05 | 0.004863661 | 0.003913243 | S100A10/CD47/RHOA/PXN | 4 | BP |
| GO:0110020 | regulation of actomyosin structure organization | 5/87 | 100/18723 | 0.000102664 | 0.004863661 | 0.003913243 | S100A10/CD47/RHOA/PXN/S1PR1 | 5 | BP |
| GO:0007160 | cell-matrix adhesion | 7/87 | 233/18723 | 0.000106018 | 0.004863661 | 0.003913243 | S100A10/ITGB1/ITGAL/ITGB7/RHOA/ITGB2/PXN | 7 | BP |
| GO:0007188 | adenylate cyclase-modulating G protein-coupled receptor signaling pathway | 7/87 | 233/18723 | 0.000106018 | 0.004863661 | 0.003913243 | RGS1/ADRB2/SLC9A3R1/S1PR5/FLNA/S1PR1/ADGRG1 | 7 | BP |
| GO:0008347 | glial cell migration | 4/87 | 53/18723 | 0.000107037 | 0.004863661 | 0.003913243 | GPR183/IDH2/ADGRG1/CX3CR1 | 4 | BP |
| GO:0043113 | receptor clustering | 4/87 | 53/18723 | 0.000107037 | 0.004863661 | 0.003913243 | ITGAL/ITGB7/ITGB2/FLNA | 4 | BP |
| GO:0042288 | MHC class I protein binding | 3/91 | 20/18368 | 0.000126161 | 0.004970748 | 0.004223077 | LILRB1/CD8B/CD8A | 3 | MF |
| GO:0009898 | cytoplasmic side of plasma membrane | 6/91 | 172/19550 | 0.000152669 | 0.005961372 | 0.004697773 | RGS1/MTSS1/CYTH1/RHOA/FGR/LITAF | 6 | CC |
| GO:0001726 | ruffle | 6/91 | 178/19550 | 0.000183966 | 0.005961372 | 0.004697773 | MTSS1/ITGB1/RHOA/SLC9A3R1/FGR/PLEK | 6 | CC |
| GO:0031256 | leading edge membrane | 6/91 | 180/19550 | 0.000195455 | 0.005961372 | 0.004697773 | ITGB1/MYO1G/RHOA/RIPOR2/FGR/PLEK | 6 | CC |
| GO:0030038 | contractile actin filament bundle assembly | 5/87 | 106/18723 | 0.00013521 | 0.005858043 | 0.004713311 | S100A10/CD47/RHOA/PXN/S1PR1 | 5 | BP |
| GO:0043149 | stress fiber assembly | 5/87 | 106/18723 | 0.00013521 | 0.005858043 | 0.004713311 | S100A10/CD47/RHOA/PXN/S1PR1 | 5 | BP |
| GO:0031532 | actin cytoskeleton reorganization | 5/87 | 107/18723 | 0.000141323 | 0.005983755 | 0.004814457 | ABI3/RHOA/FLNA/S1PR1/PLEK | 5 | BP |
| GO:0002363 | alpha-beta T cell lineage commitment | 3/87 | 23/18723 | 0.000160461 | 0.006643078 | 0.00534494 | TBX21/RHOA/SPN | 3 | BP |
| GO:0001911 | negative regulation of leukocyte mediated cytotoxicity | 3/87 | 24/18723 | 0.000182769 | 0.007350463 | 0.005914094 | LILRB1/CX3CR1/KLRD1 | 3 | BP |
| GO:0034113 | heterotypic cell-cell adhesion | 4/87 | 61/18723 | 0.000185438 | 0.007350463 | 0.005914094 | ITGB1/CD47/ITGB7/ITGB2 | 4 | BP |
| GO:0002286 | T cell activation involved in immune response | 5/87 | 114/18723 | 0.000190256 | 0.007384298 | 0.005941317 | GPR183/TBX21/ITGAL/LILRB1/SPN | 5 | BP |
| GO:0071675 | regulation of mononuclear cell migration | 5/87 | 115/18723 | 0.000198183 | 0.007534983 | 0.006062557 | CD47/RHOA/SPN/RIPOR2/CX3CR1 | 5 | BP |
| GO:0002704 | negative regulation of leukocyte mediated immunity | 4/87 | 63/18723 | 0.000210177 | 0.007831185 | 0.006300877 | TBX21/LILRB1/CX3CR1/KLRD1 | 4 | BP |
| GO:0004252 | serine-type endopeptidase activity | 6/91 | 174/18368 | 0.000227077 | 0.007455707 | 0.006334263 | GZMK/GZMA/CTSC/PRSS23/GZMB/GZMH | 6 | MF |
| GO:0098562 | cytoplasmic side of membrane | 6/91 | 197/19550 | 0.000317702 | 0.008305635 | 0.006545137 | RGS1/MTSS1/CYTH1/RHOA/FGR/LITAF | 6 | CC |
| GO:0002710 | negative regulation of T cell mediated immunity | 3/87 | 26/18723 | 0.000233211 | 0.00849845 | 0.00683775 | TBX21/LILRB1/KLRD1 | 3 | BP |
| GO:0071677 | positive regulation of mononuclear cell migration | 4/87 | 65/18723 | 0.000237208 | 0.00849845 | 0.00683775 | CD47/RHOA/SPN/CX3CR1 | 4 | BP |
| GO:0008064 | regulation of actin polymerization or depolymerization | 6/87 | 188/18723 | 0.000243511 | 0.008559635 | 0.006886979 | ARPC2/ADD3/RHOA/ACTR3/ARPC4/PLEK | 6 | BP |
| GO:0030832 | regulation of actin filament length | 6/87 | 189/18723 | 0.000250568 | 0.008644604 | 0.006955344 | ARPC2/ADD3/RHOA/ACTR3/ARPC4/PLEK | 6 | BP |

| | | | | | | | | | |
|---|---|---|---|---|---|---|---|---|---|
| GO:0002433 | immune response-regulating cell surface receptor signaling pathway involved in phagocytosis | 3/87 | 27/18723 | 0.000261483 | 0.008698979 | 0.006999094 | CD47/MYO1G/FGR | 3 | BP |
| GO:0038096 | Fc-gamma receptor signaling pathway involved in phagocytosis | 3/87 | 27/18723 | 0.000261483 | 0.008698979 | 0.006999094 | CD47/MYO1G/FGR | 3 | BP |
| GO:0042267 | natural killer cell mediated cytotoxicity | 4/87 | 68/18723 | 0.000282323 | 0.009216474 | 0.007415464 | LILRB1/SLAMF7/GZMB/KLRD1 | 4 | BP |
| GO:0031342 | negative regulation of cell killing | 3/87 | 28/18723 | 0.00029188 | 0.009216474 | 0.007415464 | LILRB1/CX3CR1/KLRD1 | 3 | BP |
| GO:0038094 | Fc-gamma receptor signaling pathway | 3/87 | 28/18723 | 0.00029188 | 0.009216474 | 0.007415464 | CD47/MYO1G/FGR | 3 | BP |
| GO:0031032 | actomyosin structure organization | 6/87 | 196/18723 | 0.000304599 | 0.009457786 | 0.007609621 | S100A10/ITGB1/CD47/RHOA/PXN/S1PR1 | 6 | BP |
| GO:0002360 | T cell lineage commitment | 3/87 | 29/18723 | 0.000324468 | 0.00990956 | 0.007973113 | TBX21/RHOA/SPN | 3 | BP |
| GO:0008236 | serine-type peptidase activity | 6/91 | 191/18368 | 0.00037441 | 0.009402221 | 0.007987995 | GZMK/GZMA/CTSC/PRSS23/GZMB/GZMH | 6 | MF |
| GO:0017171 | serine hydrolase activity | 6/91 | 195/18368 | 0.000418039 | 0.009402221 | 0.007987995 | GZMK/GZMA/CTSC/PRSS23/GZMB/GZMH | 6 | MF |
| GO:0019902 | phosphatase binding | 6/91 | 196/18368 | 0.000429543 | 0.009402221 | 0.007987995 | LILRB1/SLC9A3R1/CTSC/PPP1CA/PXN/FCRL6 | 6 | MF |
| GO:0002228 | natural killer cell mediated immunity | 4/87 | 71/18723 | 0.000333291 | 0.01001485 | 0.008057827 | LILRB1/SLAMF7/GZMB/KLRD1 | 4 | BP |
| GO:0090162 | establishment of epithelial cell polarity | 3/87 | 31/18723 | 0.000396478 | 0.011724414 | 0.009433322 | CYTH1/RHOA/SLC9A3R1 | 3 | BP |
| GO:0019956 | chemokine binding | 3/91 | 33/18368 | 0.000576318 | 0.011353461 | 0.00964574 | A2M/ITGB1/CX3CR1 | 3 | MF |
| GO:0002687 | positive regulation of leukocyte migration | 5/87 | 135/18723 | 0.00041635 | 0.012119682 | 0.00975135 | CD47/RHOA/SPN/RIPOR2/CX3CR1 | 5 | BP |
| GO:0098858 | actin-based cell projection | 6/91 | 221/19550 | 0.000584284 | 0.01336549 | 0.010532482 | ITGB1/MYO1G/SPN/SLC9A3R1/MYO1F/RIPOR2 | 6 | CC |
| GO:0002431 | Fc receptor mediated stimulatory signaling pathway | 3/87 | 33/18723 | 0.000478025 | 0.013700942 | 0.011023613 | CD47/MYO1G/FGR | 3 | BP |
| GO:0008154 | actin polymerization or depolymerization | 6/87 | 218/18723 | 0.000535932 | 0.015127911 | 0.012171735 | ARPC2/ADD3/RHOA/ACTR3/ARPC4/PLEK | 6 | BP |
| GO:0061351 | neural precursor cell proliferation | 5/87 | 145/18723 | 0.000576971 | 0.016043232 | 0.012908191 | ID2/RHOA/FLNA/ADGRG1/CX3CR1 | 5 | BP |
| GO:0007189 | adenylate cyclase-activating G protein-coupled receptor signaling pathway | 5/87 | 146/18723 | 0.00059527 | 0.016308643 | 0.013121738 | ADRB2/SLC9A3R1/S1PR5/S1PR1/ADGRG1 | 5 | BP |
| GO:0005902 | microvillus | 4/91 | 91/19550 | 0.000862001 | 0.017129317 | 0.013498513 | MYO1G/SPN/SLC9A3R1/MYO1F | 4 | CC |
| GO:0098862 | cluster of actin-based cell projections | 5/91 | 161/19550 | 0.000936028 | 0.017129317 | 0.013498513 | ADD3/SLC9A3R1/ACTR3/RIPOR2/FLNA | 5 | CC |
| GO:0002703 | regulation of leukocyte mediated immunity | 6/87 | 226/18723 | 0.00064758 | 0.017484661 | 0.014067948 | TBX21/LILRB1/ITGB2/FGR/CX3CR1/KLRD1 | 6 | BP |
| GO:0032587 | ruffle membrane | 4/91 | 99/19550 | 0.00118029 | 0.018619158 | 0.01467256 | ITGB1/RHOA/FGR/PLEK | 4 | CC |
| GO:0031234 | extrinsic component of cytoplasmic side of plasma membrane | 4/91 | 101/19550 | 0.001271166 | 0.018619158 | 0.01467256 | RGS1/CYTH1/RHOA/FGR | 4 | CC |
| GO:0019897 | extrinsic component of plasma membrane | 5/91 | 174/19550 | 0.001322672 | 0.018619158 | 0.01467256 | RGS1/S100A10/CYTH1/RHOA/FGR | 5 | CC |
| GO:0005903 | brush border | 4/91 | 106/19550 | 0.00151959 | 0.019191179 | 0.015123334 | ADD3/SLC9A3R1/ACTR3/FLNA | 4 | CC |
| GO:0030175 | filopodium | 4/91 | 107/19550 | 0.001573047 | 0.019191179 | 0.015123334 | ITGB1/MYO1G/SLC9A3R1/RIPOR2 | 4 | CC |
| GO:0008097 | 5S rRNA binding | 2/91 | 10/18368 | 0.001064555 | 0.019065219 | 0.016197541 | RPL5/GTF3A | 2 | MF |
| GO:0034314 | Arp2/3 complex-mediated actin nucleation | 3/87 | 39/18723 | 0.000784773 | 0.020781089 | 0.016720214 | ARPC2/ACTR3/ARPC4 | 3 | BP |
| GO:0002699 | positive regulation of immune effector process | 6/87 | 235/18723 | 0.000793914 | 0.020781089 | 0.016720214 | TBX21/LILRB1/SPON2/ITGB2/FGR/KLRD1 | 6 | BP |
| GO:0046631 | alpha-beta T cell activation | 5/87 | 156/18723 | 0.000803134 | 0.020781089 | 0.016720214 | GPR183/TBX21/LILRB1/RHOA/SPN | 5 | BP |
| GO:0031334 | positive regulation of protein-containing complex assembly | 6/87 | 237/18723 | 0.00082965 | 0.02117313 | 0.017035645 | ARPC2/RHOA/ACTR3/ARPC4/RAP1B/PLEK | 6 | BP |
| GO:1900026 | positive regulation of substrate adhesion-dependent cell spreading | 3/87 | 41/18723 | 0.000909274 | 0.022392411 | 0.018016664 | S100A10/ARPC2/FLNA | 3 | BP |
| GO:0001915 | negative regulation of T cell mediated cytotoxicity | 2/87 | 10/18723 | 0.000937524 | 0.022392411 | 0.018016664 | LILRB1/KLRD1 | 2 | BP |
| GO:0002291 | T cell activation via T cell receptor contact with antigen bound to MHC molecule on antigen presenting cell | 2/87 | 10/18723 | 0.000937524 | 0.022392411 | 0.018016664 | ITGAL/LILRB1 | 2 | BP |
| GO:0021932 | hindbrain radial glia guided cell migration | 2/87 | 10/18723 | 0.000937524 | 0.022392411 | 0.018016664 | ITGB1/FLNA | 2 | BP |
| GO:0040015 | negative regulation of multicellular organism growth | 2/87 | 10/18723 | 0.000937524 | 0.022392411 | 0.018016664 | ADRB2/PLAC8 | 2 | BP |
| GO:0019864 | IgG binding | 2/91 | 11/18368 | 0.001296932 | 0.021291302 | 0.01808879 | FCGR3B/FCGR3A | 2 | MF |
| GO:0010591 | regulation of lamellipodium assembly | 3/87 | 42/18723 | 0.000975946 | 0.022727333 | 0.018286139 | ARPC2/ABI3/ACTR3 | 3 | BP |
| GO:2000404 | regulation of T cell migration | 3/87 | 42/18723 | 0.000975946 | 0.022727333 | 0.018286139 | RHOA/SPN/RIPOR2 | 3 | BP |
| GO:0061383 | trabecula morphogenesis | 3/87 | 43/18723 | 0.001045636 | 0.024049638 | 0.019350049 | RHOA/TGFBR3/S1PR1 | 3 | BP |
| GO:0034767 | positive regulation of ion transmembrane transport | 5/87 | 167/18723 | 0.001089749 | 0.024491012 | 0.019705173 | ITGB1/ADRB2/ARL6IP1/SLC9A3R1/FLNA | 5 | BP |
| GO:0150076 | neuroinflammatory response | 3/87 | 44/18723 | 0.001118398 | 0.024491012 | 0.019705173 | ITGB1/CTSC/CST7 | 3 | BP |
| GO:0002357 | defense response to tumor cell | 2/87 | 11/18723 | 0.001142404 | 0.024491012 | 0.019705173 | ABI3/PRF1 | 2 | BP |
| GO:0033625 | positive regulation of integrin activation | 2/87 | 11/18723 | 0.001142404 | 0.024491012 | 0.019705173 | RAP1B/PLEK | 2 | BP |
| GO:1903961 | positive regulation of anion transmembrane transport | 2/87 | 11/18723 | 0.001142404 | 0.024491012 | 0.019705173 | ITGB1/ARL6IP1 | 2 | BP |
| GO:0002444 | myeloid leukocyte mediated immunity | 4/87 | 99/18723 | 0.001169995 | 0.024491012 | 0.019705173 | SPON2/ITGB2/FGR/CX3CR1 | 4 | BP |
| GO:0030838 | positive regulation of actin filament polymerization | 4/87 | 99/18723 | 0.001169995 | 0.024491012 | 0.019705173 | ARPC2/RHOA/ACTR3/ARPC4 | 4 | BP |
| GO:0031341 | regulation of cell killing | 4/87 | 99/18723 | 0.001169995 | 0.024491012 | 0.019705173 | LILRB1/PRF1/CX3CR1/KLRD1 | 4 | BP |
| GO:0030833 | regulation of actin filament polymerization | 5/87 | 172/18723 | 0.001242515 | 0.025720065 | 0.020694055 | ARPC2/ADD3/RHOA/ACTR3/ARPC4 | 5 | BP |
| GO:0005200 | structural constituent of cytoskeleton | 4/91 | 103/18368 | 0.001715087 | 0.025990162 | 0.022080875 | ARPC2/ADD3/ACTR3/ARPC4 | 4 | MF |
| GO:0060907 | positive regulation of macrophage cytokine production | 2/87 | 12/18723 | 0.001366748 | 0.027980792 | 0.022513008 | LILRB1/SPON2 | 2 | BP |
| GO:0042116 | macrophage activation | 4/87 | 106/18723 | 0.001506469 | 0.030209363 | 0.02430609 | ITGB2/CTSC/CST7/CX3CR1 | 4 | BP |
| GO:0009409 | response to cold | 3/87 | 49/18723 | 0.00153003 | 0.030209363 | 0.02430609 | ADRB2/PLAC8/UCP2 | 3 | BP |
| GO:0050832 | defense response to fungus | 3/87 | 49/18723 | 0.00153003 | 0.030209363 | 0.02430609 | SPON2/CX3CR1/GNLY | 3 | BP |
| GO:0032640 | tumor necrosis factor production | 5/87 | 181/18723 | 0.001556682 | 0.030209363 | 0.02430609 | CD47/LILRB1/SPON2/SPN/CX3CR1 | 5 | BP |
| GO:0032680 | regulation of tumor necrosis factor production | 5/87 | 181/18723 | 0.001556682 | 0.030209363 | 0.02430609 | CD47/LILRB1/SPON2/SPN/CX3CR1 | 5 | BP |
| GO:0010958 | regulation of amino acid import across plasma membrane | 2/87 | 13/18723 | 0.001610375 | 0.030222722 | 0.024316838 | ITGB1/ARL6IP1 | 2 | BP |
| GO:1903789 | regulation of amino acid transmembrane transport | 2/87 | 13/18723 | 0.001610375 | 0.030222722 | 0.024316838 | ITGB1/ARL6IP1 | 2 | BP |
| GO:0038093 | Fc receptor signaling pathway | 3/87 | 50/18723 | 0.001622261 | 0.030222722 | 0.024316838 | CD47/MYO1G/FGR | 3 | BP |
| GO:0045058 | T cell selection | 3/87 | 50/18723 | 0.001622261 | 0.030222722 | 0.024316838 | TBX21/RHOA/SPN | 3 | BP |
| GO:0048872 | homeostasis of number of cells | 6/87 | 272/18723 | 0.00168066 | 0.031000686 | 0.024942778 | GPR183/ID2/KLF2/TGFBR3/ADGRG1/CX3CR1 | 6 | BP |
| GO:0071706 | tumor necrosis factor superfamily cytokine production | 5/87 | 186/18723 | 0.001754616 | 0.031736398 | 0.025534724 | CD47/LILRB1/SPON2/SPN/CX3CR1 | 5 | BP |
| GO:1903555 | regulation of tumor necrosis factor superfamily cytokine production | 5/87 | 186/18723 | 0.001754616 | 0.031736398 | 0.025534724 | CD47/LILRB1/SPON2/SPN/CX3CR1 | 5 | BP |
| GO:0043090 | amino acid import | 3/87 | 52/18723 | 0.001817 | 0.032238774 | 0.025938929 | ITGB1/ARL6IP1/SLC9A3R1 | 3 | BP |
| GO:0045010 | actin nucleation | 3/87 | 52/18723 | 0.001817 | 0.032238774 | 0.025938929 | ARPC2/ACTR3/ARPC4 | 3 | BP |
| GO:0046632 | alpha-beta T cell differentiation | 4/87 | 112/18723 | 0.001844179 | 0.032412312 | 0.026078556 | GPR183/TBX21/RHOA/SPN | 4 | BP |
| GO:0003376 | sphingosine-1-phosphate receptor signaling pathway | 2/87 | 14/18723 | 0.001873106 | 0.032613048 | 0.026240066 | S1PR5/S1PR1 | 2 | BP |
| GO:0002707 | negative regulation of lymphocyte mediated immunity | 3/87 | 53/18723 | 0.001919598 | 0.033113064 | 0.026642373 | TBX21/LILRB1/KLRD1 | 3 | BP |
| GO:0030041 | actin filament polymerization | 5/87 | 191/18723 | 0.001970438 | 0.033678224 | 0.027097094 | ARPC2/ADD3/RHOA/ACTR3/ARPC4 | 5 | BP |

| | | | | | | | | | |
|---|---|---|---|---|---|---|---|---|---|
| GO:0002823 | negative regulation of adaptive immune response based on somatic recombination of immune receptors built from immunoglobulin superfamily domains | 3/87 | 54/18723 | 0.002025739 | 0.033696005 | 0.0271114 | TBX21/LILRB1/KLRD1 | 3 | BP |
| GO:1902743 | regulation of lamellipodium organization | 3/87 | 54/18723 | 0.002025739 | 0.033696005 | 0.0271114 | ARPC2/ABI3/ACTR3 | 3 | BP |
| GO:2000179 | positive regulation of neural precursor cell proliferation | 3/87 | 54/18723 | 0.002025739 | 0.033696005 | 0.0271114 | FLNA/ADGRG1/CX3CR1 | 3 | BP |
| GO:0002285 | lymphocyte activation involved in immune response | 5/87 | 194/18723 | 0.002108909 | 0.034464015 | 0.027729332 | GPR183/TBX21/ITGAL/LILRB1/SPN | 5 | BP |
| GO:0050777 | negative regulation of immune response | 5/87 | 194/18723 | 0.002108909 | 0.034464015 | 0.027729332 | A2M/TBX21/LILRB1/FGR/KLRD1 | 5 | BP |
| GO:0051938 | L-glutamate import | 2/87 | 15/18723 | 0.002154761 | 0.034606201 | 0.027843733 | ITGB1/ARL6IP1 | 2 | BP |
| GO:0098712 | L-glutamate import across plasma membrane | 2/87 | 15/18723 | 0.002154761 | 0.034606201 | 0.027843733 | ITGB1/ARL6IP1 | 2 | BP |
| GO:0002886 | regulation of myeloid leukocyte mediated immunity | 3/87 | 56/18723 | 0.002248824 | 0.035504732 | 0.02856668 | ITGB2/FGR/CX3CR1 | 3 | BP |
| GO:0015800 | acidic amino acid transport | 3/87 | 56/18723 | 0.002248824 | 0.035504732 | 0.02856668 | ITGB1/ARL6IP1/SLC9A3R1 | 3 | BP |
| GO:0045125 | bioactive lipid receptor activity | 2/91 | 15/18368 | 0.002444242 | 0.034393982 | 0.029220642 | S1PR5/S1PR1 | 2 | MF |
| GO:1900024 | regulation of substrate adhesion-dependent cell spreading | 3/87 | 57/18723 | 0.002365849 | 0.037038464 | 0.029800702 | S100A10/ARPC2/FLNA | 3 | BP |
| GO:0002693 | positive regulation of cellular extravasation | 2/87 | 16/18723 | 0.002455163 | 0.037491543 | 0.030165244 | CD47/RIPOR2 | 2 | BP |
| GO:0021535 | cell migration in hindbrain | 2/87 | 16/18723 | 0.002455163 | 0.037491543 | 0.030165244 | ITGB1/FLNA | 2 | BP |
| GO:0090520 | sphingolipid mediated signaling pathway | 2/87 | 16/18723 | 0.002455163 | 0.037491543 | 0.030165244 | S1PR5/S1PR1 | 2 | BP |
| GO:0050868 | negative regulation of T cell activation | 4/87 | 122/18723 | 0.002518132 | 0.03814048 | 0.030687371 | TBX21/LILRB1/SPN/RIPOR2 | 4 | BP |
| GO:0002820 | negative regulation of adaptive immune response | 3/87 | 59/18723 | 0.002611069 | 0.039229208 | 0.031563348 | TBX21/LILRB1/KLRD1 | 3 | BP |
| GO:0009620 | response to fungus | 3/87 | 60/18723 | 0.002739342 | 0.040063681 | 0.032234755 | SPON2/CX3CR1/GNLY | 3 | BP |
| GO:0010934 | macrophage cytokine production | 2/87 | 17/18723 | 0.002774136 | 0.040063681 | 0.032234755 | LILRB1/SPON2 | 2 | BP |
| GO:0010935 | regulation of macrophage cytokine production | 2/87 | 17/18723 | 0.002774136 | 0.040063681 | 0.032234755 | LILRB1/SPON2 | 2 | BP |
| GO:0033623 | regulation of integrin activation | 2/87 | 17/18723 | 0.002774136 | 0.040063681 | 0.032234755 | RAP1B/PLEK | 2 | BP |
| GO:0045198 | establishment of epithelial cell apical/basal polarity | 2/87 | 17/18723 | 0.002774136 | 0.040063681 | 0.032234755 | RHOA/SLC9A3R1 | 2 | BP |
| GO:2000401 | regulation of lymphocyte migration | 3/87 | 61/18723 | 0.00287144 | 0.041149945 | 0.03310875 | RHOA/SPN/RIPOR2 | 3 | BP |
| GO:0071222 | cellular response to lipopolysaccharide | 5/87 | 209/18723 | 0.002910107 | 0.041385713 | 0.033298446 | LILRB1/RHOA/SPON2/LITAF/CX3CR1 | 5 | BP |
| GO:0002685 | regulation of leukocyte migration | 5/87 | 210/18723 | 0.002970356 | 0.041922522 | 0.033730357 | CD47/RHOA/SPN/RIPOR2/CX3CR1 | 5 | BP |
| GO:0045428 | regulation of nitric oxide biosynthetic process | 3/87 | 62/18723 | 0.003007402 | 0.042126235 | 0.033894262 | CD47/KLF2/CX3CR1 | 3 | BP |
| GO:0002295 | T-helper cell lineage commitment | 2/87 | 18/18723 | 0.003111504 | 0.042623034 | 0.03429398 | TBX21/SPN | 2 | BP |
| GO:0045953 | negative regulation of natural killer cell mediated cytotoxicity | 2/87 | 18/18723 | 0.003111504 | 0.042623034 | 0.03429398 | LILRB1/KLRD1 | 2 | BP |
| GO:0071800 | podosome assembly | 2/87 | 18/18723 | 0.003111504 | 0.042623034 | 0.03429398 | RHOA/BIN2 | 2 | BP |
| GO:0005839 | proteasome core complex | 2/91 | 20/19550 | 0.003855424 | 0.044096407 | 0.034749542 | PSMA2/PSMB8 | 2 | CC |
| GO:0098631 | cell adhesion mediator activity | 3/91 | 59/18368 | 0.003129595 | 0.041102018 | 0.034919695 | ITGB1/CD47/PPP1CA | 3 | MF |
| GO:0080164 | regulation of nitric oxide metabolic process | 3/87 | 64/18723 | 0.00329106 | 0.044753605 | 0.036008212 | CD47/KLF2/CX3CR1 | 3 | BP |
| GO:0002716 | negative regulation of natural killer cell mediated immunity | 2/87 | 19/18723 | 0.003467095 | 0.045809915 | 0.036858106 | LILRB1/KLRD1 | 2 | BP |
| GO:0030050 | vesicle transport along actin filament | 2/87 | 19/18723 | 0.003467095 | 0.045809915 | 0.036858106 | MYO1G/MYO1F | 2 | BP |
| GO:0051957 | positive regulation of amino acid transport | 2/87 | 19/18723 | 0.003467095 | 0.045809915 | 0.036858106 | ITGB1/ARL6IP1 | 2 | BP |
| GO:1903978 | regulation of microglial cell activation | 2/87 | 19/18723 | 0.003467095 | 0.045809915 | 0.036858106 | CTSC/CST7 | 2 | BP |
| GO:0034764 | positive regulation of transmembrane transport | 5/87 | 219/18723 | 0.003553921 | 0.046626449 | 0.037515079 | ITGB1/ADRB2/ARL6IP1/SLC9A3R1/FLNA | 5 | BP |
| GO:0042093 | T-helper cell differentiation | 3/87 | 66/18723 | 0.003590598 | 0.046778212 | 0.037637186 | GPR183/TBX21/SPN | 3 | BP |
| GO:0071219 | cellular response to molecule of bacterial origin | 5/87 | 221/18723 | 0.00369404 | 0.047791645 | 0.038452582 | LILRB1/RHOA/SPON2/LITAF/CX3CR1 | 5 | BP |
| GO:0007266 | Rho protein signal transduction | 4/87 | 137/18723 | 0.00382016 | 0.048975039 | 0.039404726 | ITGB1/RHOA/RIPOR2/ADGRG1 | 4 | BP |
| GO:0002274 | myeloid leukocyte activation | 5/87 | 223/18723 | 0.003838087 | 0.048975039 | 0.039404726 | ITGB2/CTSC/CST7/FGR/CX3CR1 | 5 | BP |
| GO:0002294 | CD4-positive, alpha-beta T cell differentiation involved in immune response | 3/87 | 68/18723 | 0.003906289 | 0.049349987 | 0.039706405 | GPR183/TBX21/SPN | 3 | BP |
| GO:0032273 | positive regulation of protein polymerization | 4/87 | 138/18723 | 0.00392045 | 0.049349987 | 0.039706405 | ARPC2/RHOA/ACTR3/ARPC4 | 4 | BP |
