## supplementary tables for "MGPfact^XMBD^: A Model-Based Factorization Method for scRNA Data Unveils Bifurcating Transcriptional Modules Underlying Cell Fate Determination": supplementary_table_10_gse108989_enrich_go_combine.pdf

**Supplementary Table 10: GO Enrichment Analysis Results for Highly Weighted Genes Associated with Trajectory 1 of CRC (Benjamini–Hochberg-adjusted P value < 0.05)**

| ID | Description | GeneRatio | BgRatio | pvalue | p.adjust | qvalue | geneID | Count | ONTOLOGY |
|---|---|---|---|---|---|---|---|---|---|
| GO:0050870 | positive regulation of T cell activation | 19/88 | 216/18723 | 3.00383E-19 | 5.63819E-16 | 3.99984E-16 | IL7R/CCR7/LEF1/CD28/PTPN22/TNFSF4/CD74/HLA-DMA/VCAM1/HLA-DQB1/HLA-DQA1/IFNG/HLA-DPA1/HLA-DPB1/HAVCR2/HLA-DRB5/HLA-DRA/HLA-DRB1/CCL5 | 19 | BP |
| GO:0042613 | MHC class II protein complex | 9/89 | 17/19550 | 1.30739E-17 | 1.35968E-15 | 9.90861E-16 | CD74/HLA-DMA/HLA-DQB1/HLA-DQA1/HLA-DPA1/HLA-DPB1/HLA-DRB5/HLA-DRA/HLA-DRB1 | 9 | CC |
| GO:1903039 | positive regulation of leukocyte cell-cell adhesion | 19/88 | 239/18723 | 2.0523E-18 | 1.62158E-15 | 1.15038E-15 | IL7R/CCR7/LEF1/CD28/PTPN22/TNFSF4/CD74/HLA-DMA/VCAM1/HLA-DQB1/HLA-DQA1/IFNG/HLA-DPA1/HLA-DPB1/HAVCR2/HLA-DRB5/HLA-DRA/HLA-DRB1/CCL5 | 19 | BP |
| GO:0022409 | positive regulation of cell-cell adhesion | 20/88 | 284/18723 | 2.59176E-18 | 1.62158E-15 | 1.15038E-15 | IL7R/CCR7/LEF1/CD28/PTPN22/TNFSF4/CD74/HLA-DMA/VCAM1/HLA-DQB1/HLA-DQA1/IFNG/CXCL13/HLA-DPA1/HLA-DPB1/HAVCR2/HLA-DRB5/HLA-DRA/HLA-DRB1/CCL5 | 20 | BP |
| GO:0023023 | MHC protein complex binding | 11/89 | 36/18368 | 9.92323E-18 | 1.84572E-15 | 1.37881E-15 | CD74/HLA-DMA/HLA-DQB1/HLA-DQA1/HLA-DPA1/HLA-DPB1/KLRD1/HLA-DRB5/HLA-DRA/HLA-DRB1/CD8A | 11 | MF |
| GO:0042611 | MHC protein complex | 9/89 | 25/19550 | 1.06673E-15 | 5.54701E-14 | 4.04236E-14 | CD74/HLA-DMA/HLA-DQB1/HLA-DQA1/HLA-DPA1/HLA-DPB1/HLA-DRB5/HLA-DRA/HLA-DRB1 | 9 | CC |
| GO:0030217 | T cell differentiation | 18/88 | 257/18723 | 1.69074E-16 | 7.93382E-14 | 5.6284E-14 | IL7R/GPR183/CCR7/LEF1/CD28/TCF7/PTPN22/THEMIS/TNFSF4/CD74/ZNF683/EOMES/CRTAM/LAG3/IFNG/HLA-DRA/HLA-DRB1/CD8A | 18 | BP |
| GO:0001906 | cell killing | 16/88 | 188/18723 | 4.31397E-16 | 1.61946E-13 | 1.14888E-13 | IL7R/RAB27A/CTSC/SH2D1A/GNLY/CRTAM/LYST/LAG3/SLAMF7/IFNG/HAVCR2/KLRD1/PRF1/HLA-DRA/HLA-DRB1/GZMB | 16 | BP |
| GO:0001909 | leukocyte mediated cytotoxicity | 14/88 | 124/18723 | 6.26819E-16 | 1.9609E-13 | 1.3911E-13 | IL7R/RAB27A/CTSC/SH2D1A/CRTAM/LYST/LAG3/SLAMF7/HAVCR2/KLRD1/PRF1/HLA-DRA/HLA-DRB1/GZMB | 14 | BP |
| GO:0023026 | MHC class II protein complex binding | 9/89 | 27/18368 | 4.24051E-15 | 3.94368E-13 | 2.94604E-13 | CD74/HLA-DMA/HLA-DQB1/HLA-DQA1/HLA-DPA1/HLA-DPB1/HLA-DRB5/HLA-DRA/HLA-DRB1 | 9 | MF |
| GO:0002399 | MHC class II protein complex assembly | 8/88 | 16/18723 | 2.14572E-15 | 5.0344E-13 | 3.5715E-13 | HLA-DMA/HLA-DQB1/HLA-DQA1/HLA-DPA1/HLA-DPB1/HLA-DRB5/HLA-DRA/HLA-DRB1 | 8 | BP |
| GO:0002503 | peptide antigen assembly with MHC class II protein complex | 8/88 | 16/18723 | 2.14572E-15 | 5.0344E-13 | 3.5715E-13 | HLA-DMA/HLA-DQB1/HLA-DQA1/HLA-DPA1/HLA-DPB1/HLA-DRB5/HLA-DRA/HLA-DRB1 | 8 | BP |
| GO:0002501 | peptide antigen assembly with MHC protein complex | 8/88 | 18/18723 | 7.24012E-15 | 1.50997E-12 | 1.0712E-12 | HLA-DMA/HLA-DQB1/HLA-DQA1/HLA-DPA1/HLA-DPB1/HLA-DRB5/HLA-DRA/HLA-DRB1 | 8 | BP |
| GO:0019886 | antigen processing and presentation of exogenous peptide antigen via MHC class II | 9/88 | 30/18723 | 9.7041E-15 | 1.82146E-12 | 1.29218E-12 | CD74/HLA-DMA/HLA-DQB1/HLA-DQA1/HLA-DPA1/HLA-DPB1/HLA-DRB5/HLA-DRA/HLA-DRB1 | 9 | BP |
| GO:0002396 | MHC protein complex assembly | 8/88 | 19/18723 | 1.24581E-14 | 2.12581E-12 | 1.50809E-12 | HLA-DMA/HLA-DQB1/HLA-DQA1/HLA-DPA1/HLA-DPB1/HLA-DRB5/HLA-DRA/HLA-DRB1 | 8 | BP |
| GO:0002495 | antigen processing and presentation of peptide antigen via MHC class II | 9/88 | 34/18723 | 3.50384E-14 | 5.48059E-12 | 3.88803E-12 | CD74/HLA-DMA/HLA-DQB1/HLA-DQA1/HLA-DPA1/HLA-DPB1/HLA-DRB5/HLA-DRA/HLA-DRB1 | 9 | BP |
| GO:0002504 | antigen processing and presentation of peptide or polysaccharide antigen via MHC class II | 9/88 | 36/18723 | 6.24123E-14 | 9.01138E-12 | 6.39284E-12 | CD74/HLA-DMA/HLA-DQB1/HLA-DQA1/HLA-DPA1/HLA-DPB1/HLA-DRB5/HLA-DRA/HLA-DRB1 | 9 | BP |
| GO:0019882 | antigen processing and presentation | 12/88 | 106/18723 | 8.22131E-14 | 1.10224E-11 | 7.81951E-12 | CCR7/RAB27A/CD74/HLA-DMA/HLA-DQB1/HLA-DQA1/HLA-DPA1/HLA-DPB1/HLA-DRB5/HLA-DRA/HLA-DRB1/CD8A | 12 | BP |
| GO:0002478 | antigen processing and presentation of exogenous peptide antigen | 9/88 | 38/18723 | 1.07249E-13 | 1.34204E-11 | 9.52068E-12 | CD74/HLA-DMA/HLA-DQB1/HLA-DQA1/HLA-DPA1/HLA-DPB1/HLA-DRB5/HLA-DRA/HLA-DRB1 | 9 | BP |
| GO:0071556 | integral component of lumenal side of endoplasmic reticulum membrane | 8/89 | 29/19550 | 5.3081E-13 | 1.38011E-11 | 1.00574E-11 | CD74/HLA-DQB1/HLA-DQA1/HLA-DPA1/HLA-DPB1/HLA-DRB5/HLA-DRA/HLA-DRB1 | 8 | CC |
| GO:0098553 | lumenal side of endoplasmic reticulum membrane | 8/89 | 29/19550 | 5.3081E-13 | 1.38011E-11 | 1.00574E-11 | CD74/HLA-DQB1/HLA-DQA1/HLA-DPA1/HLA-DPB1/HLA-DRB5/HLA-DRA/HLA-DRB1 | 8 | CC |
| GO:0030669 | clathrin-coated endocytic vesicle membrane | 10/89 | 72/19550 | 9.67672E-13 | 2.01276E-11 | 1.46679E-11 | IL7R/LDLRAP1/CD74/HLA-DQB1/HLA-DQA1/HLA-DPA1/HLA-DPB1/HLA-DRB5/HLA-DRA/HLA-DRB1 | 10 | CC |
| GO:0098576 | lumenal side of membrane | 8/89 | 36/19550 | 3.64684E-12 | 6.32119E-11 | 4.60654E-11 | CD74/HLA-DQB1/HLA-DQA1/HLA-DPA1/HLA-DPB1/HLA-DRB5/HLA-DRA/HLA-DRB1 | 8 | CC |
| GO:0019884 | antigen processing and presentation of exogenous antigen | 9/88 | 47/18723 | 8.66313E-13 | 1.01629E-10 | 7.20977E-11 | CD74/HLA-DMA/HLA-DQB1/HLA-DQA1/HLA-DPA1/HLA-DPB1/HLA-DRB5/HLA-DRA/HLA-DRB1 | 9 | BP |
| GO:0002381 | immunoglobulin production involved in immunoglobulin-mediated immune response | 10/88 | 70/18723 | 9.79653E-13 | 1.08165E-10 | 7.67344E-11 | CD28/TNFSF4/HLA-DMA/HLA-DQB1/HLA-DQA1/HLA-DPA1/HLA-DPB1/HLA-DRB5/HLA-DRA/HLA-DRB1 | 10 | BP |
| GO:0045334 | clathrin-coated endocytic vesicle | 10/89 | 91/19550 | 1.08137E-11 | 1.60661E-10 | 1.17081E-10 | IL7R/LDLRAP1/CD74/HLA-DQB1/HLA-DQA1/HLA-DPA1/HLA-DPB1/HLA-DRB5/HLA-DRA/HLA-DRB1 | 10 | CC |
| GO:0045619 | regulation of lymphocyte differentiation | 13/88 | 174/18723 | 1.60909E-12 | 1.67792E-10 | 1.19035E-10 | IL7R/LEF1/CD28/TCF7/TNFSF4/ID2/CD74/ZNF683/CRTAM/LAG3/IFNG/HLA-DRA/HLA-DRB1 | 13 | BP |
| GO:0045580 | regulation of T cell differentiation | 12/88 | 146/18723 | 3.95472E-12 | 3.90685E-10 | 2.77159E-10 | IL7R/LEF1/CD28/TCF7/TNFSF4/CD74/ZNF683/CRTAM/LAG3/IFNG/HLA-DRA/HLA-DRB1 | 12 | BP |
| GO:0002285 | lymphocyte activation involved in immune response | 13/88 | 194/18723 | 6.44668E-12 | 6.05021E-10 | 4.29213E-10 | GPR183/LEF1/CD28/TNFSF4/RAB27A/CD244/CD74/ZNF683/EOMES/IFNG/HAVCR2/HLA-DRA/HLA-DRB1 | 13 | BP |
| GO:0048002 | antigen processing and presentation of peptide antigen | 9/88 | 62/18723 | 1.21812E-11 | 1.08877E-09 | 7.72395E-10 | CD74/HLA-DMA/HLA-DQB1/HLA-DQA1/HLA-DPA1/HLA-DPB1/HLA-DRB5/HLA-DRA/HLA-DRB1 | 9 | BP |
| GO:0030665 | clathrin-coated vesicle membrane | 10/89 | 116/19550 | 1.25165E-10 | 1.62714E-09 | 1.18577E-09 | IL7R/LDLRAP1/CD74/HLA-DQB1/HLA-DQA1/HLA-DPA1/HLA-DPB1/HLA-DRB5/HLA-DRA/HLA-DRB1 | 10 | CC |

| | | | | | | | | | |
|---|---|---|---|---|---|---|---|---|---|
| GO:0070374 | positive regulation of ERK1 and ERK2 cascade | 13/88 | 217/18723 | 2.65291E-11 | 2.26342E-09 | 1.60571E-09 | GPR183/CCR7/PTPN22/CCL3L1/CCL3L3/CD74/F2R/CCL3/HAVCR2/CCL4L1/HLA-DRB1/CCL4/CCL5 | 13 | BP |
| GO:0042267 | natural killer cell mediated cytotoxicity | 9/88 | 68/18723 | 2.89221E-11 | 2.3603E-09 | 1.67444E-09 | RAB27A/SH2D1A/CRTAM/LYST/LAG3/SLAMF7/HAVCR2/KLRD1/GZMB | 9 | BP |
| GO:0031341 | regulation of cell killing | 10/88 | 99/18723 | 3.44571E-11 | 2.69483E-09 | 1.91177E-09 | IL7R/SH2D1A/CRTAM/LAG3/IFNG/HAVCR2/KLRD1/PRF1/HLA-DRA/HLA-DRB1 | 10 | BP |
| GO:0002366 | leukocyte activation involved in immune response | 14/88 | 275/18723 | 3.80812E-11 | 2.85914E-09 | 2.02833E-09 | GPR183/LEF1/CD28/TNFSF4/RAB27A/CD244/CD74/ZNF683/EOMES/IFNG/CCL3/HAVCR2/HLA-DRA/HLA-DRB1 | 14 | BP |
| GO:0002228 | natural killer cell mediated immunity | 9/88 | 71/18723 | 4.32116E-11 | 3.09363E-09 | 2.19468E-09 | RAB27A/SH2D1A/CRTAM/LYST/LAG3/SLAMF7/HAVCR2/KLRD1/GZMB | 9 | BP |
| GO:0002263 | cell activation involved in immune response | 14/88 | 279/18723 | 4.6149E-11 | 3.09363E-09 | 2.19468E-09 | GPR183/LEF1/CD28/TNFSF4/RAB27A/CD244/CD74/ZNF683/EOMES/IFNG/CCL3/HAVCR2/HLA-DRA/HLA-DRB1 | 14 | BP |
| GO:1902105 | regulation of leukocyte differentiation | 14/88 | 279/18723 | 4.6149E-11 | 3.09363E-09 | 2.19468E-09 | IL7R/LEF1/CD28/TCF7/TNFSF4/ID2/CD74/ZNF683/CRTAM/LAG3/IFNG/CCL3/HLA-DRA/HLA-DRB1 | 14 | BP |
| GO:0030593 | neutrophil chemotaxis | 10/88 | 103/18723 | 5.13899E-11 | 3.32617E-09 | 2.35965E-09 | CCR7/CCL3L1/CCL3L3/CD74/ITGA1/CXCL13/CCL3/CCL4L1/CCL4/CCL5 | 10 | BP |
| GO:0012507 | ER to Golgi transport vesicle membrane | 8/89 | 62/19550 | 3.70189E-10 | 4.27774E-09 | 3.11738E-09 | CD74/HLA-DQB1/HLA-DQA1/HLA-DPA1/HLA-DPB1/HLA-DRB5/HLA-DRA/HLA-DRB1 | 8 | CC |
| GO:0046651 | lymphocyte proliferation | 14/88 | 288/18723 | 7.03376E-11 | 4.40079E-09 | 3.122E-09 | IL7R/GPR183/LEF1/CD28/PTPN22/TNFSF4/CD74/CRTAM/VCAM1/HLA-DPA1/HLA-DPB1/HAVCR2/HLA-DRB1/CCL5 | 14 | BP |
| GO:0030134 | COPII-coated ER to Golgi transport vesicle | 9/89 | 94/19550 | 4.30466E-10 | 4.47685E-09 | 3.26248E-09 | CTSC/CD74/HLA-DQB1/HLA-DQA1/HLA-DPA1/HLA-DPB1/HLA-DRB5/HLA-DRA/HLA-DRB1 | 9 | CC |
| GO:0032943 | mononuclear cell proliferation | 14/88 | 291/18723 | 8.0685E-11 | 4.88534E-09 | 3.46576E-09 | IL7R/GPR183/LEF1/CD28/PTPN22/TNFSF4/CD74/CRTAM/VCAM1/HLA-DPA1/HLA-DPB1/HAVCR2/HLA-DRB1/CCL5 | 14 | BP |
| GO:0002286 | T cell activation involved in immune response | 10/88 | 114/18723 | 1.42182E-10 | 8.33985E-09 | 5.91645E-09 | GPR183/LEF1/TNFSF4/RAB27A/CD74/EOMES/IFNG/HAVCR2/HLA-DRA/HLA-DRB1 | 10 | BP |
| GO:0046631 | alpha-beta T cell activation | 11/88 | 156/18723 | 1.73664E-10 | 9.87779E-09 | 7.00749E-09 | GPR183/LEF1/CD28/PTPN22/TNFSF4/ZNF683/EOMES/CRTAM/IFNG/HLA-DRA/HLA-DRB1 | 11 | BP |
| GO:0001772 | immunological synapse | 7/89 | 44/19550 | 1.06524E-09 | 1.00714E-08 | 7.33947E-09 | CD28/CRTAM/HAVCR2/HLA-DRA/HLA-DRB1/GZMA/GZMB | 7 | CC |
| GO:0072676 | lymphocyte migration | 10/88 | 117/18723 | 1.84226E-10 | 1.01704E-08 | 7.21506E-09 | GPR183/CCR7/CCL3L1/CCL3L3/CRTAM/CXCL13/CCL3/CCL4L1/CCL4/CCL5 | 10 | BP |
| GO:0030136 | clathrin-coated vesicle | 11/89 | 196/19550 | 1.43463E-09 | 1.24334E-08 | 9.06081E-09 | IL7R/LDLRAP1/RAB27A/CD74/HLA-DQB1/HLA-DQA1/HLA-DPA1/HLA-DPB1/HLA-DRB5/HLA-DRA/HLA-DRB1 | 11 | CC |
| GO:1990266 | neutrophil migration | 10/88 | 122/18723 | 2.79296E-10 | 1.49783E-08 | 1.06259E-08 | CCR7/CCL3L1/CCL3L3/CD74/ITGA1/CXCL13/CCL3/CCL4L1/CCL4/CCL5 | 10 | BP |
| GO:0070098 | chemokine-mediated signaling pathway | 9/88 | 88/18723 | 3.10726E-10 | 1.62009E-08 | 1.14932E-08 | CCR7/CCL3L1/CCL3L3/CXCR6/CXCL13/CCL3/CCL4L1/CCL4/CCL5 | 9 | BP |
| GO:0071621 | granulocyte chemotaxis | 10/88 | 125/18723 | 3.55357E-10 | 1.80272E-08 | 1.27888E-08 | CCR7/CCL3L1/CCL3L3/CD74/ITGA1/CXCL13/CCL3/CCL4L1/CCL4/CCL5 | 10 | BP |
| GO:0042605 | peptide antigen binding | 7/89 | 36/18368 | 3.67277E-10 | 2.27712E-08 | 1.70107E-08 | HLA-DQB1/HLA-DQA1/HLA-DPA1/HLA-DPB1/HLA-DRB5/HLA-DRA/HLA-DRB1 | 7 | MF |
| GO:0031343 | positive regulation of cell killing | 8/88 | 63/18723 | 5.39915E-10 | 2.66689E-08 | 1.89194E-08 | SH2D1A/CRTAM/LAG3/IFNG/KLRD1/PRF1/HLA-DRA/HLA-DRB1 | 8 | BP |
| GO:0032395 | MHC class II receptor activity | 5/89 | 10/18368 | 5.89311E-10 | 2.7403E-08 | 2.04708E-08 | HLA-DQB1/HLA-DQA1/HLA-DPA1/HLA-DRA/HLA-DRB1 | 5 | MF |
| GO:0048247 | lymphocyte chemotaxis | 8/88 | 64/18723 | 6.14701E-10 | 2.918E-08 | 2.07008E-08 | GPR183/CCL3L1/CCL3L3/CXCL13/CCL3/CCL4L1/CCL4/CCL5 | 8 | BP |
| GO:0050670 | regulation of lymphocyte proliferation | 12/88 | 225/18723 | 6.21843E-10 | 2.918E-08 | 2.07008E-08 | GPR183/CD28/PTPN22/TNFSF4/CD74/CRTAM/VCAM1/HLA-DPA1/HLA-DPB1/HAVCR2/HLA-DRB1/CCL5 | 12 | BP |
| GO:0032944 | regulation of mononuclear cell proliferation | 12/88 | 227/18723 | 6.88149E-10 | 3.15038E-08 | 2.23494E-08 | GPR183/CD28/PTPN22/TNFSF4/CD74/CRTAM/VCAM1/HLA-DPA1/HLA-DPB1/HAVCR2/HLA-DRB1/CCL5 | 12 | BP |
| GO:1990868 | response to chemokine | 9/88 | 97/18723 | 7.51038E-10 | 3.27837E-08 | 2.32573E-08 | CCR7/CCL3L1/CCL3L3/CXCR6/CXCL13/CCL3/CCL4L1/CCL4/CCL5 | 9 | BP |
| GO:1990869 | cellular response to chemokine | 9/88 | 97/18723 | 7.51038E-10 | 3.27837E-08 | 2.32573E-08 | CCR7/CCL3L1/CCL3L3/CXCR6/CXCL13/CCL3/CCL4L1/CCL4/CCL5 | 9 | BP |
| GO:0030595 | leukocyte chemotaxis | 12/88 | 230/18723 | 7.9963E-10 | 3.41115E-08 | 2.41993E-08 | GPR183/CCR7/CCL3L1/CCL3L3/CD74/ITGA1/LYST/CXCL13/CCL3/CCL4L1/CCL4/CCL5 | 12 | BP |
| GO:0050671 | positive regulation of lymphocyte proliferation | 10/88 | 137/18723 | 8.77695E-10 | 3.66096E-08 | 2.59715E-08 | GPR183/CD28/PTPN22/TNFSF4/CD74/VCAM1/HLA-DPA1/HLA-DPB1/HAVCR2/CCL5 | 10 | BP |
| GO:0032946 | positive regulation of mononuclear cell proliferation | 10/88 | 138/18723 | 9.42672E-10 | 3.84651E-08 | 2.72879E-08 | GPR183/CD28/PTPN22/TNFSF4/CD74/VCAM1/HLA-DPA1/HLA-DPB1/HAVCR2/CCL5 | 10 | BP |
| GO:0002699 | positive regulation of immune effector process | 12/88 | 235/18723 | 1.02214E-09 | 4.08202E-08 | 2.89586E-08 | CD28/PTPN22/TNFSF4/CD244/CD74/SH2D1A/CRTAM/LAG3/IFNG/KLRD1/HLA-DRA/HLA-DRB1 | 12 | BP |
| GO:0034341 | response to interferon-gamma | 10/88 | 141/18723 | 1.16402E-09 | 4.55181E-08 | 3.22914E-08 | CCL3L1/CCL3L3/CD74/FASLG/IFNG/CCL3/HLA-DPA1/CCL4L1/CCL4/CCL5 | 10 | BP |
| GO:0070663 | regulation of leukocyte proliferation | 12/88 | 245/18723 | 1.64218E-09 | 6.29056E-08 | 4.46264E-08 | GPR183/CD28/PTPN22/TNFSF4/CD74/CRTAM/VCAM1/HLA-DPA1/HLA-DPB1/HAVCR2/HLA-DRB1/CCL5 | 12 | BP |
| GO:0097530 | granulocyte migration | 10/88 | 148/18723 | 1.8693E-09 | 7.01736E-08 | 4.97825E-08 | CCR7/CCL3L1/CCL3L3/CD74/ITGA1/CXCL13/CCL3/CCL4L1/CCL4/CCL5 | 10 | BP |
| GO:0140375 | immune receptor activity | 10/89 | 144/18368 | 1.91955E-09 | 7.14074E-08 | 5.33434E-08 | IL7R/CCR7/CXCR6/CD74/HLA-DQB1/HLA-DQA1/HLA-DPA1/KLRD1/HLA-DRA/HLA-DRB1 | 10 | MF |
| GO:0070665 | positive regulation of leukocyte proliferation | 10/88 | 150/18723 | 2.13064E-09 | 7.72593E-08 | 5.48092E-08 | GPR183/CD28/PTPN22/TNFSF4/CD74/VCAM1/HLA-DPA1/HLA-DPB1/HAVCR2/CCL5 | 10 | BP |

| | | | | | | | | | |
|---|---|---|---|---|---|---|---|---|---|
| GO:0002456 | T cell mediated immunity | 9/88 | 109/18723 | 2.14037E-09 | 7.72593E-08 | 5.48092E-08 | IL7R/TNFSF4/RAB27A/CTSC/KLRD1/PRF1/HLA-DRA/HLA-DRB1/CD8A | 9 | BP |
| GO:0030662 | coated vesicle membrane | 10/89 | 181/19550 | 9.74741E-09 | 7.79793E-08 | 5.6827E-08 | IL7R/LDLRAP1/CD74/HLA-DQB1/HLA-DQA1/HLA-DPA1/HLA-DPB1/HLA-DRB5/HLA-DRA/HLA-DRB1 | 10 | CC |
| GO:0030135 | coated vesicle | 12/89 | 299/19550 | 1.09198E-08 | 8.11188E-08 | 5.91149E-08 | IL7R/LDLRAP1/RAB27A/CTSC/CD74/HLA-DQB1/HLA-DQA1/HLA-DPA1/HLA-DPB1/HLA-DRB5/HLA-DRA/HLA-DRB1 | 12 | CC |
| GO:0032609 | interferon-gamma production | 9/88 | 112/18723 | 2.7271E-09 | 9.47919E-08 | 6.72472E-08 | CCR7/PTPN22/TNFSF4/CD244/CRTAM/HLA-DPA1/HLA-DPB1/HAVCR2/HLA-DRB1 | 9 | BP |
| GO:0032649 | regulation of interferon-gamma production | 9/88 | 112/18723 | 2.7271E-09 | 9.47919E-08 | 6.72472E-08 | CCR7/PTPN22/TNFSF4/CD244/CRTAM/HLA-DPA1/HLA-DPB1/HAVCR2/HLA-DRB1 | 9 | BP |
| GO:0048020 | CCR chemokine receptor binding | 7/89 | 48/18368 | 3.09088E-09 | 9.54431E-08 | 7.12987E-08 | CCL3L1/CCL3L3/CXCL13/CCL3/CCL4L1/CCL4/CCL5 | 7 | MF |
| GO:0008009 | chemokine activity | 7/89 | 49/18368 | 3.59194E-09 | 9.54431E-08 | 7.12987E-08 | CCL3L1/CCL3L3/CXCL13/CCL3/CCL4L1/CCL4/CCL5 | 7 | MF |
| GO:0001913 | T cell mediated cytotoxicity | 7/88 | 49/18723 | 2.90911E-09 | 9.928E-08 | 7.04311E-08 | IL7R/RAB27A/CTSC/KLRD1/PRF1/HLA-DRA/HLA-DRB1 | 7 | BP |
| GO:0045589 | regulation of regulatory T cell differentiation | 6/88 | 28/18723 | 3.14398E-09 | 1.05379E-07 | 7.47581E-08 | CD28/TNFSF4/LAG3/IFNG/HLA-DRA/HLA-DRB1 | 6 | BP |
| GO:0016064 | immunoglobulin mediated immune response | 11/88 | 207/18723 | 3.51786E-09 | 1.15843E-07 | 8.21809E-08 | CD28/TNFSF4/CD74/HLA-DMA/HLA-DQB1/HLA-DQA1/HLA-DPA1/HLA-DPB1/HLA-DRB5/HLA-DRA/HLA-DRB1 | 11 | BP |
| GO:0032588 | trans-Golgi network membrane | 8/89 | 100/19550 | 1.77082E-08 | 1.17031E-07 | 8.52858E-08 | CD74/HLA-DQB1/HLA-DQA1/HLA-DPA1/HLA-DPB1/HLA-DRB5/HLA-DRA/HLA-DRB1 | 8 | CC |
| GO:0030666 | endocytic vesicle membrane | 10/89 | 193/19550 | 1.80048E-08 | 1.17031E-07 | 8.52858E-08 | IL7R/LDLRAP1/CD74/HLA-DQB1/HLA-DQA1/HLA-DPA1/HLA-DPB1/HLA-DRB5/HLA-DRA/HLA-DRB1 | 10 | CC |
| GO:0019724 | B cell mediated immunity | 11/88 | 210/18723 | 4.09052E-09 | 1.32378E-07 | 9.39112E-08 | CD28/TNFSF4/CD74/HLA-DMA/HLA-DQB1/HLA-DQA1/HLA-DPA1/HLA-DPB1/HLA-DRB5/HLA-DRA/HLA-DRB1 | 11 | BP |
| GO:0071346 | cellular response to interferon-gamma | 9/88 | 118/18723 | 4.33728E-09 | 1.37984E-07 | 9.78886E-08 | CCL3L1/CCL3L3/FASLG/IFNG/CCL3/HLA-DPA1/CCL4L1/CCL4/CCL5 | 9 | BP |
| GO:0001910 | regulation of leukocyte mediated cytotoxicity | 8/88 | 82/18723 | 4.62204E-09 | 1.44593E-07 | 1.02577E-07 | IL7R/SH2D1A/CRTAM/LAG3/HAVCR2/KLRD1/HLA-DRA/HLA-DRB1 | 8 | BP |
| GO:0045066 | regulatory T cell differentiation | 6/88 | 31/18723 | 6.07552E-09 | 1.86947E-07 | 1.32624E-07 | CD28/TNFSF4/LAG3/IFNG/HLA-DRA/HLA-DRB1 | 6 | BP |
| GO:0002706 | regulation of lymphocyte mediated immunity | 10/88 | 168/18723 | 6.38977E-09 | 1.93445E-07 | 1.37234E-07 | IL7R/CD28/TNFSF4/SH2D1A/CRTAM/LAG3/HAVCR2/KLRD1/HLA-DRA/HLA-DRB1 | 10 | BP |
| GO:0031349 | positive regulation of defense response | 12/88 | 278/18723 | 6.8249E-09 | 2.03339E-07 | 1.44252E-07 | CCR7/CD28/TNFSF4/CTSC/SH2D1A/CRTAM/LAG3/IFNG/CCL3/HAVCR2/KLRD1/CCL5 | 12 | BP |
| GO:0042129 | regulation of T cell proliferation | 10/88 | 171/18723 | 7.57732E-09 | 2.22229E-07 | 1.57653E-07 | CD28/PTPN22/TNFSF4/CRTAM/VCAM1/HLA-DPA1/HLA-DPB1/HAVCR2/HLA-DRB1/CCL5 | 10 | BP |
| GO:0003823 | antigen binding | 10/89 | 171/18368 | 1.01346E-08 | 2.3563E-07 | 1.76022E-07 | IL7R/LAG3/HLA-DQB1/HLA-DQA1/HLA-DPA1/HLA-DPB1/KLRD1/HLA-DRB5/HLA-DRA/HLA-DRB1 | 10 | MF |
| GO:0005125 | cytokine activity | 11/89 | 235/18368 | 1.80773E-08 | 3.73598E-07 | 2.79088E-07 | LTB/TNFSF4/CCL3L1/CCL3L3/FASLG/IFNG/CXCL13/CCL3/CCL4L1/CCL4/CCL5 | 11 | MF |
| GO:0042287 | MHC protein binding | 6/89 | 40/18368 | 3.66801E-08 | 6.82251E-07 | 5.09661E-07 | FCRL6/CD244/CD74/LAG3/KLRD1/CD8A | 6 | MF |
| GO:0042102 | positive regulation of T cell proliferation | 8/88 | 101/18723 | 2.43795E-08 | 7.04004E-07 | 4.99434E-07 | CD28/PTPN22/TNFSF4/VCAM1/HLA-DPA1/HLA-DPB1/HAVCR2/CCL5 | 8 | BP |
| GO:0071674 | mononuclear cell migration | 10/88 | 196/18723 | 2.79183E-08 | 7.93979E-07 | 5.63264E-07 | GPR183/CCR7/CCL3L1/CCL3L3/CRTAM/CXCL13/CCL3/CCL4L1/CCL4/CCL5 | 10 | BP |
| GO:0046634 | regulation of alpha-beta T cell activation | 8/88 | 104/18723 | 3.07194E-08 | 8.60602E-07 | 6.10527E-07 | CD28/PTPN22/TNFSF4/ZNF683/CRTAM/IFNG/HLA-DRA/HLA-DRB1 | 8 | BP |
| GO:0042098 | T cell proliferation | 10/88 | 199/18723 | 3.22442E-08 | 8.90035E-07 | 6.31407E-07 | CD28/PTPN22/TNFSF4/CRTAM/VCAM1/HLA-DPA1/HLA-DPB1/HAVCR2/HLA-DRB1/CCL5 | 10 | BP |
| GO:0002287 | alpha-beta T cell activation involved in immune response | 7/88 | 69/18723 | 3.38699E-08 | 9.08196E-07 | 6.44291E-07 | GPR183/LEF1/TNFSF4/EOMES/IFNG/HLA-DRA/HLA-DRB1 | 7 | BP |
| GO:0002293 | alpha-beta T cell differentiation involved in immune response | 7/88 | 69/18723 | 3.38699E-08 | 9.08196E-07 | 6.44291E-07 | GPR183/LEF1/TNFSF4/EOMES/IFNG/HLA-DRA/HLA-DRB1 | 7 | BP |
| GO:0042379 | chemokine receptor binding | 7/89 | 72/18368 | 5.62977E-08 | 9.51942E-07 | 7.11128E-07 | CCL3L1/CCL3L3/CXCL13/CCL3/CCL4L1/CCL4/CCL5 | 7 | MF |
| GO:0005126 | cytokine receptor binding | 11/89 | 271/18368 | 7.77352E-08 | 1.2049E-06 | 9.00092E-07 | LTB/TNFSF4/CCL3L1/CCL3L3/FASLG/IFNG/CXCL13/CCL3/CCL4L1/CCL4/CCL5 | 11 | MF |
| GO:0032729 | positive regulation of interferon-gamma production | 7/88 | 72/18723 | 4.57225E-08 | 1.20875E-06 | 8.57508E-07 | PTPN22/TNFSF4/CD244/CRTAM/HLA-DPA1/HLA-DPB1/HAVCR2 | 7 | BP |
| GO:0046632 | alpha-beta T cell differentiation | 8/88 | 112/18723 | 5.5008E-08 | 1.43403E-06 | 1.01733E-06 | GPR183/LEF1/TNFSF4/ZNF683/EOMES/IFNG/HLA-DRA/HLA-DRB1 | 8 | BP |
| GO:0002708 | positive regulation of lymphocyte mediated immunity | 8/88 | 113/18723 | 5.89742E-08 | 1.51636E-06 | 1.07574E-06 | CD28/TNFSF4/SH2D1A/CRTAM/LAG3/KLRD1/HLA-DRA/HLA-DRB1 | 8 | BP |
| GO:0002292 | T cell differentiation involved in immune response | 7/88 | 75/18723 | 6.09087E-08 | 1.54494E-06 | 1.09601E-06 | GPR183/LEF1/TNFSF4/EOMES/IFNG/HLA-DRA/HLA-DRB1 | 7 | BP |
| GO:0002377 | immunoglobulin production | 10/88 | 216/18723 | 6.98746E-08 | 1.74873E-06 | 1.24058E-06 | CD28/TNFSF4/HLA-DMA/HLA-DQB1/HLA-DQA1/HLA-DPA1/HLA-DPB1/HLA-DRB5/HLA-DRA/HLA-DRB1 | 10 | BP |
| GO:0097529 | myeloid leukocyte migration | 10/88 | 220/18723 | 8.30017E-08 | 2.04992E-06 | 1.45425E-06 | CCR7/CCL3L1/CCL3L3/CD74/ITGA1/CXCL13/CCL3/CCL4L1/CCL4/CCL5 | 10 | BP |
| GO:0002703 | regulation of leukocyte mediated immunity | 10/88 | 226/18723 | 1.0677E-07 | 2.60268E-06 | 1.84639E-06 | IL7R/CD28/TNFSF4/SH2D1A/CRTAM/LAG3/HAVCR2/KLRD1/HLA-DRA/HLA-DRB1 | 10 | BP |
| GO:0050852 | T cell receptor signaling pathway | 8/88 | 123/18723 | 1.14257E-07 | 2.7495E-06 | 1.95055E-06 | CCR7/CD28/PTPN22/THEMIS/SH2D1A/HLA-DQB1/HLA-DPB1/HLA-DRB1 | 8 | BP |
| GO:0030176 | integral component of endoplasmic reticulum membrane | 8/89 | 162/19550 | 7.47216E-07 | 4.5712E-06 | 3.33124E-06 | CD74/HLA-DQB1/HLA-DQA1/HLA-DPA1/HLA-DPB1/HLA-DRB5/HLA-DRA/HLA-DRB1 | 8 | CC |
| GO:0043383 | negative T cell selection | 4/88 | 12/18723 | 2.19115E-07 | 5.2014E-06 | 3.68997E-06 | CCR7/CD28/THEMIS/CD74 | 4 | BP |

| | | | | | | | | | |
|---|---|---|---|---|---|---|---|---|---|
| GO:0002705 | positive regulation of leukocyte mediated immunity | 8/88 | 134/18723 | 2.2169E-07 | 5.2014E-06 | 3.68997E-06 | CD28/TNFSF4/SH2D1A/CRTAM/LAG3/KLRD1/HLA-DRA/HLA-DRB1 | 8 | BP |
| GO:0045582 | positive regulation of T cell differentiation | 7/88 | 91/18723 | 2.33768E-07 | 5.41003E-06 | 3.83798E-06 | IL7R/LEF1/TNFSF4/CD74/IFNG/HLA-DRA/HLA-DRB1 | 7 | BP |
| GO:0002695 | negative regulation of leukocyte activation | 9/88 | 187/18723 | 2.36347E-07 | 5.41003E-06 | 3.83798E-06 | PTPN22/TNFSF4/ID2/CD74/CRTAM/LAG3/HAVCR2/CST7/HLA-DRB1 | 9 | BP |
| GO:0001912 | positive regulation of leukocyte mediated cytotoxicity | 6/88 | 56/18723 | 2.43895E-07 | 5.51556E-06 | 3.91284E-06 | SH2D1A/CRTAM/LAG3/KLRD1/HLA-DRA/HLA-DRB1 | 6 | BP |
| GO:0031227 | intrinsic component of endoplasmic reticulum membrane | 8/89 | 170/19550 | 1.07582E-06 | 6.21588E-06 | 4.52979E-06 | CD74/HLA-DQB1/HLA-DQA1/HLA-DPA1/HLA-DPB1/HLA-DRB5/HLA-DRA/HLA-DRB1 | 8 | CC |
| GO:0043547 | positive regulation of GTPase activity | 10/88 | 255/18723 | 3.27258E-07 | 7.31266E-06 | 5.18774E-06 | CCR7/ARAP2/CCL3L1/CCL3L3/F2R/CXCL13/CCL3/CCL4L1/CCL4/CCL5 | 10 | BP |
| GO:0032633 | interleukin-4 production | 5/88 | 33/18723 | 4.37458E-07 | 9.54778E-06 | 6.77337E-06 | LEF1/CD28/TNFSF4/HAVCR2/HLA-DRB1 | 5 | BP |
| GO:0032673 | regulation of interleukin-4 production | 5/88 | 33/18723 | 4.37458E-07 | 9.54778E-06 | 6.77337E-06 | LEF1/CD28/TNFSF4/HAVCR2/HLA-DRB1 | 5 | BP |
| GO:0002573 | myeloid leukocyte differentiation | 9/88 | 208/18723 | 5.8006E-07 | 1.24576E-05 | 8.83767E-06 | GPR183/CCR7/LEF1/RBPJ/ID2/CD74/IFNG/CCL3/HLA-DRB1 | 9 | BP |
| GO:0045621 | positive regulation of lymphocyte differentiation | 7/88 | 104/18723 | 5.84055E-07 | 1.24576E-05 | 8.83767E-06 | IL7R/LEF1/TNFSF4/CD74/IFNG/HLA-DRA/HLA-DRB1 | 7 | BP |
| GO:0002468 | dendritic cell antigen processing and presentation | 4/88 | 15/18723 | 5.97747E-07 | 1.26064E-05 | 8.94323E-06 | CCR7/CD74/HLA-DRA/HLA-DRB1 | 4 | BP |
| GO:0050866 | negative regulation of cell activation | 9/88 | 210/18723 | 6.28495E-07 | 1.31076E-05 | 9.29878E-06 | PTPN22/TNFSF4/ID2/CD74/CRTAM/LAG3/HAVCR2/CST7/HLA-DRB1 | 9 | BP |
| GO:0046635 | positive regulation of alpha-beta T cell activation | 6/88 | 67/18723 | 7.19306E-07 | 1.48367E-05 | 1.05254E-05 | CD28/PTPN22/TNFSF4/IFNG/HLA-DRA/HLA-DRB1 | 6 | BP |
| GO:0051250 | negative regulation of lymphocyte activation | 8/88 | 157/18723 | 7.44496E-07 | 1.48662E-05 | 1.05463E-05 | PTPN22/TNFSF4/ID2/CD74/CRTAM/LAG3/HAVCR2/HLA-DRB1 | 8 | BP |
| GO:1902107 | positive regulation of leukocyte differentiation | 8/88 | 157/18723 | 7.44496E-07 | 1.48662E-05 | 1.05463E-05 | IL7R/LEF1/TNFSF4/ID2/CD74/IFNG/HLA-DRA/HLA-DRB1 | 8 | BP |
| GO:1903708 | positive regulation of hemopoiesis | 8/88 | 157/18723 | 7.44496E-07 | 1.48662E-05 | 1.05463E-05 | IL7R/LEF1/TNFSF4/ID2/CD74/IFNG/HLA-DRA/HLA-DRB1 | 8 | BP |
| GO:0002294 | CD4-positive, alpha-beta T cell differentiation involved in immune response | 6/88 | 68/18723 | 7.8596E-07 | 1.55289E-05 | 1.10165E-05 | GPR183/LEF1/TNFSF4/IFNG/HLA-DRA/HLA-DRB1 | 6 | BP |
| GO:0002548 | monocyte chemotaxis | 6/88 | 70/18723 | 9.34472E-07 | 1.82709E-05 | 1.29617E-05 | CCL3L1/CCL3L3/CCL3/CCL4L1/CCL4/CCL5 | 6 | BP |
| GO:0002347 | response to tumor cell | 5/88 | 39/18723 | 1.03797E-06 | 2.00852E-05 | 1.42488E-05 | ABI3/CRTAM/HAVCR2/PRF1/HLA-DRB1 | 5 | BP |
| GO:0002833 | positive regulation of response to biotic stimulus | 8/88 | 168/18723 | 1.24221E-06 | 2.3792E-05 | 1.68785E-05 | SH2D1A/CRTAM/LAG3/OASL/HAVCR2/KLRD1/HLA-DRB1/CCL5 | 8 | BP |
| GO:0005770 | late endosome | 9/89 | 282/19550 | 5.51901E-06 | 2.8981E-05 | 2.11198E-05 | RAB27A/CD74/F2R/HLA-DMA/CD63/CST7/HLA-DRB5/HLA-DRA/HLA-DRB1 | 9 | CC |
| GO:0030658 | transport vesicle membrane | 8/89 | 212/19550 | 5.57327E-06 | 2.8981E-05 | 2.11198E-05 | CD74/HLA-DQB1/HLA-DQA1/HLA-DPA1/HLA-DPB1/HLA-DRB5/HLA-DRA/HLA-DRB1 | 8 | CC |
| GO:0048245 | eosinophil chemotaxis | 4/88 | 19/18723 | 1.67313E-06 | 3.14046E-05 | 2.2279E-05 | CCL3/CCL4L1/CCL4/CCL5 | 4 | BP |
| GO:0140131 | positive regulation of lymphocyte chemotaxis | 4/88 | 19/18723 | 1.67313E-06 | 3.14046E-05 | 2.2279E-05 | CXCL13/CCL3/CCL4/CCL5 | 4 | BP |
| GO:0050868 | negative regulation of T cell activation | 7/88 | 122/18723 | 1.72024E-06 | 3.19692E-05 | 2.26795E-05 | PTPN22/TNFSF4/CD74/CRTAM/LAG3/HAVCR2/HLA-DRB1 | 7 | BP |
| GO:0042269 | regulation of natural killer cell mediated cytotoxicity | 5/88 | 44/18723 | 1.92201E-06 | 3.53688E-05 | 2.50913E-05 | SH2D1A/CRTAM/LAG3/HAVCR2/KLRD1 | 5 | BP |
| GO:0043367 | CD4-positive, alpha-beta T cell differentiation | 6/88 | 83/18723 | 2.56199E-06 | 4.66879E-05 | 3.31212E-05 | GPR183/LEF1/TNFSF4/IFNG/HLA-DRA/HLA-DRB1 | 6 | BP |
| GO:0002715 | regulation of natural killer cell mediated immunity | 5/88 | 48/18723 | 2.986E-06 | 5.38915E-05 | 3.82317E-05 | SH2D1A/CRTAM/LAG3/HAVCR2/KLRD1 | 5 | BP |
| GO:0030101 | natural killer cell activation | 6/88 | 88/18723 | 3.61009E-06 | 6.45346E-05 | 4.57821E-05 | PTPN22/RAB27A/CD244/ZNF683/SLAMF7/HAVCR2 | 6 | BP |
| GO:0072677 | eosinophil migration | 4/88 | 23/18723 | 3.76789E-06 | 6.672E-05 | 4.73324E-05 | CCL3/CCL4L1/CCL4/CCL5 | 4 | BP |
| GO:1903038 | negative regulation of leukocyte cell-cell adhesion | 7/88 | 141/18723 | 4.51627E-06 | 7.92246E-05 | 5.62034E-05 | PTPN22/TNFSF4/CD74/CRTAM/LAG3/HAVCR2/HLA-DRB1 | 7 | BP |
| GO:0002690 | positive regulation of leukocyte chemotaxis | 6/88 | 94/18723 | 5.30321E-06 | 9.20281E-05 | 6.52864E-05 | CCR7/CD74/CXCL13/CCL3/CCL4/CCL5 | 6 | BP |
| GO:1901623 | regulation of lymphocyte chemotaxis | 4/88 | 25/18723 | 5.3442E-06 | 9.20281E-05 | 6.52864E-05 | CXCL13/CCL3/CCL4/CCL5 | 4 | BP |
| GO:0006968 | cellular defense response | 5/88 | 54/18723 | 5.39413E-06 | 9.20435E-05 | 6.52974E-05 | FCMR/KLRG1/SH2D1A/GNLY/PRF1 | 5 | BP |
| GO:1903900 | regulation of viral life cycle | 7/88 | 148/18723 | 6.21905E-06 | 0.000104535 | 7.41589E-05 | CD28/CD74/OASL/APOBEC3C/APOBEC3G/HLA-DRB1/CCL5 | 7 | BP |
| GO:0002418 | immune response to tumor cell | 4/88 | 26/18723 | 6.29325E-06 | 0.000104535 | 7.41589E-05 | CRTAM/HAVCR2/PRF1/HLA-DRB1 | 4 | BP |
| GO:0045954 | positive regulation of natural killer cell mediated cytotoxicity | 4/88 | 26/18723 | 6.29325E-06 | 0.000104535 | 7.41589E-05 | SH2D1A/CRTAM/LAG3/KLRD1 | 4 | BP |
| GO:0005802 | trans-Golgi network | 8/89 | 259/19550 | 2.38337E-05 | 0.000118033 | 8.60163E-05 | CD74/HLA-DQB1/HLA-DQA1/HLA-DPA1/HLA-DPB1/HLA-DRB5/HLA-DRA/HLA-DRB1 | 8 | CC |
| GO:0002763 | positive regulation of myeloid leukocyte differentiation | 5/88 | 58/18723 | 7.70096E-06 | 0.000126796 | 8.99512E-05 | LEF1/ID2/CD74/IFNG/HLA-DRB1 | 5 | BP |
| GO:0035710 | CD4-positive, alpha-beta T cell activation | 6/88 | 102/18723 | 8.51121E-06 | 0.000138467 | 9.82314E-05 | GPR183/LEF1/TNFSF4/IFNG/HLA-DRA/HLA-DRB1 | 6 | BP |
| GO:0010818 | T cell chemotaxis | 4/88 | 28/18723 | 8.55739E-06 | 0.000138467 | 9.82314E-05 | GPR183/CXCL13/CCL3/CCL5 | 4 | BP |
| GO:0002429 | immune response-activating cell surface receptor signaling pathway | 9/88 | 291/18723 | 9.12996E-06 | 0.000145228 | 0.000103028 | CCR7/CD28/PTPN22/THEMIS/SH2D1A/HLA-DQB1/HLA-DPB1/KLRD1/HLA-DRB1 | 9 | BP |
| GO:0002757 | immune response-activating signal transduction | 9/88 | 291/18723 | 9.12996E-06 | 0.000145228 | 0.000103028 | CCR7/CD28/PTPN22/THEMIS/SH2D1A/HLA-DQB1/HLA-DPB1/KLRD1/HLA-DRB1 | 9 | BP |
| GO:0043030 | regulation of macrophage activation | 5/88 | 61/18723 | 9.88841E-06 | 0.000155971 | 0.000110649 | CTSC/CD74/CCL3/HAVCR2/CST7 | 5 | BP |
| GO:0002274 | myeloid leukocyte activation | 8/88 | 223/18723 | 1.01301E-05 | 0.000158451 | 0.000112408 | RBPJ/CTSC/CD74/IFNG/CCL3/HAVCR2/CST7/CCL5 | 8 | BP |
| GO:0035821 | modulation of process of other organism | 6/88 | 106/18723 | 1.06221E-05 | 0.000162095 | 0.000114993 | LEF1/IFNG/CCL3/PRF1/CCL4/CCL5 | 6 | BP |
| GO:0042116 | macrophage activation | 6/88 | 106/18723 | 1.06221E-05 | 0.000162095 | 0.000114993 | CTSC/CD74/IFNG/CCL3/HAVCR2/CST7 | 6 | BP |
| GO:0071887 | leukocyte apoptotic process | 6/88 | 106/18723 | 1.06221E-05 | 0.000162095 | 0.000114993 | IL7R/CCR7/PDCD1/CD74/FASLG/CCL5 | 6 | BP |
| GO:0035747 | natural killer cell chemotaxis | 3/88 | 10/18723 | 1.1756E-05 | 0.000177952 | 0.000126242 | CCL3/CCL4/CCL5 | 3 | BP |
| GO:0050792 | regulation of viral process | 7/88 | 164/18723 | 1.21811E-05 | 0.000182911 | 0.00012976 | CD28/CD74/OASL/APOBEC3C/APOBEC3G/HLA-DRB1/CCL5 | 7 | BP |
| GO:0002717 | positive regulation of natural killer cell mediated immunity | 4/88 | 31/18723 | 1.30098E-05 | 0.000193805 | 0.000137489 | SH2D1A/CRTAM/LAG3/KLRD1 | 4 | BP |
| GO:0071677 | positive regulation of mononuclear cell migration | 5/88 | 65/18723 | 1.35281E-05 | 0.00019994 | 0.000141841 | CCR7/CXCL13/CCL3/CCL4/CCL5 | 5 | BP |
| GO:0002822 | regulation of adaptive immune response based on somatic recombination of immune receptors built from immunoglobulin superfamily domains | 7/88 | 168/18723 | 1.42472E-05 | 0.000208922 | 0.000148213 | IL7R/CD28/TNFSF4/HAVCR2/KLRD1/HLA-DRA/HLA-DRB1 | 7 | BP |
| GO:0042093 | T-helper cell differentiation | 5/88 | 66/18723 | 1.45831E-05 | 0.00021219 | 0.000150532 | GPR183/LEF1/TNFSF4/HLA-DRA/HLA-DRB1 | 5 | BP |
| GO:0019835 | cytolysis | 4/88 | 32/18723 | 1.48152E-05 | 0.000212275 | 0.000150592 | PRF1/GZMA/GZMB/GZMH | 4 | BP |

| | | | | | | | | | |
|---|---|---|---|---|---|---|---|---|---|
| GO:0043372 | positive regulation of CD4-positive, alpha-beta T cell differentiation | 4/88 | 32/18723 | 1.48152E-05 | 0.000212275 | 0.000150592 | TNFSF4/IFNG/HLA-DRA/HLA-DRB1 | 4 | BP |
| GO:0071347 | cellular response to interleukin-1 | 6/88 | 113/18723 | 1.53247E-05 | 0.000217912 | 0.000154591 | CCL3L1/CCL3L3/CCL3/CCL4L1/CCL4/CCL5 | 6 | BP |
| GO:0045060 | negative thymic T cell selection | 3/88 | 11/18723 | 1.61095E-05 | 0.000227351 | 0.000161287 | CCR7/CD28/CD74 | 3 | BP |
| GO:0046637 | regulation of alpha-beta T cell differentiation | 5/88 | 68/18723 | 1.68849E-05 | 0.000236515 | 0.000167788 | TNFSF4/ZNF683/IFNG/HLA-DRA/HLA-DRB1 | 5 | BP |
| GO:0050851 | antigen receptor-mediated signaling pathway | 8/88 | 240/18723 | 1.72477E-05 | 0.000239807 | 0.000170123 | CCR7/CD28/PTPN22/THEMIS/SH2D1A/HLA-DQB1/HLA-DPB1/HLA-DRB1 | 8 | BP |
| GO:0031902 | late endosome membrane | 6/89 | 146/19550 | 5.4631E-05 | 0.000258256 | 0.000188203 | RAB27A/HLA-DMA/CD63/HLA-DRB5/HLA-DRA/HLA-DRB1 | 6 | CC |
| GO:2000403 | positive regulation of lymphocyte migration | 4/88 | 35/18723 | 2.1341E-05 | 0.000293283 | 0.00020806 | CXCL13/CCL3/CCL4/CCL5 | 4 | BP |
| GO:0043380 | regulation of memory T cell differentiation | 3/88 | 12/18723 | 2.14064E-05 | 0.000293283 | 0.00020806 | TNFSF4/HLA-DRA/HLA-DRB1 | 3 | BP |
| GO:0002761 | regulation of myeloid leukocyte differentiation | 6/88 | 120/18723 | 2.15818E-05 | 0.000293544 | 0.000208245 | LEF1/ID2/CD74/IFNG/CCL3/HLA-DRB1 | 6 | BP |
| GO:0070227 | lymphocyte apoptotic process | 5/88 | 72/18723 | 2.23316E-05 | 0.000301557 | 0.00021393 | IL7R/PDCD1/CD74/FASLG/CCL5 | 5 | BP |
| GO:0002688 | regulation of leukocyte chemotaxis | 6/88 | 122/18723 | 2.37041E-05 | 0.000317804 | 0.000225456 | CCR7/CD74/CXCL13/CCL3/CCL4/CCL5 | 6 | BP |
| GO:0002819 | regulation of adaptive immune response | 7/88 | 183/18723 | 2.47538E-05 | 0.000329524 | 0.00023377 | IL7R/CD28/TNFSF4/HAVCR2/KLRD1/HLA-DRA/HLA-DRB1 | 7 | BP |
| GO:0043379 | memory T cell differentiation | 3/88 | 13/18723 | 2.77337E-05 | 0.000366592 | 0.000260067 | TNFSF4/HLA-DRA/HLA-DRB1 | 3 | BP |
| GO:0001914 | regulation of T cell mediated cytotoxicity | 4/88 | 39/18723 | 3.3047E-05 | 0.000430758 | 0.000305588 | IL7R/KLRD1/HLA-DRA/HLA-DRB1 | 4 | BP |
| GO:2000516 | positive regulation of CD4-positive, alpha-beta T cell activation | 4/88 | 39/18723 | 3.3047E-05 | 0.000430758 | 0.000305588 | TNFSF4/IFNG/HLA-DRA/HLA-DRB1 | 4 | BP |
| GO:0051607 | defense response to virus | 8/88 | 265/18723 | 3.50426E-05 | 0.00044047 | 0.000312477 | SERINC5/PTPN22/LYST/OASL/APOBEC3C/IFNG/APOBEC3G/PRF1 | 8 | BP |
| GO:0140546 | defense response to symbiont | 8/88 | 265/18723 | 3.50426E-05 | 0.00044047 | 0.000312477 | SERINC5/PTPN22/LYST/OASL/APOBEC3C/IFNG/APOBEC3G/PRF1 | 8 | BP |
| GO:0001771 | immunological synapse formation | 3/88 | 14/18723 | 3.51774E-05 | 0.00044047 | 0.000312477 | CCR7/HAVCR2/PRF1 | 3 | BP |
| GO:0043922 | negative regulation by host of viral transcription | 3/88 | 14/18723 | 3.51774E-05 | 0.00044047 | 0.000312477 | CCL3/CCL4/CCL5 | 3 | BP |
| GO:0090715 | immunological memory formation process | 3/88 | 14/18723 | 3.51774E-05 | 0.00044047 | 0.000312477 | TNFSF4/HLA-DRA/HLA-DRB1 | 3 | BP |
| GO:0019079 | viral genome replication | 6/88 | 131/18723 | 3.54347E-05 | 0.00044047 | 0.000312477 | CD28/CXCR6/OASL/APOBEC3C/APOBEC3G/CCL5 | 6 | BP |
| GO:0045089 | positive regulation of innate immune response | 6/88 | 131/18723 | 3.54347E-05 | 0.00044047 | 0.000312477 | SH2D1A/CRTAM/LAG3/HAVCR2/KLRD1/CCL5 | 6 | BP |
| GO:0022408 | negative regulation of cell-cell adhesion | 7/88 | 196/18723 | 3.8398E-05 | 0.000474164 | 0.000336381 | PTPN22/TNFSF4/CD74/CRTAM/LAG3/HAVCR2/HLA-DRB1 | 7 | BP |
| GO:2000106 | regulation of leukocyte apoptotic process | 5/88 | 81/18723 | 3.956E-05 | 0.000485321 | 0.000344296 | IL7R/CCR7/PDCD1/CD74/CCL5 | 5 | BP |
| GO:0032735 | positive regulation of interleukin-12 production | 4/88 | 41/18723 | 4.0398E-05 | 0.000492383 | 0.000349306 | LTB/CCR7/TNFSF4/IFNG | 4 | BP |
| GO:0002687 | positive regulation of leukocyte migration | 6/88 | 135/18723 | 4.19578E-05 | 0.000508096 | 0.000360453 | CCR7/CD74/CXCL13/CCL3/CCL4/CCL5 | 6 | BP |
| GO:0042088 | T-helper 1 type immune response | 4/88 | 43/18723 | 4.88785E-05 | 0.000588109 | 0.000417215 | LEF1/TNFSF4/HAVCR2/HLA-DRB1 | 4 | BP |
| GO:0002709 | regulation of T cell mediated immunity | 5/88 | 85/18723 | 4.99051E-05 | 0.00059286 | 0.000420586 | IL7R/TNFSF4/KLRD1/HLA-DRA/HLA-DRB1 | 5 | BP |
| GO:0045069 | regulation of viral genome replication | 5/88 | 85/18723 | 4.99051E-05 | 0.00059286 | 0.000420586 | CD28/OASL/APOBEC3C/APOBEC3G/CCL5 | 5 | BP |
| GO:0050921 | positive regulation of chemotaxis | 6/88 | 141/18723 | 5.35154E-05 | 0.000628492 | 0.000445864 | CCR7/CD74/CXCL13/CCL3/CCL4/CCL5 | 6 | BP |
| GO:0150076 | neuroinflammatory response | 4/88 | 44/18723 | 5.35741E-05 | 0.000628492 | 0.000445864 | CTSC/IFNG/CCL3/CST7 | 4 | BP |
| GO:0050729 | positive regulation of inflammatory response | 6/88 | 142/18723 | 5.56676E-05 | 0.000648994 | 0.000460408 | CCR7/CD28/TNFSF4/CTSC/IFNG/CCL3 | 6 | BP |
| GO:0071222 | cellular response to lipopolysaccharide | 7/88 | 209/18723 | 5.76969E-05 | 0.000666605 | 0.000472902 | PTPN22/TNFSF4/TNIP3/CXCL13/CCL3/HAVCR2/CCL5 | 7 | BP |
| GO:0070555 | response to interleukin-1 | 6/88 | 143/18723 | 5.78885E-05 | 0.000666605 | 0.000472902 | CCL3L1/CCL3L3/CCL3/CCL4L1/CCL4/CCL5 | 6 | BP |
| GO:2000107 | negative regulation of leukocyte apoptotic process | 4/88 | 46/18723 | 6.39414E-05 | 0.000731818 | 0.000519165 | IL7R/CCR7/CD74/CCL5 | 4 | BP |
| GO:0045591 | positive regulation of regulatory T cell differentiation | 3/88 | 17/18723 | 6.50484E-05 | 0.000735517 | 0.00052179 | IFNG/HLA-DRA/HLA-DRB1 | 3 | BP |
| GO:0090713 | immunological memory process | 3/88 | 17/18723 | 6.50484E-05 | 0.000735517 | 0.00052179 | TNFSF4/HLA-DRA/HLA-DRB1 | 3 | BP |
| GO:0001774 | microglial cell activation | 4/88 | 47/18723 | 6.96397E-05 | 0.000782717 | 0.000555274 | CTSC/IFNG/CCL3/CST7 | 4 | BP |
| GO:0045088 | regulation of innate immune response | 7/88 | 218/18723 | 7.52297E-05 | 0.000840513 | 0.000596276 | PTPN22/SH2D1A/CRTAM/LAG3/HAVCR2/KLRD1/CCL5 | 7 | BP |
| GO:0002544 | chronic inflammatory response | 3/88 | 18/18723 | 7.7793E-05 | 0.000864008 | 0.000612944 | VCAM1/CXCL13/CCL5 | 3 | BP |
| GO:0071219 | cellular response to molecule of bacterial origin | 7/88 | 221/18723 | 8.19588E-05 | 0.000904922 | 0.000641968 | PTPN22/TNFSF4/TNIP3/CXCL13/CCL3/HAVCR2/CCL5 | 7 | BP |
| GO:0005771 | multivesicular body | 4/89 | 64/19550 | 0.000206985 | 0.000935932 | 0.000682056 | RAB27A/CD74/CD63/CST7 | 4 | CC |
| GO:0050920 | regulation of chemotaxis | 7/88 | 223/18723 | 8.67119E-05 | 0.000951803 | 0.000675227 | GPR183/CCR7/CD74/CXCL13/CCL3/CCL4/CCL5 | 7 | BP |
| GO:0019083 | viral transcription | 4/88 | 50/18723 | 8.89566E-05 | 0.000954123 | 0.000676873 | LEF1/CCL3/CCL4/CCL5 | 4 | BP |
| GO:0045058 | T cell selection | 4/88 | 50/18723 | 8.89566E-05 | 0.000954123 | 0.000676873 | CCR7/CD28/THEMIS/CD74 | 4 | BP |
| GO:0046638 | positive regulation of alpha-beta T cell differentiation | 4/88 | 50/18723 | 8.89566E-05 | 0.000954123 | 0.000676873 | TNFSF4/IFNG/HLA-DRA/HLA-DRB1 | 4 | BP |
| GO:0070231 | T cell apoptotic process | 4/88 | 50/18723 | 8.89566E-05 | 0.000954123 | 0.000676873 | IL7R/PDCD1/FASLG/CCL5 | 4 | BP |
| GO:1903975 | regulation of glial cell migration | 3/88 | 19/18723 | 9.20655E-05 | 0.00097631 | 0.000692613 | GPR183/IDH2/CCL3 | 3 | BP |
| GO:1903978 | regulation of microglial cell activation | 3/88 | 19/18723 | 9.20655E-05 | 0.00097631 | 0.000692613 | CTSC/CCL3/CST7 | 3 | BP |
| GO:0043370 | regulation of CD4-positive, alpha-beta T cell differentiation | 4/88 | 51/18723 | 9.61826E-05 | 0.00101424 | 0.000719521 | TNFSF4/IFNG/HLA-DRA/HLA-DRB1 | 4 | BP |
| GO:0071356 | cellular response to tumor necrosis factor | 7/88 | 229/18723 | 0.000102339 | 0.001073127 | 0.000761296 | CCL3L1/CCL3L3/VCAM1/CCL3/CCL4L1/CCL4/CCL5 | 7 | BP |
| GO:0002834 | regulation of response to tumor cell | 3/88 | 20/18723 | 0.000107945 | 0.001119403 | 0.000794125 | CRTAM/HAVCR2/HLA-DRB1 | 3 | BP |
| GO:0002837 | regulation of immune response to tumor cell | 3/88 | 20/18723 | 0.000107945 | 0.001119403 | 0.000794125 | CRTAM/HAVCR2/HLA-DRB1 | 3 | BP |
| GO:0008347 | glial cell migration | 4/88 | 53/18723 | 0.0001119 | 0.001147741 | 0.000814229 | GPR183/IDH2/ADGRG1/CCL3 | 4 | BP |
| GO:0033059 | cellular pigmentation | 4/88 | 53/18723 | 0.0001119 | 0.001147741 | 0.000814229 | RAB27A/CD63/ZEB2/LYST | 4 | BP |
| GO:1902106 | negative regulation of leukocyte differentiation | 5/88 | 102/18723 | 0.000119035 | 0.001214291 | 0.000861441 | TNFSF4/ID2/CD74/LAG3/CCL3 | 5 | BP |
| GO:0002823 | negative regulation of adaptive immune response based on somatic recombination of immune receptors built from immunoglobulin superfamily domains | 4/88 | 54/18723 | 0.000120421 | 0.001215213 | 0.000862095 | IL7R/TNFSF4/HAVCR2/KLRD1 | 4 | BP |
| GO:0070228 | regulation of lymphocyte apoptotic process | 4/88 | 54/18723 | 0.000120421 | 0.001215213 | 0.000862095 | IL7R/PDCD1/CD74/CCL5 | 4 | BP |
| GO:0045639 | positive regulation of myeloid cell differentiation | 5/88 | 103/18723 | 0.000124649 | 0.00125116 | 0.000887596 | LEF1/ID2/CD74/IFNG/HLA-DRB1 | 5 | BP |
| GO:0046641 | positive regulation of alpha-beta T cell proliferation | 3/88 | 21/18723 | 0.000125508 | 0.001253075 | 0.000888955 | CD28/PTPN22/TNFSF4 | 3 | BP |
| GO:0032635 | interleukin-6 production | 6/88 | 165/18723 | 0.00012763 | 0.001260854 | 0.000894473 | PTPN22/TNFSF4/CD74/F2R/IFNG/HAVCR2 | 6 | BP |
| GO:0032675 | regulation of interleukin-6 production | 6/88 | 165/18723 | 0.00012763 | 0.001260854 | 0.000894473 | PTPN22/TNFSF4/CD74/F2R/IFNG/HAVCR2 | 6 | BP |
| GO:0045620 | negative regulation of lymphocyte differentiation | 4/88 | 55/18723 | 0.000129402 | 0.00127166 | 0.000902139 | TNFSF4/ID2/CD74/LAG3 | 4 | BP |
| GO:0045071 | negative regulation of viral genome replication | 4/88 | 56/18723 | 0.000138858 | 0.001357484 | 0.000963024 | OASL/APOBEC3C/APOBEC3G/CCL5 | 4 | BP |
| GO:1903707 | negative regulation of hemopoiesis | 5/88 | 106/18723 | 0.000142715 | 0.00138796 | 0.000984644 | TNFSF4/ID2/CD74/LAG3/CCL3 | 5 | BP |
| GO:0045061 | thymic T cell selection | 3/88 | 22/18723 | 0.000144832 | 0.001401283 | 0.000994096 | CCR7/CD28/CD74 | 3 | BP |
| GO:0071216 | cellular response to biotic stimulus | 7/88 | 246/18723 | 0.00015946 | 0.001534907 | 0.001088892 | PTPN22/TNFSF4/TNIP3/CXCL13/CCL3/HAVCR2/CCL5 | 7 | BP |
| GO:0002820 | negative regulation of adaptive immune response | 4/88 | 59/18723 | 0.00017023 | 0.001630211 | 0.001156502 | IL7R/TNFSF4/HAVCR2/KLRD1 | 4 | BP |

| | | | | | | | | | |
|---|---|---|---|---|---|---|---|---|---|
| GO:0051851 | modulation by host of symbiont process | 4/88 | 60/18723 | 0.000181738 | 0.001731588 | 0.001228421 | LEF1/CCL3/CCL4/CCL5 | 4 | BP |
| GO:0001911 | negative regulation of leukocyte mediated cytotoxicity | 3/88 | 24/18723 | 0.000189061 | 0.001787004 | 0.001267734 | IL7R/HAVCR2/KLRD1 | 3 | BP |
| GO:0034612 | response to tumor necrosis factor | 7/88 | 253/18723 | 0.000189459 | 0.001787004 | 0.001267734 | CCL3L1/CCL3L3/VCAM1/CCL3/CCL4L1/CCL4/CCL5 | 7 | BP |
| GO:2000401 | regulation of lymphocyte migration | 4/88 | 61/18723 | 0.000193798 | 0.001818794 | 0.001290286 | CXCL13/CCL3/CCL4/CCL5 | 4 | BP |
| GO:0032615 | interleukin-12 production | 4/88 | 62/18723 | 0.000206424 | 0.001918111 | 0.001360744 | LTB/CCR7/TNFSF4/IFNG | 4 | BP |
| GO:0032655 | regulation of interleukin-12 production | 4/88 | 62/18723 | 0.000206424 | 0.001918111 | 0.001360744 | LTB/CCR7/TNFSF4/IFNG | 4 | BP |
| GO:0071675 | regulation of mononuclear cell migration | 5/88 | 115/18723 | 0.000209101 | 0.001933411 | 0.001371598 | CCR7/CXCL13/CCL3/CCL4/CCL5 | 5 | BP |
| GO:0032753 | positive regulation of interleukin-4 production | 3/88 | 25/18723 | 0.000214113 | 0.001970048 | 0.001397589 | CD28/TNFSF4/HAVCR2 | 3 | BP |
| GO:0002702 | positive regulation of production of molecular mediator of immune response | 5/88 | 117/18723 | 0.000226604 | 0.002074804 | 0.001471905 | CD28/PTPN22/TNFSF4/CD244/CD74 | 5 | BP |
| GO:0002407 | dendritic cell chemotaxis | 3/88 | 26/18723 | 0.00024122 | 0.002187292 | 0.001551706 | GPR183/CCR7/CCL5 | 3 | BP |
| GO:0019883 | antigen processing and presentation of endogenous antigen | 3/88 | 26/18723 | 0.00024122 | 0.002187292 | 0.001551706 | CD74/HLA-DRA/HLA-DRB1 | 3 | BP |
| GO:0048524 | positive regulation of viral process | 4/88 | 65/18723 | 0.00024786 | 0.002236697 | 0.001586755 | CD28/CD74/HLA-DRB1/CCL5 | 4 | BP |
| GO:0002562 | somatic diversification of immune receptors via germline recombination within a single locus | 4/88 | 66/18723 | 0.00026291 | 0.002338778 | 0.001659173 | LEF1/CD28/TCF7/TNFSF4 | 4 | BP |
| GO:0016444 | somatic cell DNA recombination | 4/88 | 66/18723 | 0.00026291 | 0.002338778 | 0.001659173 | LEF1/CD28/TCF7/TNFSF4 | 4 | BP |
| GO:0072678 | T cell migration | 4/88 | 66/18723 | 0.00026291 | 0.002338778 | 0.001659173 | GPR183/CXCL13/CCL3/CCL5 | 4 | BP |
| GO:0036037 | CD8-positive, alpha-beta T cell activation | 3/88 | 27/18723 | 0.000270452 | 0.002394521 | 0.001698718 | PTPN22/EOMES/CRTAM | 3 | BP |
| GO:2000514 | regulation of CD4-positive, alpha-beta T cell activation | 4/88 | 67/18723 | 0.000278606 | 0.002455132 | 0.001741717 | TNFSF4/IFNG/HLA-DRA/HLA-DRB1 | 4 | BP |
| GO:0098636 | protein complex involved in cell adhesion | 3/89 | 36/19550 | 0.000584098 | 0.00253109 | 0.001844519 | CD28/ITGA1/ITGAE | 3 | CC |
| GO:0048872 | homeostasis of number of cells | 7/88 | 272/18723 | 0.000294486 | 0.002582946 | 0.00183239 | IL7R/GPR183/CCR7/ID2/CD74/F2R/ADGRG1 | 7 | BP |
| GO:0031342 | negative regulation of cell killing | 3/88 | 28/18723 | 0.000301879 | 0.002635477 | 0.001869657 | IL7R/HAVCR2/KLRD1 | 3 | BP |
| GO:0050777 | negative regulation of immune response | 6/88 | 194/18723 | 0.000306773 | 0.002665796 | 0.001891166 | IL7R/TNFSF4/PDCD1/HAVCR2/KLRD1/HLA-DRB1 | 6 | BP |
| GO:0032612 | interleukin-1 production | 5/88 | 128/18723 | 0.000343644 | 0.00295881 | 0.002099035 | CCR7/F2R/IFNG/CCL3/HAVCR2 | 5 | BP |
| GO:0032652 | regulation of interleukin-1 production | 5/88 | 128/18723 | 0.000343644 | 0.00295881 | 0.002099035 | CCR7/F2R/IFNG/CCL3/HAVCR2 | 5 | BP |
| GO:0032722 | positive regulation of chemokine production | 4/88 | 71/18723 | 0.000348169 | 0.002984075 | 0.002116958 | TNFSF4/CD74/IFNG/HAVCR2 | 4 | BP |
| GO:0043032 | positive regulation of macrophage activation | 3/88 | 30/18723 | 0.000371592 | 0.003156013 | 0.002238935 | CTSC/CCL3/HAVCR2 | 3 | BP |
| GO:0070229 | negative regulation of lymphocyte apoptotic process | 3/88 | 30/18723 | 0.000371592 | 0.003156013 | 0.002238935 | IL7R/CD74/CCL5 | 3 | BP |
| GO:0019722 | calcium-mediated signaling | 6/88 | 202/18723 | 0.000380623 | 0.003218153 | 0.002283018 | CCR7/CXCR6/PLEK/VCAM1/CCL3/CCL4 | 6 | BP |
| GO:0002437 | inflammatory response to antigenic stimulus | 4/88 | 74/18723 | 0.000407915 | 0.003433441 | 0.002435747 | CCR7/CD28/RBPJ/HLA-DRB1 | 4 | BP |
| GO:0002323 | natural killer cell activation involved in immune response | 3/88 | 31/18723 | 0.000410012 | 0.00343568 | 0.002437336 | RAB27A/CD244/ZNF683 | 3 | BP |
| GO:0033077 | T cell differentiation in thymus | 4/88 | 75/18723 | 0.000429362 | 0.003581831 | 0.002541018 | IL7R/CCR7/CD28/CD74 | 4 | BP |
| GO:0002685 | regulation of leukocyte migration | 6/88 | 210/18723 | 0.000467719 | 0.003867437 | 0.002743632 | CCR7/CD74/CXCL13/CCL3/CCL4/CCL5 | 6 | BP |
| GO:0045637 | regulation of myeloid cell differentiation | 6/88 | 210/18723 | 0.000467719 | 0.003867437 | 0.002743632 | LEF1/ID2/CD74/IFNG/CCL3/HLA-DRB1 | 6 | BP |
| GO:0002200 | somatic diversification of immune receptors | 4/88 | 77/18723 | 0.00047465 | 0.003907534 | 0.002772078 | LEF1/CD28/TCF7/TNFSF4 | 4 | BP |
| GO:0036336 | dendritic cell migration | 3/88 | 33/18723 | 0.000494304 | 0.004051566 | 0.002874256 | GPR183/CCR7/CCL5 | 3 | BP |
| GO:0010922 | positive regulation of phosphatase activity | 3/88 | 34/18723 | 0.000540304 | 0.004371339 | 0.00310111 | PLEK/ITGA1/IFNG | 3 | BP |
| GO:0050931 | pigment cell differentiation | 3/88 | 34/18723 | 0.000540304 | 0.004371339 | 0.00310111 | RAB27A/CD63/ZEB2 | 3 | BP |
| GO:0070232 | regulation of T cell apoptotic process | 3/88 | 34/18723 | 0.000540304 | 0.004371339 | 0.00310111 | IL7R/PDCD1/CCL5 | 3 | BP |
| GO:0051817 | modulation of process of other organism involved in symbiotic interaction | 4/88 | 81/18723 | 0.000575277 | 0.004634312 | 0.003287668 | LEF1/CCL3/CCL4/CCL5 | 4 | BP |
| GO:0032814 | regulation of natural killer cell activation | 3/88 | 35/18723 | 0.000588957 | 0.004704136 | 0.003337202 | PTPN22/ZNF683/HAVCR2 | 3 | BP |
| GO:0046640 | regulation of alpha-beta T cell proliferation | 3/88 | 35/18723 | 0.000588957 | 0.004704136 | 0.003337202 | CD28/PTPN22/TNFSF4 | 3 | BP |
| GO:0046633 | alpha-beta T cell proliferation | 3/88 | 38/18723 | 0.000751485 | 0.005976485 | 0.004239831 | CD28/PTPN22/TNFSF4 | 3 | BP |
| GO:0005164 | tumor necrosis factor receptor binding | 3/89 | 31/18368 | 0.000448059 | 0.006410695 | 0.004788974 | LTB/TNFSF4/FASLG | 3 | MF |
| GO:0045622 | regulation of T-helper cell differentiation | 3/88 | 39/18723 | 0.000811305 | 0.006425397 | 0.004558297 | TNFSF4/HLA-DRA/HLA-DRB1 | 3 | BP |
| GO:1990907 | beta-catenin-TCF complex | 2/89 | 13/19550 | 0.001547121 | 0.006436024 | 0.00469022 | LEF1/TCF7 | 2 | CC |
| GO:0150077 | regulation of neuroinflammatory response | 3/88 | 40/18723 | 0.00087412 | 0.006893792 | 0.004890585 | CTSC/CCL3/CST7 | 3 | BP |
| GO:0048525 | negative regulation of viral process | 4/88 | 92/18723 | 0.000929006 | 0.007296005 | 0.005175922 | OASL/APOBEC3C/APOBEC3G/CCL5 | 4 | BP |
| GO:0050856 | regulation of T cell receptor signaling pathway | 3/88 | 41/18723 | 0.00093994 | 0.007351112 | 0.005215017 | CCR7/PTPN22/SH2D1A | 3 | BP |
| GO:0001915 | negative regulation of T cell mediated cytotoxicity | 2/88 | 10/18723 | 0.000959054 | 0.007408001 | 0.005255375 | IL7R/KLRD1 | 2 | BP |
| GO:0070383 | DNA cytosine deamination | 2/88 | 10/18723 | 0.000959054 | 0.007408001 | 0.005255375 | APOBEC3C/APOBEC3G | 2 | BP |
| GO:1903980 | positive regulation of microglial cell activation | 2/88 | 10/18723 | 0.000959054 | 0.007408001 | 0.005255375 | CTSC/CCL3 | 2 | BP |
| GO:0032755 | positive regulation of interleukin-6 production | 4/88 | 93/18723 | 0.000967309 | 0.007441143 | 0.005278886 | TNFSF4/CD74/F2R/IFNG | 4 | BP |
| GO:0001664 | G protein-coupled receptor binding | 7/89 | 295/18368 | 0.000574839 | 0.007637144 | 0.005705167 | CCL3L1/CCL3L3/CXCL13/CCL3/CCL4L1/CCL4/CCL5 | 7 | MF |
| GO:0019080 | viral gene expression | 4/88 | 94/18723 | 0.00100671 | 0.007666219 | 0.005438559 | LEF1/CCL3/CCL4/CCL5 | 4 | BP |
| GO:0051702 | biological process involved in interaction with symbiont | 4/88 | 94/18723 | 0.00100671 | 0.007666219 | 0.005438559 | LEF1/CCL3/CCL4/CCL5 | 4 | BP |
| GO:0032689 | negative regulation of interferon-gamma production | 3/88 | 42/18723 | 0.00100882 | 0.007666219 | 0.005438559 | TNFSF4/HAVCR2/HLA-DRB1 | 3 | BP |
| GO:0061629 | RNA polymerase II-specific DNA-binding transcription factor binding | 7/89 | 299/18368 | 0.000622519 | 0.007719231 | 0.005766489 | LEF1/GABARAPL1/BHLHE40/RBPJ/ID2/EOMES/OASL | 7 | MF |
| GO:0002700 | regulation of production of molecular mediator of immune response | 5/88 | 164/18723 | 0.00105817 | 0.008008814 | 0.005681603 | CD28/PTPN22/TNFSF4/CD244/CD74 | 5 | BP |
| GO:0002861 | regulation of inflammatory response to antigenic stimulus | 3/88 | 43/18723 | 0.001080816 | 0.008147357 | 0.005779888 | CCR7/CD28/HLA-DRB1 | 3 | BP |
| GO:1901216 | positive regulation of neuron death | 4/88 | 97/18723 | 0.001131673 | 0.008390392 | 0.005952301 | ITGA1/FASLG/IFNG/CCL3 | 4 | BP |
| GO:0043122 | regulation of I-kappaB kinase/NF-kappaB signaling | 6/88 | 249/18723 | 0.001136716 | 0.008390392 | 0.005952301 | CCR7/TNIP3/CD74/F2R/FASLG/HLA-DRB1 | 6 | BP |
| GO:0031295 | T cell costimulation | 3/88 | 44/18723 | 0.00115598 | 0.008390392 | 0.005952301 | CCR7/CD28/TNFSF4 | 3 | BP |
| GO:0048066 | developmental pigmentation | 3/88 | 44/18723 | 0.00115598 | 0.008390392 | 0.005952301 | RAB27A/CD63/ZEB2 | 3 | BP |
| GO:0002357 | defense response to tumor cell | 2/88 | 11/18723 | 0.001168597 | 0.008390392 | 0.005952301 | ABI3/PRF1 | 2 | BP |
| GO:0002604 | regulation of dendritic cell antigen processing and presentation | 2/88 | 11/18723 | 0.001168597 | 0.008390392 | 0.005952301 | CCR7/CD74 | 2 | BP |
| GO:0033625 | positive regulation of integrin activation | 2/88 | 11/18723 | 0.001168597 | 0.008390392 | 0.005952301 | PLEK/CXCL13 | 2 | BP |
| GO:0033632 | regulation of cell-cell adhesion mediated by integrin | 2/88 | 11/18723 | 0.001168597 | 0.008390392 | 0.005952301 | CXCL13/CCL5 | 2 | BP |
| GO:0045657 | positive regulation of monocyte differentiation | 2/88 | 11/18723 | 0.001168597 | 0.008390392 | 0.005952301 | CD74/HLA-DRB1 | 2 | BP |
| GO:0046598 | positive regulation of viral entry into host cell | 2/88 | 11/18723 | 0.001168597 | 0.008390392 | 0.005952301 | CD74/HLA-DRB1 | 2 | BP |
| GO:0046643 | regulation of gamma-delta T cell activation | 2/88 | 11/18723 | 0.001168597 | 0.008390392 | 0.005952301 | LEF1/TCF7 | 2 | BP |
| GO:0075294 | positive regulation by symbiont of entry into host | 2/88 | 11/18723 | 0.001168597 | 0.008390392 | 0.005952301 | CD74/HLA-DRB1 | 2 | BP |

| | | | | | | | | | |
|---|---|---|---|---|---|---|---|---|---|
| GO:0032642 | regulation of chemokine production | 4/88 | 98/18723 | 0.001175638 | 0.008390392 | 0.005952301 | TNFSF4/CD74/IFNG/HAVCR2 | 4 | BP |
| GO:0043473 | pigmentation | 4/88 | 98/18723 | 0.001175638 | 0.008390392 | 0.005952301 | RAB27A/CD63/ZEB2/LYST | 4 | BP |
| GO:0032602 | chemokine production | 4/88 | 99/18723 | 0.001220787 | 0.008646859 | 0.006134244 | TNFSF4/CD74/IFNG/HAVCR2 | 4 | BP |
| GO:0042100 | B cell proliferation | 4/88 | 99/18723 | 0.001220787 | 0.008646859 | 0.006134244 | IL7R/GPR183/LEF1/CD74 | 4 | BP |
| GO:0031294 | lymphocyte costimulation | 3/88 | 46/18723 | 0.001316023 | 0.00928637 | 0.006587925 | CCR7/CD28/TNFSF4 | 3 | BP |
| GO:0032585 | multivesicular body membrane | 2/89 | 16/19550 | 0.002359148 | 0.00943659 | 0.006876868 | RAB27A/CD63 | 2 | CC |
| GO:0032677 | regulation of interleukin-8 production | 4/88 | 102/18723 | 0.001363508 | 0.009562484 | 0.006783805 | PTPN22/CD244/CD74/F2R | 4 | BP |
| GO:0002863 | positive regulation of inflammatory response to antigenic stimulus | 2/88 | 12/18723 | 0.001398035 | 0.009562484 | 0.006783805 | CCR7/CD28 | 2 | BP |
| GO:0006216 | cytidine catabolic process | 2/88 | 12/18723 | 0.001398035 | 0.009562484 | 0.006783805 | APOBEC3C/APOBEC3G | 2 | BP |
| GO:0009972 | cytidine deamination | 2/88 | 12/18723 | 0.001398035 | 0.009562484 | 0.006783805 | APOBEC3C/APOBEC3G | 2 | BP |
| GO:0016554 | cytidine to uridine editing | 2/88 | 12/18723 | 0.001398035 | 0.009562484 | 0.006783805 | APOBEC3C/APOBEC3G | 2 | BP |
| GO:0034154 | toll-like receptor 7 signaling pathway | 2/88 | 12/18723 | 0.001398035 | 0.009562484 | 0.006783805 | PTPN22/HAVCR2 | 2 | BP |
| GO:0046087 | cytidine metabolic process | 2/88 | 12/18723 | 0.001398035 | 0.009562484 | 0.006783805 | APOBEC3C/APOBEC3G | 2 | BP |
| GO:0070493 | thrombin-activated receptor signaling pathway | 2/88 | 12/18723 | 0.001398035 | 0.009562484 | 0.006783805 | PLEK/F2R | 2 | BP |
| GO:0045581 | negative regulation of T cell differentiation | 3/88 | 47/18723 | 0.001401003 | 0.009562484 | 0.006783805 | TNFSF4/CD74/LAG3 | 3 | BP |
| GO:0032637 | interleukin-8 production | 4/88 | 103/18723 | 0.001413563 | 0.00961325 | 0.006819819 | PTPN22/CD244/CD74/F2R | 4 | BP |
| GO:0030225 | macrophage differentiation | 3/88 | 49/18723 | 0.001581132 | 0.010675487 | 0.00757339 | ID2/IFNG/HLA-DRB1 | 3 | BP |
| GO:1904894 | positive regulation of receptor signaling pathway via STAT | 3/88 | 49/18723 | 0.001581132 | 0.010675487 | 0.00757339 | IL7R/F2R/CCL5 | 3 | BP |
| GO:0002824 | positive regulation of adaptive immune response based on somatic recombination of immune receptors built from immunoglobulin superfamily domains | 4/88 | 107/18723 | 0.001626642 | 0.010886475 | 0.007723069 | CD28/TNFSF4/HLA-DRA/HLA-DRB1 | 4 | BP |
| GO:0007229 | integrin-mediated signaling pathway | 4/88 | 107/18723 | 0.001626642 | 0.010886475 | 0.007723069 | PLEK/ITGA1/CD63/ITGAE | 4 | BP |
| GO:0001768 | establishment of T cell polarity | 2/88 | 13/18723 | 0.001647181 | 0.010886475 | 0.007723069 | CCR7/CRTAM | 2 | BP |
| GO:0042492 | gamma-delta T cell differentiation | 2/88 | 13/18723 | 0.001647181 | 0.010886475 | 0.007723069 | LEF1/TCF7 | 2 | BP |
| GO:0051709 | regulation of killing of cells of other organism | 2/88 | 13/18723 | 0.001647181 | 0.010886475 | 0.007723069 | IFNG/PRF1 | 2 | BP |
| GO:0070234 | positive regulation of T cell apoptotic process | 2/88 | 13/18723 | 0.001647181 | 0.010886475 | 0.007723069 | PDCD1/CCL5 | 2 | BP |
| GO:0071622 | regulation of granulocyte chemotaxis | 3/88 | 51/18723 | 0.00177514 | 0.011691008 | 0.00829382 | CCR7/CD74/CCL5 | 3 | BP |
| GO:0002698 | negative regulation of immune effector process | 4/88 | 110/18723 | 0.00180043 | 0.011707161 | 0.008305279 | IL7R/TNFSF4/HAVCR2/KLRD1 | 4 | BP |
| GO:0032611 | interleukin-1 beta production | 4/88 | 110/18723 | 0.00180043 | 0.011707161 | 0.008305279 | CCR7/F2R/IFNG/CCL3 | 4 | BP |
| GO:0032651 | regulation of interleukin-1 beta production | 4/88 | 110/18723 | 0.00180043 | 0.011707161 | 0.008305279 | CCR7/F2R/IFNG/CCL3 | 4 | BP |
| GO:0043433 | negative regulation of DNA-binding transcription factor activity | 5/88 | 185/18723 | 0.001802541 | 0.011707161 | 0.008305279 | TNFSF4/BHLHE40/ID2/EOMES/HAVCR2 | 5 | BP |
| GO:0047844 | deoxycytidine deaminase activity | 2/89 | 10/18368 | 0.001018613 | 0.011841381 | 0.008845853 | APOBEC3C/APOBEC3G | 2 | MF |
| GO:0043123 | positive regulation of I-kappaB kinase/NF-kappaB signaling | 5/88 | 186/18723 | 0.001845576 | 0.011945331 | 0.008474242 | CCR7/CD74/F2R/FASLG/HLA-DRB1 | 5 | BP |
| GO:0014066 | regulation of phosphatidylinositol 3-kinase signaling | 4/88 | 111/18723 | 0.001861111 | 0.012004487 | 0.008516208 | CD28/PIK3IP1/F2R/CCL5 | 4 | BP |
| GO:0001767 | establishment of lymphocyte polarity | 2/88 | 14/18723 | 0.001915848 | 0.012189991 | 0.008647808 | CCR7/CRTAM | 2 | BP |
| GO:0045006 | DNA deamination | 2/88 | 14/18723 | 0.001915848 | 0.012189991 | 0.008647808 | APOBEC3C/APOBEC3G | 2 | BP |
| GO:0046131 | pyrimidine ribonucleoside metabolic process | 2/88 | 14/18723 | 0.001915848 | 0.012189991 | 0.008647808 | APOBEC3C/APOBEC3G | 2 | BP |
| GO:0046133 | pyrimidine ribonucleoside catabolic process | 2/88 | 14/18723 | 0.001915848 | 0.012189991 | 0.008647808 | APOBEC3C/APOBEC3G | 2 | BP |
| GO:0002821 | positive regulation of adaptive immune response | 4/88 | 112/18723 | 0.001923197 | 0.012195406 | 0.008651649 | CD28/TNFSF4/HLA-DRA/HLA-DRB1 | 4 | BP |
| GO:0002707 | negative regulation of lymphocyte mediated immunity | 3/88 | 53/18723 | 0.001983399 | 0.012534815 | 0.008892432 | IL7R/HAVCR2/KLRD1 | 3 | BP |
| GO:1904892 | regulation of receptor signaling pathway via STAT | 4/88 | 114/18723 | 0.002051647 | 0.012922622 | 0.00916755 | IL7R/F2R/IFNG/CCL5 | 4 | BP |
| GO:0042608 | T cell receptor binding | 2/89 | 11/18368 | 0.001241052 | 0.013081646 | 0.009772367 | HLA-DRA/HLA-DRB1 | 2 | MF |
| GO:0004896 | cytokine receptor activity | 4/89 | 97/18368 | 0.001265966 | 0.013081646 | 0.009772367 | IL7R/CCR7/CXCR6/CD74 | 4 | MF |
| GO:0007249 | I-kappaB kinase/NF-kappaB signaling | 6/88 | 281/18723 | 0.002098905 | 0.013176068 | 0.00934735 | CCR7/TNIP3/CD74/F2R/FASLG/HLA-DRB1 | 6 | BP |
| GO:0051209 | release of sequestered calcium ion into cytosol | 4/88 | 115/18723 | 0.002118045 | 0.013251902 | 0.009401147 | CCR7/F2R/FASLG/CCL3 | 4 | BP |
| GO:0051283 | negative regulation of sequestering of calcium ion | 4/88 | 116/18723 | 0.002185914 | 0.013607338 | 0.009653301 | CCR7/F2R/FASLG/CCL3 | 4 | BP |
| GO:0002836 | positive regulation of response to tumor cell | 2/88 | 15/18723 | 0.002203852 | 0.013607338 | 0.009653301 | CRTAM/HLA-DRB1 | 2 | BP |
| GO:0002839 | positive regulation of immune response to tumor cell | 2/88 | 15/18723 | 0.002203852 | 0.013607338 | 0.009653301 | CRTAM/HLA-DRB1 | 2 | BP |
| GO:0010820 | positive regulation of T cell chemotaxis | 2/88 | 15/18723 | 0.002203852 | 0.013607338 | 0.009653301 | CXCL13/CCL5 | 2 | BP |
| GO:0002711 | positive regulation of T cell mediated immunity | 3/88 | 56/18723 | 0.002323293 | 0.014221973 | 0.010089334 | TNFSF4/HLA-DRA/HLA-DRB1 | 3 | BP |
| GO:0031529 | ruffle organization | 3/88 | 56/18723 | 0.002323293 | 0.014221973 | 0.010089334 | CCR7/PLEK/ABI3 | 3 | BP |
| GO:0051282 | regulation of sequestering of calcium ion | 4/88 | 118/18723 | 0.00232613 | 0.014221973 | 0.010089334 | CCR7/F2R/FASLG/CCL3 | 4 | BP |
| GO:0004126 | cytidine deaminase activity | 2/89 | 12/18368 | 0.001484574 | 0.014389734 | 0.010749547 | APOBEC3C/APOBEC3G | 2 | MF |
| GO:0004252 | serine-type endopeptidase activity | 5/89 | 174/18368 | 0.001573719 | 0.014389734 | 0.010749547 | CTSC/GZMK/GZMA/GZMB/GZMH | 5 | MF |
| GO:0015026 | coreceptor activity | 3/89 | 48/18368 | 0.001624647 | 0.014389734 | 0.010749547 | CD28/CXCR6/CD8A | 3 | MF |
| GO:0032813 | tumor necrosis factor receptor superfamily binding | 3/89 | 49/18368 | 0.001724578 | 0.014580526 | 0.010892074 | LTB/TNFSF4/FASLG | 3 | MF |
| GO:0050864 | regulation of B cell activation | 5/88 | 198/18723 | 0.002422645 | 0.014763974 | 0.010473839 | GPR183/CD28/TNFSF4/ID2/CD74 | 5 | BP |
| GO:0044403 | biological process involved in symbiotic interaction | 6/88 | 290/18723 | 0.002456635 | 0.014868028 | 0.010547658 | LEF1/CD74/CCL3/HLA-DRB1/CCL4/CCL5 | 6 | BP |
| GO:0002830 | positive regulation of type 2 immune response | 2/88 | 16/18723 | 0.00251101 | 0.014868028 | 0.010547658 | TNFSF4/CD74 | 2 | BP |
| GO:0010819 | regulation of T cell chemotaxis | 2/88 | 16/18723 | 0.00251101 | 0.014868028 | 0.010547658 | CXCL13/CCL5 | 2 | BP |
| GO:0010919 | regulation of inositol phosphate biosynthetic process | 2/88 | 16/18723 | 0.00251101 | 0.014868028 | 0.010547658 | CD244/PLEK | 2 | BP |
| GO:0033631 | cell-cell adhesion mediated by integrin | 2/88 | 16/18723 | 0.00251101 | 0.014868028 | 0.010547658 | CXCL13/CCL5 | 2 | BP |
| GO:0034138 | toll-like receptor 3 signaling pathway | 2/88 | 16/18723 | 0.00251101 | 0.014868028 | 0.010547658 | PTPN22/HAVCR2 | 2 | BP |
| GO:0045869 | negative regulation of single stranded viral RNA replication via double stranded DNA intermediate | 2/88 | 16/18723 | 0.00251101 | 0.014868028 | 0.010547658 | APOBEC3C/APOBEC3G | 2 | BP |
| GO:0046135 | pyrimidine nucleoside catabolic process | 2/88 | 16/18723 | 0.00251101 | 0.014868028 | 0.010547658 | APOBEC3C/APOBEC3G | 2 | BP |
| GO:0070230 | positive regulation of lymphocyte apoptotic process | 2/88 | 16/18723 | 0.00251101 | 0.014868028 | 0.010547658 | PDCD1/CCL5 | 2 | BP |
| GO:0043525 | positive regulation of neuron apoptotic process | 3/88 | 58/18723 | 0.002568726 | 0.015161943 | 0.010756166 | ITGA1/FASLG/CCL3 | 3 | BP |
| GO:0051208 | sequestering of calcium ion | 4/88 | 122/18723 | 0.002624936 | 0.015396888 | 0.010922841 | CCR7/F2R/FASLG/CCL3 | 4 | BP |
| GO:0051928 | positive regulation of calcium ion transport | 4/88 | 122/18723 | 0.002624936 | 0.015396888 | 0.010922841 | F2R/CCL3/CCL4/CCL5 | 4 | BP |
| GO:0042827 | platelet dense granule | 2/89 | 21/19550 | 0.004067948 | 0.015669134 | 0.011418802 | CD63/CTSW | 2 | CC |
| GO:0035306 | positive regulation of dephosphorylation | 3/88 | 59/18723 | 0.002697216 | 0.015771574 | 0.01118865 | PLEK/ITGA1/IFNG | 3 | BP |
| GO:0031663 | lipopolysaccharide-mediated signaling pathway | 3/88 | 60/18723 | 0.00282961 | 0.016436134 | 0.011660101 | PTPN22/CCL3/CCL5 | 3 | BP |

| | | | | | | | | | |
|---|---|---|---|---|---|---|---|---|---|
| GO:0033151 | V(D)J recombination | 2/88 | 17/18723 | 0.002837138 | 0.016436134 | 0.011660101 | LEF1/TCF7 | 2 | BP |
| GO:0033623 | regulation of integrin activation | 2/88 | 17/18723 | 0.002837138 | 0.016436134 | 0.011660101 | PLEK/CXCL13 | 2 | BP |
| GO:0030246 | carbohydrate binding | 6/89 | 271/18368 | 0.002039232 | 0.016491181 | 0.012319388 | SELL/KLRG1/CLEC2B/KLRD1/HLA-DRA/HLA-DRB1 | 6 | MF |
| GO:0032613 | interleukin-10 production | 3/88 | 62/18723 | 0.00310626 | 0.017721731 | 0.012572128 | CD28/TNFSF4/HLA-DRB1 | 3 | BP |
| GO:0032623 | interleukin-2 production | 3/88 | 62/18723 | 0.00310626 | 0.017721731 | 0.012572128 | CD28/LAG3/HAVCR2 | 3 | BP |
| GO:0032653 | regulation of interleukin-10 production | 3/88 | 62/18723 | 0.00310626 | 0.017721731 | 0.012572128 | CD28/TNFSF4/HLA-DRB1 | 3 | BP |
| GO:0032663 | regulation of interleukin-2 production | 3/88 | 62/18723 | 0.00310626 | 0.017721731 | 0.012572128 | CD28/LAG3/HAVCR2 | 3 | BP |
| GO:0032757 | positive regulation of interleukin-8 production | 3/88 | 62/18723 | 0.00310626 | 0.017721731 | 0.012572128 | CD244/CD74/F2R | 3 | BP |
| GO:0034162 | toll-like receptor 9 signaling pathway | 2/88 | 18/18723 | 0.003182056 | 0.017936092 | 0.0127242 | PTPN22/HAVCR2 | 2 | BP |
| GO:0045953 | negative regulation of natural killer cell mediated cytotoxicity | 2/88 | 18/18723 | 0.003182056 | 0.017936092 | 0.0127242 | HAVCR2/KLRD1 | 2 | BP |
| GO:0048535 | lymph node development | 2/88 | 18/18723 | 0.003182056 | 0.017936092 | 0.0127242 | IL7R/LTB | 2 | BP |
| GO:0150078 | positive regulation of neuroinflammatory response | 2/88 | 18/18723 | 0.003182056 | 0.017936092 | 0.0127242 | CTSC/CCL3 | 2 | BP |
| GO:0002704 | negative regulation of leukocyte mediated immunity | 3/88 | 63/18723 | 0.003250591 | 0.018213013 | 0.012920653 | IL7R/HAVCR2/KLRD1 | 3 | BP |
| GO:0050854 | regulation of antigen receptor-mediated signaling pathway | 3/88 | 63/18723 | 0.003250591 | 0.018213013 | 0.012920653 | CCR7/PTPN22/SH2D1A | 3 | BP |
| GO:0008236 | serine-type peptidase activity | 5/89 | 191/18368 | 0.002364674 | 0.01832622 | 0.013690215 | CTSC/GZMK/GZMA/GZMB/GZMH | 5 | MF |
| GO:0017171 | serine hydrolase activity | 5/89 | 195/18368 | 0.002586471 | 0.019068064 | 0.014244394 | CTSC/GZMK/GZMA/GZMB/GZMH | 5 | MF |
| GO:0016004 | phospholipase activator activity | 2/89 | 16/18368 | 0.002665428 | 0.019068064 | 0.014244394 | CCL3/CCL5 | 2 | MF |
| GO:0002281 | macrophage activation involved in immune response | 2/88 | 19/18723 | 0.003545584 | 0.019402511 | 0.013764505 | IFNG/HAVCR2 | 2 | BP |
| GO:0002483 | antigen processing and presentation of endogenous peptide antigen | 2/88 | 19/18723 | 0.003545584 | 0.019402511 | 0.013764505 | HLA-DRA/HLA-DRB1 | 2 | BP |
| GO:0002716 | negative regulation of natural killer cell mediated immunity | 2/88 | 19/18723 | 0.003545584 | 0.019402511 | 0.013764505 | HAVCR2/KLRD1 | 2 | BP |
| GO:0032616 | interleukin-13 production | 2/88 | 19/18723 | 0.003545584 | 0.019402511 | 0.013764505 | LEF1/TNFSF4 | 2 | BP |
| GO:0032656 | regulation of interleukin-13 production | 2/88 | 19/18723 | 0.003545584 | 0.019402511 | 0.013764505 | LEF1/TNFSF4 | 2 | BP |
| GO:0045091 | regulation of single stranded viral RNA replication via double stranded DNA intermediate | 2/88 | 19/18723 | 0.003545584 | 0.019402511 | 0.013764505 | APOBEC3C/APOBEC3G | 2 | BP |
| GO:0070233 | negative regulation of T cell apoptotic process | 2/88 | 19/18723 | 0.003545584 | 0.019402511 | 0.013764505 | IL7R/CCL5 | 2 | BP |
| GO:2001185 | regulation of CD8-positive, alpha-beta T cell activation | 2/88 | 19/18723 | 0.003545584 | 0.019402511 | 0.013764505 | PTPN22/CRTAM | 2 | BP |
| GO:0007596 | blood coagulation | 5/88 | 217/18723 | 0.003590057 | 0.019588772 | 0.013896641 | DGKA/RAB27A/PLEK/ANXA5/F2R | 5 | BP |
| GO:0042130 | negative regulation of T cell proliferation | 3/88 | 67/18723 | 0.003868801 | 0.021002849 | 0.014899815 | CRTAM/HAVCR2/HLA-DRB1 | 3 | BP |
| GO:0002577 | regulation of antigen processing and presentation | 2/88 | 20/18723 | 0.003927544 | 0.021002849 | 0.014899815 | CCR7/CD74 | 2 | BP |
| GO:0016553 | base conversion or substitution editing | 2/88 | 20/18723 | 0.003927544 | 0.021002849 | 0.014899815 | APOBEC3C/APOBEC3G | 2 | BP |
| GO:0039692 | single stranded viral RNA replication via double stranded DNA intermediate | 2/88 | 20/18723 | 0.003927544 | 0.021002849 | 0.014899815 | APOBEC3C/APOBEC3G | 2 | BP |
| GO:0045063 | T-helper 1 cell differentiation | 2/88 | 20/18723 | 0.003927544 | 0.021002849 | 0.014899815 | LEF1/TNFSF4 | 2 | BP |
| GO:0046629 | gamma-delta T cell activation | 2/88 | 20/18723 | 0.003927544 | 0.021002849 | 0.014899815 | LEF1/TCF7 | 2 | BP |
| GO:1990182 | exosomal secretion | 2/88 | 20/18723 | 0.003927544 | 0.021002849 | 0.014899815 | RAB27A/IFNG | 2 | BP |
| GO:0007599 | hemostasis | 5/88 | 222/18723 | 0.003954658 | 0.021028026 | 0.014917676 | DGKA/RAB27A/PLEK/ANXA5/F2R | 5 | BP |
| GO:0050817 | coagulation | 5/88 | 222/18723 | 0.003954658 | 0.021028026 | 0.014917676 | DGKA/RAB27A/PLEK/ANXA5/F2R | 5 | BP |
| GO:0009988 | cell-cell recognition | 3/88 | 68/18723 | 0.004033747 | 0.021327729 | 0.01513029 | CCR7/HAVCR2/PRF1 | 3 | BP |
| GO:0031640 | killing of cells of other organism | 3/88 | 68/18723 | 0.004033747 | 0.021327729 | 0.01513029 | GNLY/IFNG/PRF1 | 3 | BP |
| GO:0033630 | positive regulation of cell adhesion mediated by integrin | 2/88 | 21/18723 | 0.004327758 | 0.022564451 | 0.016007644 | CXCL13/CCL5 | 2 | BP |
| GO:0042454 | ribonucleoside catabolic process | 2/88 | 21/18723 | 0.004327758 | 0.022564451 | 0.016007644 | APOBEC3C/APOBEC3G | 2 | BP |
| GO:0045655 | regulation of monocyte differentiation | 2/88 | 21/18723 | 0.004327758 | 0.022564451 | 0.016007644 | CD74/HLA-DRB1 | 2 | BP |
| GO:0070269 | pyroptosis | 2/88 | 21/18723 | 0.004327758 | 0.022564451 | 0.016007644 | GZMA/GZMB | 2 | BP |
| GO:0097734 | extracellular exosome biogenesis | 2/88 | 21/18723 | 0.004327758 | 0.022564451 | 0.016007644 | RAB27A/IFNG | 2 | BP |
| GO:0045123 | cellular extravasation | 3/88 | 70/18723 | 0.004376344 | 0.022754565 | 0.016142514 | SELL/ITGA1/VCAM1 | 3 | BP |
| GO:0030183 | B cell differentiation | 4/88 | 141/18723 | 0.004408035 | 0.022856025 | 0.016214492 | GPR183/RBPJ/ID2/VCAM1 | 4 | BP |
| GO:0060229 | lipase activator activity | 2/89 | 18/18368 | 0.003377101 | 0.023264476 | 0.017379235 | CCL3/CCL5 | 2 | MF |
| GO:0097553 | calcium ion transmembrane import into cytosol | 4/88 | 142/18723 | 0.004519779 | 0.023370869 | 0.016579732 | CCR7/F2R/FASLG/CCL3 | 4 | BP |
| GO:0062013 | positive regulation of small molecule metabolic process | 4/88 | 143/18723 | 0.004633419 | 0.023892659 | 0.0169499 | LDLRAP1/CD244/CD74/IFNG | 4 | BP |
| GO:0032069 | regulation of nuclease activity | 2/88 | 22/18723 | 0.004746051 | 0.024156683 | 0.017137204 | OASL/GZMA | 2 | BP |
| GO:0032700 | negative regulation of interleukin-17 production | 2/88 | 22/18723 | 0.004746051 | 0.024156683 | 0.017137204 | TNFSF4/IFNG | 2 | BP |
| GO:0050860 | negative regulation of T cell receptor signaling pathway | 2/88 | 22/18723 | 0.004746051 | 0.024156683 | 0.017137204 | PTPN22/SH2D1A | 2 | BP |
| GO:0014065 | phosphatidylinositol 3-kinase signaling | 4/88 | 144/18723 | 0.00474897 | 0.024156683 | 0.017137204 | CD28/PIK3IP1/F2R/CCL5 | 4 | BP |
| GO:0031644 | regulation of nervous system process | 4/88 | 144/18723 | 0.00474897 | 0.024156683 | 0.017137204 | CTSC/F2R/CCL3/CST7 | 4 | BP |
| GO:0032732 | positive regulation of interleukin-1 production | 3/88 | 73/18723 | 0.004922468 | 0.024904239 | 0.017667533 | IFNG/CCL3/HAVCR2 | 3 | BP |
| GO:0050848 | regulation of calcium-mediated signaling | 3/88 | 73/18723 | 0.004922468 | 0.024904239 | 0.017667533 | PLEK/CCL3/CCL4 | 3 | BP |
| GO:0034695 | response to prostaglandin E | 2/88 | 23/18723 | 0.005182247 | 0.026077957 | 0.018500191 | CCR7/TNFSF4 | 2 | BP |
| GO:0140112 | extracellular vesicle biogenesis | 2/88 | 23/18723 | 0.005182247 | 0.026077957 | 0.018500191 | RAB27A/IFNG | 2 | BP |
| GO:0045834 | positive regulation of lipid metabolic process | 4/88 | 149/18723 | 0.005355886 | 0.026879672 | 0.019068943 | CCR7/LDLRAP1/CD74/IFNG | 4 | BP |
| GO:0042288 | MHC class I protein binding | 2/89 | 20/18368 | 0.004167496 | 0.027684078 | 0.020680805 | CD244/CD8A | 2 | MF |
| GO:0006213 | pyrimidine nucleoside metabolic process | 2/88 | 24/18723 | 0.005636173 | 0.028135897 | 0.019960132 | APOBEC3C/APOBEC3G | 2 | BP |
| GO:0090023 | positive regulation of neutrophil chemotaxis | 2/88 | 24/18723 | 0.005636173 | 0.028135897 | 0.019960132 | CCR7/CD74 | 2 | BP |
| GO:0050871 | positive regulation of B cell activation | 4/88 | 152/18723 | 0.005743881 | 0.028597517 | 0.020287614 | GPR183/CD28/TNFSF4/CD74 | 4 | BP |
| GO:0140416 | transcription regulator inhibitor activity | 2/89 | 21/18368 | 0.004591727 | 0.029450385 | 0.022000288 | LEF1/ID2 | 2 | MF |
| GO:0030318 | melanocyte differentiation | 2/88 | 25/18723 | 0.006107657 | 0.030217361 | 0.021436762 | RAB27A/ZEB2 | 2 | BP |
| GO:0033622 | integrin activation | 2/88 | 25/18723 | 0.006107657 | 0.030217361 | 0.021436762 | PLEK/CXCL13 | 2 | BP |
| GO:0014068 | positive regulation of phosphatidylinositol 3-kinase signaling | 3/88 | 79/18723 | 0.006133625 | 0.030217361 | 0.021436762 | CD28/F2R/CCL5 | 3 | BP |
| GO:1901184 | regulation of ERBB signaling pathway | 3/88 | 79/18723 | 0.006133625 | 0.030217361 | 0.021436762 | RBPJ/ITGA1/FASLG | 3 | BP |
| GO:0033002 | muscle cell proliferation | 5/88 | 248/18723 | 0.006288885 | 0.030901143 | 0.02192185 | LDLRAP1/RBPJ/ID2/IFNG/CCL5 | 5 | BP |
| GO:0002710 | negative regulation of T cell mediated immunity | 2/88 | 26/18723 | 0.006596527 | 0.032076892 | 0.022755948 | IL7R/KLRD1 | 2 | BP |
| GO:0009164 | nucleoside catabolic process | 2/88 | 26/18723 | 0.006596527 | 0.032076892 | 0.022755948 | APOBEC3C/APOBEC3G | 2 | BP |
| GO:1905523 | positive regulation of macrophage migration | 2/88 | 26/18723 | 0.006596527 | 0.032076892 | 0.022755948 | CCL3/CCL5 | 2 | BP |
| GO:2000108 | positive regulation of leukocyte apoptotic process | 2/88 | 26/18723 | 0.006596527 | 0.032076892 | 0.022755948 | PDCD1/CCL5 | 2 | BP |

| | | | | | | | | | |
|---|---|---|---|---|---|---|---|---|---|
| GO:0008305 | integrin complex | 2/89 | 31/19550 | 0.00874594 | 0.032484921 | 0.023673222 | ITGA1/ITGAE | 2 | CC |
| GO:0001892 | embryonic placenta development | 3/88 | 82/18723 | 0.006800104 | 0.032896377 | 0.023337306 | LEF1/RBPJ/EOMES | 3 | BP |
| GO:0002312 | B cell activation involved in immune response | 3/88 | 82/18723 | 0.006800104 | 0.032896377 | 0.023337306 | GPR183/CD28/TNFSF4 | 3 | BP |
| GO:0060402 | calcium ion transport into cytosol | 4/88 | 160/18723 | 0.006869229 | 0.033145352 | 0.023513933 | CCR7/F2R/FASLG/CCL3 | 4 | BP |
| GO:0050672 | negative regulation of lymphocyte proliferation | 3/88 | 83/18723 | 0.007031449 | 0.033665666 | 0.023883054 | CRTAM/HAVCR2/HLA-DRB1 | 3 | BP |
| GO:0002825 | regulation of T-helper 1 type immune response | 2/88 | 27/18723 | 0.007102613 | 0.033665666 | 0.023883054 | TNFSF4/HAVCR2 | 2 | BP |
| GO:0010528 | regulation of transposition | 2/88 | 27/18723 | 0.007102613 | 0.033665666 | 0.023883054 | APOBEC3C/APOBEC3G | 2 | BP |
| GO:0010529 | negative regulation of transposition | 2/88 | 27/18723 | 0.007102613 | 0.033665666 | 0.023883054 | APOBEC3C/APOBEC3G | 2 | BP |
| GO:0032703 | negative regulation of interleukin-2 production | 2/88 | 27/18723 | 0.007102613 | 0.033665666 | 0.023883054 | LAG3/HAVCR2 | 2 | BP |
| GO:0045830 | positive regulation of isotype switching | 2/88 | 27/18723 | 0.007102613 | 0.033665666 | 0.023883054 | CD28/TNFSF4 | 2 | BP |
| GO:0071624 | positive regulation of granulocyte chemotaxis | 2/88 | 27/18723 | 0.007102613 | 0.033665666 | 0.023883054 | CCR7/CD74 | 2 | BP |
| GO:0016493 | C-C chemokine receptor activity | 2/89 | 23/18368 | 0.005497296 | 0.034083236 | 0.025461161 | CCR7/CXCR6 | 2 | MF |
| GO:0010921 | regulation of phosphatase activity | 3/88 | 84/18723 | 0.007267426 | 0.034273766 | 0.024314451 | PLEK/ITGA1/IFNG | 3 | BP |
| GO:0032945 | negative regulation of mononuclear cell proliferation | 3/88 | 84/18723 | 0.007267426 | 0.034273766 | 0.024314451 | CRTAM/HAVCR2/HLA-DRB1 | 3 | BP |
| GO:0031348 | negative regulation of defense response | 5/88 | 258/18723 | 0.007401428 | 0.034818247 | 0.024700717 | AOAH/HAVCR2/CST7/KLRD1/HLA-DRB1 | 5 | BP |
| GO:0007205 | protein kinase C-activating G protein-coupled receptor signaling pathway | 2/88 | 28/18723 | 0.007625746 | 0.035429517 | 0.025134363 | DGKA/F2R | 2 | BP |
| GO:0032438 | melanosome organization | 2/88 | 28/18723 | 0.007625746 | 0.035429517 | 0.025134363 | ZEB2/LYST | 2 | BP |
| GO:0032958 | inositol phosphate biosynthetic process | 2/88 | 28/18723 | 0.007625746 | 0.035429517 | 0.025134363 | CD244/PLEK | 2 | BP |
| GO:0080111 | DNA demethylation | 2/88 | 28/18723 | 0.007625746 | 0.035429517 | 0.025134363 | APOBEC3C/APOBEC3G | 2 | BP |
| GO:1902624 | positive regulation of neutrophil migration | 2/88 | 28/18723 | 0.007625746 | 0.035429517 | 0.025134363 | CCR7/CD74 | 2 | BP |
| GO:0019957 | C-C chemokine binding | 2/89 | 24/18368 | 0.005978254 | 0.035869525 | 0.02679557 | CCR7/CXCR6 | 2 | MF |
| GO:0001773 | myeloid dendritic cell activation | 2/88 | 29/18723 | 0.008165759 | 0.037383242 | 0.026520372 | RBPJ/HAVCR2 | 2 | BP |
| GO:0001916 | positive regulation of T cell mediated cytotoxicity | 2/88 | 29/18723 | 0.008165759 | 0.037383242 | 0.026520372 | HLA-DRA/HLA-DRB1 | 2 | BP |
| GO:0043153 | entrainment of circadian clock by photoperiod | 2/88 | 29/18723 | 0.008165759 | 0.037383242 | 0.026520372 | BHLHE40/ID2 | 2 | BP |
| GO:0048753 | pigment granule organization | 2/88 | 29/18723 | 0.008165759 | 0.037383242 | 0.026520372 | ZEB2/LYST | 2 | BP |
| GO:1903902 | positive regulation of viral life cycle | 2/88 | 29/18723 | 0.008165759 | 0.037383242 | 0.026520372 | CD74/HLA-DRB1 | 2 | BP |
| GO:2000406 | positive regulation of T cell migration | 2/88 | 29/18723 | 0.008165759 | 0.037383242 | 0.026520372 | CXCL13/CCL5 | 2 | BP |
| GO:0030247 | polysaccharide binding | 2/89 | 25/18368 | 0.006477743 | 0.037651882 | 0.028127042 | HLA-DRA/HLA-DRB1 | 2 | MF |
| GO:0001637 | G protein-coupled chemoattractant receptor activity | 2/89 | 26/18368 | 0.006995576 | 0.038269918 | 0.028588733 | CCR7/CXCR6 | 2 | MF |
| GO:0004950 | chemokine receptor activity | 2/89 | 26/18368 | 0.006995576 | 0.038269918 | 0.028588733 | CCR7/CXCR6 | 2 | MF |
| GO:0050708 | regulation of protein secretion | 5/88 | 268/18723 | 0.008644809 | 0.039450854 | 0.027987175 | F2R/IDH2/IFNG/HLA-DRB1/CCL5 | 5 | BP |
| GO:0034694 | response to prostaglandin | 2/88 | 30/18723 | 0.008722485 | 0.039450854 | 0.027987175 | CCR7/TNFSF4 | 2 | BP |
| GO:0045070 | positive regulation of viral genome replication | 2/88 | 30/18723 | 0.008722485 | 0.039450854 | 0.027987175 | CD28/CCL5 | 2 | BP |
| GO:0045940 | positive regulation of steroid metabolic process | 2/88 | 30/18723 | 0.008722485 | 0.039450854 | 0.027987175 | LDLRAP1/IFNG | 2 | BP |
| GO:0050858 | negative regulation of antigen receptor-mediated signaling pathway | 2/88 | 30/18723 | 0.008722485 | 0.039450854 | 0.027987175 | PTPN22/SH2D1A | 2 | BP |
| GO:0070664 | negative regulation of leukocyte proliferation | 3/88 | 90/18723 | 0.008781728 | 0.039623327 | 0.02810953 | CRTAM/HAVCR2/HLA-DRB1 | 3 | BP |
| GO:0031904 | endosome lumen | 2/89 | 35/19550 | 0.011060171 | 0.03966406 | 0.028904983 | CD63/PRF1 | 2 | CC |
| GO:0002221 | pattern recognition receptor signaling pathway | 4/88 | 172/18723 | 0.008816756 | 0.039685973 | 0.028153972 | PTPN22/TNIP3/OASL/HAVCR2 | 4 | BP |
| GO:0002275 | myeloid cell activation involved in immune response | 3/88 | 91/18723 | 0.009050704 | 0.040641557 | 0.028831882 | IFNG/CCL3/HAVCR2 | 3 | BP |
| GO:0002230 | positive regulation of defense response to virus by host | 2/88 | 31/18723 | 0.009295759 | 0.041248557 | 0.029262499 | PTPN22/APOBEC3G | 2 | BP |
| GO:0002828 | regulation of type 2 immune response | 2/88 | 31/18723 | 0.009295759 | 0.041248557 | 0.029262499 | TNFSF4/CD74 | 2 | BP |
| GO:0034656 | nucleobase-containing small molecule catabolic process | 2/88 | 31/18723 | 0.009295759 | 0.041248557 | 0.029262499 | APOBEC3C/APOBEC3G | 2 | BP |
| GO:0035510 | DNA dealkylation | 2/88 | 31/18723 | 0.009295759 | 0.041248557 | 0.029262499 | APOBEC3C/APOBEC3G | 2 | BP |
| GO:0070528 | protein kinase C signaling | 2/88 | 31/18723 | 0.009295759 | 0.041248557 | 0.029262499 | PLEK/ADGRG1 | 2 | BP |
| GO:1905475 | regulation of protein localization to membrane | 4/88 | 175/18723 | 0.00935446 | 0.041411135 | 0.029377835 | LDLRAP1/ABI3/IFNG/GZMB | 4 | BP |
| GO:0001540 | amyloid-beta binding | 3/89 | 84/18368 | 0.007897887 | 0.04197163 | 0.031354019 | LDLRAP1/ITM2C/CD74 | 3 | MF |
| GO:0043270 | positive regulation of ion transport | 5/88 | 275/18723 | 0.009596898 | 0.042384418 | 0.0300683 | F2R/IFNG/CCL3/CCL4/CCL5 | 5 | BP |
| GO:0032196 | transposition | 2/88 | 32/18723 | 0.009885417 | 0.04325604 | 0.030686644 | APOBEC3C/APOBEC3G | 2 | BP |
| GO:0050850 | positive regulation of calcium-mediated signaling | 2/88 | 32/18723 | 0.009885417 | 0.04325604 | 0.030686644 | CCL3/CCL4 | 2 | BP |
| GO:0090022 | regulation of neutrophil chemotaxis | 2/88 | 32/18723 | 0.009885417 | 0.04325604 | 0.030686644 | CCR7/CD74 | 2 | BP |
| GO:0030316 | osteoclast differentiation | 3/88 | 94/18723 | 0.009886436 | 0.04325604 | 0.030686644 | GPR183/IFNG/CCL3 | 3 | BP |
| GO:0048015 | phosphatidylinositol-mediated signaling | 4/88 | 178/18723 | 0.009913133 | 0.04327198 | 0.030697953 | CD28/PIK3IP1/F2R/CCL5 | 4 | BP |
| GO:0042982 | amyloid precursor protein metabolic process | 3/88 | 95/18723 | 0.010174671 | 0.044310576 | 0.031434752 | LDLRAP1/ITM2C/IFNG | 3 | BP |
| GO:0048660 | regulation of smooth muscle cell proliferation | 4/88 | 180/18723 | 0.010297383 | 0.044741177 | 0.031740228 | LDLRAP1/ID2/IFNG/CCL5 | 4 | BP |
| GO:0009648 | photoperiodism | 2/88 | 33/18723 | 0.010491295 | 0.044864496 | 0.031827714 | BHLHE40/ID2 | 2 | BP |
| GO:0035025 | positive regulation of Rho protein signal transduction | 2/88 | 33/18723 | 0.010491295 | 0.044864496 | 0.031827714 | F2R/ADGRG1 | 2 | BP |
| GO:0050901 | leukocyte tethering or rolling | 2/88 | 33/18723 | 0.010491295 | 0.044864496 | 0.031827714 | SELL/VCAM1 | 2 | BP |
| GO:0071353 | cellular response to interleukin-4 | 2/88 | 33/18723 | 0.010491295 | 0.044864496 | 0.031827714 | LEF1/TCF7 | 2 | BP |
| GO:0032640 | tumor necrosis factor production | 4/88 | 181/18723 | 0.010493081 | 0.044864496 | 0.031827714 | PTPN22/IFNG/CCL3/HAVCR2 | 4 | BP |
| GO:0032680 | regulation of tumor necrosis factor production | 4/88 | 181/18723 | 0.010493081 | 0.044864496 | 0.031827714 | PTPN22/IFNG/CCL3/HAVCR2 | 4 | BP |
| GO:0097696 | receptor signaling pathway via STAT | 4/88 | 181/18723 | 0.010493081 | 0.044864496 | 0.031827714 | IL7R/F2R/IFNG/CCL5 | 4 | BP |
| GO:0048017 | inositol lipid-mediated signaling | 4/88 | 182/18723 | 0.010691177 | 0.045504169 | 0.032281509 | CD28/PIK3IP1/F2R/CCL5 | 4 | BP |
| GO:0060401 | cytosolic calcium ion transport | 4/88 | 182/18723 | 0.010691177 | 0.045504169 | 0.032281509 | CCR7/F2R/FASLG/CCL3 | 4 | BP |
| GO:0048659 | smooth muscle cell proliferation | 4/88 | 184/18723 | 0.0110946 | 0.047086989 | 0.033404392 | LDLRAP1/ID2/IFNG/CCL5 | 4 | BP |
| GO:0009649 | entrainment of circadian clock | 2/88 | 34/18723 | 0.011113232 | 0.047086989 | 0.033404392 | BHLHE40/ID2 | 2 | BP |
| GO:0071706 | tumor necrosis factor superfamily cytokine production | 4/88 | 186/18723 | 0.011507738 | 0.048539379 | 0.034434744 | PTPN22/IFNG/CCL3/HAVCR2 | 4 | BP |
| GO:1903555 | regulation of tumor necrosis factor superfamily cytokine production | 4/88 | 186/18723 | 0.011507738 | 0.048539379 | 0.034434744 | PTPN22/IFNG/CCL3/HAVCR2 | 4 | BP |
| GO:0009119 | ribonucleoside metabolic process | 2/88 | 35/18723 | 0.011751067 | 0.049233822 | 0.034927395 | APOBEC3C/APOBEC3G | 2 | BP |
| GO:0039694 | viral RNA genome replication | 2/88 | 35/18723 | 0.011751067 | 0.049233822 | 0.034927395 | APOBEC3C/APOBEC3G | 2 | BP |
| GO:1903514 | release of sequestered calcium ion into cytosol by endoplasmic reticulum | 2/88 | 35/18723 | 0.011751067 | 0.049233822 | 0.034927395 | FASLG/CCL3 | 2 | BP |

**Supplementary Table 10: GO Enrichment Analysis Results for Highly Weighted Genes Associated with Trajectory 2 of CRC (Benjamini–Hochberg-adjusted P value < 0.05)**

| ID | Description | GeneRatio | BgRatio | pvalue | p.adjust | qvalue | geneID | Count | ONTOLOGY |
|---|---|---|---|---|---|---|---|---|---|
| GO:0022409 | positive regulation of cell-cell adhesion | 16/101 | 284/18723 | 2.59232E-12 | 5.16908E-09 | 3.79297E-09 | CCL5/CD160/ANXA1/TFRC/HLA-DQA1/TNFSF4/IL6ST/CD28/VCAM1/SIRPG/HLA-DRA/ADAM19/ICOS/CXCL13/HAVCR2/CD27 | 16 | BP |
| GO:0050870 | positive regulation of T cell activation | 14/101 | 216/18723 | 1.02724E-11 | 1.02416E-08 | 7.51506E-09 | CCL5/CD160/ANXA1/TFRC/HLA-DQA1/TNFSF4/IL6ST/CD28/VCAM1/SIRPG/HLA-DRA/ICOS/HAVCR2/CD27 | 14 | BP |
| GO:0002366 | leukocyte activation involved in immune response | 15/101 | 275/18723 | 2.09776E-11 | 1.28348E-08 | 9.41796E-09 | CX3CR1/FGR/KLRC2/CD244/ANXA1/PTGDR/CD300A/TFRC/LGALS3/TNFSF4/BATF/ITM2A/CD28/HLA-DRA/HAVCR2 | 15 | BP |
| GO:0002263 | cell activation involved in immune response | 15/101 | 279/18723 | 2.57469E-11 | 1.28348E-08 | 9.41796E-09 | CX3CR1/FGR/KLRC2/CD244/ANXA1/PTGDR/CD300A/TFRC/LGALS3/TNFSF4/BATF/ITM2A/CD28/HLA-DRA/HAVCR2 | 15 | BP |
| GO:0050777 | negative regulation of immune response | 13/101 | 194/18723 | 3.89688E-11 | 1.32764E-08 | 9.74197E-09 | KLRD1/CD160/FGR/A2M/LYAR/ANXA1/CD300A/LGALS3/TNFSF4/SAMSN1/PDCD1/CTLA4/HAVCR2 | 13 | BP |
| GO:1903039 | positive regulation of leukocyte cell-cell adhesion | 14/101 | 239/18723 | 3.99491E-11 | 1.32764E-08 | 9.74197E-09 | CCL5/CD160/ANXA1/TFRC/HLA-DQA1/TNFSF4/IL6ST/CD28/VCAM1/SIRPG/HLA-DRA/ICOS/HAVCR2/CD27 | 14 | BP |
| GO:0042129 | regulation of T cell proliferation | 11/101 | 171/18723 | 2.08564E-09 | 5.9411E-07 | 4.35947E-07 | CCL5/ANXA1/TFRC/PELI1/LGALS3/TNFSF4/IL6ST/CD28/VCAM1/CTLA4/HAVCR2 | 11 | BP |
| GO:0050670 | regulation of lymphocyte proliferation | 12/101 | 225/18723 | 3.14689E-09 | 7.7056E-07 | 5.65422E-07 | CCL5/ANXA1/CD300A/TFRC/PELI1/LGALS3/TNFSF4/IL6ST/CD28/VCAM1/CTLA4/HAVCR2 | 12 | BP |
| GO:0032944 | regulation of mononuclear cell proliferation | 12/101 | 227/18723 | 3.47795E-09 | 7.7056E-07 | 5.65422E-07 | CCL5/ANXA1/CD300A/TFRC/PELI1/LGALS3/TNFSF4/IL6ST/CD28/VCAM1/CTLA4/HAVCR2 | 12 | BP |
| GO:0002695 | negative regulation of leukocyte activation | 11/101 | 187/18723 | 5.34802E-09 | 1.0664E-06 | 7.825E-07 | FGR/CST7/ANXA1/CD300A/PELI1/PAG1/LGALS3/TNFSF4/SAMSN1/CTLA4/HAVCR2 | 11 | BP |
| GO:0002285 | lymphocyte activation involved in immune response | 11/101 | 194/18723 | 7.85358E-09 | 1.36326E-06 | 1.00033E-06 | KLRC2/CD244/ANXA1/TFRC/LGALS3/TNFSF4/BATF/ITM2A/CD28/HLA-DRA/HAVCR2 | 11 | BP |
| GO:0070663 | regulation of leukocyte proliferation | 12/101 | 245/18723 | 8.20415E-09 | 1.36326E-06 | 1.00033E-06 | CCL5/ANXA1/CD300A/TFRC/PELI1/LGALS3/TNFSF4/IL6ST/CD28/VCAM1/CTLA4/HAVCR2 | 12 | BP |
| GO:0042098 | T cell proliferation | 11/101 | 199/18723 | 1.0238E-08 | 1.57036E-06 | 1.1523E-06 | CCL5/ANXA1/TFRC/PELI1/LGALS3/TNFSF4/IL6ST/CD28/VCAM1/CTLA4/HAVCR2 | 11 | BP |
| GO:0051250 | negative regulation of lymphocyte activation | 10/101 | 157/18723 | 1.28744E-08 | 1.83368E-06 | 1.34552E-06 | FGR/ANXA1/CD300A/PELI1/PAG1/LGALS3/TNFSF4/SAMSN1/CTLA4/HAVCR2 | 10 | BP |
| GO:0050866 | negative regulation of cell activation | 11/101 | 210/18723 | 1.78888E-08 | 2.37802E-06 | 1.74495E-06 | FGR/CST7/ANXA1/CD300A/PELI1/PAG1/LGALS3/TNFSF4/SAMSN1/CTLA4/HAVCR2 | 11 | BP |
| GO:0001909 | leukocyte mediated cytotoxicity | 9/101 | 124/18723 | 2.28815E-08 | 2.85161E-06 | 2.09245E-06 | KLRD1/SLAMF7/CD160/CX3CR1/KLRC2/PRF1/LYST/HLA-DRA/HAVCR2 | 9 | BP |
| GO:0002703 | regulation of leukocyte mediated immunity | 11/101 | 226/18723 | 3.80959E-08 | 4.46842E-06 | 3.27884E-06 | KLRD1/CD160/CX3CR1/FGR/KLRC2/CD300A/TFRC/TNFSF4/CD28/HLA-DRA/HAVCR2 | 11 | BP |
| GO:0046651 | lymphocyte proliferation | 12/101 | 288/18723 | 4.92182E-08 | 5.03509E-06 | 3.69465E-06 | CCL5/ANXA1/CD300A/TFRC/PELI1/LGALS3/TNFSF4/IL6ST/CD28/VCAM1/CTLA4/HAVCR2 | 12 | BP |
| GO:0050671 | positive regulation of lymphocyte proliferation | 9/101 | 137/18723 | 5.45314E-08 | 5.03509E-06 | 3.69465E-06 | CCL5/ANXA1/TFRC/PELI1/TNFSF4/IL6ST/CD28/VCAM1/HAVCR2 | 9 | BP |
| GO:0032943 | mononuclear cell proliferation | 12/101 | 291/18723 | 5.5134E-08 | 5.03509E-06 | 3.69465E-06 | CCL5/ANXA1/CD300A/TFRC/PELI1/LGALS3/TNFSF4/IL6ST/CD28/VCAM1/CTLA4/HAVCR2 | 12 | BP |
| GO:0002819 | regulation of adaptive immune response | 10/101 | 183/18723 | 5.54002E-08 | 5.03509E-06 | 3.69465E-06 | KLRD1/CD160/ANXA1/TFRC/TNFSF4/IL6ST/CD28/HLA-DRA/SAMSN1/HAVCR2 | 10 | BP |
| GO:0002699 | positive regulation of immune effector process | 11/101 | 235/18723 | 5.67905E-08 | 5.03509E-06 | 3.69465E-06 | KLRD1/CD160/FGR/KLRC2/CD244/ANXA1/CD300A/TFRC/TNFSF4/CD28/HLA-DRA | 11 | BP |
| GO:0032946 | positive regulation of mononuclear cell proliferation | 9/101 | 138/18723 | 5.80778E-08 | 5.03509E-06 | 3.69465E-06 | CCL5/ANXA1/TFRC/PELI1/TNFSF4/IL6ST/CD28/VCAM1/HAVCR2 | 9 | BP |
| GO:0042102 | positive regulation of T cell proliferation | 8/101 | 101/18723 | 7.2369E-08 | 6.01266E-06 | 4.41197E-06 | CCL5/ANXA1/TFRC/TNFSF4/IL6ST/CD28/VCAM1/HAVCR2 | 8 | BP |
| GO:0022408 | negative regulation of cell-cell adhesion | 10/101 | 196/18723 | 1.05641E-07 | 8.42596E-06 | 6.18281E-06 | MBP/ANXA1/CD300A/PELI1/PAG1/LGALS3/TNFSF4/RGCC/CTLA4/HAVCR2 | 10 | BP |
| GO:0070665 | positive regulation of leukocyte proliferation | 9/101 | 150/18723 | 1.19191E-07 | 9.14104E-06 | 6.70752E-06 | CCL5/ANXA1/TFRC/PELI1/TNFSF4/IL6ST/CD28/VCAM1/HAVCR2 | 9 | BP |
| GO:0002698 | negative regulation of immune effector process | 8/101 | 110/18723 | 1.40988E-07 | 1.04122E-05 | 7.64027E-06 | KLRD1/CX3CR1/A2M/ANXA1/CD300A/LGALS3/TNFSF4/HAVCR2 | 8 | BP |
| GO:0050868 | negative regulation of T cell activation | 8/101 | 122/18723 | 3.14225E-07 | 2.23773E-05 | 1.642E-05 | ANXA1/CD300A/PELI1/PAG1/LGALS3/TNFSF4/CTLA4/HAVCR2 | 8 | BP |
| GO:0002274 | myeloid leukocyte activation | 10/101 | 223/18723 | 3.50532E-07 | 2.41021E-05 | 1.76856E-05 | CCL5/CX3CR1/FGR/CST7/PTGDR/CD300A/NAMPT/BATF/RBPJ/HAVCR2 | 10 | BP |
| GO:0006968 | cellular defense response | 6/101 | 54/18723 | 4.44228E-07 | 2.95264E-05 | 2.16659E-05 | CX3CR1/KLRG1/KLRC3/KLRC2/PRF1/TRAT1 | 6 | BP |
| GO:0045066 | regulatory T cell differentiation | 5/101 | 31/18723 | 6.28368E-07 | 4.04183E-05 | 2.96581E-05 | TOX/TNFSF4/CD28/HLA-DRA/CTLA4 | 5 | BP |
| GO:0001906 | cell killing | 9/101 | 188/18723 | 8.08106E-07 | 5.03551E-05 | 3.69496E-05 | KLRD1/SLAMF7/CD160/CX3CR1/KLRC2/PRF1/LYST/HLA-DRA/HAVCR2 | 9 | BP |
| GO:1903038 | negative regulation of leukocyte cell-cell adhesion | 8/101 | 141/18723 | 9.4973E-07 | 5.73867E-05 | 4.21093E-05 | ANXA1/CD300A/PELI1/PAG1/LGALS3/TNFSF4/CTLA4/HAVCR2 | 8 | BP |
| GO:0070232 | regulation of T cell apoptotic process | 5/101 | 34/18723 | 1.01594E-06 | 5.9582E-05 | 4.37201E-05 | CCL5/HIF1A/LGALS3/PDCD1/CD27 | 5 | BP |
| GO:0031341 | regulation of cell killing | 7/101 | 99/18723 | 1.06866E-06 | 6.0883E-05 | 4.46748E-05 | KLRD1/CD160/CX3CR1/KLRC2/PRF1/HLA-DRA/HAVCR2 | 7 | BP |
| GO:0042267 | natural killer cell mediated cytotoxicity | 6/101 | 68/18723 | 1.7714E-06 | 9.8116E-05 | 7.19956E-05 | KLRD1/SLAMF7/CD160/KLRC2/LYST/HAVCR2 | 6 | BP |
| GO:0046631 | alpha-beta T cell activation | 8/101 | 156/18723 | 2.03532E-06 | 0.000109687 | 8.04865E-05 | CD160/ANXA1/CD300A/TOX/TNFSF4/BATF/CD28/HLA-DRA | 8 | BP |
| GO:0002381 | immunoglobulin production involved in immunoglobulin-mediated immune response | 6/101 | 70/18723 | 2.10362E-06 | 0.000110385 | 8.09982E-05 | TFRC/HLA-DQA1/TNFSF4/BATF/CD28/HLA-DRA | 6 | BP |
| GO:0002228 | natural killer cell mediated immunity | 6/101 | 71/18723 | 2.28784E-06 | 0.000116973 | 8.58325E-05 | KLRD1/SLAMF7/CD160/KLRC2/LYST/HAVCR2 | 6 | BP |
| GO:0002708 | positive regulation of lymphocyte mediated immunity | 7/101 | 113/18723 | 2.60697E-06 | 0.000129958 | 9.53603E-05 | KLRD1/CD160/KLRC2/TFRC/TNFSF4/CD28/HLA-DRA | 7 | BP |
| GO:0045088 | regulation of innate immune response | 9/101 | 218/18723 | 2.75524E-06 | 0.000133999 | 9.83258E-05 | KLRD1/CCL5/CD160/FGR/A2M/KLRC2/LYAR/BIRC3/HAVCR2 | 9 | BP |
| GO:0002706 | regulation of lymphocyte mediated immunity | 8/101 | 168/18723 | 3.53856E-06 | 0.000167997 | 0.000123273 | KLRD1/CD160/KLRC2/TFRC/TNFSF4/CD28/HLA-DRA/HAVCR2 | 8 | BP |

| | | | | | | | | | |
|---|---|---|---|---|---|---|---|---|---|
| GO:2000106 | regulation of leukocyte apoptotic process | 6/101 | 81/18723 | 4.96387E-06 | 0.000230185 | 0.000168905 | CCL5/ANXA1/HIF1A/LGALS3/PDCD1/CD27 | 6 | BP |
| GO:0001910 | regulation of leukocyte mediated cytotoxicity | 6/101 | 82/18723 | 5.33254E-06 | 0.000241661 | 0.000177326 | KLRD1/CD160/CX3CR1/KLRC2/HLA-DRA/HAVCR2 | 6 | BP |
| GO:0070231 | T cell apoptotic process | 5/101 | 50/18723 | 7.22487E-06 | 0.000320142 | 0.000234914 | CCL5/HIF1A/LGALS3/PDCD1/CD27 | 5 | BP |
| GO:0030101 | natural killer cell activation | 6/101 | 88/18723 | 8.04065E-06 | 0.000335641 | 0.000246287 | SLAMF7/FGR/KLRC2/CD244/TOX/HAVCR2 | 6 | BP |
| GO:0070098 | chemokine-mediated signaling pathway | 6/101 | 88/18723 | 8.04065E-06 | 0.000335641 | 0.000246287 | CCL5/CX3CR1/XCL2/HIF1A/CXCR6/CXCL13 | 6 | BP |
| GO:0002705 | positive regulation of leukocyte mediated immunity | 7/101 | 134/18723 | 8.07962E-06 | 0.000335641 | 0.000246287 | KLRD1/CD160/KLRC2/TFRC/TNFSF4/CD28/HLA-DRA | 7 | BP |
| GO:0043547 | positive regulation of GTPase activity | 9/101 | 255/18723 | 9.82509E-06 | 0.000399821 | 0.000293381 | CCL5/XCL2/S1PR1/GPR65/SNX9/TIAM1/RGS1/TBC1D4/CXCL13 | 9 | BP |
| GO:0030217 | T cell differentiation | 9/101 | 257/18723 | 1.04595E-05 | 0.000414498 | 0.00030415 | CD8A/ANXA1/TOX/TNFSF4/BATF/CD28/HLA-DRA/CTLA4/CD27 | 9 | BP |
| GO:0070228 | regulation of lymphocyte apoptotic process | 5/101 | 54/18723 | 1.06015E-05 | 0.000414498 | 0.00030415 | CCL5/HIF1A/LGALS3/PDCD1/CD27 | 5 | BP |
| GO:0071674 | mononuclear cell migration | 8/101 | 196/18723 | 1.0998E-05 | 0.000421732 | 0.000309459 | CCL5/CX3CR1/GPR15/ANXA1/XCL2/S1PR1/LGALS3/CXCL13 | 8 | BP |
| GO:0050864 | regulation of B cell activation | 8/101 | 198/18723 | 1.18412E-05 | 0.000445496 | 0.000326896 | CD300A/TFRC/PELI1/TNFSF4/CD28/SAMSN1/CTLA4/CD27 | 8 | BP |
| GO:1990868 | response to chemokine | 6/101 | 97/18723 | 1.40997E-05 | 0.000504159 | 0.000369942 | CCL5/CX3CR1/XCL2/HIF1A/CXCR6/CXCL13 | 6 | BP |
| GO:1990869 | cellular response to chemokine | 6/101 | 97/18723 | 1.40997E-05 | 0.000504159 | 0.000369942 | CCL5/CX3CR1/XCL2/HIF1A/CXCR6/CXCL13 | 6 | BP |
| GO:0045580 | regulation of T cell differentiation | 7/101 | 146/18723 | 1.41589E-05 | 0.000504159 | 0.000369942 | ANXA1/TOX/TNFSF4/CD28/HLA-DRA/CTLA4/CD27 | 7 | BP |
| GO:0045589 | regulation of regulatory T cell differentiation | 4/101 | 28/18723 | 1.47846E-05 | 0.000517202 | 0.000379513 | TNFSF4/CD28/HLA-DRA/CTLA4 | 4 | BP |
| GO:0002820 | negative regulation of adaptive immune response | 5/101 | 59/18723 | 1.64285E-05 | 0.0005648 | 0.000414439 | KLRD1/CD160/TNFSF4/SAMSN1/HAVCR2 | 5 | BP |
| GO:0002685 | regulation of leukocyte migration | 8/101 | 210/18723 | 1.81305E-05 | 0.000612751 | 0.000449625 | CCL5/CX3CR1/ANXA1/XCL2/CD300A/LGALS3/TNFRSF18/CXCL13 | 8 | BP |
| GO:0035710 | CD4-positive, alpha-beta T cell activation | 6/101 | 102/18723 | 1.88007E-05 | 0.000624809 | 0.000458473 | CD160/ANXA1/TOX/TNFSF4/BATF/HLA-DRA | 6 | BP |
| GO:0061384 | heart trabecula morphogenesis | 4/101 | 30/18723 | 1.96254E-05 | 0.000641527 | 0.00047074 | TGFBR3/S1PR1/FKBP1A/RBPJ | 4 | BP |
| GO:0046634 | regulation of alpha-beta T cell activation | 6/101 | 104/18723 | 2.10028E-05 | 0.000675477 | 0.000495652 | CD160/ANXA1/CD300A/TNFSF4/CD28/HLA-DRA | 6 | BP |
| GO:0031343 | positive regulation of cell killing | 5/101 | 63/18723 | 2.26751E-05 | 0.000717686 | 0.000526624 | KLRD1/CD160/KLRC2/PRF1/HLA-DRA | 5 | BP |
| GO:0071887 | leukocyte apoptotic process | 6/101 | 106/18723 | 2.34082E-05 | 0.000729312 | 0.000535155 | CCL5/ANXA1/HIF1A/LGALS3/PDCD1/CD27 | 6 | BP |
| GO:0071677 | positive regulation of mononuclear cell migration | 5/101 | 65/18723 | 2.64203E-05 | 0.000810493 | 0.000594724 | CCL5/CX3CR1/XCL2/LGALS3/CXCL13 | 5 | BP |
| GO:0002429 | immune response-activating cell surface receptor signaling pathway | 9/101 | 291/18723 | 2.79873E-05 | 0.000832935 | 0.000611191 | KLRD1/CD160/FGR/KLRC2/CD300A/TRAT1/LGALS3/CD28/CTLA4 | 9 | BP |
| GO:0002757 | immune response-activating signal transduction | 9/101 | 291/18723 | 2.79873E-05 | 0.000832935 | 0.000611191 | KLRD1/CD160/FGR/KLRC2/CD300A/TRAT1/LGALS3/CD28/CTLA4 | 9 | BP |
| GO:0072678 | T cell migration | 5/101 | 66/18723 | 2.84643E-05 | 0.000834673 | 0.000612467 | CCL5/GPR15/XCL2/S1PR1/CXCL13 | 5 | BP |
| GO:0046635 | positive regulation of alpha-beta T cell activation | 5/101 | 67/18723 | 3.0629E-05 | 0.000885133 | 0.000649493 | CD160/ANXA1/TNFSF4/CD28/HLA-DRA | 5 | BP |
| GO:0030595 | leukocyte chemotaxis | 8/101 | 230/18723 | 3.47765E-05 | 0.000978918 | 0.000718311 | CCL5/CX3CR1/ANXA1/XCL2/S1PR1/LGALS3/LYST/CXCL13 | 8 | BP |
| GO:0002822 | regulation of adaptive immune response based on somatic recombination of immune receptors built from immunoglobulin superfamily domains | 7/101 | 168/18723 | 3.50134E-05 | 0.000978918 | 0.000718311 | KLRD1/ANXA1/TFRC/TNFSF4/CD28/HLA-DRA/HAVCR2 | 7 | BP |
| GO:0002286 | T cell activation involved in immune response | 6/101 | 114/18723 | 3.53471E-05 | 0.000978918 | 0.000718311 | ANXA1/LGALS3/TNFSF4/BATF/HLA-DRA/HAVCR2 | 6 | BP |
| GO:0032814 | regulation of natural killer cell activation | 4/101 | 35/18723 | 3.67284E-05 | 0.001003239 | 0.000736157 | FGR/KLRC2/TOX/HAVCR2 | 4 | BP |
| GO:0002548 | monocyte chemotaxis | 5/101 | 70/18723 | 3.78968E-05 | 0.001021166 | 0.000749311 | CCL5/CX3CR1/ANXA1/XCL2/LGALS3 | 5 | BP |
| GO:0070227 | lymphocyte apoptotic process | 5/101 | 72/18723 | 4.34385E-05 | 0.001149008 | 0.00084312 | CCL5/HIF1A/LGALS3/PDCD1/CD27 | 5 | BP |
| GO:0045619 | regulation of lymphocyte differentiation | 7/101 | 174/18723 | 4.37937E-05 | 0.001149008 | 0.00084312 | ANXA1/TOX/TNFSF4/CD28/HLA-DRA/CTLA4/CD27 | 7 | BP |
| GO:0001771 | immunological synapse formation | 3/101 | 14/18723 | 5.31164E-05 | 0.001375507 | 0.00100932 | PRF1/LGALS3/HAVCR2 | 3 | BP |
| GO:0050852 | T cell receptor signaling pathway | 6/101 | 123/18723 | 5.41656E-05 | 0.001384696 | 0.001016062 | CD160/CD300A/TRAT1/LGALS3/CD28/CTLA4 | 6 | BP |
| GO:0002347 | response to tumor cell | 4/101 | 39/18723 | 5.6749E-05 | 0.001414468 | 0.001037909 | CD160/PRF1/ABI3/HAVCR2 | 4 | BP |
| GO:2000516 | positive regulation of CD4-positive, alpha-beta T cell activation | 4/101 | 39/18723 | 5.6749E-05 | 0.001414468 | 0.001037909 | CD160/ANXA1/TNFSF4/HLA-DRA | 4 | BP |
| GO:0048644 | muscle organ morphogenesis | 5/101 | 77/18723 | 6.00481E-05 | 0.001478222 | 0.00108469 | TGFBR3/S1PR1/FKBP1A/ARID5B/RBPJ | 5 | BP |
| GO:0010820 | positive regulation of T cell chemotaxis | 3/101 | 15/18723 | 6.61354E-05 | 0.001588843 | 0.001165862 | CCL5/XCL2/CXCL13 | 3 | BP |
| GO:0045064 | T-helper 2 cell differentiation | 3/101 | 15/18723 | 6.61354E-05 | 0.001588843 | 0.001165862 | ANXA1/TNFSF4/BATF | 3 | BP |
| GO:0050856 | regulation of T cell receptor signaling pathway | 4/101 | 41/18723 | 6.92957E-05 | 0.001644949 | 0.001207031 | CD160/CD300A/TRAT1/LGALS3 | 4 | BP |
| GO:0031348 | negative regulation of defense response | 8/101 | 258/18723 | 7.80946E-05 | 0.001832007 | 0.001344291 | KLRD1/AOAH/FGR/A2M/LYAR/CST7/ACP5/HAVCR2 | 8 | BP |
| GO:0010819 | regulation of T cell chemotaxis | 3/101 | 16/18723 | 8.10787E-05 | 0.001860593 | 0.001365266 | CCL5/XCL2/CXCL13 | 3 | BP |
| GO:0002312 | B cell activation involved in immune response | 5/101 | 82/18723 | 8.11793E-05 | 0.001860593 | 0.001365266 | TFRC/TNFSF4/BATF/ITM2A/CD28 | 5 | BP |
| GO:0061383 | trabecula morphogenesis | 4/101 | 43/18723 | 8.37502E-05 | 0.001897703 | 0.001392497 | TGFBR3/S1PR1/FKBP1A/RBPJ | 4 | BP |
| GO:0043367 | CD4-positive, alpha-beta T cell differentiation | 5/101 | 83/18723 | 8.60163E-05 | 0.001927151 | 0.001414105 | ANXA1/TOX/TNFSF4/BATF/HLA-DRA | 5 | BP |
| GO:0002687 | positive regulation of leukocyte migration | 6/101 | 135/18723 | 9.08991E-05 | 0.001988479 | 0.001459107 | CCL5/CX3CR1/XCL2/LGALS3/TNFRSF18/CXCL13 | 6 | BP |
| GO:0031295 | T cell costimulation | 4/101 | 44/18723 | 9.17453E-05 | 0.001988479 | 0.001459107 | CD160/TNFSF4/CD28/ICOS | 4 | BP |
| GO:0042269 | regulation of natural killer cell mediated cytotoxicity | 4/101 | 44/18723 | 9.17453E-05 | 0.001988479 | 0.001459107 | KLRD1/CD160/KLRC2/HAVCR2 | 4 | BP |
| GO:0090713 | immunological memory process | 3/101 | 17/18723 | 9.80672E-05 | 0.002102646 | 0.00154288 | CD160/TNFSF4/HLA-DRA | 3 | BP |
| GO:0031294 | lymphocyte costimulation | 4/101 | 46/18723 | 0.000109379 | 0.002320221 | 0.001702532 | CD160/TNFSF4/CD28/ICOS | 4 | BP |
| GO:0050921 | positive regulation of chemotaxis | 6/101 | 141/18723 | 0.00011553 | 0.00242492 | 0.001779359 | CCL5/CX3CR1/XCL2/S1PR1/TIAM1/CXCL13 | 6 | BP |
| GO:0002544 | chronic inflammatory response | 3/101 | 18/18723 | 0.00011722 | 0.002434759 | 0.001786578 | CCL5/VCAM1/CXCL13 | 3 | BP |
| GO:0002715 | regulation of natural killer cell mediated immunity | 4/101 | 48/18723 | 0.000129349 | 0.002627444 | 0.001927967 | KLRD1/CD160/KLRC2/HAVCR2 | 4 | BP |
| GO:0016064 | immunoglobulin mediated immune response | 7/101 | 207/18723 | 0.000130355 | 0.002627444 | 0.001927967 | TFRC/HLA-DQA1/TNFSF4/BATF/CD28/HLA-DRA/CD27 | 7 | BP |
| GO:0031349 | positive regulation of defense response | 8/101 | 278/18723 | 0.000130969 | 0.002627444 | 0.001927967 | KLRD1/CCL5/CD160/KLRC2/TNFSF4/IL6ST/CD28/HAVCR2 | 8 | BP |
| GO:0002275 | myeloid cell activation involved in immune response | 5/101 | 91/18723 | 0.000133162 | 0.002627444 | 0.001927967 | CX3CR1/FGR/PTGDR/CD300A/HAVCR2 | 5 | BP |
| GO:0070233 | negative regulation of T cell apoptotic process | 3/101 | 19/18723 | 0.000138654 | 0.002627444 | 0.001927967 | CCL5/HIF1A/CD27 | 3 | BP |
| GO:0140131 | positive regulation of lymphocyte chemotaxis | 3/101 | 19/18723 | 0.000138654 | 0.002627444 | 0.001927967 | CCL5/XCL2/CXCL13 | 3 | BP |
| GO:0007189 | adenylate cyclase-activating G protein-coupled receptor signaling pathway | 6/101 | 146/18723 | 0.000139869 | 0.002627444 | 0.001927967 | PTGER2/S1PR5/S1PR1/ADGRG1/GPR65/ADRB2 | 6 | BP |
| GO:0002204 | somatic recombination of immunoglobulin genes involved in immune response | 4/101 | 49/18723 | 0.000140266 | 0.002627444 | 0.001927967 | TFRC/TNFSF4/BATF/CD28 | 4 | BP |
| GO:0002208 | somatic diversification of immunoglobulins involved in immune response | 4/101 | 49/18723 | 0.000140266 | 0.002627444 | 0.001927967 | TFRC/TNFSF4/BATF/CD28 | 4 | BP |
| GO:0045190 | isotype switching | 4/101 | 49/18723 | 0.000140266 | 0.002627444 | 0.001927967 | TFRC/TNFSF4/BATF/CD28 | 4 | BP |
| GO:0007249 | I-kappaB kinase/NF-kappaB signaling | 8/101 | 281/18723 | 0.000140991 | 0.002627444 | 0.001927967 | CX3CR1/TFRC/PELI1/TNFSF10/FKBP1A/TANK/BIRC3/NDFIP2 | 8 | BP |
| GO:0019724 | B cell mediated immunity | 7/101 | 210/18723 | 0.000142498 | 0.002630933 | 0.001930527 | TFRC/HLA-DQA1/TNFSF4/BATF/CD28/HLA-DRA/CD27 | 7 | BP |
| GO:0097530 | granulocyte migration | 6/101 | 148/18723 | 0.000150672 | 0.00275633 | 0.00202254 | CCL5/ANXA1/XCL2/CD300A/LGALS3/CXCL13 | 6 | BP |

| | | | | | | | | | |
|---|---|---|---|---|---|---|---|---|---|
| GO:0097529 | myeloid leukocyte migration | 7/101 | 220/18723 | 0.000189775 | 0.003440098 | 0.002524276 | CCL5/CX3CR1/ANXA1/XCL2/CD300A/LGALS3/CXCL13 | 7 | BP |
| GO:2000179 | positive regulation of neural precursor cell proliferation | 4/101 | 54/18723 | 0.000205087 | 0.003684178 | 0.002703378 | CX3CR1/ADGRG1/TOX/HIF1A | 4 | BP |
| GO:0003229 | ventricular cardiac muscle tissue development | 4/101 | 55/18723 | 0.000220262 | 0.003921445 | 0.002877479 | HOPX/TGFBR3/FKBP1A/RBPJ | 4 | BP |
| GO:0001912 | positive regulation of leukocyte mediated cytotoxicity | 4/101 | 56/18723 | 0.000236228 | 0.004168488 | 0.003058754 | KLRD1/CD160/KLRC2/HLA-DRA | 4 | BP |
| GO:0045621 | positive regulation of lymphocyte differentiation | 5/101 | 104/18723 | 0.000249173 | 0.004288321 | 0.003146686 | ANXA1/TOX/TNFSF4/HLA-DRA/CD27 | 5 | BP |
| GO:0034695 | response to prostaglandin E | 3/101 | 23/18723 | 0.000249471 | 0.004288321 | 0.003146686 | PTGER2/TGFBR3/TNFSF4 | 3 | BP |
| GO:0072677 | eosinophil migration | 3/101 | 23/18723 | 0.000249471 | 0.004288321 | 0.003146686 | CCL5/CD300A/LGALS3 | 3 | BP |
| GO:0016447 | somatic recombination of immunoglobulin gene segments | 4/101 | 57/18723 | 0.000253011 | 0.004311999 | 0.00316406 | TFRC/TNFSF4/BATF/CD28 | 4 | BP |
| GO:0007188 | adenylate cyclase-modulating G protein-coupled receptor signaling pathway | 7/101 | 233/18723 | 0.00026942 | 0.004552736 | 0.003340708 | PTGER2/S1PR5/S1PR1/ADGRG1/GPR65/ADRB2/RGS1 | 7 | BP |
| GO:0001911 | negative regulation of leukocyte mediated cytotoxicity | 3/101 | 24/18723 | 0.000283995 | 0.004725725 | 0.003467644 | KLRD1/CX3CR1/HAVCR2 | 3 | BP |
| GO:0031532 | actin cytoskeleton reorganization | 5/101 | 107/18723 | 0.000284397 | 0.004725725 | 0.003467644 | PLEK/ANXA1/ABI3/S1PR1/GPR65 | 5 | BP |
| GO:0055008 | cardiac muscle tissue morphogenesis | 4/101 | 60/18723 | 0.000308498 | 0.005065022 | 0.003716613 | TGFBR3/S1PR1/FKBP1A/RBPJ | 4 | BP |
| GO:0002456 | T cell mediated immunity | 5/101 | 109/18723 | 0.000309896 | 0.005065022 | 0.003716613 | CD8A/KLRD1/PRF1/TNFSF4/HLA-DRA | 5 | BP |
| GO:0032753 | positive regulation of interleukin-4 production | 3/101 | 25/18723 | 0.000321461 | 0.005169297 | 0.003793129 | TNFSF4/CD28/HAVCR2 | 3 | BP |
| GO:1901623 | regulation of lymphocyte chemotaxis | 3/101 | 25/18723 | 0.000321461 | 0.005169297 | 0.003793129 | CCL5/XCL2/CXCL13 | 3 | BP |
| GO:0002821 | positive regulation of adaptive immune response | 5/101 | 112/18723 | 0.00035135 | 0.005560248 | 0.004080001 | TFRC/TNFSF4/IL6ST/CD28/HLA-DRA | 5 | BP |
| GO:0046632 | alpha-beta T cell differentiation | 5/101 | 112/18723 | 0.00035135 | 0.005560248 | 0.004080001 | ANXA1/TOX/TNFSF4/BATF/HLA-DRA | 5 | BP |
| GO:0002418 | immune response to tumor cell | 3/101 | 26/18723 | 0.000361971 | 0.005595113 | 0.004105584 | CD160/PRF1/HAVCR2 | 3 | BP |
| GO:0045954 | positive regulation of natural killer cell mediated cytotoxicity | 3/101 | 26/18723 | 0.000361971 | 0.005595113 | 0.004105584 | KLRD1/CD160/KLRC2 | 3 | BP |
| GO:2000108 | positive regulation of leukocyte apoptotic process | 3/101 | 26/18723 | 0.000361971 | 0.005595113 | 0.004105584 | CCL5/ANXA1/PDCD1 | 3 | BP |
| GO:0002704 | negative regulation of leukocyte mediated immunity | 4/101 | 63/18723 | 0.000372211 | 0.005624428 | 0.004127094 | KLRD1/CX3CR1/CD300A/HAVCR2 | 4 | BP |
| GO:0050854 | regulation of antigen receptor-mediated signaling pathway | 4/101 | 63/18723 | 0.000372211 | 0.005624428 | 0.004127094 | CD160/CD300A/TRAT1/LGALS3 | 4 | BP |
| GO:0048771 | tissue remodeling | 6/101 | 175/18723 | 0.000372329 | 0.005624428 | 0.004127094 | S1PR1/ADRB2/TFRC/HIF1A/ACP5/RBPJ | 6 | BP |
| GO:0030888 | regulation of B cell proliferation | 4/101 | 64/18723 | 0.000395392 | 0.005903789 | 0.004332084 | CD300A/TFRC/PELI1/CTLA4 | 4 | BP |
| GO:0071675 | regulation of mononuclear cell migration | 5/101 | 115/18723 | 0.00039687 | 0.005903789 | 0.004332084 | CCL5/CX3CR1/XCL2/LGALS3/CXCL13 | 5 | BP |
| GO:0043122 | regulation of I-kappaB kinase/NF-kappaB signaling | 7/101 | 249/18723 | 0.000402265 | 0.005903789 | 0.004332084 | TFRC/PELI1/TNFSF10/FKBP1A/TANK/BIRC3/NDFIP2 | 7 | BP |
| GO:0002825 | regulation of T-helper 1 type immune response | 3/101 | 27/18723 | 0.000405626 | 0.005903789 | 0.004332084 | ANXA1/TNFSF4/HAVCR2 | 3 | BP |
| GO:0045830 | positive regulation of isotype switching | 3/101 | 27/18723 | 0.000405626 | 0.005903789 | 0.004332084 | TFRC/TNFSF4/CD28 | 3 | BP |
| GO:0045453 | bone resorption | 4/101 | 65/18723 | 0.000419587 | 0.006062733 | 0.004448714 | S1PR1/ADRB2/TFRC/ACP5 | 4 | BP |
| GO:0002702 | positive regulation of production of molecular mediator of immune response | 5/101 | 117/18723 | 0.000429601 | 0.006118748 | 0.004489817 | CD160/CD244/TFRC/TNFSF4/CD28 | 5 | BP |
| GO:0072676 | lymphocyte migration | 5/101 | 117/18723 | 0.000429601 | 0.006118748 | 0.004489817 | CCL5/GPR15/XCL2/S1PR1/CXCL13 | 5 | BP |
| GO:0002562 | somatic diversification of immune receptors via germline recombination within a single locus | 4/101 | 66/18723 | 0.000444821 | 0.00620261 | 0.004551353 | TFRC/TNFSF4/BATF/CD28 | 4 | BP |
| GO:0016444 | somatic cell DNA recombination | 4/101 | 66/18723 | 0.000444821 | 0.00620261 | 0.004551353 | TFRC/TNFSF4/BATF/CD28 | 4 | BP |
| GO:0042093 | T-helper cell differentiation | 4/101 | 66/18723 | 0.000444821 | 0.00620261 | 0.004551353 | ANXA1/TNFSF4/BATF/HLA-DRA | 4 | BP |
| GO:0010818 | T cell chemotaxis | 3/101 | 28/18723 | 0.000452527 | 0.006223032 | 0.004566339 | CCL5/XCL2/CXCL13 | 3 | BP |
| GO:0031342 | negative regulation of cell killing | 3/101 | 28/18723 | 0.000452527 | 0.006223032 | 0.004566339 | KLRD1/CX3CR1/HAVCR2 | 3 | BP |
| GO:0016445 | somatic diversification of immunoglobulins | 4/101 | 67/18723 | 0.000471119 | 0.006390553 | 0.004689262 | TFRC/TNFSF4/BATF/CD28 | 4 | BP |
| GO:2000514 | regulation of CD4-positive, alpha-beta T cell activation | 4/101 | 67/18723 | 0.000471119 | 0.006390553 | 0.004689262 | CD160/ANXA1/TNFSF4/HLA-DRA | 4 | BP |
| GO:0002294 | CD4-positive, alpha-beta T cell differentiation involved in immune response | 4/101 | 68/18723 | 0.000498506 | 0.006683509 | 0.004904227 | ANXA1/TNFSF4/BATF/HLA-DRA | 4 | BP |
| GO:0001773 | myeloid dendritic cell activation | 3/101 | 29/18723 | 0.000502771 | 0.006683509 | 0.004904227 | BATF/RBPJ/HAVCR2 | 3 | BP |
| GO:2000406 | positive regulation of T cell migration | 3/101 | 29/18723 | 0.000502771 | 0.006683509 | 0.004904227 | CCL5/XCL2/CXCL13 | 3 | BP |
| GO:0043123 | positive regulation of I-kappaB kinase/NF-kappaB signaling | 6/101 | 186/18723 | 0.000514501 | 0.006794145 | 0.00498541 | TFRC/PELI1/TNFSF10/FKBP1A/BIRC3/NDFIP2 | 6 | BP |
| GO:0002287 | alpha-beta T cell activation involved in immune response | 4/101 | 69/18723 | 0.000527009 | 0.00686834 | 0.005039852 | ANXA1/TNFSF4/BATF/HLA-DRA | 4 | BP |
| GO:0002293 | alpha-beta T cell differentiation involved in immune response | 4/101 | 69/18723 | 0.000527009 | 0.00686834 | 0.005039852 | ANXA1/TNFSF4/BATF/HLA-DRA | 4 | BP |
| GO:0034694 | response to prostaglandin | 3/101 | 30/18723 | 0.000556454 | 0.007069837 | 0.005187707 | PTGER2/TGFBR3/TNFSF4 | 3 | BP |
| GO:0050858 | negative regulation of antigen receptor-mediated signaling pathway | 3/101 | 30/18723 | 0.000556454 | 0.007069837 | 0.005187707 | CD160/CD300A/LGALS3 | 3 | BP |
| GO:0070229 | negative regulation of lymphocyte apoptotic process | 3/101 | 30/18723 | 0.000556454 | 0.007069837 | 0.005187707 | CCL5/HIF1A/CD27 | 3 | BP |
| GO:0060415 | muscle tissue morphogenesis | 4/101 | 70/18723 | 0.000556652 | 0.007069837 | 0.005187707 | TGFBR3/S1PR1/FKBP1A/RBPJ | 4 | BP |
| GO:0071621 | granulocyte chemotaxis | 5/101 | 125/18723 | 0.000581282 | 0.007275767 | 0.005338814 | CCL5/ANXA1/XCL2/LGALS3/CXCL13 | 5 | BP |
| GO:0003208 | cardiac ventricle morphogenesis | 4/101 | 71/18723 | 0.000587462 | 0.007275767 | 0.005338814 | TGFBR3/HIF1A/FKBP1A/RBPJ | 4 | BP |
| GO:0032722 | positive regulation of chemokine production | 4/101 | 71/18723 | 0.000587462 | 0.007275767 | 0.005338814 | MBP/HIF1A/TNFSF4/HAVCR2 | 4 | BP |
| GO:0045824 | negative regulation of innate immune response | 4/101 | 71/18723 | 0.000587462 | 0.007275767 | 0.005338814 | KLRD1/A2M/LYAR/HAVCR2 | 4 | BP |
| GO:0002717 | positive regulation of natural killer cell mediated immunity | 3/101 | 31/18723 | 0.00061367 | 0.007553447 | 0.00554257 | KLRD1/CD160/KLRC2 | 3 | BP |
| GO:0032729 | positive regulation of interferon-gamma production | 4/101 | 72/18723 | 0.000619463 | 0.00757797 | 0.005560565 | CD160/CD244/TNFSF4/HAVCR2 | 4 | BP |
| GO:0043299 | leukocyte degranulation | 4/101 | 73/18723 | 0.000652682 | 0.007935658 | 0.00582303 | FGR/KLRC2/PTGDR/CD300A | 4 | BP |
| GO:0043372 | positive regulation of CD4-positive, alpha-beta T cell differentiation | 3/101 | 32/18723 | 0.000674511 | 0.008151367 | 0.005981312 | ANXA1/TNFSF4/HLA-DRA | 3 | BP |
| GO:0045089 | positive regulation of innate immune response | 5/101 | 131/18723 | 0.000719083 | 0.008631231 | 0.006333428 | KLRD1/CCL5/CD160/KLRC2/HAVCR2 | 5 | BP |
| GO:0002292 | T cell differentiation involved in immune response | 4/101 | 75/18723 | 0.000722876 | 0.008631231 | 0.006333428 | ANXA1/TNFSF4/BATF/HLA-DRA | 4 | BP |
| GO:0032633 | interleukin-4 production | 3/101 | 33/18723 | 0.000739068 | 0.00872013 | 0.00639866 | TNFSF4/CD28/HAVCR2 | 3 | BP |
| GO:0032673 | regulation of interleukin-4 production | 3/101 | 33/18723 | 0.000739068 | 0.00872013 | 0.00639866 | TNFSF4/CD28/HAVCR2 | 3 | BP |
| GO:1902105 | regulation of leukocyte differentiation | 7/101 | 279/18723 | 0.000789657 | 0.009262207 | 0.006796425 | ANXA1/TOX/TNFSF4/CD28/HLA-DRA/CTLA4/CD27 | 7 | BP |
| GO:0002200 | somatic diversification of immune receptors | 4/101 | 77/18723 | 0.000798252 | 0.00930827 | 0.006830225 | TFRC/TNFSF4/BATF/CD28 | 4 | BP |
| GO:0050869 | negative regulation of B cell activation | 3/101 | 34/18723 | 0.00080743 | 0.009360551 | 0.006868588 | CD300A/SAMSN1/CTLA4 | 3 | BP |
| GO:2000403 | positive regulation of lymphocyte migration | 3/101 | 35/18723 | 0.000879683 | 0.010139236 | 0.007439972 | CCL5/XCL2/CXCL13 | 3 | BP |
| GO:1903322 | positive regulation of protein modification by small protein conjugation or removal | 5/101 | 138/18723 | 0.000909204 | 0.010419266 | 0.007645452 | PELI1/FKBP1A/TANK/BIRC3/NDFIP2 | 5 | BP |
| GO:0042092 | type 2 immune response | 3/101 | 36/18723 | 0.000955913 | 0.010830064 | 0.007946887 | ANXA1/TNFSF4/BATF | 3 | BP |

| | | | | | | | | | |
|---|---|---|---|---|---|---|---|---|---|
| GO:0045191 | regulation of isotype switching | 3/101 | 36/18723 | 0.000955913 | 0.010830064 | 0.007946887 | TFRC/TNFSF4/CD28 | 3 | BP |
| GO:0050672 | negative regulation of lymphocyte proliferation | 4/101 | 83/18723 | 0.001057539 | 0.011913744 | 0.008742071 | CD300A/PELI1/CTLA4/HAVCR2 | 4 | BP |
| GO:0070555 | response to interleukin-1 | 5/101 | 143/18723 | 0.001066296 | 0.011944916 | 0.008764944 | CCL5/ANXA1/XCL2/HIF1A/TANK | 5 | BP |
| GO:0010921 | regulation of phosphatase activity | 4/101 | 84/18723 | 0.001105862 | 0.012250497 | 0.008989173 | PLEK/CD300A/FKBP1A/LGALS3 | 4 | BP |
| GO:0032945 | negative regulation of mononuclear cell proliferation | 4/101 | 84/18723 | 0.001105862 | 0.012250497 | 0.008989173 | CD300A/PELI1/CTLA4/HAVCR2 | 4 | BP |
| GO:0002377 | immunoglobulin production | 6/101 | 216/18723 | 0.001121443 | 0.012354464 | 0.009065462 | TFRC/HLA-DQA1/TNFSF4/BATF/CD28/HLA-DRA | 6 | BP |
| GO:0045622 | regulation of T-helper cell differentiation | 3/101 | 39/18723 | 0.001209296 | 0.013249102 | 0.00972193 | ANXA1/TNFSF4/HLA-DRA | 3 | BP |
| GO:2000177 | regulation of neural precursor cell proliferation | 4/101 | 87/18723 | 0.00126011 | 0.013369665 | 0.009810397 | CX3CR1/ADGRG1/TOX/HIF1A | 4 | BP |
| GO:0002291 | T cell activation via T cell receptor contact with antigen bound to MHC molecule on antigen presenting cell | 2/101 | 10/18723 | 0.00126053 | 0.013369665 | 0.009810397 | LGALS3/HAVCR2 | 2 | BP |
| GO:0002887 | negative regulation of myeloid leukocyte mediated immunity | 2/101 | 10/18723 | 0.00126053 | 0.013369665 | 0.009810397 | CX3CR1/CD300A | 2 | BP |
| GO:0040015 | negative regulation of multicellular organism growth | 2/101 | 10/18723 | 0.00126053 | 0.013369665 | 0.009810397 | PLAC8/ADRB2 | 2 | BP |
| GO:0045625 | regulation of T-helper 1 cell differentiation | 2/101 | 10/18723 | 0.00126053 | 0.013369665 | 0.009810397 | ANXA1/TNFSF4 | 2 | BP |
| GO:0070099 | regulation of chemokine-mediated signaling pathway | 2/101 | 10/18723 | 0.00126053 | 0.013369665 | 0.009810397 | CCL5/HIF1A | 2 | BP |
| GO:0002714 | positive regulation of B cell mediated immunity | 3/101 | 40/18723 | 0.001302256 | 0.013666832 | 0.010028452 | TFRC/TNFSF4/CD28 | 3 | BP |
| GO:0002891 | positive regulation of immunoglobulin mediated immune response | 3/101 | 40/18723 | 0.001302256 | 0.013666832 | 0.010028452 | TFRC/TNFSF4/CD28 | 3 | BP |
| GO:0050920 | regulation of chemotaxis | 6/101 | 223/18723 | 0.001320527 | 0.013786022 | 0.010115911 | CCL5/CX3CR1/XCL2/S1PR1/TIAM1/CXCL13 | 6 | BP |
| GO:0050871 | positive regulation of B cell activation | 5/101 | 152/18723 | 0.001398673 | 0.014525803 | 0.010658748 | TFRC/PELI1/TNFSF4/CD28/CD27 | 5 | BP |
| GO:0046849 | bone remodeling | 4/101 | 90/18723 | 0.001428818 | 0.014685897 | 0.010776222 | S1PR1/ADRB2/TFRC/ACP5 | 4 | BP |
| GO:0070664 | negative regulation of leukocyte proliferation | 4/101 | 90/18723 | 0.001428818 | 0.014685897 | 0.010776222 | CD300A/PELI1/CTLA4/HAVCR2 | 4 | BP |
| GO:0045582 | positive regulation of T cell differentiation | 4/101 | 91/18723 | 0.001488389 | 0.01521973 | 0.011167938 | ANXA1/TNFSF4/HLA-DRA/CD27 | 4 | BP |
| GO:2000404 | regulation of T cell migration | 3/101 | 42/18723 | 0.001501389 | 0.015230132 | 0.011175571 | CCL5/XCL2/CXCL13 | 3 | BP |
| GO:0071356 | cellular response to tumor necrosis factor | 6/101 | 229/18723 | 0.00151177 | 0.015230132 | 0.011175571 | CCL5/XCL2/TANK/BIRC3/VCAM1/TNFRSF18 | 6 | BP |
| GO:0002357 | defense response to tumor cell | 2/101 | 11/18723 | 0.001535234 | 0.015230132 | 0.011175571 | PRF1/ABI3 | 2 | BP |
| GO:0033625 | positive regulation of integrin activation | 2/101 | 11/18723 | 0.001535234 | 0.015230132 | 0.011175571 | PLEK/CXCL13 | 2 | BP |
| GO:0033632 | regulation of cell-cell adhesion mediated by integrin | 2/101 | 11/18723 | 0.001535234 | 0.015230132 | 0.011175571 | CCL5/CXCL13 | 2 | BP |
| GO:0045628 | regulation of T-helper 2 cell differentiation | 2/101 | 11/18723 | 0.001535234 | 0.015230132 | 0.011175571 | ANXA1/TNFSF4 | 2 | BP |
| GO:0042088 | T-helper 1 type immune response | 3/101 | 43/18723 | 0.001607712 | 0.015617181 | 0.011459579 | ANXA1/TNFSF4/HAVCR2 | 3 | BP |
| GO:0046636 | negative regulation of alpha-beta T cell activation | 3/101 | 43/18723 | 0.001607712 | 0.015617181 | 0.011459579 | ANXA1/CD300A/TNFSF4 | 3 | BP |
| GO:0001837 | epithelial to mesenchymal transition | 5/101 | 157/18723 | 0.00161341 | 0.015617181 | 0.011459579 | TGFBR3/HIF1A/RGCC/TIAM1/RBPJ | 5 | BP |
| GO:1902107 | positive regulation of leukocyte differentiation | 5/101 | 157/18723 | 0.00161341 | 0.015617181 | 0.011459579 | ANXA1/TOX/TNFSF4/HLA-DRA/CD27 | 5 | BP |
| GO:1903708 | positive regulation of hemopoiesis | 5/101 | 157/18723 | 0.00161341 | 0.015617181 | 0.011459579 | ANXA1/TOX/TNFSF4/HLA-DRA/CD27 | 5 | BP |
| GO:0002690 | positive regulation of leukocyte chemotaxis | 4/101 | 94/18723 | 0.001677464 | 0.016158755 | 0.011856976 | CCL5/CX3CR1/XCL2/CXCL13 | 4 | BP |
| GO:0050764 | regulation of phagocytosis | 4/101 | 95/18723 | 0.001744029 | 0.016719196 | 0.012268217 | FGR/LYAR/CD300A/SIRPG | 4 | BP |
| GO:1904646 | cellular response to amyloid-beta | 3/101 | 45/18723 | 0.001834233 | 0.017431448 | 0.012790853 | ADRB2/NAMPT/VCAM1 | 3 | BP |
| GO:0043380 | regulation of memory T cell differentiation | 2/101 | 12/18723 | 0.00183581 | 0.017431448 | 0.012790853 | TNFSF4/HLA-DRA | 2 | BP |
| GO:0050851 | antigen receptor-mediated signaling pathway | 6/101 | 240/18723 | 0.001916463 | 0.018044789 | 0.013240911 | CD160/CD300A/TRAT1/LGALS3/CD28/CTLA4 | 6 | BP |
| GO:0002700 | regulation of production of molecular mediator of immune response | 5/101 | 164/18723 | 0.001953551 | 0.018044789 | 0.013240911 | CD160/CD244/TFRC/TNFSF4/CD28 | 5 | BP |
| GO:0043300 | regulation of leukocyte degranulation | 3/101 | 46/18723 | 0.001954571 | 0.018044789 | 0.013240911 | FGR/KLRC2/CD300A | 3 | BP |
| GO:0055010 | ventricular cardiac muscle tissue morphogenesis | 3/101 | 46/18723 | 0.001954571 | 0.018044789 | 0.013240911 | TGFBR3/FKBP1A/RBPJ | 3 | BP |
| GO:2000107 | negative regulation of leukocyte apoptotic process | 3/101 | 46/18723 | 0.001954571 | 0.018044789 | 0.013240911 | CCL5/HIF1A/CD27 | 3 | BP |
| GO:0032642 | regulation of chemokine production | 4/101 | 98/18723 | 0.001954701 | 0.018044789 | 0.013240911 | MBP/HIF1A/TNFSF4/HAVCR2 | 4 | BP |
| GO:1903320 | regulation of protein modification by small protein conjugation or removal | 6/101 | 242/18723 | 0.001998094 | 0.018360362 | 0.013472472 | HIF1A/PELI1/FKBP1A/TANK/BIRC3/NDFIP2 | 6 | BP |
| GO:0002444 | myeloid leukocyte mediated immunity | 4/101 | 99/18723 | 0.00202867 | 0.018387124 | 0.013492109 | CX3CR1/FGR/PTGDR/CD300A | 4 | BP |
| GO:0032602 | chemokine production | 4/101 | 99/18723 | 0.00202867 | 0.018387124 | 0.013492109 | MBP/HIF1A/TNFSF4/HAVCR2 | 4 | BP |
| GO:0042100 | B cell proliferation | 4/101 | 99/18723 | 0.00202867 | 0.018387124 | 0.013492109 | CD300A/TFRC/PELI1/CTLA4 | 4 | BP |
| GO:0001774 | microglial cell activation | 3/101 | 47/18723 | 0.00207972 | 0.018596239 | 0.013645554 | CX3CR1/CST7/NAMPT | 3 | BP |
| GO:0045581 | negative regulation of T cell differentiation | 3/101 | 47/18723 | 0.00207972 | 0.018596239 | 0.013645554 | ANXA1/TNFSF4/CTLA4 | 3 | BP |
| GO:0045911 | positive regulation of DNA recombination | 3/101 | 47/18723 | 0.00207972 | 0.018596239 | 0.013645554 | TFRC/TNFSF4/CD28 | 3 | BP |
| GO:0034112 | positive regulation of homotypic cell-cell adhesion | 2/101 | 13/18723 | 0.002161975 | 0.01881291 | 0.013804543 | CCL5/IL6ST | 2 | BP |
| GO:0043379 | memory T cell differentiation | 2/101 | 13/18723 | 0.002161975 | 0.01881291 | 0.013804543 | TNFSF4/HLA-DRA | 2 | BP |
| GO:0060347 | heart trabecula formation | 2/101 | 13/18723 | 0.002161975 | 0.01881291 | 0.013804543 | TGFBR3/FKBP1A | 2 | BP |
| GO:0061418 | regulation of transcription from RNA polymerase II promoter in response to hypoxia | 2/101 | 13/18723 | 0.002161975 | 0.01881291 | 0.013804543 | HIF1A/RBPJ | 2 | BP |
| GO:0070234 | positive regulation of T cell apoptotic process | 2/101 | 13/18723 | 0.002161975 | 0.01881291 | 0.013804543 | CCL5/PDCD1 | 2 | BP |
| GO:1903977 | positive regulation of glial cell migration | 2/101 | 13/18723 | 0.002161975 | 0.01881291 | 0.013804543 | CX3CR1/TIAM1 | 2 | BP |
| GO:0002833 | positive regulation of response to biotic stimulus | 5/101 | 168/18723 | 0.002169995 | 0.01881291 | 0.013804543 | KLRD1/CCL5/CD160/KLRC2/HAVCR2 | 5 | BP |
| GO:0043303 | mast cell degranulation | 3/101 | 48/18723 | 0.002209745 | 0.019074591 | 0.013996559 | FGR/PTGDR/CD300A | 3 | BP |
| GO:0030593 | neutrophil chemotaxis | 4/101 | 103/18723 | 0.002343937 | 0.01998014 | 0.014661033 | CCL5/XCL2/LGALS3/CXCL13 | 4 | BP |
| GO:0001913 | T cell mediated cytotoxicity | 3/101 | 49/18723 | 0.002344711 | 0.01998014 | 0.014661033 | KLRD1/PRF1/HLA-DRA | 3 | BP |
| GO:0002279 | mast cell activation involved in immune response | 3/101 | 49/18723 | 0.002344711 | 0.01998014 | 0.014661033 | FGR/PTGDR/CD300A | 3 | BP |
| GO:0002448 | mast cell mediated immunity | 3/101 | 50/18723 | 0.002484681 | 0.020625651 | 0.015134696 | FGR/PTGDR/CD300A | 3 | BP |
| GO:0002639 | positive regulation of immunoglobulin production | 3/101 | 50/18723 | 0.002484681 | 0.020625651 | 0.015134696 | TFRC/TNFSF4/CD28 | 3 | BP |
| GO:0045058 | T cell selection | 3/101 | 50/18723 | 0.002484681 | 0.020625651 | 0.015134696 | TOX/BATF/CD28 | 3 | BP |
| GO:0046638 | positive regulation of alpha-beta T cell differentiation | 3/101 | 50/18723 | 0.002484681 | 0.020625651 | 0.015134696 | ANXA1/TNFSF4/HLA-DRA | 3 | BP |
| GO:0034612 | response to tumor necrosis factor | 6/101 | 253/18723 | 0.002495107 | 0.020625651 | 0.015134696 | CCL5/XCL2/TANK/BIRC3/VCAM1/TNFRSF18 | 6 | BP |
| GO:0003376 | sphingosine-1-phosphate receptor signaling pathway | 2/101 | 14/18723 | 0.002513452 | 0.020625651 | 0.015134696 | S1PR5/S1PR1 | 2 | BP |
| GO:0060576 | intestinal epithelial cell development | 2/101 | 14/18723 | 0.002513452 | 0.020625651 | 0.015134696 | HIF1A/IL6ST | 2 | BP |
| GO:0090715 | immunological memory formation process | 2/101 | 14/18723 | 0.002513452 | 0.020625651 | 0.015134696 | TNFSF4/HLA-DRA | 2 | BP |
| GO:0032231 | regulation of actin filament bundle assembly | 4/101 | 105/18723 | 0.002513557 | 0.020625651 | 0.015134696 | PLEK/S1PR1/GPR65/RGCC | 4 | BP |
| GO:0035821 | modulation of process of other organism | 4/101 | 106/18723 | 0.002601451 | 0.021172627 | 0.015536056 | CCL5/CX3CR1/STOM/PRF1 | 4 | BP |

| | | | | | | | | | |
|---|---|---|---|---|---|---|---|---|---|
| GO:0042116 | macrophage activation | 4/101 | 106/18723 | 0.002601451 | 0.021172627 | 0.015536056 | CX3CR1/CST7/NAMPT/HAVCR2 | 4 | BP |
| GO:0043370 | regulation of CD4-positive, alpha-beta T cell differentiation | 3/101 | 51/18723 | 0.002629719 | 0.02131569 | 0.015641033 | ANXA1/TNFSF4/HLA-DRA | 3 | BP |
| GO:0002824 | positive regulation of adaptive immune response based on somatic recombination of immune receptors built from immunoglobulin superfamily domains | 4/101 | 107/18723 | 0.002691434 | 0.021727608 | 0.01594329 | TFRC/TNFSF4/CD28/HLA-DRA | 4 | BP |
| GO:0002832 | negative regulation of response to biotic stimulus | 4/101 | 108/18723 | 0.00278353 | 0.022380478 | 0.016422353 | KLRD1/A2M/LYAR/HAVCR2 | 4 | BP |
| GO:0042976 | activation of Janus kinase activity | 2/101 | 15/18723 | 0.002889962 | 0.023050336 | 0.016913882 | CCL5/CD300A | 2 | BP |
| GO:0071380 | cellular response to prostaglandin E stimulus | 2/101 | 15/18723 | 0.002889962 | 0.023050336 | 0.016913882 | PTGER2/TNFSF4 | 2 | BP |
| GO:0008347 | glial cell migration | 3/101 | 53/18723 | 0.002935237 | 0.023133841 | 0.016975156 | CX3CR1/ADGRG1/TIAM1 | 3 | BP |
| GO:0033059 | cellular pigmentation | 3/101 | 53/18723 | 0.002935237 | 0.023133841 | 0.016975156 | ZEB2/MYO7A/LYST | 3 | BP |
| GO:0045744 | negative regulation of G protein-coupled receptor signaling pathway | 3/101 | 53/18723 | 0.002935237 | 0.023133841 | 0.016975156 | CCL5/PLEK/ADRB2 | 3 | BP |
| GO:0002823 | negative regulation of adaptive immune response based on somatic recombination of immune receptors built from immunoglobulin superfamily domains | 3/101 | 54/18723 | 0.003095834 | 0.024208205 | 0.017763503 | KLRD1/TNFSF4/HAVCR2 | 3 | BP |
| GO:0043647 | inositol phosphate metabolic process | 3/101 | 54/18723 | 0.003095834 | 0.024208205 | 0.017763503 | PLEK/CD244/INPP4B | 3 | BP |
| GO:0032609 | interferon-gamma production | 4/101 | 112/18723 | 0.003173533 | 0.024579454 | 0.018035919 | CD160/CD244/TNFSF4/HAVCR2 | 4 | BP |
| GO:0032649 | regulation of interferon-gamma production | 4/101 | 112/18723 | 0.003173533 | 0.024579454 | 0.018035919 | CD160/CD244/TNFSF4/HAVCR2 | 4 | BP |
| GO:0045620 | negative regulation of lymphocyte differentiation | 3/101 | 55/18723 | 0.003261732 | 0.024579454 | 0.018035919 | ANXA1/TNFSF4/CTLA4 | 3 | BP |
| GO:1905517 | macrophage migration | 3/101 | 55/18723 | 0.003261732 | 0.024579454 | 0.018035919 | CCL5/CX3CR1/LGALS3 | 3 | BP |
| GO:0071347 | cellular response to interleukin-1 | 4/101 | 113/18723 | 0.00327656 | 0.024579454 | 0.018035919 | CCL5/XCL2/HIF1A/TANK | 4 | BP |
| GO:0002399 | MHC class II protein complex assembly | 2/101 | 16/18723 | 0.003291231 | 0.024579454 | 0.018035919 | HLA-DQA1/HLA-DRA | 2 | BP |
| GO:0002503 | peptide antigen assembly with MHC class II protein complex | 2/101 | 16/18723 | 0.003291231 | 0.024579454 | 0.018035919 | HLA-DQA1/HLA-DRA | 2 | BP |
| GO:0010919 | regulation of inositol phosphate biosynthetic process | 2/101 | 16/18723 | 0.003291231 | 0.024579454 | 0.018035919 | PLEK/CD244 | 2 | BP |
| GO:0033631 | cell-cell adhesion mediated by integrin | 2/101 | 16/18723 | 0.003291231 | 0.024579454 | 0.018035919 | CCL5/CXCL13 | 2 | BP |
| GO:0034138 | toll-like receptor 3 signaling pathway | 2/101 | 16/18723 | 0.003291231 | 0.024579454 | 0.018035919 | PELI1/HAVCR2 | 2 | BP |
| GO:0070230 | positive regulation of lymphocyte apoptotic process | 2/101 | 16/18723 | 0.003291231 | 0.024579454 | 0.018035919 | CCL5/PDCD1 | 2 | BP |
| GO:0090520 | sphingolipid mediated signaling pathway | 2/101 | 16/18723 | 0.003291231 | 0.024579454 | 0.018035919 | S1PR5/S1PR1 | 2 | BP |
| GO:0002886 | regulation of myeloid leukocyte mediated immunity | 3/101 | 56/18723 | 0.003432987 | 0.025447494 | 0.018672869 | CX3CR1/FGR/CD300A | 3 | BP |
| GO:1904645 | response to amyloid-beta | 3/101 | 56/18723 | 0.003432987 | 0.025447494 | 0.018672869 | ADRB2/NAMPT/VCAM1 | 3 | BP |
| GO:0030278 | regulation of ossification | 4/101 | 115/18723 | 0.003489411 | 0.025674858 | 0.018839704 | S1PR1/ADRB2/HIF1A/RBPJ | 4 | BP |
| GO:0090630 | activation of GTPase activity | 4/101 | 115/18723 | 0.003489411 | 0.025674858 | 0.018839704 | GPR65/TIAM1/TBC1D4/CXCL13 | 4 | BP |
| GO:0048872 | homeostasis of number of cells | 6/101 | 272/18723 | 0.003565774 | 0.026140269 | 0.019181214 | CX3CR1/TGFBR3/LYAR/ANXA1/ADGRG1/HIF1A | 6 | BP |
| GO:0033623 | regulation of integrin activation | 2/101 | 17/18723 | 0.003716986 | 0.027148975 | 0.019921382 | PLEK/CXCL13 | 2 | BP |
| GO:0043666 | regulation of phosphoprotein phosphatase activity | 3/101 | 58/18723 | 0.00379178 | 0.027594198 | 0.020248079 | CD300A/FKBP1A/LGALS3 | 3 | BP |
| GO:0030282 | bone mineralization | 4/101 | 119/18723 | 0.003942967 | 0.02848651 | 0.020902839 | FGR/S1PR1/ADRB2/HIF1A | 4 | BP |
| GO:0031398 | positive regulation of protein ubiquitination | 4/101 | 119/18723 | 0.003942967 | 0.02848651 | 0.020902839 | PELI1/FKBP1A/BIRC3/NDFIP2 | 4 | BP |
| GO:0002223 | stimulatory C-type lectin receptor signaling pathway | 2/101 | 18/18723 | 0.004166957 | 0.028868455 | 0.021183104 | KLRD1/KLRC2 | 2 | BP |
| GO:0002501 | peptide antigen assembly with MHC protein complex | 2/101 | 18/18723 | 0.004166957 | 0.028868455 | 0.021183104 | HLA-DQA1/HLA-DRA | 2 | BP |
| GO:0030889 | negative regulation of B cell proliferation | 2/101 | 18/18723 | 0.004166957 | 0.028868455 | 0.021183104 | CD300A/CTLA4 | 2 | BP |
| GO:0045623 | negative regulation of T-helper cell differentiation | 2/101 | 18/18723 | 0.004166957 | 0.028868455 | 0.021183104 | ANXA1/TNFSF4 | 2 | BP |
| GO:0045953 | negative regulation of natural killer cell mediated cytotoxicity | 2/101 | 18/18723 | 0.004166957 | 0.028868455 | 0.021183104 | KLRD1/HAVCR2 | 2 | BP |
| GO:0060546 | negative regulation of necroptotic process | 2/101 | 18/18723 | 0.004166957 | 0.028868455 | 0.021183104 | PELI1/BIRC3 | 2 | BP |
| GO:1990840 | response to lectin | 2/101 | 18/18723 | 0.004166957 | 0.028868455 | 0.021183104 | KLRD1/KLRC2 | 2 | BP |
| GO:1990858 | cellular response to lectin | 2/101 | 18/18723 | 0.004166957 | 0.028868455 | 0.021183104 | KLRD1/KLRC2 | 2 | BP |
| GO:0002712 | regulation of B cell mediated immunity | 3/101 | 60/18723 | 0.004172629 | 0.028868455 | 0.021183104 | TFRC/TNFSF4/CD28 | 3 | BP |
| GO:0002889 | regulation of immunoglobulin mediated immune response | 3/101 | 60/18723 | 0.004172629 | 0.028868455 | 0.021183104 | TFRC/TNFSF4/CD28 | 3 | BP |
| GO:0051851 | modulation by host of symbiont process | 3/101 | 60/18723 | 0.004172629 | 0.028868455 | 0.021183104 | CCL5/CX3CR1/STOM | 3 | BP |
| GO:0002224 | toll-like receptor signaling pathway | 4/101 | 121/18723 | 0.004184044 | 0.028868455 | 0.021183104 | CD300A/PELI1/BIRC3/HAVCR2 | 4 | BP |
| GO:0003206 | cardiac chamber morphogenesis | 4/101 | 121/18723 | 0.004184044 | 0.028868455 | 0.021183104 | TGFBR3/HIF1A/FKBP1A/RBPJ | 4 | BP |
| GO:0043393 | regulation of protein binding | 5/101 | 196/18723 | 0.004204207 | 0.02890755 | 0.021211791 | HOPX/TGFBR3/ADRB2/FKBP1A/TIAM1 | 5 | BP |
| GO:0002688 | regulation of leukocyte chemotaxis | 4/101 | 122/18723 | 0.004308237 | 0.02941995 | 0.021587779 | CCL5/CX3CR1/XCL2/CXCL13 | 4 | BP |
| GO:1990266 | neutrophil migration | 4/101 | 122/18723 | 0.004308237 | 0.02941995 | 0.021587779 | CCL5/XCL2/LGALS3/CXCL13 | 4 | BP |
| GO:2000401 | regulation of lymphocyte migration | 3/101 | 61/18723 | 0.004371449 | 0.029749723 | 0.021829761 | CCL5/XCL2/CXCL13 | 3 | BP |
| GO:0003231 | cardiac ventricle development | 4/101 | 123/18723 | 0.004434898 | 0.030078865 | 0.022071278 | TGFBR3/HIF1A/FKBP1A/RBPJ | 4 | BP |
| GO:0010573 | vascular endothelial growth factor production | 3/101 | 62/18723 | 0.004575928 | 0.030340669 | 0.022263385 | ADGRG1/HIF1A/IL6ST | 3 | BP |
| GO:0032623 | interleukin-2 production | 3/101 | 62/18723 | 0.004575928 | 0.030340669 | 0.022263385 | ANXA1/CD28/HAVCR2 | 3 | BP |
| GO:0032663 | regulation of interleukin-2 production | 3/101 | 62/18723 | 0.004575928 | 0.030340669 | 0.022263385 | ANXA1/CD28/HAVCR2 | 3 | BP |
| GO:0045576 | mast cell activation | 3/101 | 62/18723 | 0.004575928 | 0.030340669 | 0.022263385 | FGR/PTGDR/CD300A | 3 | BP |
| GO:0002281 | macrophage activation involved in immune response | 2/101 | 19/18723 | 0.004640875 | 0.030340669 | 0.022263385 | CX3CR1/HAVCR2 | 2 | BP |
| GO:0002396 | MHC protein complex assembly | 2/101 | 19/18723 | 0.004640875 | 0.030340669 | 0.022263385 | HLA-DQA1/HLA-DRA | 2 | BP |
| GO:0002716 | negative regulation of natural killer cell mediated immunity | 2/101 | 19/18723 | 0.004640875 | 0.030340669 | 0.022263385 | KLRD1/HAVCR2 | 2 | BP |
| GO:0030050 | vesicle transport along actin filament | 2/101 | 19/18723 | 0.004640875 | 0.030340669 | 0.022263385 | MYO1F/MYO7A | 2 | BP |
| GO:0048245 | eosinophil chemotaxis | 2/101 | 19/18723 | 0.004640875 | 0.030340669 | 0.022263385 | CCL5/LGALS3 | 2 | BP |
| GO:0062099 | negative regulation of programmed necrotic cell death | 2/101 | 19/18723 | 0.004640875 | 0.030340669 | 0.022263385 | PELI1/BIRC3 | 2 | BP |
| GO:1903975 | regulation of glial cell migration | 2/101 | 19/18723 | 0.004640875 | 0.030340669 | 0.022263385 | CX3CR1/TIAM1 | 2 | BP |
| GO:0051054 | positive regulation of DNA metabolic process | 5/101 | 201/18723 | 0.004675255 | 0.030465549 | 0.022355019 | TFRC/TOX/TNFSF4/RGCC/CD28 | 5 | BP |
| GO:0019722 | calcium-mediated signaling | 5/101 | 202/18723 | 0.00477379 | 0.030985428 | 0.022736496 | PLEK/CX3CR1/TRAT1/VCAM1/CXCR6 | 5 | BP |
| GO:0032233 | positive regulation of actin filament bundle assembly | 3/101 | 63/18723 | 0.004786114 | 0.030985428 | 0.022736496 | PLEK/GPR65/RGCC | 3 | BP |
| GO:0048247 | lymphocyte chemotaxis | 3/101 | 64/18723 | 0.005002051 | 0.032220492 | 0.023642762 | CCL5/XCL2/CXCL13 | 3 | BP |
| GO:0032612 | interleukin-1 production | 4/101 | 128/18723 | 0.005105986 | 0.032220492 | 0.023642762 | CX3CR1/ANXA1/ACP5/HAVCR2 | 4 | BP |
| GO:0032652 | regulation of interleukin-1 production | 4/101 | 128/18723 | 0.005105986 | 0.032220492 | 0.023642762 | CX3CR1/ANXA1/ACP5/HAVCR2 | 4 | BP |
| GO:0035303 | regulation of dephosphorylation | 4/101 | 128/18723 | 0.005105986 | 0.032220492 | 0.023642762 | PLEK/CD300A/FKBP1A/LGALS3 | 4 | BP |

| | | | | | | | | | |
|---|---|---|---|---|---|---|---|---|---|
| GO:0002834 | regulation of response to tumor cell | 2/101 | 20/18723 | 0.005138474 | 0.032220492 | 0.023642762 | CD160/HAVCR2 | 2 | BP |
| GO:0002837 | regulation of immune response to tumor cell | 2/101 | 20/18723 | 0.005138474 | 0.032220492 | 0.023642762 | CD160/HAVCR2 | 2 | BP |
| GO:0043011 | myeloid dendritic cell differentiation | 2/101 | 20/18723 | 0.005138474 | 0.032220492 | 0.023642762 | BATF/RBPJ | 2 | BP |
| GO:0045063 | T-helper 1 cell differentiation | 2/101 | 20/18723 | 0.005138474 | 0.032220492 | 0.023642762 | ANXA1/TNFSF4 | 2 | BP |
| GO:0071379 | cellular response to prostaglandin stimulus | 2/101 | 20/18723 | 0.005138474 | 0.032220492 | 0.023642762 | PTGER2/TNFSF4 | 2 | BP |
| GO:0097320 | plasma membrane tubulation | 2/101 | 20/18723 | 0.005138474 | 0.032220492 | 0.023642762 | BIN2/SNX9 | 2 | BP |
| GO:0048524 | positive regulation of viral process | 3/101 | 65/18723 | 0.005223783 | 0.03265274 | 0.023959937 | CCL5/STOM/CD28 | 3 | BP |
| GO:0050918 | positive chemotaxis | 3/101 | 66/18723 | 0.005451353 | 0.033968744 | 0.024925595 | CCL5/S1PR1/LGALS3 | 3 | BP |
| GO:0071222 | cellular response to lipopolysaccharide | 5/101 | 209/18723 | 0.005505343 | 0.034198302 | 0.02509404 | CCL5/CX3CR1/TNFSF4/CXCL13/HAVCR2 | 5 | BP |
| GO:0019079 | viral genome replication | 4/101 | 131/18723 | 0.005539659 | 0.034304598 | 0.025172038 | CCL5/STOM/CD28/CXCR6 | 4 | BP |
| GO:0033630 | positive regulation of cell adhesion mediated by integrin | 2/101 | 21/18723 | 0.005659489 | 0.034405555 | 0.025246118 | CCL5/CXCL13 | 2 | BP |
| GO:0043373 | CD4-positive, alpha-beta T cell lineage commitment | 2/101 | 21/18723 | 0.005659489 | 0.034405555 | 0.025246118 | TOX/BATF | 2 | BP |
| GO:0046641 | positive regulation of alpha-beta T cell proliferation | 2/101 | 21/18723 | 0.005659489 | 0.034405555 | 0.025246118 | TNFSF4/CD28 | 2 | BP |
| GO:0090026 | positive regulation of monocyte chemotaxis | 2/101 | 21/18723 | 0.005659489 | 0.034405555 | 0.025246118 | CCL5/CX3CR1 | 2 | BP |
| GO:0099515 | actin filament-based transport | 2/101 | 21/18723 | 0.005659489 | 0.034405555 | 0.025246118 | MYO1F/MYO7A | 2 | BP |
| GO:2000047 | regulation of cell-cell adhesion mediated by cadherin | 2/101 | 21/18723 | 0.005659489 | 0.034405555 | 0.025246118 | RGCC/ADAM19 | 2 | BP |
| GO:0042130 | negative regulation of T cell proliferation | 3/101 | 67/18723 | 0.005684802 | 0.03445439 | 0.025281952 | PELI1/CTLA4/HAVCR2 | 3 | BP |
| GO:0009988 | cell-cell recognition | 3/101 | 68/18723 | 0.005924169 | 0.035580702 | 0.026108418 | PRF1/LGALS3/HAVCR2 | 3 | BP |
| GO:0042531 | positive regulation of tyrosine phosphorylation of STAT protein | 3/101 | 68/18723 | 0.005924169 | 0.035580702 | 0.026108418 | CCL5/IL6ST/TNFRSF18 | 3 | BP |
| GO:0046637 | regulation of alpha-beta T cell differentiation | 3/101 | 68/18723 | 0.005924169 | 0.035580702 | 0.026108418 | ANXA1/TNFSF4/HLA-DRA | 3 | BP |
| GO:0008277 | regulation of G protein-coupled receptor signaling pathway | 4/101 | 134/18723 | 0.005997306 | 0.035804278 | 0.026272473 | CCL5/PLEK/ADRB2/RGS1 | 4 | BP |
| GO:0032147 | activation of protein kinase activity | 4/101 | 134/18723 | 0.005997306 | 0.035804278 | 0.026272473 | CCL5/ADRB2/CD300A/RGCC | 4 | BP |
| GO:0002220 | innate immune response activating cell surface receptor signaling pathway | 2/101 | 22/18723 | 0.00620366 | 0.036169878 | 0.026540743 | KLRD1/KLRC2 | 2 | BP |
| GO:0032816 | positive regulation of natural killer cell activation | 2/101 | 22/18723 | 0.00620366 | 0.036169878 | 0.026540743 | KLRC2/TOX | 2 | BP |
| GO:0043371 | negative regulation of CD4-positive, alpha-beta T cell differentiation | 2/101 | 22/18723 | 0.00620366 | 0.036169878 | 0.026540743 | ANXA1/TNFSF4 | 2 | BP |
| GO:0045061 | thymic T cell selection | 2/101 | 22/18723 | 0.00620366 | 0.036169878 | 0.026540743 | TOX/CD28 | 2 | BP |
| GO:0045624 | positive regulation of T-helper cell differentiation | 2/101 | 22/18723 | 0.00620366 | 0.036169878 | 0.026540743 | ANXA1/TNFSF4 | 2 | BP |
| GO:0050860 | negative regulation of T cell receptor signaling pathway | 2/101 | 22/18723 | 0.00620366 | 0.036169878 | 0.026540743 | CD160/LGALS3 | 2 | BP |
| GO:0060343 | trabecula formation | 2/101 | 22/18723 | 0.00620366 | 0.036169878 | 0.026540743 | TGFBR3/FKBP1A | 2 | BP |
| GO:1901522 | positive regulation of transcription from RNA polymerase II promoter involved in cellular response to chemical stimulus | 2/101 | 22/18723 | 0.00620366 | 0.036169878 | 0.026540743 | HIF1A/RBPJ | 2 | BP |
| GO:0002052 | positive regulation of neuroblast proliferation | 2/101 | 23/18723 | 0.006770726 | 0.038684318 | 0.02838579 | CX3CR1/HIF1A | 2 | BP |
| GO:0002363 | alpha-beta T cell lineage commitment | 2/101 | 23/18723 | 0.006770726 | 0.038684318 | 0.02838579 | TOX/BATF | 2 | BP |
| GO:0002758 | innate immune response-activating signal transduction | 2/101 | 23/18723 | 0.006770726 | 0.038684318 | 0.02838579 | KLRD1/KLRC2 | 2 | BP |
| GO:0031338 | regulation of vesicle fusion | 2/101 | 23/18723 | 0.006770726 | 0.038684318 | 0.02838579 | ANXA1/TBC1D4 | 2 | BP |
| GO:0043369 | CD4-positive or CD8-positive, alpha-beta T cell lineage commitment | 2/101 | 23/18723 | 0.006770726 | 0.038684318 | 0.02838579 | TOX/BATF | 2 | BP |
| GO:0060547 | negative regulation of necrotic cell death | 2/101 | 23/18723 | 0.006770726 | 0.038684318 | 0.02838579 | PELI1/BIRC3 | 2 | BP |
| GO:0060575 | intestinal epithelial cell differentiation | 2/101 | 23/18723 | 0.006770726 | 0.038684318 | 0.02838579 | HIF1A/IL6ST | 2 | BP |
| GO:0071219 | cellular response to molecule of bacterial origin | 5/101 | 221/18723 | 0.006938497 | 0.039529606 | 0.029006046 | CCL5/CX3CR1/TNFSF4/CXCL13/HAVCR2 | 5 | BP |
| GO:0030183 | B cell differentiation | 4/101 | 141/18723 | 0.007161528 | 0.040684008 | 0.029853123 | ITM2A/VCAM1/RBPJ/CD27 | 4 | BP |
| GO:0002637 | regulation of immunoglobulin production | 3/101 | 73/18723 | 0.007211116 | 0.040849335 | 0.029974437 | TFRC/TNFSF4/CD28 | 3 | BP |
| GO:0036003 | positive regulation of transcription from RNA polymerase II promoter in response to stress | 2/101 | 24/18723 | 0.007360428 | 0.041342801 | 0.030336532 | HIF1A/RBPJ | 2 | BP |
| GO:0043302 | positive regulation of leukocyte degranulation | 2/101 | 24/18723 | 0.007360428 | 0.041342801 | 0.030336532 | FGR/KLRC2 | 2 | BP |
| GO:2000353 | positive regulation of endothelial cell apoptotic process | 2/101 | 24/18723 | 0.007360428 | 0.041342801 | 0.030336532 | CD160/RGCC | 2 | BP |
| GO:0002437 | inflammatory response to antigenic stimulus | 3/101 | 74/18723 | 0.007486777 | 0.041816901 | 0.030684418 | FGR/CD28/RBPJ | 3 | BP |
| GO:0003151 | outflow tract morphogenesis | 3/101 | 74/18723 | 0.007486777 | 0.041816901 | 0.030684418 | TGFBR3/HIF1A/RBPJ | 3 | BP |
| GO:0014065 | phosphatidylinositol 3-kinase signaling | 4/101 | 144/18723 | 0.007703089 | 0.042904915 | 0.031482781 | CCL5/CD160/FGR/CD28 | 4 | BP |
| GO:0034121 | regulation of toll-like receptor signaling pathway | 3/101 | 75/18723 | 0.007768606 | 0.043149307 | 0.031662111 | CD300A/PELI1/BIRC3 | 3 | BP |
| GO:0061351 | neural precursor cell proliferation | 4/101 | 145/18723 | 0.007889433 | 0.043698693 | 0.032065239 | CX3CR1/ADGRG1/TOX/HIF1A | 4 | BP |
| GO:0033622 | integrin activation | 2/101 | 25/18723 | 0.007972513 | 0.04391489 | 0.032223881 | PLEK/CXCL13 | 2 | BP |
| GO:0042832 | defense response to protozoan | 2/101 | 25/18723 | 0.007972513 | 0.04391489 | 0.032223881 | BATF/LYST | 2 | BP |
| GO:0045055 | regulated exocytosis | 5/101 | 230/18723 | 0.008171872 | 0.044889015 | 0.032938674 | PLEK/FGR/KLRC2/PTGDR/CD300A | 5 | BP |
| GO:0032418 | lysosome localization | 3/101 | 77/18723 | 0.008350894 | 0.045746381 | 0.033567793 | FGR/PTGDR/CD300A | 3 | BP |
| GO:0010212 | response to ionizing radiation | 4/101 | 148/18723 | 0.008466199 | 0.046250964 | 0.033938045 | ANXA1/NAMPT/TANK/VCAM1 | 4 | BP |
| GO:0001562 | response to protozoan | 2/101 | 26/18723 | 0.008606724 | 0.046383266 | 0.034035126 | BATF/LYST | 2 | BP |
| GO:0032515 | negative regulation of phosphoprotein phosphatase activity | 2/101 | 26/18723 | 0.008606724 | 0.046383266 | 0.034035126 | FKBP1A/LGALS3 | 2 | BP |
| GO:0046639 | negative regulation of alpha-beta T cell differentiation | 2/101 | 26/18723 | 0.008606724 | 0.046383266 | 0.034035126 | ANXA1/TNFSF4 | 2 | BP |
| GO:0060544 | regulation of necroptotic process | 2/101 | 26/18723 | 0.008606724 | 0.046383266 | 0.034035126 | PELI1/BIRC3 | 2 | BP |
| GO:1905523 | positive regulation of macrophage migration | 2/101 | 26/18723 | 0.008606724 | 0.046383266 | 0.034035126 | CCL5/CX3CR1 | 2 | BP |
| GO:0030500 | regulation of bone mineralization | 3/101 | 78/18723 | 0.008651412 | 0.046498424 | 0.034119627 | S1PR1/ADRB2/HIF1A | 3 | BP |
| GO:0014068 | positive regulation of phosphatidylinositol 3-kinase signaling | 3/101 | 79/18723 | 0.008958217 | 0.04801797 | 0.03523464 | CCL5/FGR/CD28 | 3 | BP |
| GO:0048738 | cardiac muscle tissue development | 5/101 | 236/18723 | 0.009074106 | 0.048379059 | 0.0354996 | HOPX/TGFBR3/S1PR1/FKBP1A/RBPJ | 5 | BP |
| GO:0048762 | mesenchymal cell differentiation | 5/101 | 236/18723 | 0.009074106 | 0.048379059 | 0.0354996 | TGFBR3/HIF1A/RGCC/TIAM1/RBPJ | 5 | BP |
| GO:0031334 | positive regulation of protein-containing complex assembly | 5/101 | 237/18723 | 0.009230902 | 0.049083783 | 0.036016712 | PLEK/SNX9/TFRC/LGALS3/CXCL13 | 5 | BP |
| GO:0001772 | immunological synapse | 5/103 | 44/19550 | 3.39547E-06 | 0.000482157 | 0.000396734 | GZMA/LGALS3/CD28/HLA-DRA/HAVCR2 | 5 | CC |
| GO:0045334 | clathrin-coated endocytic vesicle | 5/103 | 91/19550 | 0.000119542 | 0.008487473 | 0.006983762 | ADRB2/TFRC/HLA-DQA1/HLA-DRA/CTLA4 | 5 | CC |
| GO:0030669 | clathrin-coated endocytic vesicle membrane | 4/103 | 72/19550 | 0.000567875 | 0.02132241 | 0.017544756 | ADRB2/TFRC/HLA-DQA1/HLA-DRA | 4 | CC |
| GO:0030136 | clathrin-coated vesicle | 6/103 | 196/19550 | 0.000600631 | 0.02132241 | 0.017544756 | ADRB2/SNX9/TFRC/HLA-DQA1/HLA-DRA/CTLA4 | 6 | CC |
| GO:0023023 | MHC protein complex binding | 6/102 | 36/18368 | 4.30133E-08 | 9.54896E-06 | 7.78768E-06 | CD8A/KLRD1/CD160/KLRC2/HLA-DQA1/HLA-DRA | 6 | MF |

| | | | | | | | | | |
|---|---|---|---|---|---|---|---|---|---|
| GO:0140375 | immune receptor activity | 8/102 | 144/18368 | 1.38333E-06 | 0.00015355 | 0.000125228 | KLRD1/CD160/CX3CR1/KLRC2/HLA-DQA1/IL6ST/HLA-DRA/CXCR6 | 8 | MF |
| GO:0015026 | coreceptor activity | 5/102 | 48/18368 | 6.77713E-06 | 0.000501507 | 0.000409005 | CD8A/TGFBR3/GPR15/CD28/CXCR6 | 5 | MF |
| GO:0019955 | cytokine binding | 7/102 | 139/18368 | 1.23948E-05 | 0.000687912 | 0.000561028 | CX3CR1/A2M/TGFBR3/PLP2/IL6ST/CXCR6/TNFRSF9 | 7 | MF |
| GO:0005126 | cytokine receptor binding | 9/102 | 271/18368 | 2.00777E-05 | 0.00089145 | 0.000727025 | CCL5/CX3CR1/TGFBR3/XCL2/TNFSF10/FKBP1A/TNFSF4/IL6ST/CXCL13 | 9 | MF |
| GO:0019956 | chemokine binding | 4/102 | 33/18368 | 3.24045E-05 | 0.001198965 | 0.000977819 | CX3CR1/A2M/PLP2/CXCR6 | 4 | MF |
| GO:0042287 | MHC protein binding | 4/102 | 40/18368 | 7.02455E-05 | 0.002227785 | 0.001816876 | CD8A/KLRD1/FCRL6/CD244 | 4 | MF |
| GO:0019865 | immunoglobulin binding | 3/102 | 23/18368 | 0.00027159 | 0.007536622 | 0.00614651 | FCGR3A/FCGR3B/LGALS3 | 3 | MF |
| GO:0042379 | chemokine receptor binding | 4/102 | 72/18368 | 0.000690262 | 0.017026453 | 0.013885964 | CCL5/CX3CR1/XCL2/CXCL13 | 4 | MF |
| GO:0030246 | carbohydrate binding | 7/102 | 271/18368 | 0.000789146 | 0.017519043 | 0.014287697 | KLRD1/KLRG1/KLRC3/KLRC2/GALM/LGALS3/HLA-DRA | 7 | MF |
| GO:0019838 | growth factor binding | 5/102 | 141/18368 | 0.001138768 | 0.022794365 | 0.018589999 | FGFBP2/A2M/TGFBR3/IL6ST/CXCL13 | 5 | MF |
| GO:0004955 | prostaglandin receptor activity | 2/102 | 10/18368 | 0.001334805 | 0.022794365 | 0.018589999 | PTGER2/PTGDR | 2 | MF |
| GO:0032395 | MHC class II receptor activity | 2/102 | 10/18368 | 0.001334805 | 0.022794365 | 0.018589999 | HLA-DQA1/HLA-DRA | 2 | MF |
| GO:0004954 | prostanoid receptor activity | 2/102 | 11/18368 | 0.001625526 | 0.024057789 | 0.019620388 | PTGER2/PTGDR | 2 | MF |
| GO:0019864 | IgG binding | 2/102 | 11/18368 | 0.001625526 | 0.024057789 | 0.019620388 | FCGR3A/FCGR3B | 2 | MF |
| GO:0005125 | cytokine activity | 6/102 | 235/18368 | 0.001994094 | 0.027668049 | 0.022564744 | CCL5/XCL2/TNFSF10/NAMPT/TNFSF4/CXCL13 | 6 | MF |
| GO:0048020 | CCR chemokine receptor binding | 3/102 | 48/18368 | 0.002398737 | 0.031324681 | 0.025546919 | CCL5/XCL2/CXCL13 | 3 | MF |
| GO:0008009 | chemokine activity | 3/102 | 49/18368 | 0.002544954 | 0.031387769 | 0.02559837 | CCL5/XCL2/CXCL13 | 3 | MF |
| GO:0004953 | icosanoid receptor activity | 2/102 | 15/18368 | 0.003058659 | 0.032334394 | 0.026370393 | PTGER2/PTGDR | 2 | MF |
| GO:0045125 | bioactive lipid receptor activity | 2/102 | 15/18368 | 0.003058659 | 0.032334394 | 0.026370393 | S1PR5/S1PR1 | 2 | MF |
| GO:0048185 | activin binding | 2/102 | 15/18368 | 0.003058659 | 0.032334394 | 0.026370393 | TGFBR3/FKBP1A | 2 | MF |
| GO:0004869 | cysteine-type endopeptidase inhibitor activity | 3/102 | 56/18368 | 0.003723189 | 0.037570359 | 0.030640596 | CST7/BIRC3/CD27 | 3 | MF |
