## supplementary tables for "MGPfact^XMBD^: A Model-Based Factorization Method for scRNA Data Unveils Bifurcating Transcriptional Modules Underlying Cell Fate Determination": supplementary_table_11_ZNF683_seg4_t1_seg5_t2.pdf

**Supplementary Table 11: List of genes specifically expressed in CD8-ZNF683.**
**CD8-ZNF683-T1 cells (n = 204) vs. CD8-ZNF683-T2 cells (n = 395), two-sided moderated t-test with limma**

| Gene | logFC | AveExpr | t | P.Value | adj.P.Val | B |
|---|---|---|---|---|---|---|
| RGS1 | 1.102098067 | 0.975184147 | 3.055245056 | 0.002348263 | 0.012792992 | -2.111227395 |
| ATP2B1 | 1.003820358 | -0.11068835 | 3.346154662 | 0.000870428 | 0.005900309 | -1.208678331 |
| PHLDA1 | 0.686307242 | 0.510725831 | 2.858257113 | 0.004407116 | 0.02170987 | -2.6772241 |
| CTLA4 | 0.680995713 | -0.648740959 | 3.721565676 | 0.000216545 | 0.001831352 | 0.072473752 |
| HNRNPLL | 0.676642653 | -0.449823189 | 2.632590516 | 0.008691106 | 0.037464573 | -3.280485051 |
| TOX | 0.600989969 | 0.207470701 | 2.282262771 | 0.022821835 | 0.075058479 | -4.120827791 |
| SLA | 0.563040962 | 0.77430623 | 2.128074811 | 0.033736479 | 0.09799461 | -4.45338428 |
| U2AF1L4 | 0.531913448 | -0.096628794 | 2.176493955 | 0.029906309 | 0.092211119 | -4.351416898 |
| TNFRSF18 | 0.311448886 | -0.798021165 | 2.180092503 | 0.029637238 | 0.092182639 | -4.343748465 |
| LAYN | 0.301937347 | -0.257175755 | 2.941069684 | 0.003396489 | 0.017333804 | -2.44374363 |
| LOC606724 | -0.320647856 | 0.448975948 | -2.219713022 | 0.026810073 | 0.085026232 | -4.258493902 |
| EEF1A1 | -0.322459545 | -0.426172562 | -3.347544643 | 0.000866155 | 0.005900309 | -1.204175985 |
| RPL10 | -0.350068167 | -0.211721679 | -2.996282143 | 0.002845417 | 0.014984013 | -2.284480712 |
| RPL6 | -0.368913644 | -0.363475637 | -2.564054377 | 0.010587271 | 0.043324868 | -3.454112621 |
| RPL13 | -0.384008584 | -0.396957072 | -3.700582695 | 0.000234823 | 0.001948816 | -0.00257005 |
| CSF1 | -0.397245164 | 0.235493457 | -2.146886194 | 0.032201156 | 0.096376921 | -4.414037071 |
| ANAPC1P1 | -0.404950203 | 2.585123656 | -2.275416441 | 0.023231588 | 0.075296261 | -4.136079163 |
| RPL15 | -0.41000521 | -0.345501492 | -2.453062823 | 0.014447085 | 0.055060137 | -3.72578853 |
| RPS27 | -0.417453595 | -0.390793924 | -2.168778646 | 0.030490302 | 0.093363409 | -4.367816045 |
| PLEK | -0.430959834 | -1.339371747 | -2.978788575 | 0.003010459 | 0.015633259 | -2.335252869 |
| LINC02067 | -0.445303495 | -0.275318579 | -4.052759859 | 5.73E-05 | 0.000591186 | 1.310377639 |
| PFN1 | -0.456637937 | 0.361740338 | -5.691068703 | 1.97E-08 | 5.31E-07 | 8.870206096 |
| S1PR1 | -0.462888497 | -2.388199944 | -2.350726245 | 0.019058471 | 0.066615727 | -3.965838731 |
| RPS6 | -0.466900317 | -0.48775123 | -3.503456926 | 0.000493337 | 0.003744301 | -0.687746746 |
| RAD9A | -0.469366198 | 0.470775349 | -2.135410833 | 0.033130429 | 0.096775727 | -4.438080301 |
| SEM1 | -0.471557442 | 0.39201276 | -2.245481883 | 0.025099272 | 0.080173214 | -4.202234585 |
| RPL8 | -0.472222873 | -0.227424503 | -3.139902907 | 0.001772839 | 0.010316889 | -1.856756201 |
| CD96 | -0.476127357 | 2.189759297 | -2.138256718 | 0.03289785 | 0.096775727 | -4.432129422 |
| RPS7 | -0.48825005 | -0.23480328 | -3.1341769 | 0.001807229 | 0.010316889 | -1.874180101 |
| CMC1 | -0.496485763 | -0.125994362 | -2.270062972 | 0.023556441 | 0.075790287 | -4.147973562 |
| CDC42SE1 | -0.507100518 | -0.904140398 | -2.151948086 | 0.031798415 | 0.096376921 | -4.403391047 |
| SH3BGRL3 | -0.508513012 | 0.877085833 | -5.048325365 | 5.91E-07 | 1.05E-05 | 5.625285909 |
| RPSAP58 | -0.509587933 | -0.347715333 | -2.92838691 | 0.003536111 | 0.017841286 | -2.479921438 |
| TGFBR3 | -0.520112184 | -0.4527867 | -2.274300087 | 0.023299006 | 0.075296261 | -4.138561762 |
| APOBEC3F | -0.528029801 | 1.035152785 | -2.411277864 | 0.016195141 | 0.059187839 | -3.825014779 |
| TPT1 | -0.529332325 | -0.012749944 | -3.051929828 | 0.002373949 | 0.012854065 | -2.121055503 |
| CD8A | -0.537902112 | 6.324107241 | -4.525031331 | 7.28E-06 | 9.79E-05 | 3.24745671 |
| ID2 | -0.540120944 | 2.661005379 | -2.179391975 | 0.029689453 | 0.092182639 | -4.345242252 |
| TMEM256-PLSCR | -0.542186416 | -0.620780743 | -2.53723974 | 0.01142428 | 0.04632311 | -3.520825076 |
| HSPA1B | -0.546663522 | 0.444758517 | -2.525697917 | 0.011802331 | 0.046997624 | -3.549328854 |
| KIAA1671 | -0.548078846 | 0.762413651 | -2.318161141 | 0.020774768 | 0.070144463 | -4.040121269 |
| EEF1D | -0.551205328 | 0.002423798 | -2.363833217 | 0.018403565 | 0.064594332 | -3.935654041 |
| CEP78 | -0.554104264 | -0.301326987 | -3.13801044 | 0.001784138 | 0.010316889 | -1.862518267 |
| DRAP1 | -0.564728103 | 1.116799808 | -2.164513821 | 0.030817314 | 0.094040463 | -4.376856475 |
| OXNAD1 | -0.569489738 | 1.506520703 | -2.146376516 | 0.032241949 | 0.096376921 | -4.415107644 |
| FBXO10 | -0.569813984 | 0.216666416 | -2.275697439 | 0.023214645 | 0.075296261 | -4.135454077 |
| GAS5-AS1 | -0.574319283 | -0.063970496 | -4.068428436 | 5.36E-05 | 0.000560322 | 1.371419135 |
| GFI1 | -0.575425077 | 0.468513215 | -2.617335002 | 0.009084656 | 0.038784493 | -3.319520213 |
| FBXO32 | -0.575672772 | -0.264074721 | -2.390712649 | 0.017121991 | 0.062058483 | -3.873236948 |
| C1orf56 | -0.583107591 | -0.64775449 | -2.932764675 | 0.003487336 | 0.017695739 | -2.467450958 |
| CCL4L1 | -0.583817092 | 1.968920382 | -2.519409459 | 0.012012955 | 0.047622787 | -3.564805349 |

| | | | | | | |
|---|---|---|---|---|---|---|
| CLIC3 | -0.58527446 | 1.293767591 | -2.148955971 | 0.032035951 | 0.096376921 | -4.40968696 |
| ANAPC11 | -0.586148207 | 0.47118453 | -2.444118065 | 0.014806584 | 0.055855776 | -3.747170095 |
| XCL2 | -0.588735846 | 1.971543445 | -2.375116677 | 0.017855676 | 0.063423361 | -3.90953689 |
| POMP | -0.597968507 | 0.591200223 | -2.263081324 | 0.023986024 | 0.076893826 | -4.163444098 |
| RPL19 | -0.599594994 | -0.345523214 | -4.238980273 | 2.60E-05 | 0.000296523 | 2.050228392 |
| RAC2 | -0.601226604 | 0.624150443 | -3.795354418 | 0.00016235 | 0.001471086 | 0.339585902 |
| NACA | -0.603841628 | -0.07136324 | -2.532396876 | 0.011581576 | 0.046747451 | -3.532800538 |
| HDLBP | -0.60664236 | 0.498267865 | -2.136662206 | 0.033027988 | 0.096775727 | -4.435464582 |
| RPS3 | -0.60801639 | -0.33766131 | -5.053021543 | 5.78E-07 | 1.05E-05 | 5.647709764 |
| FIBP | -0.617815421 | 0.451921926 | -2.369163559 | 0.018142929 | 0.064186935 | -3.923331425 |
| HAPLN3 | -0.624494055 | -0.366766716 | -2.64298283 | 0.008431831 | 0.036524225 | -3.253766737 |
| EFHD2 | -0.627088123 | -0.25518663 | -2.607442336 | 0.00934831 | 0.039719137 | -3.344714662 |
| CD7 | -0.629798226 | 1.734268216 | -3.09665436 | 0.002048227 | 0.011584875 | -1.987597319 |
| RBX1 | -0.631817806 | 0.214360298 | -2.664364349 | 0.007920091 | 0.034587526 | -3.198472202 |
| EEF1B2 | -0.638103024 | -0.464757071 | -2.66326174 | 0.007945783 | 0.034587526 | -3.201334287 |
| ADGRE5 | -0.641708711 | 1.892592405 | -2.34455659 | 0.019373754 | 0.066858209 | -3.979990112 |
| RPL39 | -0.642315818 | -0.326124942 | -4.630630203 | 4.47E-06 | 6.11E-05 | 3.707869053 |
| OAS1 | -0.643487638 | -0.11512497 | -2.44318458 | 0.014844553 | 0.055855776 | -3.749397088 |
| RPS25 | -0.644083125 | -0.298659463 | -4.490858722 | 8.50E-06 | 0.000112719 | 3.100585596 |
| ACTN4 | -0.645078974 | 1.431670485 | -2.145120021 | 0.032342706 | 0.096376921 | -4.417745826 |
| SRRT | -0.645339175 | 0.703494791 | -2.322453291 | 0.02054109 | 0.069926737 | -4.030388886 |
| EIF5A | -0.64799698 | 0.429442061 | -2.481174799 | 0.013366969 | 0.051608123 | -3.658091442 |
| R3HDM1 | -0.648814937 | 0.092694787 | -2.785375592 | 0.00551488 | 0.02563986 | -2.877337029 |
| SLC1A5 | -0.651730737 | 0.858568044 | -2.273984439 | 0.023318099 | 0.075296261 | -4.139263498 |
| RPL12 | -0.653057215 | -0.249355225 | -3.756779221 | 0.000188846 | 0.001628112 | 0.19932161 |
| EPSTI1 | -0.654066146 | 0.159979935 | -2.367149103 | 0.018241046 | 0.064277971 | -3.927991619 |
| ACP5 | -0.654364186 | 0.520283094 | -2.470231699 | 0.013778613 | 0.052738831 | -3.684533596 |
| APOBEC3G | -0.65625425 | 2.680645619 | -2.297884971 | 0.021910342 | 0.073029339 | -4.085858105 |
| PTPN22 | -0.658301241 | 2.603375321 | -2.127691407 | 0.033768413 | 0.09799461 | -4.45418269 |
| TOMM5 | -0.659417818 | 0.185791696 | -2.574052065 | 0.010289473 | 0.042696504 | -3.429063695 |
| ITGAE | -0.660600253 | 3.586034762 | -2.124416693 | 0.034042222 | 0.098147704 | -4.460996251 |
| PDE4D | -0.660764949 | 0.949702864 | -2.349328231 | 0.019129516 | 0.066615727 | -3.969048564 |
| SUSD3 | -0.66226851 | 0.999554882 | -2.176632368 | 0.029895921 | 0.092211119 | -4.351122174 |
| ITGAL | -0.662698475 | 0.116256205 | -2.145886314 | 0.032281226 | 0.096376921 | -4.41613707 |
| MYO1F | -0.662900853 | 0.487180187 | -2.125469463 | 0.033953989 | 0.098147704 | -4.458806923 |
| PGK1 | -0.663607811 | 1.83298509 | -2.856947092 | 0.004425097 | 0.02170987 | -2.680865455 |
| RAB1B | -0.665068566 | 0.943286188 | -2.438554563 | 0.015034152 | 0.056330495 | -3.760430486 |
| IRF7 | -0.665988916 | 0.604905734 | -2.322237363 | 0.020552791 | 0.069926737 | -4.030878922 |
| DEF6 | -0.669538009 | 0.623950915 | -2.143108928 | 0.032504535 | 0.096535208 | -4.421965225 |
| DMAC1 | -0.670283379 | -0.095256277 | -3.072886915 | 0.002215818 | 0.012221405 | -2.058753617 |
| RNF31 | -0.670311323 | 0.252587667 | -2.135600907 | 0.033114851 | 0.096775727 | -4.43768309 |
| RPL22 | -0.681640341 | -0.263257616 | -3.292757457 | 0.001050222 | 0.006748382 | -1.380275481 |
| LARS | -0.6817383 | 0.094608445 | -2.405990531 | 0.016429113 | 0.0597912 | -3.837451419 |
| RPSA | -0.682128201 | -0.271752067 | -3.521018151 | 0.000462408 | 0.003539816 | -0.628164489 |
| RPL13A | -0.68323901 | -0.73272838 | -4.048836242 | 5.82E-05 | 0.000594008 | 1.295126934 |
| PTGER2 | -0.687497023 | -0.024013311 | -2.433812356 | 0.015230559 | 0.056826624 | -3.771709972 |
| COX8A | -0.6882912 | 0.411103277 | -3.168890577 | 0.001607816 | 0.009639176 | -1.768076489 |
| PRDX2 | -0.688381558 | -0.340574894 | -2.315700944 | 0.020909753 | 0.070332805 | -4.045691757 |
| RPS3A | -0.69413786 | -0.336014312 | -4.285138334 | 2.13E-05 | 0.000248407 | 2.238453766 |
| RIN3 | -0.696042361 | 0.70905968 | -3.136266067 | 0.001794611 | 0.010316889 | -1.867826454 |
| DDX60 | -0.700459914 | 0.81107094 | -2.202266196 | 0.028024885 | 0.088248575 | -4.296221729 |
| GBP1 | -0.701168763 | 0.61766608 | -2.186014514 | 0.029198979 | 0.091298217 | -4.331101638 |
| TNFSF10 | -0.701325273 | 0.661530436 | -2.304889129 | 0.021512095 | 0.072085812 | -4.070103485 |
| RPS15 | -0.705615817 | -0.219109996 | -5.112784714 | 4.27E-07 | 8.82E-06 | 5.934742643 |

| | | | | | | |
|---|---|---|---|---|---|---|
| GPANK1 | -0.705893883 | 0.011911139 | -2.967682868 | 0.00311973 | 0.016106512 | -2.367335597 |
| RASSF1 | -0.706689013 | 0.390536873 | -2.413525067 | 0.016096592 | 0.059187839 | -3.819720896 |
| PSMD8 | -0.707796036 | 0.372833237 | -2.225697815 | 0.026403989 | 0.084038502 | -4.24548466 |
| APOBEC3C | -0.709548064 | 1.586722728 | -2.425189813 | 0.015593486 | 0.057695899 | -3.792163807 |
| RPL17 | -0.710696191 | -0.293827496 | -3.81231924 | 0.000151843 | 0.001404552 | 0.401704269 |
| KHDRBS1 | -0.711290494 | 0.152642563 | -2.493890428 | 0.012902365 | 0.050031878 | -3.627222485 |
| SNRPB | -0.714031591 | 0.48942477 | -2.339450614 | 0.019638126 | 0.067330719 | -3.991674115 |
| GZMK | -0.71519632 | 0.632988906 | -2.196038616 | 0.028469882 | 0.089333058 | -4.309617611 |
| PLAC8 | -0.715878626 | -1.903585127 | -3.362112431 | 0.000822527 | 0.005662044 | -1.156880558 |
| CD63 | -0.719509468 | 2.759521401 | -2.503252831 | 0.012569494 | 0.0493881 | -3.604395187 |
| RPL10A | -0.720709444 | -0.125287672 | -3.756907895 | 0.000188752 | 0.001628112 | 0.199787214 |
| LY6E | -0.72215723 | 0.879630378 | -2.686962198 | 0.007409713 | 0.03273545 | -3.139559377 |
| PDIA6 | -0.723180383 | 1.02007375 | -2.215383046 | 0.027107234 | 0.085662719 | -4.267884562 |
| HM13 | -0.724919235 | 0.402767688 | -2.318379667 | 0.020762815 | 0.070144463 | -4.039626192 |
| RPL38 | -0.726406663 | -0.302494602 | -4.753443416 | 2.50E-06 | 3.77E-05 | 4.255747869 |
| RPS23 | -0.730744867 | -0.36190466 | -4.381662064 | 1.39E-05 | 0.000171369 | 2.638248175 |
| PTP4A2 | -0.734231824 | 0.348642263 | -2.381471088 | 0.017553475 | 0.062852766 | -3.894775006 |
| COX4I1 | -0.734290877 | -0.074209326 | -2.941991329 | 0.003386541 | 0.017333804 | -2.441108706 |
| RAB27A | -0.734510936 | 1.404793375 | -2.387324482 | 0.017279088 | 0.062373293 | -3.881142792 |
| ST13 | -0.735417058 | 0.023881391 | -2.622378113 | 0.008952828 | 0.038406333 | -3.306640641 |
| WDR1 | -0.737156432 | 0.653552936 | -2.503576505 | 0.012558124 | 0.0493881 | -3.60360451 |
| UBE2D2 | -0.737547732 | 0.322845783 | -2.577913424 | 0.010176472 | 0.042625975 | -3.41936365 |
| GZMH | -0.739583063 | 2.913105118 | -2.297051696 | 0.021958146 | 0.073029339 | -4.087729275 |
| AOAH | -0.740016061 | 2.860759439 | -2.274705318 | 0.023274514 | 0.075296261 | -4.13766073 |
| TMEM123 | -0.74021041 | -0.083593502 | -2.151792543 | 0.031810726 | 0.096376921 | -4.403718547 |
| IFI44L | -0.742411642 | 0.674273833 | -2.379968094 | 0.017624545 | 0.0628538 | -3.898270094 |
| RPL37A | -0.743087647 | -0.278717431 | -4.641397752 | 4.25E-06 | 5.93E-05 | 3.755371045 |
| SYNC | -0.744514268 | 0.007117056 | -2.825296046 | 0.004880318 | 0.02355284 | -2.768349442 |
| C12orf75 | -0.745108801 | 1.580363561 | -2.2845525 | 0.022686206 | 0.074889781 | -4.115716978 |
| NAP1L1 | -0.746897301 | 0.012502312 | -2.38405864 | 0.017431711 | 0.06266947 | -3.888752796 |
| CMTM6 | -0.747999869 | 0.008189234 | -2.286080554 | 0.022596085 | 0.074870611 | -4.112303471 |
| DARS | -0.748092698 | -0.02870682 | -2.411243051 | 0.016196672 | 0.059187839 | -3.825096751 |
| LYAR | -0.748172903 | 0.160568847 | -2.772833628 | 0.005729131 | 0.026497231 | -2.911266208 |
| COPB1 | -0.748694431 | 0.440988562 | -2.343561713 | 0.01942502 | 0.066858209 | -3.98226865 |
| RPS11 | -0.748915312 | -0.252633743 | -5.101135119 | 4.53E-07 | 8.94E-06 | 5.878548928 |
| RPS10-NUDT3 | -0.752328698 | -0.304957477 | -4.643792433 | 4.20E-06 | 5.93E-05 | 3.765949335 |
| IKZF3 | -0.754316179 | 1.249824155 | -2.807093605 | 0.005160958 | 0.02450765 | -2.818231585 |
| EPHA1 | -0.755015858 | 1.204273307 | -2.726927781 | 0.006579009 | 0.029457573 | -3.034180211 |
| TCIRG1 | -0.759742162 | 0.558203041 | -2.529855109 | 0.0116649 | 0.04687073 | -3.539076874 |
| RPS4X | -0.760624517 | -0.279609319 | -4.140639463 | 3.96E-05 | 0.000423489 | 1.655614284 |
| IDH2 | -0.761684554 | 1.6460772 | -2.347267856 | 0.019234644 | 0.066720172 | -3.973775758 |
| CKLF-CMTM1 | -0.768509435 | 1.921374875 | -3.611709719 | 0.000329561 | 0.002612948 | -0.315920277 |
| PLTP | -0.768510357 | 1.578760143 | -3.076921455 | 0.002186509 | 0.012135127 | -2.046712203 |
| S100A10 | -0.773695774 | 0.854929712 | -3.426080333 | 0.000654073 | 0.004722088 | -0.946865605 |
| RPL23A | -0.774021157 | -0.284849274 | -5.113553786 | 4.25E-07 | 8.82E-06 | 5.938456516 |
| TRNT1 | -0.774115318 | -0.335255642 | -2.904953028 | 0.00380794 | 0.019104243 | -2.546367288 |
| RPL18 | -0.774213316 | -0.466695884 | -4.408844146 | 1.23E-05 | 0.000153942 | 2.752340993 |
| SIT1 | -0.77530757 | 0.52184561 | -2.477118012 | 0.013518284 | 0.051966389 | -3.667907337 |
| RPLP0 | -0.777128518 | -0.367885572 | -3.857190622 | 0.000127062 | 0.001187699 | 0.567276158 |
| SAMHD1 | -0.77889842 | -1.089608087 | -2.495103671 | 0.012858793 | 0.050031878 | -3.624269094 |
| NDUFB11 | -0.779200536 | 0.193112213 | -3.331231021 | 0.00091756 | 0.006080549 | -1.256904655 |
| GNPTAB | -0.779227029 | 0.941571985 | -2.579591104 | 0.010127721 | 0.042622826 | -3.415144753 |
| RPS21 | -0.780753686 | -0.337484014 | -5.048566897 | 5.91E-07 | 1.05E-05 | 5.626438739 |
| DOK2 | -0.786131199 | 0.037328102 | -2.566499203 | 0.010513746 | 0.043223179 | -3.447995984 |

| | | | | | | |
|---|---|---|---|---|---|---|
| TOMM7 | -0.786862208 | -0.207846346 | -3.933982031 | 9.33E-05 | 0.000920414 | 0.854902097 |
| ANAPC16 | -0.787269481 | 0.215985372 | -2.743192993 | 0.006265683 | 0.028680034 | -2.99085853 |
| ARHGAP9 | -0.788771606 | 0.762344901 | -2.58548885 | 0.009957993 | 0.042108083 | -3.400292265 |
| SNRK | -0.789446716 | -0.129915161 | -2.995600779 | 0.002851687 | 0.014984013 | -2.286463656 |
| JUN | -0.792332332 | 2.420941568 | -2.427838579 | 0.015481195 | 0.057520088 | -3.78588815 |
| PTPN7 | -0.792743207 | 1.64024332 | -2.542164874 | 0.011266266 | 0.045891945 | -3.508623218 |
| ATP6V1C2 | -0.793982877 | 0.793564338 | -2.74140397 | 0.006299474 | 0.028684891 | -2.995635808 |
| LINC00861 | -0.795299468 | -1.2189289 | -2.785496793 | 0.005512845 | 0.02563986 | -2.877008422 |
| RPS24 | -0.795728874 | -0.219917232 | -4.640064745 | 4.28E-06 | 5.93E-05 | 3.749484801 |
| RPS16 | -0.797402171 | -0.218204832 | -4.92552365 | 1.09E-06 | 1.82E-05 | 5.045721938 |
| RPS14P3 | -0.800340809 | -0.401342545 | -6.147021364 | 1.44E-09 | 5.11E-08 | 11.38269744 |
| IRF9 | -0.801151882 | 0.200961442 | -2.574088238 | 0.010288409 | 0.042696504 | -3.428972892 |
| C7orf25 | -0.812014565 | 0.212367145 | -2.663458314 | 0.007941197 | 0.034587526 | -3.200824116 |
| C9orf16 | -0.812174981 | 0.097692494 | -3.279697491 | 0.001099157 | 0.006971799 | -1.42184 |
| CKLF | -0.812369367 | 1.921172657 | -3.778367376 | 0.000173554 | 0.001556725 | 0.277650612 |
| PELO | -0.815700441 | 2.63564694 | -2.82573428 | 0.004873736 | 0.02355284 | -2.767144632 |
| UBA52 | -0.817660831 | -0.192618659 | -5.059893273 | 5.58E-07 | 1.05E-05 | 5.68055616 |
| MT1A | -0.822253223 | 0.541678958 | -4.069646062 | 5.34E-05 | 0.000560322 | 1.37617208 |
| RPS29 | -0.823373041 | -0.330813388 | -6.219538955 | 9.33E-10 | 3.60E-08 | 11.79805062 |
| ARF6 | -0.826793626 | 1.32605041 | -2.802424351 | 0.005235265 | 0.024633384 | -2.830976658 |
| RPS14 | -0.82735321 | -0.496022094 | -6.75456808 | 3.38E-11 | 2.00E-09 | 14.99319188 |
| SURF4 | -0.827600238 | 1.132008761 | -2.445645977 | 0.014744621 | 0.055855776 | -3.743523192 |
| CSNK2B | -0.829899316 | 0.068590231 | -2.711067149 | 0.006898073 | 0.030627442 | -3.076182345 |
| ATP6V1G1 | -0.830932735 | -0.069069731 | -2.731542532 | 0.006488707 | 0.029248588 | -3.02191455 |
| SRGAP3 | -0.833428192 | 1.005477851 | -3.310192415 | 0.000988041 | 0.006451326 | -1.324538985 |
| CLSTN3 | -0.840929505 | 0.696858086 | -3.427293905 | 0.000651213 | 0.004722088 | -0.942844506 |
| SUMO2 | -0.841248747 | 0.282140749 | -3.379208094 | 0.000773937 | 0.005454413 | -1.101126104 |
| CDK2AP2 | -0.843379791 | 0.176414393 | -2.752551482 | 0.00609157 | 0.028027535 | -2.965818771 |
| SKIL | -0.846078398 | 1.260448009 | -2.739724297 | 0.00633135 | 0.028684891 | -3.000118321 |
| CYBA | -0.847929299 | 0.757439893 | -4.020594378 | 6.54E-05 | 0.000660208 | 1.185766142 |
| CAPG | -0.847985368 | 3.572769707 | -2.526160327 | 0.011786974 | 0.046997624 | -3.548189327 |
| HINT1 | -0.851504287 | 0.091331476 | -3.291095761 | 0.001056335 | 0.006748382 | -1.385572824 |
| CXCR3 | -0.852174158 | 2.938304978 | -2.725791552 | 0.006601415 | 0.029457573 | -3.037197118 |
| ITGA1 | -0.856031359 | 4.161391781 | -2.570368802 | 0.010398304 | 0.042947414 | -3.438303098 |
| RPS27A | -0.860286235 | -0.265238664 | -6.477263141 | 1.94E-10 | 9.08E-09 | 13.30861972 |
| C4orf3 | -0.862775555 | 0.432047937 | -3.079844395 | 0.002165499 | 0.012094107 | -2.03797889 |
| LY6G5B | -0.864009817 | 0.045620039 | -2.810109895 | 0.005113466 | 0.02450765 | -2.809987415 |
| ISG15 | -0.87590092 | 0.703486457 | -3.25906503 | 0.001180792 | 0.007384105 | -1.487180005 |
| CCL4 | -0.880320784 | 3.215608325 | -2.868752813 | 0.004265428 | 0.021160338 | -2.64799147 |
| TPI1 | -0.885711709 | 1.126168425 | -3.094400844 | 0.002063602 | 0.011597963 | -1.994366801 |
| MT2A | -0.886847678 | 0.577856341 | -4.149203503 | 3.82E-05 | 0.00041336 | 1.689632199 |
| SNORA33 | -0.888140268 | -0.271118478 | -5.861614788 | 7.56E-09 | 2.24E-07 | 9.789838941 |
| RPS20 | -0.888864841 | -0.487913074 | -5.002699149 | 7.43E-07 | 1.29E-05 | 5.408421047 |
| RPS15A | -0.890408873 | -0.250225363 | -5.511178549 | 5.29E-08 | 1.24E-06 | 7.926603903 |
| RPL31 | -0.89076277 | -0.47120165 | -6.201536427 | 1.04E-09 | 3.85E-08 | 11.69453977 |
| RPL11 | -0.891897801 | -0.410653603 | -5.618993818 | 2.94E-08 | 7.46E-07 | 8.488870704 |
| ARPC4 | -0.892760848 | 0.278488055 | -3.336469366 | 0.000900753 | 0.00601405 | -1.240000389 |
| FLNA | -0.89578452 | 0.040910289 | -2.984888142 | 0.002951944 | 0.015419566 | -2.317582652 |
| RNASEK-C17orf4 | -0.897429127 | 0.681450994 | -3.894692614 | 0.000109335 | 0.001043969 | 0.707068726 |
| LDHA | -0.898813575 | 1.253002854 | -3.022270411 | 0.002615482 | 0.013991254 | -2.208521693 |
| RPL27A | -0.899659512 | -0.347597332 | -6.496723588 | 1.72E-10 | 8.49E-09 | 13.42484663 |
| PSMA2 | -0.90067579 | 0.259483952 | -2.891426056 | 0.003973385 | 0.019822282 | -2.584486282 |
| RPL27 | -0.903626805 | -0.135372704 | -5.422749105 | 8.51E-08 | 1.89E-06 | 7.472777807 |
| PATL2 | -0.905608967 | -0.209760419 | -3.443261335 | 0.00061465 | 0.004577449 | -0.889809908 |

| | | | | | | |
|---|---|---|---|---|---|---|
| RPS10 | -0.906046575 | -0.377315377 | -5.967809035 | 4.11E-09 | 1.26E-07 | 10.37467596 |
| ACTR3 | -0.9113939 | 0.928631506 | -3.145876362 | 0.001737607 | 0.010233892 | -1.838546543 |
| ZYX | -0.918025988 | 2.430329984 | -3.067210859 | 0.002257662 | 0.012375334 | -2.075668359 |
| SELENOH | -0.919476814 | 0.321540288 | -3.539397314 | 0.000431988 | 0.003348919 | -0.565501173 |
| CISH | -0.924844008 | 1.945434276 | -2.494767193 | 0.012870864 | 0.050031878 | -3.625088323 |
| GPI | -0.92926968 | 0.87352267 | -2.808676157 | 0.005135991 | 0.02450765 | -2.813907203 |
| KLRC2 | -0.930703635 | 1.951900825 | -3.440047018 | 0.000621853 | 0.004577449 | -0.900505058 |
| GNG5 | -0.934215405 | 0.265238084 | -3.93005222 | 9.48E-05 | 0.000924925 | 0.840052084 |
| MIR497HG | -0.937957532 | 0.486248734 | -3.313025169 | 0.000978267 | 0.006434821 | -1.315456379 |
| BLOC1S1-RDH5 | -0.938901155 | 0.60371902 | -3.951254548 | 8.70E-05 | 0.000867641 | 0.920338654 |
| IFI6 | -0.943388622 | 0.532086701 | -3.218602279 | 0.001357425 | 0.008256117 | -1.614164171 |
| KLRC3 | -0.944442873 | 1.91669786 | -3.246512914 | 0.001233166 | 0.007657704 | -1.526736095 |
| RPS12 | -0.950801087 | -0.27795787 | -6.258179411 | 7.40E-10 | 2.99E-08 | 12.02111357 |
| IVNS1ABP | -0.951586323 | 1.577470923 | -2.828704083 | 0.004829339 | 0.02355284 | -2.758975139 |
| NCL | -0.953231174 | -0.020717908 | -3.220003468 | 0.00135092 | 0.008256117 | -1.609792421 |
| RPS18 | -0.95596358 | -0.44729534 | -7.03555503 | 5.42E-12 | 4.01E-10 | 16.76165059 |
| LASP1 | -0.95650558 | 0.660837484 | -3.012722334 | 0.002697905 | 0.014345744 | -2.236502934 |
| EWSR1 | -0.956981813 | 0.335962169 | -3.047500821 | 0.002408666 | 0.012963004 | -2.134169312 |
| RPLP1 | -0.956994458 | -0.085997249 | -7.970701953 | 7.95E-15 | 3.53E-12 | 23.07794499 |
| RPS19 | -0.963404662 | -0.256349509 | -7.727168949 | 4.62E-14 | 7.92E-12 | 21.37075339 |
| SENP3-EIF4A1 | -0.96545936 | 0.104742413 | -3.207446377 | 0.001410254 | 0.008519083 | -1.64890534 |
| RPL36 | -0.974452463 | -0.355347471 | -6.603576711 | 8.83E-11 | 4.61E-09 | 14.06839783 |
| ATP5L | -0.97460158 | 0.318060445 | -4.755518499 | 2.48E-06 | 3.77E-05 | 4.265119216 |
| RPL30 | -0.979605398 | -0.384862621 | -6.723700308 | 4.12E-11 | 2.29E-09 | 14.80267233 |
| NDFIP2 | -0.982484984 | 0.532382477 | -3.729031889 | 0.000210371 | 0.001796241 | 0.099273687 |
| RPS13 | -0.992865268 | -0.492875918 | -6.015907424 | 3.11E-09 | 1.02E-07 | 10.64262837 |
| PFDN5 | -0.994086319 | -0.349220268 | -4.474650702 | 9.15E-06 | 0.000117801 | 3.031288447 |
| RBBP4 | -0.996181388 | 0.061968795 | -3.167136499 | 0.001617384 | 0.009639176 | -1.773464999 |
| PXN | -0.996869716 | 0.057703178 | -3.368869964 | 0.000802991 | 0.005570748 | -1.134874595 |
| RAP1B | -1.001550249 | 0.415489805 | -4.194111648 | 3.15E-05 | 0.000349831 | 1.869100478 |
| GAS5 | -1.002645691 | -0.034050413 | -3.624886573 | 0.000313547 | 0.002508376 | -0.269920939 |
| COX6C | -1.004100173 | 0.249773479 | -4.99514846 | 7.71E-07 | 1.32E-05 | 5.372706475 |
| NDUFB3 | -1.013090053 | 0.212963799 | -4.767340441 | 2.34E-06 | 3.72E-05 | 4.318580953 |
| TSPAN14 | -1.01836234 | 0.830307652 | -3.374665142 | 0.000786582 | 0.005499878 | -1.11596868 |
| RPL28 | -1.023124502 | -0.082307046 | -5.853899455 | 7.90E-09 | 2.26E-07 | 9.747712674 |
| CTSC | -1.02332917 | 0.605051925 | -3.219253752 | 0.001354397 | 0.008256117 | -1.612131786 |
| RPL32 | -1.023705395 | -0.34184361 | -6.908448169 | 1.25E-11 | 8.54E-10 | 15.95407718 |
| RPL34 | -1.02584737 | -0.489725072 | -7.38222129 | 5.20E-13 | 5.78E-11 | 19.02705388 |
| PARP9 | -1.035854822 | 0.316368016 | -3.26383881 | 0.001161419 | 0.00731447 | -1.472097506 |
| RPLP2 | -1.040664598 | -0.326975752 | -7.745389142 | 4.06E-14 | 7.92E-12 | 21.49698776 |
| TARP | -1.045083005 | 2.247995034 | -3.133320182 | 0.001812427 | 0.010316889 | -1.876784399 |
| F2R | -1.046440926 | 1.012410126 | -3.297805617 | 0.001031855 | 0.006688227 | -1.364166584 |
| CD52 | -1.046442427 | 0.623485832 | -8.079602477 | 3.57E-15 | 3.17E-12 | 23.85516817 |
| RPL35A | -1.057560557 | -0.277949782 | -6.32680655 | 4.89E-10 | 2.17E-08 | 12.42026294 |
| GIMAP1-GIMAP5 | -1.064387862 | -0.548591869 | -3.411669979 | 0.000688943 | 0.004894252 | -0.99450883 |
| RPL23 | -1.076158215 | -0.166865901 | -6.831164187 | 2.07E-11 | 1.31E-09 | 15.46917718 |
| LAP3 | -1.080222825 | 0.521152598 | -3.772173225 | 0.000177818 | 0.001579022 | 0.255132408 |
| ARL6IP1 | -1.087379152 | -0.018935297 | -3.422008389 | 0.000663756 | 0.004753348 | -0.960347751 |
| GZMA | -1.097759941 | 2.849517433 | -3.630329824 | 0.000307147 | 0.002479517 | -0.250872215 |
| YARS | -1.100820324 | 0.829638434 | -3.439215442 | 0.000623729 | 0.004577449 | -0.903270438 |
| FAU | -1.10429526 | -0.192983073 | -7.662403462 | 7.33E-14 | 9.30E-12 | 20.92400844 |
| IL2RB | -1.107167139 | 1.706993214 | -3.580633182 | 0.000370418 | 0.002910899 | -0.423770962 |
| BLOC1S1 | -1.109213547 | 0.658241241 | -4.479753418 | 8.94E-06 | 0.000116801 | 3.053079783 |
| RPL24 | -1.110323299 | -0.380084137 | -6.084279186 | 2.08E-09 | 7.11E-08 | 11.02679449 |

| | | | | | | |
|---|---|---|---|---|---|---|
| GIMAP5 | -1.12192543 | -0.691952593 | -3.341153902 | 0.000885965 | 0.00596013 | -1.224861608 |
| PABPC1 | -1.143496408 | -0.588303184 | -4.75830587 | 2.45E-06 | 3.77E-05 | 4.277713317 |
| RPS9 | -1.151369435 | -0.424956737 | -5.473748038 | 6.47E-08 | 1.47E-06 | 7.733698145 |
| OAS2 | -1.156161236 | 0.181246859 | -3.637465899 | 0.000298943 | 0.002435427 | -0.22585802 |
| RPL35 | -1.161555651 | -0.291045132 | -7.09102875 | 3.75E-12 | 3.33E-10 | 17.11800587 |
| SYNGR2 | -1.179205077 | 0.284178849 | -3.76558477 | 0.000182461 | 0.001604215 | 0.231219405 |
| CYTH4 | -1.181144859 | 0.616547488 | -3.704265326 | 0.000231514 | 0.001939475 | 0.010571252 |
| ABI3 | -1.183849832 | 2.051130457 | -3.666296378 | 0.000267846 | 0.002202291 | -0.124319973 |
| CTSW | -1.186963633 | 3.430412744 | -4.299563252 | 2.00E-05 | 0.000236339 | 2.297669305 |
| CTSA | -1.187531461 | 2.350829438 | -3.486142664 | 0.000525715 | 0.003956228 | -0.746211256 |
| RPL37 | -1.189278128 | -0.435980849 | -7.706679595 | 5.35E-14 | 7.92E-12 | 21.2290877 |
| ITM2C | -1.194374364 | 2.568674481 | -3.538331196 | 0.0004337 | 0.003348919 | -0.569144619 |
| PPP1CC | -1.195038448 | 0.669777534 | -3.799839443 | 0.000159507 | 0.001460231 | 0.355982607 |
| RPL29 | -1.196323158 | -0.39039994 | -5.805653994 | 1.04E-08 | 2.88E-07 | 9.485410491 |
| CCL5 | -1.199009288 | 4.376940116 | -7.841823111 | 2.03E-14 | 6.01E-12 | 22.16913734 |
| ATP5B | -1.20994843 | 0.419340569 | -4.328434058 | 1.76E-05 | 0.000211022 | 2.416747833 |
| RPL26 | -1.21807537 | -0.102159878 | -7.1453313 | 2.61E-12 | 2.57E-10 | 17.46912217 |
| SLC9A3R1 | -1.248115692 | 0.785015985 | -4.198035934 | 3.10E-05 | 0.000348349 | 1.88486978 |
| SLAMF7 | -1.26913226 | 1.398325675 | -3.924091884 | 9.71E-05 | 0.000937234 | 0.817555988 |
| B3GNTL1 | -1.286861363 | 1.647885806 | -5.232205395 | 2.32E-07 | 5.02E-06 | 6.517541448 |
| GZMB | -1.298750094 | 4.091821009 | -4.183257605 | 3.30E-05 | 0.000361943 | 1.825557106 |
| PSME2 | -1.308673675 | 0.94444695 | -5.090977046 | 4.77E-07 | 9.21E-06 | 5.829645628 |
| STAT3 | -1.330845815 | 0.491239289 | -4.238401199 | 2.60E-05 | 0.000296523 | 2.047879199 |
| PSMA1 | -1.375003338 | 0.732060965 | -4.695781432 | 3.29E-06 | 4.87E-05 | 3.99685572 |
| RPS8 | -1.378654532 | -0.724334484 | -6.289132886 | 6.14E-10 | 2.60E-08 | 12.20067392 |
| FASLG | -1.414897522 | 1.988083008 | -4.365103525 | 1.50E-05 | 0.000181919 | 2.569070258 |
| SAMD3 | -1.430947055 | 0.642661083 | -4.664827863 | 3.81E-06 | 5.54E-05 | 3.859089415 |
| GLUL | -1.486766507 | 2.754598261 | -4.416788945 | 1.19E-05 | 0.000150687 | 2.785812923 |
| ABRACL | -1.495453009 | 0.74717677 | -5.105401963 | 4.43E-07 | 8.94E-06 | 5.899117121 |
| SPN | -1.512336068 | 0.740102579 | -4.840038508 | 1.65E-06 | 2.72E-05 | 4.650038806 |
| NKG7 | -1.616918928 | 3.682129314 | -7.067568678 | 4.38E-12 | 3.54E-10 | 16.96701336 |
| CALR | -1.629480956 | 0.82234196 | -5.607403441 | 3.13E-08 | 7.73E-07 | 8.427955562 |
| KLRD1 | -1.743037756 | 2.742874342 | -4.832085404 | 1.72E-06 | 2.77E-05 | 4.613551679 |
| TBCD | -1.783359322 | 2.612362132 | -5.577307822 | 3.69E-08 | 8.86E-07 | 8.27031137 |
| PRF1 | -1.831068111 | 2.17321645 | -5.97383699 | 3.97E-09 | 1.26E-07 | 10.40815279 |
| GNLY | -2.15211353 | 1.412724078 | -5.643718507 | 2.56E-08 | 6.70E-07 | 8.619193023 |
