## supplementary tables for "MGPfact^XMBD^: A Model-Based Factorization Method for scRNA Data Unveils Bifurcating Transcriptional Modules Underlying Cell Fate Determination": supplementary_table_12_GZMK_seg5_t1_seg6_t2_BH.pdf

**Supplementary Table 12: List of genes specifically expressed in CD8-GZMK.**
**CD8-GZMK-T1 cells (n = 155) vs. CD8-GZMK-T2 cells (n = 221), two-sided moderated t-test with limma**

| Gene | logFC | AveExpr | t | P.Value | adj.P.Val | B |
|---|---|---|---|---|---|---|
| GZMB | 3.064255203 | 1.463824435 | 8.157775663 | 5.15E-15 | 3.61E-12 | 23.47790692 |
| GZMH | 3.045848722 | 1.769473459 | 7.917729732 | 2.74E-14 | 9.59E-12 | 21.87051672 |
| HLA-DRA | 2.286718664 | 1.597606853 | 5.643591339 | 3.28E-08 | 4.59E-06 | 8.478352178 |
| BHLHE40 | 2.029695679 | 0.789764309 | 5.479849867 | 7.81E-08 | 7.73E-06 | 7.65530549 |
| HSPH1 | 1.956779687 | 0.966465148 | 5.013370783 | 8.25E-07 | 5.25E-05 | 5.424541449 |
| C12orf75 | 1.883882346 | 0.39784373 | 5.048388588 | 6.95E-07 | 4.87E-05 | 5.586043086 |
| ITGB1 | 1.877480555 | -0.682836097 | 5.456159153 | 8.83E-08 | 7.73E-06 | 7.537923878 |
| DUSP4 | 1.617222268 | 0.619065825 | 4.190037537 | 3.48E-05 | 0.000869544 | 1.915719215 |
| SMC4 | 1.605598843 | -0.241977709 | 4.689507791 | 3.84E-06 | 0.000191798 | 3.977715384 |
| ZEB2 | 1.594165841 | 1.946306382 | 4.068659275 | 5.76E-05 | 0.001222399 | 1.446370233 |
| PGK1 | 1.580202197 | -0.146768515 | 4.9649188 | 1.04E-06 | 6.09E-05 | 5.202699057 |
| HSPA1A | 1.552642873 | 0.686070584 | 4.124466908 | 4.58E-05 | 0.001019768 | 1.660607234 |
| PRF1 | 1.546885414 | 1.650882068 | 4.136910139 | 4.34E-05 | 0.001013812 | 1.708737738 |
| LDHA | 1.492464044 | 0.853987801 | 4.644346467 | 4.72E-06 | 0.000194405 | 3.782746729 |
| IFNG | 1.490052969 | 1.623446953 | 3.798456266 | 0.000169695 | 0.002474717 | 0.447014831 |
| HSPA1L | 1.485695117 | 0.425110903 | 4.385847862 | 1.50E-05 | 0.000500303 | 2.699232831 |
| ANXA1 | 1.455170938 | 2.232492758 | 3.766629208 | 0.000191942 | 0.002742034 | 0.333470066 |
| SLA2 | 1.452035521 | 0.928371345 | 3.852293564 | 0.000137508 | 0.002169758 | 0.641091399 |
| HLA-DQB1 | 1.436724019 | 1.52917212 | 4.398385671 | 1.42E-05 | 0.000497444 | 2.750501355 |
| CAPG | 1.422554621 | -0.495704026 | 4.375271585 | 1.57E-05 | 0.000500303 | 2.656087718 |
| APOBEC3C | 1.422391141 | 0.543674898 | 3.965986658 | 8.75E-05 | 0.001655266 | 1.059203654 |
| CTSC | 1.406359289 | -0.473650929 | 3.615032731 | 0.000341136 | 0.0042642 | -0.195187158 |
| IL6ST | 1.356489254 | -1.321466034 | 3.920529054 | 0.000104964 | 0.001836878 | 0.890688722 |
| BHLHE40-AS1 | 1.356366537 | 0.059333998 | 5.544854315 | 5.55E-08 | 6.47E-06 | 7.979599925 |
| HLA-DQA1 | 1.3353184 | 0.570275153 | 3.998066535 | 7.69E-05 | 0.00153727 | 1.179199333 |
| RGCC | 1.294918616 | 1.268635425 | 3.42106864 | 0.000692123 | 0.00654711 | -0.842033871 |
| TNFAIP3 | 1.284745089 | 1.492533644 | 4.558940418 | 6.96E-06 | 0.000270699 | 3.418627739 |
| HLA-DMA | 1.282595014 | 0.547811218 | 3.567857423 | 0.000406379 | 0.004733731 | -0.355576582 |
| DDIT4 | 1.25915988 | 0.929603147 | 3.16568983 | 0.001673141 | 0.01270741 | -1.642511031 |
| CLEC2B | 1.238198403 | 0.798387942 | 3.48240763 | 0.000555284 | 0.005716156 | -0.641082893 |
| HLA-DRB6 | 1.235917554 | 2.076549075 | 4.297821624 | 2.20E-05 | 0.000669237 | 2.343001384 |
| TANK | 1.234218208 | -0.218156579 | 3.13711838 | 0.001840419 | 0.013560984 | -1.72841779 |
| LOC100130476 | 1.224478304 | 0.468809342 | 3.387177462 | 0.000780632 | 0.007285903 | -0.951626642 |
| CHST12 | 1.219998258 | 0.42381607 | 3.563796551 | 0.000412511 | 0.004733731 | -0.369291121 |
| JUN | 1.205112062 | 2.014382818 | 3.046679267 | 0.002476916 | 0.017513551 | -1.995471018 |
| ICOS | 1.191832911 | -0.942110202 | 2.983883498 | 0.003031703 | 0.020211355 | -2.176527673 |
| JMJD6 | 1.187578637 | 1.52874252 | 2.966279894 | 0.003206485 | 0.020977002 | -2.226638934 |
| IRF4 | 1.177705459 | 0.049550764 | 3.676804321 | 0.000270504 | 0.003572694 | 0.017792127 |
| HMGB2 | 1.176028961 | 0.47069036 | 3.32133546 | 0.000983484 | 0.008605487 | -1.16160496 |
| PFKFB3 | 1.1736649 | 0.1598721 | 3.436529394 | 0.000654938 | 0.00645714 | -0.791698888 |
| PTTG1 | 1.171658901 | -0.22134956 | 4.193946098 | 3.42E-05 | 0.000869544 | 1.931041595 |
| HLA-DRB5 | 1.169996171 | 2.115586473 | 4.119984002 | 4.66E-05 | 0.001019768 | 1.643299704 |
| MCL1 | 1.164845135 | 1.044424461 | 3.378258554 | 0.000805619 | 0.007420178 | -0.980297065 |
| NR3C1 | 1.146573961 | -0.436733814 | 2.991829993 | 0.0029557 | 0.019894135 | -2.153814256 |
| HLA-DRB1 | 1.141924333 | 2.409244054 | 3.871444056 | 0.000127521 | 0.002075921 | 0.710734056 |
| HSPA1B | 1.125775421 | 0.912199629 | 3.232537409 | 0.001335129 | 0.010620342 | -1.438639274 |
| OASL | 1.117438357 | 1.363124844 | 2.884546618 | 0.00414512 | 0.025661571 | -2.455597658 |
| JAKMIP1 | 1.116340039 | 0.205761821 | 3.756539381 | 0.000199551 | 0.002756568 | 0.297658661 |
| TNFSF10 | 1.105561811 | -0.487352388 | 3.935231508 | 9.90E-05 | 0.001776575 | 0.944996555 |
| CKS2 | 1.10541405 | 0.488947462 | 3.667967688 | 0.000279687 | 0.003625574 | -0.012880926 |
| PDE4B | 1.086770881 | 1.463000662 | 3.168306852 | 0.001658544 | 0.01270741 | -1.634605461 |
| LGALS3 | 1.0749792 | -0.210883973 | 3.256104788 | 0.001231896 | 0.010027062 | -1.36580373 |

| | | | | | | |
|---|---|---|---|---|---|---|
| SAMSN1 | 1.060310064 | 0.294091308 | 3.119161163 | 0.001953311 | 0.014242896 | -1.782032538 |
| CD6 | 1.046408265 | 0.429244798 | 2.81649107 | 0.005110728 | 0.029566197 | -2.641574333 |
| KLRD1 | 1.013049444 | 0.684919679 | 2.810862462 | 0.005199088 | 0.029588304 | -2.65676541 |
| HSP90AB1 | 1.006140626 | 0.788263218 | 3.574098536 | 0.000397121 | 0.004733731 | -0.334470438 |
| LEPROTL1 | 0.998699082 | 0.274276334 | 3.000542454 | 0.00287438 | 0.019726135 | -2.128845363 |
| CD55 | 0.986870533 | 0.27586005 | 2.654860495 | 0.008270755 | 0.042652585 | -3.066188737 |
| CLIC1 | 0.981826055 | 0.385099173 | 3.162999805 | 0.00168827 | 0.01270741 | -1.650630679 |
| TYMP | 0.977040042 | -0.266268553 | 3.327531614 | 0.000962487 | 0.008528367 | -1.142009992 |
| IQGAP1 | 0.96797086 | 0.194196092 | 2.611080842 | 0.009386257 | 0.046931284 | -3.177047523 |
| HSPD1 | 0.940108165 | 0.96872061 | 2.379006506 | 0.017857021 | 0.075300691 | -3.734955386 |
| CAMK4 | 0.933589878 | -0.066243096 | 2.531164713 | 0.011774635 | 0.055506173 | -3.374826813 |
| GNLY | 0.932635198 | 0.379481584 | 2.663997927 | 0.008053535 | 0.042253005 | -3.042827133 |
| RBPJ | 0.927735207 | -0.341548256 | 2.58783999 | 0.010031612 | 0.048764782 | -3.235176221 |
| SLC2A3 | 0.922933453 | 1.146465175 | 2.551263739 | 0.011127829 | 0.053352606 | -3.325643375 |
| PMAIP1 | 0.917451017 | 0.734125688 | 2.315453085 | 0.021124829 | 0.086450029 | -3.878966964 |
| ARID5B | 0.898952334 | -0.199490716 | 2.305651348 | 0.021672819 | 0.087189501 | -3.900840627 |
| SLA | 0.893373669 | 0.690822486 | 2.457821346 | 0.01442798 | 0.064328575 | -3.551112601 |
| DNAJA1 | 0.885455218 | 1.526265937 | 2.599511764 | 0.009702709 | 0.047830255 | -3.206046061 |
| TNFRSF1B | 0.88490335 | -0.219675264 | 2.284736694 | 0.022883786 | 0.089026226 | -3.947212536 |
| TPI1 | 0.871413713 | -0.37142242 | 2.419559387 | 0.016012199 | 0.068764045 | -3.641086961 |
| SLAMF1 | 0.855701365 | -1.38334135 | 2.778554906 | 0.005733608 | 0.031853379 | -2.743397091 |
| HSP90AA1 | 0.855585286 | 0.804704168 | 3.627369238 | 0.000325775 | 0.004146226 | -0.152921048 |
| SERINC5 | 0.849271042 | -1.085136579 | 2.607193962 | 0.009491529 | 0.047121067 | -3.186804048 |
| HAVCR2 | 0.849184972 | -0.46088317 | 3.813229903 | 0.000160217 | 0.002438084 | 0.500020739 |
| GZMA | 0.847108027 | 3.240779406 | 2.901272114 | 0.003934789 | 0.024813984 | -2.409241765 |
| PELO | 0.83238065 | 0.375584183 | 2.510181693 | 0.012485405 | 0.057681094 | -3.425772535 |
| PTPN22 | 0.831880803 | 0.893712873 | 2.288428669 | 0.022665824 | 0.089026226 | -3.939056548 |
| LINC-PINT | 0.830505437 | 0.882438505 | 2.574744967 | 0.010412459 | 0.050267042 | -3.267708016 |
| S100A4 | 0.830470672 | -1.78504197 | 2.762760117 | 0.006012628 | 0.033140468 | -2.785400812 |
| CTLA4 | 0.829557765 | -1.61511814 | 3.524141768 | 0.000477123 | 0.005138253 | -0.502448187 |
| SLC7A5 | 0.796743881 | 1.387796558 | 2.284590442 | 0.022892458 | 0.089026226 | -3.947535358 |
| MTHFD2 | 0.793436185 | 0.131911895 | 2.220391591 | 0.026986796 | 0.099951096 | -4.087300915 |
| HLA-DQB2 | 0.776537986 | 0.557274589 | 2.995527385 | 0.002920935 | 0.019851016 | -2.143226409 |
| EMP3 | 0.762853252 | -0.230983145 | 2.613328932 | 0.009325848 | 0.046931284 | -3.171398159 |
| IFI6 | 0.759347497 | -0.036111488 | 2.452147951 | 0.014653756 | 0.064921702 | -3.564540082 |
| ACP5 | 0.758173171 | -0.758779846 | 3.114242293 | 0.00198533 | 0.014327125 | -1.796667849 |
| VCAM1 | 0.741431698 | 0.253289452 | 3.515991224 | 0.000491528 | 0.005166992 | -0.529644464 |
| ADAM19 | 0.734596116 | -1.172372768 | 2.223957754 | 0.026743739 | 0.099577751 | -4.079638805 |
| SLC27A2 | 0.721825073 | 0.118542759 | 3.31664339 | 0.000999667 | 0.008639096 | -1.176420464 |
| AHR | 0.691983231 | -0.46263813 | 2.29605865 | 0.022221108 | 0.088379408 | -3.922160505 |
| ARL3 | 0.653856618 | -0.354332162 | 3.426860163 | 0.000677969 | 0.006539336 | -0.823203569 |
| CD38 | 0.645888474 | -0.099306888 | 2.738441413 | 0.00646629 | 0.035351644 | -2.849623285 |
| PERP | 0.632241612 | -0.059681767 | 2.249019098 | 0.025088576 | 0.094929746 | -4.025455503 |
| HLA-DPA1 | 0.585331529 | 1.92839567 | 2.404514133 | 0.016676026 | 0.071178162 | -3.676091866 |
| MIR4435-2HG | 0.581289159 | -0.507690933 | 3.347446113 | 0.000897792 | 0.008161741 | -1.078798791 |
| GFPT2 | 0.547149562 | 0.183884248 | 2.535939683 | 0.011618012 | 0.055323866 | -3.363176211 |
| CD44 | 0.491607333 | 1.296681441 | 2.427115681 | 0.015687684 | 0.06778629 | -3.623426389 |
| GAPDH | 0.4524342 | -0.097439681 | 2.447373722 | 0.014846165 | 0.064951972 | -3.575816193 |
| LAIR2 | 0.29827726 | -0.762675971 | 2.811502964 | 0.005188964 | 0.029588304 | -2.655038226 |
| SCML1 | -0.291584254 | -0.124096982 | -2.465798207 | 0.014115748 | 0.063339897 | -3.532182617 |
| PTCH1 | -0.303157695 | -0.116010112 | -2.300676147 | 0.021955692 | 0.08782277 | -3.911908848 |
| NT5E | -0.309603559 | -0.083199376 | -2.631150582 | 0.00885915 | 0.044937717 | -3.126447312 |
| KLRF1 | -0.312532082 | -0.149080626 | -2.499571145 | 0.012859143 | 0.058832682 | -3.451378406 |
| RPL37 | -0.328268176 | -0.331278807 | -2.311121094 | 0.021365507 | 0.086450029 | -3.888645397 |
| RPS6 | -0.359088192 | -0.285278091 | -2.509043237 | 0.012525038 | 0.057681094 | -3.428524931 |
| RPL3 | -0.37290412 | -0.178185497 | -2.735932864 | 0.006514803 | 0.035351644 | -2.856217019 |

| | | | | | | |
|---|---|---|---|---|---|---|
| RPLP1 | -0.383004312 | -0.225866531 | -3.268700282 | 0.001179808 | 0.009831732 | -1.326672266 |
| LINC02067 | -0.39973542 | 0.124227911 | -3.106706254 | 0.002035324 | 0.01453803 | -1.819047623 |
| RPS3 | -0.402411161 | 0.082224141 | -3.958501079 | 9.02E-05 | 0.001660956 | 1.031331296 |
| RPS14P3 | -0.405636582 | -0.096651912 | -3.232753573 | 0.001334147 | 0.010620342 | -1.437973484 |
| RPL37A | -0.421871714 | -0.262767382 | -2.524558517 | 0.011994424 | 0.055973977 | -3.390910512 |
| RPS14 | -0.424762688 | -0.076620335 | -3.807339611 | 0.000163934 | 0.002441574 | 0.478864334 |
| RPL27A | -0.426388804 | 0.143691853 | -3.341538131 | 0.00091655 | 0.008225444 | -1.097588524 |
| SPINT2 | -0.430228333 | -0.555921125 | -2.654194626 | 0.008286788 | 0.042652585 | -3.067888141 |
| EEF1A1 | -0.434904342 | -0.037527357 | -3.457328862 | 0.000607851 | 0.006166604 | -0.723647166 |
| RPL13A | -0.462593491 | 0.166034035 | -2.875257977 | 0.004266304 | 0.025661571 | -2.48123115 |
| ID3 | -0.481104815 | -0.448051438 | -2.529946741 | 0.011814885 | 0.055506173 | -3.377795185 |
| RPL34 | -0.503456135 | -0.358793584 | -3.566146812 | 0.000408951 | 0.004733731 | -0.361355503 |
| RPL32 | -0.505017142 | -0.424743903 | -3.271100206 | 0.001170118 | 0.009831732 | -1.319200055 |
| RPLP2 | -0.525752921 | -0.086962416 | -4.277746557 | 2.40E-05 | 0.000699039 | 2.262674032 |
| UBASH3B | -0.527594318 | -0.579228191 | -2.588169445 | 0.010022193 | 0.048764782 | -3.234355707 |
| RPL30 | -0.530890783 | -0.099277029 | -3.883920614 | 0.000121388 | 0.002023133 | 0.756277513 |
| RPL19 | -0.536130806 | -0.333009452 | -3.193304325 | 0.001524918 | 0.011860471 | -1.558780844 |
| RASGRP2 | -0.551968886 | -0.701538261 | -2.276812623 | 0.023357778 | 0.089837608 | -3.964674475 |
| RPL36 | -0.560417305 | -0.405506104 | -3.283765561 | 0.001120191 | 0.009562607 | -1.279680533 |
| SNORA33 | -0.566835329 | -0.03408419 | -3.538827783 | 0.000452162 | 0.005079029 | -0.453296178 |
| RPL13 | -0.569144169 | -0.129046811 | -5.398431072 | 1.19E-07 | 9.27E-06 | 7.253707783 |
| RPS2 | -0.569353104 | 0.124066878 | -2.960679199 | 0.003263996 | 0.021155531 | -2.242522872 |
| A2MP1 | -0.585659039 | 0.328865335 | -2.855943524 | 0.004528677 | 0.026865031 | -2.534279361 |
| FAM117B | -0.599287814 | 0.202832427 | -2.31916953 | 0.020920244 | 0.086142179 | -3.870649739 |
| RPS12 | -0.60021615 | -0.048495265 | -3.749822696 | 0.000204774 | 0.002756568 | 0.273868729 |
| RPS29 | -0.616125493 | -0.009358306 | -4.258800097 | 2.60E-05 | 0.000713074 | 2.187174926 |
| RPL10 | -0.633382909 | 0.062254629 | -4.145273159 | 4.20E-05 | 0.001012831 | 1.741160175 |
| RPS18 | -0.64109019 | -0.201583556 | -4.661464228 | 4.36E-06 | 0.000194405 | 3.856449639 |
| IKZF2 | -0.641335925 | -0.883409039 | -2.827072929 | 0.004948292 | 0.028865035 | -2.612936111 |
| RPS16 | -0.642684088 | -0.196218529 | -3.437212606 | 0.000653339 | 0.00645714 | -0.78946967 |
| PDCD4 | -0.645268729 | 0.994410928 | -2.372796309 | 0.018155456 | 0.076100713 | -3.749194416 |
| EEF1G | -0.66015868 | -0.557852687 | -2.797095666 | 0.005421072 | 0.030602824 | -2.693797992 |
| ABCB1 | -0.68704011 | 0.241807352 | -2.339877817 | 0.019811723 | 0.082548848 | -3.824068452 |
| LINC00612 | -0.692153834 | 1.07113983 | -2.227471718 | 0.026506098 | 0.099220689 | -4.07207714 |
| LDLRAP1 | -0.714662654 | -0.549383166 | -2.269762685 | 0.023786652 | 0.090831131 | -3.980160543 |
| CCL3L1 | -0.759589924 | 1.561258159 | -2.673757544 | 0.0078272 | 0.041507878 | -3.017789501 |
| TOX | -0.760427745 | -0.211341491 | -2.27747181 | 0.023318024 | 0.089837608 | -3.963224102 |
| CCL3L3 | -0.787741119 | 1.647691987 | -2.709743697 | 0.007041259 | 0.037625048 | -2.924708785 |
| NR4A2 | -0.810679064 | 2.601734296 | -2.480841243 | 0.013543183 | 0.061250973 | -3.496322352 |
| FAM53C | -0.824451802 | 0.822740975 | -2.268314222 | 0.023875612 | 0.090831131 | -3.98333649 |
| LGR6 | -0.829207228 | 0.485421214 | -3.533402722 | 0.000461237 | 0.005079029 | -0.471475332 |
| EEF1B2 | -0.830907241 | -0.478101392 | -2.874632519 | 0.004274579 | 0.025661571 | -2.482954362 |
| LINC00861 | -0.851213122 | -0.34100291 | -2.480318864 | 0.013562715 | 0.061250973 | -3.497571159 |
| BBIP1 | -0.861297145 | 0.571645609 | -2.662515121 | 0.008088432 | 0.042253005 | -3.046623463 |
| RIPOR2 | -0.904176219 | -0.150391737 | -2.449314769 | 0.01476767 | 0.064951972 | -3.571234256 |
| ATM | -0.914416712 | 0.753010716 | -2.312809652 | 0.02127141 | 0.086450029 | -3.884874955 |
| PTGDR | -0.933534358 | 0.303616671 | -3.531553827 | 0.000464368 | 0.005079029 | -0.477664949 |
| IL7R | -0.951444839 | -0.332192712 | -2.237597058 | 0.025831562 | 0.097215555 | -4.050223543 |
| RAB37 | -0.954878689 | 0.002515779 | -2.88722115 | 0.004110813 | 0.025661571 | -2.448202205 |
| PLAC8 | -0.961236562 | -0.707603615 | -3.202350752 | 0.001479065 | 0.011633093 | -1.531201464 |
| CCND3 | -0.962107651 | -0.658486234 | -2.43682284 | 0.01527933 | 0.066431869 | -3.600660558 |
| TNFRSF9 | -0.980583889 | 0.317897573 | -2.87353395 | 0.004289148 | 0.025661571 | -2.485980184 |
| FOSB | -0.981628997 | 2.490912749 | -2.849987147 | 0.004612502 | 0.027132362 | -2.550569879 |
| CLDND1 | -0.982862051 | 1.311871686 | -2.394501653 | 0.017131133 | 0.072677533 | -3.69927005 |
| GIMAP1-GIMAP5 | -0.994267407 | -0.054624732 | -2.649704257 | 0.008395641 | 0.042897437 | -3.079337559 |
| LTB | -1.017106717 | -2.030540996 | -2.949786256 | 0.003378562 | 0.021499942 | -2.273333997 |

| | | | | | | |
|---|---|---|---|---|---|---|
| PATJ | -1.03421608 | 0.233866642 | -2.954960854 | 0.003323688 | 0.021344785 | -2.258710943 |
| GIMAP5 | -1.102743321 | -0.134137026 | -2.787567371 | 0.005579705 | 0.031246348 | -2.719326997 |
| CD28 | -1.104628062 | -0.180367227 | -2.877033693 | 0.004242891 | 0.025661571 | -2.476336888 |
| GIMAP7 | -1.120011717 | -0.292312057 | -3.042793546 | 0.002508334 | 0.01755834 | -2.006778684 |
| DGKA | -1.152990596 | -1.465900866 | -2.968727651 | 0.003181642 | 0.020977002 | -2.219687947 |
| LIMD2 | -1.180950089 | -0.541566759 | -3.425216069 | 0.000681959 | 0.006539336 | -0.828552143 |
| TCF7 | -1.196780944 | -0.205223236 | -3.750399143 | 0.00020432 | 0.002756568 | 0.275908909 |
| TRAT1 | -1.29005052 | 0.891645272 | -3.14015794 | 0.001821917 | 0.013560984 | -1.71931372 |
| TXNIP | -1.293903245 | -1.136246652 | -3.848659433 | 0.000139484 | 0.002169758 | 0.62791148 |
| CCR7 | -1.294724148 | 0.225620844 | -3.011138092 | 0.002778247 | 0.01925518 | -2.098386351 |
| CD27-AS1 | -1.35951252 | 0.004550657 | -3.514306872 | 0.000494555 | 0.005166992 | -0.535257374 |
| CRTAM | -1.375790917 | 2.85517877 | -3.264876315 | 0.001195402 | 0.009844485 | -1.338567572 |
| CMC1 | -1.415333078 | 1.282728897 | -4.25435919 | 2.65E-05 | 0.000713074 | 2.169522422 |
| GZMK | -1.474190269 | 6.076355183 | -4.481495119 | 9.85E-06 | 0.00036281 | 3.09366943 |
| PZP | -1.519438666 | 1.35327647 | -4.648893992 | 4.62E-06 | 0.000194405 | 3.802303226 |
| CD27 | -1.571196205 | 0.019905171 | -3.90179342 | 0.000113087 | 0.001930758 | 0.821754139 |
| EOMES | -1.577500169 | 1.979477122 | -4.009983181 | 7.32E-05 | 0.001507818 | 1.223999625 |
| KLRG1 | -1.694735623 | 2.298785498 | -3.980725029 | 8.24E-05 | 0.001603134 | 1.114222828 |
| FCMR | -1.790832811 | -0.559057465 | -4.840192129 | 1.90E-06 | 0.000102092 | 4.640312076 |
| FCRL3 | -2.055573419 | 0.767114253 | -6.045777533 | 3.58E-09 | 8.36E-07 | 10.58553378 |
| A2M | -2.126694261 | 1.828769934 | -5.725825074 | 2.11E-08 | 3.69E-06 | 8.899381188 |
